# Neurological Research in 47 Asian Countries

**DOI:** 10.1101/2024.04.06.588406

**Authors:** Waseem Hassan, Mehreen Zafar

## Abstract

Neurology research in the Asian region has seen significant growth and development over the past decade, yet comprehensive analyses of its progress remain scarce. This study aims to address this gap by providing a detailed examination of neurology research in 47 Asian countries from 2013 to 2022. Data was extracted from the Scopus database, encompassing scholarly output, citations, and citation impact metrics. Findings reveal a total of 128,306 research documents published with 1,725,678 total citations, demonstrating a commendable level of research activity. However, disparities exist among countries, with China, Japan, and Australia emerging as leading contributors in terms of both quantity and citation impact. Despite this, a substantial number of countries and universities have published fewer than 100 papers, indicating room for improvement and the need for targeted capacity-building efforts.

## 1.0 Introduction

In a recent study details about “Neurosurgical research in Southeast Asia” is provided (1). The authors meticulously provided descriptive insights into this domain and conducted a thorough bibliometric analysis (1). Their focus centered on Eleven (n=11) Southeast Asian (SEA) countries, namely Brunei, Cambodia, Indonesia, Laos, Malaysia, Myanmar, Philippines, Singapore, Thailand, Timor* OR East Timor, and Vietnam. Employing various bibliometric indicators such as the number of publications, citations, and average citations per publication, they meticulously presented the research productivity of each country. They presented the most productive universities and sources (where the papers were published) with only number of publications. Further the quality of journals was described with its impact factor.

In the present study, we tried to provide a holistic overview of neurological research across all 47 Asian countries from 2013 to 2022. By expanding the scope beyond conventional metrics and incorporating a broader array of indicators, we tried to offer a nuanced understanding of research progress and trends in this dynamic field. Through this endeavor, we aim to fill a crucial gap in the literature and contribute valuable insights to the discourse on neurological research in Asia.

## 2.0 Materials and Methods

Data for this study were obtained (On July 01, 2023) from the Scopus database, a comprehensive bibliographic database covering a wide range of academic disciplines, including neurology. Scopus provides access to scholarly publications, citation metrics, and other bibliometric indicators, making it a valuable resource for analyzing research trends and productivity. We included all scholarly documents categorized under the field of neurology published between the years 2013 and 2022. This encompassed research articles, reviews, conference papers, and other document types related to neurological sciences. Key metrics such as total scholarly output, total citations, citations per paper, and Field-Weighted Citation Impact (FWCI) were used to assess research productivity and impact.

## 3.0 Results and Discussion

The entire Asian region has published 128306 research documents or scholarly output (SO) with 1725678 total citations (TC), 13.4 citations per paper (CPP) and 0.95 Field-Weighted Citation Impact (FWCI). The per year scholarly output, TC, CPP and FWCI is presented in supplementary table 1. A marked increase in publications can be observed in the stated supplementary table 1. While a decrease in number of citations was noted for the last years. The data for each individual country with SO, growth rate (GR), TC, CPP, and FWCI is presented in supplementary table 2. The highest documents with highest citations were recorded for China (SO= 42471, TC= 522913), followed by Japan (SO= 29797, TC= 368137), and Australia (SO=17949, TC= 487042). Only 13 countries have published at least 500 papers in neurology. The highest CPP and FWCI was noted for Hong Kong (CPP= 29, FWCI=2.05), New Zealand (CPP=28.5, FWCI=1.89), Australia (CPP=27.1, FWCI=1.69) and Singapore (CPP=25.5, FWCI=1.69). While, 26 countries/regions have published less than 100 paper in neurology. This is alarming, and concrete measures are needed to improve the overall research culture and productivity.

We also presented the SO, GR, TC, CPP, and FWCI of three thousand (n=3000) universities/organization involved in neurology research. The data is presented in supplementary table 3. For instance, the highest documents are published by Capital Medical University (from China, n=5974), University of Melbourne (from Australia, n=4624), and University of Sydney (from Australia, n=3684). The highest TC was noted for three Australian Universities i.e. University of Melbourne (TC= 154108), University of Sydney (n= 123102) and University of New South Wales (n= 84917). Total 41 universities (14 from China, 8 from Australia, 7 from Japan, 6 from South Korea, 3 from India, 2 from Taiwan and 1 from Singapore) have published at least 1000 research paper in this field. While, 65 universities published 500-1000 papers, while 307 universities published 100-500 papers.

In order to acknowledge the research performance of authors, we also presented some relevant descriptive data i.e. SO, TC, CPP, FWCI and H-Index for the top 500 scientists (supplementary table 4). Total 224 researchers have published at least 100 papers. In the last decade (from 2013 to 2022), the highest documents are published by Tominaga, Teiji from Tohoku University School of Medicine, Sendai, Japan (n=401), followed by Wang, Yongjun from Beijing Tiantan Hospital, Capital Medical University, Beijing, China (n=387) and Hattori, Nobutaka from Juntendo University School of Medicine, Tokyo, Japan (n=320). Interestingly the highest TC was recorded for Scheffer, Ingrid E. from University of Melbourne, Parkville, Australia (n= 18328), Feigin, Valery L. from Auckland University of Technology, Auckland, New Zealand (n= 15920) and Rowe, Christopher C. from University of Melbourne, Parkville, Australia (n= 15271). While, the highest h-index was noted for Masters, Colin L. from The Florey Institute of Neuroscience and Mental Health, Melbourne, Australia (n=135), Hodges, John R. from The University of Sydney, Sydney, Australia (n=133) and Berkovic, Samuel F. from University of Melbourne, Parkville, Australia (n=117).

The quality of research can be decoded from the quality of journals. For the purpose, we tried to distribute all publications (n=128306) in different quartile groups. 3565 papers are published in those journals which do not have citescore data or are not included in top 100% journals. The entire Asian region published 29.98% papers (n= 37407) in Q1 (top 25%) research journals, while 31.06%, 26.62% and 12.31 % documents are published in Q2 (top 26% - 50%), Q3 (top 51% - 75%), Q4 (top 76% - 100%), respectively. This shows that 61% papers are published in top 50 % journals. The per year (from 2013 to 2022) publications in Q1-Q4 journals are presented in supplementary table 5. Furthermore, the highest documents are published in World Neurosurgery (n= 6439), followed by Frontiers in Neurology (n=5011), and Neurology India (n= 3462). In the present study the TC, CPP, Source-Normalized Impact per Paper (SNIP), CiteScore (2022) and SCImago Journal Rank (SJR) for 100 journals are presented in supplementary table 6.

Based on the data provided, several key conclusions can be drawn regarding the state of neurology research in the Asian region.

1. The Asian region has demonstrated significant research output in neurology, with a total of 128,306 scholarly documents published over the past decade. This reflects a substantial commitment to advancing knowledge in the field.
2. While the total number of citations is substantial at 1,725,678, the average citations per paper (CPP) and Field-Weighted Citation Impact (FWCI) suggest room for improvement in citation impact and influence of the research produced.
3. There are notable disparities in research productivity among Asian countries, with China, Japan, and Australia emerging as the top contributors in terms of both quantity and citation impact. However, a significant number of countries and universities have published fewer than 100 papers, indicating a need for broader participation and capacity building.
4. Several universities, particularly those from China, Australia, Japan, and South Korea, have made substantial contributions to neurology research. However, there is also a wide variation in research output among universities, emphasizing the need for targeted support and collaboration to enhance research capacity across institutions.
5. The top-performing authors, as indicated by the number of publications, citations, and H-index, hail from diverse institutions across Asia, underscoring the global nature of neurology research collaboration and excellence.
6. While a significant proportion of papers are published in high-quality journals (Q1 and Q2 quartiles), there is still a considerable portion published in lower-ranking journals. This highlights the importance of ensuring research is disseminated in reputable outlets to maximize visibility and impact.

While the Asian region has made significant strides in neurology research, there are opportunities for further growth and enhancement, particularly in improving citation impact, addressing regional disparities, fostering collaboration among institutions, and prioritizing publication in high-quality journals. Our data (on neurology research in the Asian region) can help in several ways:

1. Government agencies and funding bodies can utilize this data to inform their policies and funding allocations in the field of neurology research. Understanding the strengths and weaknesses of different countries and institutions can help in directing resources where they are most needed and where they can have the greatest impact.
2. Researchers and institutions can use this data to identify potential collaborators and partners across the Asian region. By understanding which universities and countries are leading in neurology research, researchers can initiate collaborations to share resources, expertise, and data, ultimately accelerating the pace of scientific discovery.
3. Analysis of the data can reveal areas of neurology research that are underrepresented or underfunded in certain regions or institutions. This information can guide stakeholders in identifying research gaps and setting research priorities to address pressing issues in neurology.
4. Institutions and researchers can use metrics such as citations per paper and Field-Weighted Citation Impact to assess the impact and influence of their research outputs. This can be valuable for performance evaluations, grant applications, and tenure decisions, providing a quantitative measure of research quality and significance.
5. Institutions and countries can benchmark their neurology research output and impact against regional and global peers. By identifying best practices and successful strategies employed by leading institutions, countries can learn from each other and implement initiatives to improve their own research productivity and impact.

Overall, the data on neurology research in the Asian region serves as a valuable resource for various stakeholders involved in scientific research, policy-making, funding allocation, and collaboration efforts. By leveraging this data effectively, stakeholders can work towards advancing knowledge and improving outcomes in the field of neurology. Despite its value, the data on neurology research in the Asian region also comes with several limitations:

1. The data was retrieved from the Scopus database, which serves as a widely used and reputable source for academic publications. However, it’s important to acknowledge that the completeness and accuracy of the data may be influenced by factors such as variations in data collection methods, indexing criteria, and publication practices across different journals and regions. Additionally, updates and revisions to the database may impact the consistency and reliability of the data over time.
2. The data covers a diverse range of countries and institutions in the Asian region, each with its own cultural, economic, and institutional contexts. These variations may influence research productivity, impact, and publication patterns, making direct comparisons challenging.
3. The data primarily focuses on quantitative metrics such as publication counts and citation counts, lacking contextual information such as research methodologies, collaborations, funding sources, and societal impact. This limits the depth of understanding and interpretation of research activities and outcomes.
4. One notable limitation of our study is that we did not conduct a comparative analysis of the progress of neurology research in the Asian region with that of other advanced countries or continents. For instance, comparing research outputs, citation impact, funding allocation, and collaboration patterns between Asia and regions such as North America or Europe could provide valuable benchmarks and identify areas where the Asian region excels or lags behind. Furthermore, comparative analyses with other regions could offer insights into factors driving research productivity and impact, such as investment in research infrastructure, funding mechanisms, policy frameworks, and cultural attitudes towards scientific inquiry. Understanding these factors can inform evidence-based policy decisions, resource allocation strategies, and collaborative initiatives aimed at fostering excellence and innovation in neurology research across the globe. In future research endeavors, it would be prudent to include comparative analyses with other advanced countries or continents to provide a more comprehensive understanding of the strengths, weaknesses, and opportunities for improvement in neurology research within the Asian region. This would enhance the relevance and utility of our findings and contribute to the broader discourse on global neurology research trends and dynamics.

In conclusion, our comprehensive analysis of neurology research in the Asian region may shed light on key trends and challenges, revealing disparities in research output, citation impact, and publication practices. Despite notable contributions from leading countries and institutions, there remains a need for greater inclusivity, collaboration, and investment to address regional disparities and advance neurological research agendas. By providing a nuanced understanding of the landscape, our work underscores the importance of fostering interdisciplinary collaboration, leveraging multilingual data sources, and engaging diverse stakeholders to drive innovation and improve neurological care across Asia. Moving forward, our findings may serve as a foundation for targeted interventions, policy reforms, and capacity-building initiatives aimed at enhancing research quality, visibility, and impact in the dynamic field of neurology.

## Reference

1. Abdelsimar T. Omar, Kevin Ivan P. Chan, Erika P. Ong, Louie F. Dy, Daniel Alexander D. Go, Michael Paolo Capistrano, Sean Kendrich N. Cua, Jose Danilo B. Diestro, Adrian I. Espiritu, Julian Spears, Neurosurgical research in Southeast Asia: A bibliometric analysis, Journal of Clinical Neuroscience, Volume 106, 2022, Pages 159–165.

