## Supplementary Tables for "Neurological Research in 47 Asian Countries"

**List of Supplementary Tables**

**Table 1:**	The overall research productivity of Asia in neurology.

**Table 2:**	The list of 47 countries/regions with scholarly output, growth rate (%), total citations, citations per publications and field-weighted citation impact.

**Table 3:**	The list of 3000 universities with sector, scholarly output, growth rate (%), total citations, citations per publications and field-weighted citation impact.

**Table 4:**	The list of top 100 researchers with scholarly output, most recent publications, total citations, citations per publications, field-weighted citation impact and h-index.

**Table 5:**	The publications distribution in four quartile group (Q1-Q4).

**Table 6:**	The list of 100 sources with most publications, total citations, citations per publication, Source-Normalized Impact per Paper (SNIP), CiteScore 2022 and SCImago Journal Rank (SJR).

| **S#** | **Title** | **Overall** | **2013** | **2014** | **2015** | **2016** | **2017** | **2018** | **2019** | **2020** | **2021** | **2022** |
| --- | --- | --- | --- | --- | --- | --- | --- | --- | --- | --- | --- | --- |
| **1** | Scholarly Output | 128306 | 9752 | 10205 | 10347 | 10898 | 11520 | 12383 | 13806 | 15617 | 16775 | 17003 |
| **2** | Citations | 1725678 | 230950 | 232117 | 229200 | 207767 | 212673 | 188048 | 168195 | 144326 | 81254 | 31148 |
| **3** | Citations per Publication | 13.4 | 23.7 | 22.7 | 22.2 | 19.1 | 18.5 | 15.2 | 12.2 | 9.2 | 4.8 | 1.8 |
| **4** | Field-Weighted Citation Impact | 0.95 | 0.87 | 0.92 | 0.97 | 0.94 | 0.98 | 0.96 | 0.98 | 0.97 | 0.94 | 0.99 |

**Table 1:**	The overall research productivity of Asia in neurology.

| **S#** | **Country/Region** | **Scholarly Output** | **Growth Rate (%)** | **Citations** | **Citations per Publication** | **Field-Weighted Citation Impact** |
| --- | --- | --- | --- | --- | --- | --- |
| **1** | China | 42471 | 198.8 | 522913 | 12.3 | 0.94 |
| **2** | Japan | 29797 | 4.6 | 368137 | 12.4 | 0.85 |
| **3** | Australia | 17949 | 55.6 | 487042 | 27.1 | 1.69 |
| **4** | India | 15160 | 113.2 | 129209 | 8.5 | 0.72 |
| **5** | South Korea | 13787 | 19.1 | 207203 | 15 | 1.03 |
| **6** | Taiwan | 4709 | -0.4 | 75756 | 16.1 | 1.13 |
| **7** | Singapore | 2348 | 49.7 | 59888 | 25.5 | 1.79 |
| **8** | New Zealand | 2024 | 59.2 | 57751 | 28.5 | 1.89 |
| **9** | Hong Kong | 2003 | 125.5 | 58063 | 29 | 2.05 |
| **10** | Thailand | 1563 | 105.4 | 21569 | 13.8 | 1.08 |
| **11** | Malaysia | 1182 | 159.4 | 24792 | 21 | 1.65 |
| **12** | Pakistan | 809 | 269.2 | 16728 | 20.7 | 1.76 |
| **13** | Indonesia | 501 | 1110 | 9511 | 19 | 1.73 |
| **14** | Philippines | 434 | 468.8 | 11369 | 26.2 | 2.46 |
| **15** | Nepal | 282 | 983.3 | 6803 | 24.1 | 2.51 |
| **16** | Bangladesh | 270 | 562.5 | 10012 | 37.1 | 3.32 |
| **17** | Viet Nam | 241 | 642.9 | 11638 | 48.3 | 3.88 |
| **18** | Georgia | 238 | 583.3 | 6923 | 29.1 | 3.16 |
| **19** | Kazakhstan | 186 | 400 | 3256 | 17.5 | 2.07 |
| **20** | Sri Lanka | 174 | -21.1 | 10243 | 58.9 | 4.3 |
| **21** | Macao | 104 | 300 | 2639 | 25.4 | 2.18 |
| **22** | Armenia | 71 | 1200 | 425 | 6 | 0.84 |
| **23** | Myanmar | 49 | - | 449 | 9.2 | 1.1 |
| **24** | Azerbaijan | 47 | - | 459 | 9.8 | 1.71 |
| **25** | Cambodia | 30 | -66.7 | 356 | 11.9 | 0.87 |
| **26** | Mongolia | 29 | - | 530 | 18.3 | 2.6 |
| **27** | Uzbekistan | 25 | 50 | 88 | 3.5 | 0.32 |
| **28** | Kyrgyzstan | 20 | 33.3 | 5226 | 261.3 | 26.16 |
| **29** | Laos | 18 | -100 | 170 | 9.4 | 1.25 |
| **30** | Brunei Darussalam | 12 | - | 120 | 10 | 0.7 |
| **31** | Bhutan | 12 | - | 102 | 8.5 | 0.76 |
| **32** | Fiji | 9 | - | 1070 | 118.9 | 3.75 |
| **33** | New Caledonia | 5 | - | 75 | 15 | 1.24 |
| **34** | Tajikistan | 5 | - | 29 | 5.8 | 0.31 |
| **35** | Guam | 4 | - | 77 | 19.3 | 0.94 |
| **36** | Papua New Guinea | 4 | - | 59 | 14.8 | 1.04 |
| **37** | French Polynesia | 4 | - | 25 | 6.3 | 0.51 |
| **38** | North Korea | 4 | - | 14 | 3.5 | 0.19 |
| **39** | Maldives | 3 | - | 104 | 34.7 | 1.06 |
| **40** | Vanuatu | 3 | - | 7 | 2.3 | 1.9 |
| **41** | Palau | 1 | -100 | 32 | 32 | 0.55 |
| **42** | United States Minor Outlying Islands | 1 | - | 31 | 31 | 1.42 |
| **43** | Norfolk Island | 1 | -100 | 24 | 24 | 2.31 |
| **44** | Federated States of Micronesia | 1 | - | 21 | 21 | 1.06 |
| **45** | Marshall Islands | 1 | - | 11 | 11 | 0.52 |
| **46** | Turkmenistan | 1 | - | 10 | 10 | 0.76 |
| **47** | Tonga | 1 | - | 0 | 0 | 0 |

**Table 2:**	The list of 47 countries/regions with scholarly output, growth rate (%), total citations, citations per publications and field-weighted citation impact.

| **S#** | **Institution** | **Sector** | **Country/Region** | **Scholarly Output** | **Growth Rate (%)** | **Citations** | **Citations per Publication** | **Field-Weighted Citation Impact** |
| --- | --- | --- | --- | --- | --- | --- | --- | --- |
| **1** | Capital Medical University | academic | China | 5974 | 235.2 | 83373 | 14 | 1.05 |
| **2** | University of Melbourne | academic | Australia | 4624 | 83.8 | 154108 | 33.3 | 2.16 |
| **3** | University of Sydney | academic | Australia | 3684 | 92.6 | 123102 | 33.4 | 2.05 |
| **4** | Seoul National University | academic | South Korea | 3205 | 31.7 | 57303 | 17.9 | 1.24 |
| **5** | Monash University | academic | Australia | 2771 | 146.9 | 74217 | 26.8 | 1.84 |
| **6** | University of New South Wales | academic | Australia | 2497 | 82.8 | 84917 | 34 | 2.07 |
| **7** | Fudan University | academic | China | 2387 | 179.9 | 38012 | 15.9 | 1.18 |
| **8** | Sichuan University | academic | China | 2368 | 261.2 | 27137 | 11.5 | 0.91 |
| **9** | Shanghai Jiao Tong University | academic | China | 2320 | 153.6 | 36932 | 15.9 | 1.14 |
| **10** | University of Queensland | academic | Australia | 2034 | 70.1 | 50305 | 24.7 | 1.89 |
| **11** | Yonsei University | academic | South Korea | 1978 | 41.3 | 34273 | 17.3 | 1.25 |
| **12** | Peking University | academic | China | 1924 | 358.9 | 28391 | 14.8 | 1.12 |
| **13** | All India Institute of Medical Sciences, New Delhi | government | India | 1823 | 161.4 | 16886 | 9.3 | 0.84 |
| **14** | National Institute of Mental Health and Neurosciences | medical | India | 1638 | 94.1 | 14509 | 8.9 | 0.74 |
| **15** | University of Ulsan | academic | South Korea | 1553 | 69.6 | 22387 | 14.4 | 0.99 |
| **16** | Florey Institute of Neuroscience and Mental Health | academic | Australia | 1535 | 71.8 | 60934 | 39.7 | 2.46 |
| **17** | Zhejiang University | academic | China | 1519 | 322.2 | 20827 | 13.7 | 1.19 |
| **18** | Royal Melbourne Hospital | medical | Australia | 1497 | 88.2 | 36997 | 24.7 | 1.75 |
| **19** | Sungkyunkwan University | academic | South Korea | 1481 | 17.6 | 26070 | 17.6 | 1.22 |
| **20** | Tohoku University | academic | Japan | 1478 | 11.4 | 30401 | 20.6 | 1.25 |
| **21** | The University of Tokyo | academic | Japan | 1468 | 58 | 29191 | 19.9 | 1.48 |
| **22** | Sun Yat-Sen University | academic | China | 1456 | 127.8 | 19782 | 13.6 | 1.06 |
| **23** | Chinese Academy of Medical Sciences | government | China | 1424 | 256.7 | 16903 | 11.9 | 1.06 |
| **24** | Nanjing Medical University | academic | China | 1348 | 323.4 | 18786 | 13.9 | 1.08 |
| **25** | Central South University | academic | China | 1334 | 177.1 | 19691 | 14.8 | 1.18 |
| **26** | National University of Singapore | academic | Singapore | 1304 | 61.9 | 36940 | 28.3 | 1.99 |
| **27** | Catholic University of Korea | academic | South Korea | 1285 | 6.6 | 14577 | 11.3 | 0.86 |
| **28** | Postgraduate Institute of Medical Education and Research | academic | India | 1246 | 83 | 10027 | 8 | 0.79 |
| **29** | Nagoya University | academic | Japan | 1233 | 34.8 | 27407 | 22.2 | 1.46 |
| **30** | Southern Medical University | academic | China | 1176 | 231 | 14136 | 12 | 1.02 |
| **31** | Korea University | academic | South Korea | 1164 | 16.3 | 22061 | 19 | 1.26 |
| **32** | Kyoto University | academic | Japan | 1143 | 34.1 | 21407 | 18.7 | 1.34 |
| **33** | National Yang Ming Chiao Tung University | academic | Taiwan | 1128 | 21.8 | 21931 | 19.4 | 1.18 |
| **34** | Zhengzhou University | academic | China | 1106 | 916 | 12680 | 11.5 | 1.22 |
| **35** | General Hospital of People's Liberation Army | medical | China | 1086 | 100 | 12155 | 11.2 | 0.81 |
| **36** | Osaka University | academic | Japan | 1073 | 29.1 | 16712 | 15.6 | 1.28 |
| **37** | Shandong University | academic | China | 1049 | 227.9 | 13177 | 12.6 | 0.96 |
| **38** | Chang Gung University | academic | Taiwan | 1048 | -12.7 | 15166 | 14.5 | 1.01 |
| **39** | University of Western Australia | academic | Australia | 1037 | 55.7 | 34971 | 33.7 | 2.23 |
| **40** | Juntendo University | academic | Japan | 1030 | 52.7 | 14312 | 13.9 | 1.14 |
| **41** | Keio University | academic | Japan | 1029 | 23.9 | 13913 | 13.5 | 1 |
| **42** | Huazhong University of Science and Technology | academic | China | 999 | 253.8 | 20562 | 20.6 | 1.88 |
| **43** | National Center of Neurology and Psychiatry Kodaira | government | Japan | 989 | 0 | 22989 | 23.2 | 1.63 |
| **44** | Tianjin Medical University | academic | China | 940 | 186.8 | 15679 | 16.7 | 1.15 |
| **45** | Kyushu University | academic | Japan | 940 | -5.4 | 12938 | 13.8 | 0.96 |
| **46** | Veterans General Hospital-Taipei | medical | Taiwan | 939 | 54.5 | 19154 | 20.4 | 1.34 |
| **47** | Hallym University | academic | South Korea | 935 | 103.7 | 13407 | 14.3 | 1 |
| **48** | Inje University | academic | South Korea | 928 | 40 | 11596 | 12.5 | 0.83 |
| **49** | Tokyo Women's Medical University | academic | Japan | 923 | -21.2 | 11842 | 12.8 | 0.94 |
| **50** | Nanjing University | academic | China | 918 | 87.9 | 18731 | 20.4 | 1.42 |
| **51** | Soochow University | academic | China | 917 | 248.1 | 11265 | 12.3 | 0.95 |
| **52** | Second Military Medical University | academic | China | 907 | 2.4 | 10800 | 11.9 | 0.93 |
| **53** | Queensland Health | government | Australia | 897 | 127.3 | 18125 | 20.2 | 1.55 |
| **54** | Chiba University | academic | Japan | 894 | 27.4 | 12843 | 14.4 | 1.15 |
| **55** | Jilin University | academic | China | 877 | 114.8 | 10872 | 12.4 | 0.87 |
| **56** | Chinese Academy of Sciences | government | China | 875 | 214.6 | 17882 | 20.4 | 1.46 |
| **57** | National Taiwan University | academic | Taiwan | 852 | 22.7 | 16142 | 18.9 | 1.42 |
| **58** | Chinese University of Hong Kong | academic | Hong Kong | 846 | 27 | 28092 | 33.2 | 2.36 |
| **59** | Murdoch Children's Research Institute | academic | Australia | 846 | 151.2 | 22530 | 26.6 | 1.94 |
| **60** | Kyung Hee University | academic | South Korea | 816 | 11.8 | 13615 | 16.7 | 1.15 |
| **61** | The University of Hong Kong | academic | Hong Kong | 793 | 141.5 | 20937 | 26.4 | 1.81 |
| **62** | Wenzhou Medical University | academic | China | 777 | 450 | 9000 | 11.6 | 0.93 |
| **63** | Neuroscience Research Australia | academic | Australia | 760 | -37.5 | 33947 | 44.7 | 2.07 |
| **64** | Royal Children's Hospital | medical | Australia | 747 | 53.7 | 37421 | 50.1 | 2.94 |
| **65** | Xi'an Jiaotong University | academic | China | 746 | 460.9 | 9806 | 13.1 | 1.25 |
| **66** | University of Newcastle | academic | Australia | 736 | 62 | 18422 | 25 | 1.84 |
| **67** | Royal Prince Alfred Hospital | medical | Australia | 722 | 86.7 | 19388 | 26.9 | 1.8 |
| **68** | China Medical University | academic | China | 717 | 106.5 | 10103 | 14.1 | 1.07 |
| **69** | Fujian Medical University | academic | China | 715 | 562.5 | 7020 | 9.8 | 0.98 |
| **70** | University of Tsukuba | academic | Japan | 711 | 112.5 | 9599 | 13.5 | 1.22 |
| **71** | University of Otago | academic | New Zealand | 706 | 38.6 | 22007 | 31.2 | 1.92 |
| **72** | Macquarie University | academic | Australia | 705 | 103.8 | 16101 | 22.8 | 1.86 |
| **73** | Hokkaido University | academic | Japan | 704 | 13.2 | 9708 | 13.8 | 1.04 |
| **74** | Fourth Military Medical University | academic | China | 672 | 83.1 | 8064 | 12 | 0.9 |
| **75** | Taipei Medical University | academic | Taiwan | 665 | 65.4 | 12730 | 19.1 | 1.52 |
| **76** | Niigata University | academic | Japan | 662 | 9.5 | 11721 | 17.7 | 1.13 |
| **77** | Singapore Health Services | other | Singapore | 651 | 95.3 | 10586 | 16.3 | 1.29 |
| **78** | National Cerebral and Cardiovascular Center | government | Japan | 643 | -4.5 | 17645 | 27.4 | 1.85 |
| **79** | La Trobe University | academic | Australia | 641 | 187.1 | 21244 | 33.1 | 2.41 |
| **80** | Kyungpook National University | academic | South Korea | 641 | 17.6 | 9322 | 14.5 | 0.95 |
| **81** | Tokyo Medical and Dental University | academic | Japan | 633 | 49.2 | 7135 | 11.3 | 0.95 |
| **82** | The University of Auckland | academic | New Zealand | 631 | 78.6 | 22164 | 35.1 | 2.42 |
| **83** | Sanjay Gandhi Postgraduate Institute of Medical Sciences | academic | India | 630 | 27.3 | 7783 | 12.4 | 0.93 |
| **84** | Pusan National University | academic | South Korea | 628 | 65.9 | 7053 | 11.2 | 0.76 |
| **85** | Hiroshima University | academic | Japan | 626 | 37.5 | 7078 | 11.3 | 0.87 |
| **86** | Alfred Health | medical | Australia | 608 | 26 | 17868 | 29.4 | 1.69 |
| **87** | Soonchunhyang University | academic | South Korea | 606 | 100 | 7344 | 12.1 | 0.83 |
| **88** | Nippon Medical School | academic | Japan | 594 | 10.6 | 7344 | 12.4 | 0.83 |
| **89** | National Hospital Organization, Japan | medical | Japan | 593 | 47.6 | 7085 | 11.9 | 1.07 |
| **90** | Saitama Medical University | academic | Japan | 586 | 4 | 6951 | 11.9 | 0.92 |
| **91** | Sree Chitra Tirunal Institute for Medical Sciences and Technology | government | India | 582 | 120.5 | 10329 | 17.7 | 1.54 |
| **92** | Wuhan University | academic | China | 576 | 361.5 | 16245 | 28.2 | 2.32 |
| **93** | Anhui Medical University | academic | China | 575 | 323.1 | 6434 | 11.2 | 0.97 |
| **94** | Okayama University | academic | Japan | 568 | -18.2 | 6955 | 12.2 | 0.85 |
| **95** | Kanazawa University | academic | Japan | 559 | 62.2 | 8469 | 15.2 | 1.17 |
| **96** | Tongji University | academic | China | 555 | 245.2 | 6289 | 11.3 | 0.94 |
| **97** | Chonnam National University | academic | South Korea | 547 | 9.3 | 7927 | 14.5 | 0.92 |
| **98** | Third Military Medical University | academic | China | 543 | 22.2 | 12598 | 23.2 | 1.45 |
| **99** | Kobe University | academic | Japan | 542 | 142.9 | 8829 | 16.3 | 1.24 |
| **100** | Hanyang University | academic | South Korea | 542 | 25.5 | 8537 | 15.8 | 1.02 |
| **101** | Qingdao University | academic | China | 534 | 307.7 | 9669 | 18.1 | 1.34 |
| **102** | Chongqing Medical University | academic | China | 529 | 592.9 | 7210 | 13.6 | 1.13 |
| **103** | University of Adelaide | academic | Australia | 527 | 56.8 | 19442 | 36.9 | 2.6 |
| **104** | Fujita Health University | academic | Japan | 521 | 60.5 | 9833 | 18.9 | 1.62 |
| **105** | Yokohama City University | academic | Japan | 509 | 47.4 | 9577 | 18.8 | 1.24 |
| **106** | Hamamatsu University School of Medicine | academic | Japan | 501 | 186.2 | 7089 | 14.1 | 1.22 |
| **107** | Mahidol University | academic | Thailand | 499 | 111.4 | 8630 | 17.3 | 1.21 |
| **108** | Guangzhou Medical College | academic | China | 495 | 330 | 7076 | 14.3 | 1.07 |
| **109** | The Jikei University School of Medicine | academic | Japan | 495 | 93 | 8576 | 17.3 | 1.18 |
| **110** | Royal North Shore Hospital | medical | Australia | 493 | 44.4 | 12619 | 25.6 | 1.56 |
| **111** | University of Malaya | academic | Malaysia | 493 | 93.5 | 7757 | 15.7 | 1.18 |
| **112** | Shinshu University | academic | Japan | 491 | 80 | 4422 | 9 | 0.78 |
| **113** | Griffith University Queensland | academic | Australia | 488 | 180.8 | 12861 | 26.4 | 2.07 |
| **114** | Aichi Medical University | academic | Japan | 488 | 141.7 | 6534 | 13.4 | 1.07 |
| **115** | Tokushima University | academic | Japan | 476 | 16.7 | 6228 | 13.1 | 0.98 |
| **116** | Tsinghua University | academic | China | 475 | 492.3 | 6522 | 13.7 | 1.16 |
| **117** | Hunter New England Health | medical | Australia | 475 | 103 | 12448 | 26.2 | 1.77 |
| **118** | Christian Medical College | academic | India | 472 | 103 | 11656 | 24.7 | 1.7 |
| **119** | Tokyo Medical University | academic | Japan | 468 | -20.9 | 7161 | 15.3 | 1.03 |
| **120** | University of Tasmania | academic | Australia | 467 | 305.3 | 11357 | 24.3 | 1.88 |
| **121** | Hebei Medical University | academic | China | 465 | 389.5 | 5708 | 12.3 | 0.84 |
| **122** | Nanchang University | academic | China | 465 | 841.7 | 4338 | 9.3 | 0.9 |
| **123** | The Children's Hospital at Westmead | medical | Australia | 463 | 75.7 | 15894 | 34.3 | 2.13 |
| **124** | Shandong First Medical University & Shandong Academy of Medical Sciences | academic | China | 460 | 363.2 | 6083 | 13.2 | 1.06 |
| **125** | Fukuoka University | academic | Japan | 458 | 25 | 5702 | 12.4 | 0.97 |
| **126** | Ewha Womans University | academic | South Korea | 453 | 54.1 | 7437 | 16.4 | 1.05 |
| **127** | Kumamoto University | academic | Japan | 448 | -10.8 | 5599 | 12.5 | 0.92 |
| **128** | Flinders University | academic | Australia | 443 | 109.1 | 12211 | 27.6 | 1.64 |
| **129** | Keimyung University | academic | South Korea | 435 | 200 | 6699 | 15.4 | 1.09 |
| **130** | Harbin Medical University | academic | China | 431 | 143.8 | 6403 | 14.9 | 1.06 |
| **131** | Jichi Medical University | academic | Japan | 430 | -17.6 | 6382 | 14.8 | 1.04 |
| **132** | China Medical University Taichung | academic | Taiwan | 429 | -2.5 | 7415 | 17.3 | 1.42 |
| **133** | Southeast University, Nanjing | academic | China | 428 | 347.1 | 5662 | 13.2 | 1.08 |
| **134** | Royal Brisbane and Women's Hospital | medical | Australia | 427 | 44.1 | 10259 | 24 | 1.71 |
| **135** | Westmead Hospital | medical | Australia | 425 | 62.5 | 13698 | 32.2 | 1.93 |
| **136** | Chung-Ang University | academic | South Korea | 422 | 32.3 | 6156 | 14.6 | 0.93 |
| **137** | Kitasato University | academic | Japan | 418 | 20.5 | 10284 | 24.6 | 1.18 |
| **138** | University of Electronic Science and Technology of China | academic | China | 417 | 925 | 7127 | 17.1 | 1.45 |
| **139** | Ajou University | academic | South Korea | 417 | 100 | 5666 | 13.6 | 1.07 |
| **140** | Osaka Metropolitan University | academic | Japan | 416 | 103.3 | 4626 | 11.1 | 0.96 |
| **141** | Seth GS Medical College and KEM Hospital | medical | India | 415 | 72.7 | 3191 | 7.7 | 1.02 |
| **142** | St. Vincent's Hospital Melbourne | medical | Australia | 414 | 112.5 | 8308 | 20.1 | 1.34 |
| **143** | Gachon University | academic | South Korea | 407 | 25 | 5329 | 13.1 | 0.95 |
| **144** | Curtin University | academic | Australia | 406 | 143.5 | 14186 | 34.9 | 2.88 |
| **145** | MOH Holdings Pte Ltd. | corporate | Singapore | 403 | 180 | 6785 | 16.8 | 1.52 |
| **146** | Western Sydney University | academic | Australia | 399 | 63.6 | 16345 | 41 | 2.98 |
| **147** | Nara Medical University | academic | Japan | 398 | 195 | 3379 | 8.5 | 0.74 |
| **148** | Chulalongkorn University | academic | Thailand | 398 | 162.5 | 6293 | 15.8 | 1.16 |
| **149** | Dokkyo Medical University | academic | Japan | 396 | 14.3 | 4653 | 11.8 | 0.96 |
| **150** | Kyoto Prefectural University of Medicine | academic | Japan | 396 | 17.1 | 5860 | 14.8 | 1.06 |
| **151** | Deakin University | academic | Australia | 395 | 238.9 | 13740 | 34.8 | 2.86 |
| **152** | The George Institute for Global Health | academic | Australia | 391 | 163.2 | 13168 | 33.7 | 2.26 |
| **153** | National Defense Medical Center Taiwan | academic | Taiwan | 390 | -20.7 | 3956 | 10.1 | 0.73 |
| **154** | Jinan University | academic | China | 387 | 461.5 | 4552 | 11.8 | 1 |
| **155** | Fukushima Medical University | academic | Japan | 382 | 85.7 | 11968 | 31.3 | 1.76 |
| **156** | Yeungnam University | academic | South Korea | 380 | 23.5 | 4884 | 12.9 | 0.93 |
| **157** | Konkuk University | academic | South Korea | 379 | -5.6 | 5708 | 15.1 | 0.94 |
| **158** | Shantou University | academic | China | 376 | 211.8 | 6053 | 16.1 | 1.19 |
| **159** | Teikyo University | academic | Japan | 375 | 5.7 | 8527 | 22.7 | 1.58 |
| **160** | University of Toyama | academic | Japan | 375 | 57.1 | 5177 | 13.8 | 1.17 |
| **161** | Sapporo Medical University | academic | Japan | 374 | 26.9 | 3728 | 10 | 0.76 |
| **162** | Manipal Academy of Higher Education | academic | India | 369 | 147.4 | 3659 | 9.9 | 1.13 |
| **163** | Kaohsiung Medical University | academic | Taiwan | 365 | 4.4 | 4426 | 12.1 | 0.99 |
| **164** | Shenzhen University | academic | China | 360 | 750 | 5380 | 14.9 | 1.79 |
| **165** | Xuzhou Medical University | academic | China | 360 | 257.9 | 4568 | 12.7 | 1.03 |
| **166** | Showa University | academic | Japan | 358 | 13.3 | 4449 | 12.4 | 0.84 |
| **167** | University of Chinese Academy of Sciences | academic | China | 357 | 1450 | 4901 | 13.7 | 1.36 |
| **168** | Yamaguchi University | academic | Japan | 357 | 10.3 | 3940 | 11 | 0.75 |
| **169** | Chungnam National University | academic | South Korea | 357 | 275 | 4210 | 11.8 | 0.92 |
| **170** | Hyogo Medical University | academic | Japan | 351 | 7.4 | 4308 | 12.3 | 0.97 |
| **171** | Eulji University | academic | South Korea | 351 | 135 | 4649 | 13.2 | 0.99 |
| **172** | Dalian Medical University | academic | China | 350 | 491.7 | 4735 | 13.5 | 1.07 |
| **173** | International University of Health and Welfare | academic | Japan | 350 | 343.8 | 4329 | 12.4 | 1.13 |
| **174** | University of South Australia | academic | Australia | 348 | -5.3 | 10587 | 30.4 | 1.67 |
| **175** | Prince of Wales Hospital | medical | Australia | 346 | 75 | 12499 | 36.1 | 2.32 |
| **176** | Nagoya City University | academic | Japan | 341 | 76.9 | 4269 | 12.5 | 1.19 |
| **177** | Gunma University | academic | Japan | 338 | -2.6 | 4090 | 12.1 | 0.84 |
| **178** | Nanfang Hospital | medical | China | 334 | 200 | 4314 | 12.9 | 0.97 |
| **179** | Singapore General Hospital | medical | Singapore | 333 | 18.5 | 5282 | 15.9 | 1.34 |
| **180** | National Cheng Kung University | academic | Taiwan | 331 | -2.6 | 4690 | 14.2 | 0.99 |
| **181** | Sir Charles Gairdner Hospital | medical | Australia | 329 | -5.7 | 16871 | 51.3 | 3.34 |
| **182** | Beijing Normal University | academic | China | 324 | 176.9 | 6698 | 20.7 | 1.27 |
| **183** | Toho University | academic | Japan | 323 | 34.5 | 3873 | 12 | 0.94 |
| **184** | Nihon University | academic | Japan | 322 | 5.4 | 4615 | 14.3 | 0.92 |
| **185** | Samsung | corporate | South Korea | 322 | -3.2 | 5165 | 16 | 1 |
| **186** | Nantong University | academic | China | 319 | 108.3 | 5424 | 17 | 1.19 |
| **187** | Inha University | academic | South Korea | 319 | 21.7 | 5118 | 16 | 1.15 |
| **188** | King George's Medical University | academic | India | 319 | 117.4 | 4127 | 12.9 | 0.87 |
| **189** | University of Fukui | academic | Japan | 314 | 57.6 | 3760 | 12 | 1.19 |
| **190** | Kagoshima University | academic | Japan | 313 | 20 | 3415 | 10.9 | 0.74 |
| **191** | South China University of Technology | academic | China | 312 | 391.7 | 3484 | 11.2 | 0.97 |
| **192** | Kindai University | academic | Japan | 312 | -22.2 | 5306 | 17 | 1.35 |
| **193** | Tokyo Metropolitan Institute of Medical Science | medical | Japan | 311 | -44.7 | 5984 | 19.2 | 1.31 |
| **194** | Aga Khan University | academic | Pakistan | 311 | 200 | 6739 | 21.7 | 2.12 |
| **195** | Auckland District Health Board | medical | New Zealand | 311 | 94.7 | 6230 | 20 | 1.42 |
| **196** | Shanxi Medical University | academic | China | 310 | 340 | 3247 | 10.5 | 0.86 |
| **197** | Queensland University of Technology | academic | Australia | 309 | 290 | 8311 | 26.9 | 2.5 |
| **198** | Australian Catholic University | academic | Australia | 308 | 133.3 | 13162 | 42.7 | 2.64 |
| **199** | Gifu University | academic | Japan | 307 | 27.3 | 3028 | 9.9 | 0.8 |
| **200** | Kurume University | academic | Japan | 303 | 253.8 | 3395 | 11.2 | 0.88 |
| **201** | Henan Provincial People's Hospital | medical | China | 303 | 1383.3 | 3410 | 11.3 | 1.32 |
| **202** | Chongqing University | academic | China | 302 | 90.9 | 4999 | 16.6 | 1.05 |
| **203** | University of Delhi | academic | India | 302 | 58.1 | 2246 | 7.4 | 0.62 |
| **204** | Princess Alexandra Hospital Brisbane | medical | Australia | 297 | 111.8 | 4759 | 16 | 1.21 |
| **205** | University of Yamanashi | academic | Japan | 295 | 16.7 | 3169 | 10.7 | 1.12 |
| **206** | Mie University | academic | Japan | 294 | 25 | 4070 | 13.8 | 1.12 |
| **207** | National Defense Medical College Tokorozawa | academic | Japan | 291 | -34.9 | 5575 | 19.2 | 1.37 |
| **208** | Iwate Medical University | academic | Japan | 287 | 92.3 | 4141 | 14.4 | 1.32 |
| **209** | Kyorin University | academic | Japan | 287 | 181.3 | 4073 | 14.2 | 1.23 |
| **210** | Tokai University | academic | Japan | 285 | 38.1 | 3272 | 11.5 | 0.93 |
| **211** | Nagasaki University | academic | Japan | 283 | 11.5 | 3042 | 10.7 | 0.78 |
| **212** | Dongguk University | academic | South Korea | 281 | 150 | 5656 | 20.1 | 1.39 |
| **213** | Kobe City Medical Center General Hospital | medical | Japan | 281 | 141.2 | 3157 | 11.2 | 1.02 |
| **214** | Hirosaki University | academic | Japan | 280 | 63.6 | 3245 | 11.6 | 1.06 |
| **215** | St. Marianna University School of Medicine | academic | Japan | 279 | 87.5 | 2446 | 8.8 | 0.66 |
| **216** | Gyeongsang National University | academic | South Korea | 276 | 23.8 | 3816 | 13.8 | 0.94 |
| **217** | Kawasaki Medical School | academic | Japan | 274 | -30.3 | 3186 | 11.6 | 0.79 |
| **218** | Dong-A University | academic | South Korea | 274 | 31.3 | 3911 | 14.3 | 0.93 |
| **219** | Tokyo Metropolitan Neurological Hospital | medical | Japan | 271 | -17.2 | 3723 | 13.7 | 1.3 |
| **220** | Ehime University | academic | Japan | 270 | -20.7 | 3284 | 12.2 | 0.8 |
| **221** | Yangzhou University | academic | China | 269 | 400 | 3149 | 11.7 | 1.28 |
| **222** | Hong Kong Polytechnic University | academic | Hong Kong | 269 | 338.5 | 6219 | 23.1 | 1.73 |
| **223** | Hunter Medical Research Institute | academic | Australia | 268 | 112.5 | 6685 | 24.9 | 1.62 |
| **224** | Osaka Medical and Pharmaceutical University | academic | Japan | 267 | 20 | 3010 | 11.3 | 0.77 |
| **225** | Jeonbuk National University | academic | South Korea | 263 | 17.4 | 2706 | 10.3 | 0.68 |
| **226** | University of Technology Sydney | academic | Australia | 262 | 169.2 | 10350 | 39.5 | 3.35 |
| **227** | Kunming Medical College | academic | China | 260 | 533.3 | 2994 | 11.5 | 0.81 |
| **228** | Tottori University | academic | Japan | 259 | -12.5 | 3203 | 12.4 | 0.86 |
| **229** | Xiamen University | academic | China | 257 | 161.1 | 14262 | 55.5 | 3.66 |
| **230** | Guangxi Medical University | academic | China | 251 | 266.7 | 3381 | 13.5 | 1.05 |
| **231** | University of Science and Technology of China | academic | China | 251 | 2166.7 | 2795 | 11.1 | 1.34 |
| **232** | Sydney Children's Hospital | medical | Australia | 249 | 377.8 | 5550 | 22.3 | 1.85 |
| **233** | National Hospital Organization Shizuoka Institute of Epilepsy and Neurological Disorders | medical | Japan | 249 | -13.6 | 3444 | 13.8 | 0.89 |
| **234** | University of the Philippines | academic | Philippines | 247 | 1140 | 8487 | 34.4 | 3.41 |
| **235** | Nanjing Brain Hospital | medical | China | 245 | 410 | 2398 | 9.8 | 0.92 |
| **236** | Royal Adelaide Hospital | medical | Australia | 241 | 133.3 | 4417 | 18.3 | 1.61 |
| **237** | Tokyo Metropolitan Geriatric Hospital and Institute of Gerontology | medical | Japan | 241 | 121.4 | 5617 | 23.3 | 1.68 |
| **238** | Edith Cowan University | academic | Australia | 238 | 111.1 | 8248 | 34.7 | 2.46 |
| **239** | RIKEN | government | Japan | 237 | -20 | 7270 | 30.7 | 1.52 |
| **240** | Guangzhou University of Chinese Medicine | academic | China | 234 | 900 | 2614 | 11.2 | 1.11 |
| **241** | Kangwon National University | academic | South Korea | 234 | -5 | 3150 | 13.5 | 0.96 |
| **242** | Auckland University of Technology | academic | New Zealand | 232 | 250 | 17938 | 77.3 | 5.03 |
| **243** | Shiga University of Medical Science | academic | Japan | 231 | 128.6 | 3096 | 13.4 | 1.12 |
| **244** | Wakayama Medical University | academic | Japan | 231 | 166.7 | 2804 | 12.1 | 1.06 |
| **245** | All India Institute of Medical Sciences, Rishikesh | academic | India | 231 | 5800 | 754 | 3.3 | 0.73 |
| **246** | Nanyang Technological University | academic | Singapore | 230 | 350 | 8608 | 37.4 | 2.79 |
| **247** | Osaka City General Hospital | medical | Japan | 228 | 75 | 2380 | 10.4 | 0.89 |
| **248** | CHA University | academic | South Korea | 228 | 42.1 | 3188 | 14 | 1.05 |
| **249** | University of Notre Dame Australia | academic | Australia | 227 | 94.1 | 5208 | 22.9 | 1.36 |
| **250** | Eisai Co., Ltd. | corporate | Japan | 226 | 253.8 | 6960 | 30.8 | 2.33 |
| **251** | Nizam's Institute of Medical Sciences | medical | India | 226 | -21.4 | 2529 | 11.2 | 0.69 |
| **252** | National Cancer Center Japan | government | Japan | 225 | 71.4 | 4641 | 20.6 | 1.51 |
| **253** | Concord Repatriation General Hospital | medical | Australia | 223 | 146.7 | 4842 | 21.7 | 1.55 |
| **254** | Murdoch University | academic | Australia | 220 | 280 | 4574 | 20.8 | 1.6 |
| **255** | Tzu Chi University | academic | Taiwan | 220 | 10 | 2976 | 13.5 | 0.91 |
| **256** | Philippine General Hospital | medical | Philippines | 220 | 783.3 | 3265 | 14.8 | 1.42 |
| **257** | Australian National University | academic | Australia | 219 | -29.2 | 8604 | 39.3 | 2.49 |
| **258** | St. Vincent's Hospital Sydney | medical | Australia | 218 | 23.5 | 4042 | 18.5 | 1.25 |
| **259** | Kohnan Hospital | medical | Japan | 216 | 29.4 | 2381 | 11 | 1.07 |
| **260** | Northern Sydney Local Health District | government | Australia | 216 | 300 | 7294 | 33.8 | 1.9 |
| **261** | Shanghai University of Traditional Chinese Medicine | academic | China | 213 | 900 | 1914 | 9 | 1.08 |
| **262** | China-Japan Friendship Hospital | medical | China | 213 | 58.3 | 2214 | 10.4 | 0.8 |
| **263** | Beijing University of Chinese Medicine | academic | China | 210 | 1633.3 | 2561 | 12.2 | 1.01 |
| **264** | Yonsei University Mirae Campus | academic | South Korea | 210 | -13 | 2547 | 12.1 | 0.75 |
| **265** | National Center for Child Health and Development | government | Japan | 209 | 86.7 | 2262 | 10.8 | 0.76 |
| **266** | Southwest Medical University | academic | China | 208 | 925 | 2071 | 10 | 1.03 |
| **267** | Jeju National University | academic | South Korea | 208 | 4.8 | 2889 | 13.9 | 0.9 |
| **268** | Ministry of Health of People's Republic of China | government | China | 206 | 0 | 3554 | 17.3 | 1.03 |
| **269** | Epworth HealthCare | medical | Australia | 205 | 5.3 | 4658 | 22.7 | 1.45 |
| **270** | Kolling Institute of Medical Research | academic | Australia | 204 | 250 | 7207 | 35.3 | 1.96 |
| **271** | Lanzhou University | academic | China | 203 | 2900 | 1756 | 8.7 | 0.83 |
| **272** | Jawaharlal Institute of Postgraduate Medical Education and Research | academic | India | 203 | 16.7 | 2237 | 11 | 1.35 |
| **273** | Box Hill Hospital | medical | Australia | 200 | 1700 | 4245 | 21.2 | 1.79 |
| **274** | Canterbury District Health Board | medical | New Zealand | 198 | 383.3 | 3768 | 19 | 1.75 |
| **275** | Veterans General Hospital-Taichung Taiwan | medical | Taiwan | 193 | 41.7 | 2294 | 11.9 | 0.92 |
| **276** | Guangdong Provincial People's Hospital | medical | China | 192 | 700 | 1720 | 9 | 0.86 |
| **277** | Yamagata University | academic | Japan | 192 | -34.5 | 3564 | 18.6 | 1.15 |
| **278** | Kansai Medical University | academic | Japan | 189 | 68.4 | 2499 | 13.2 | 1 |
| **279** | Sichuan Provincial People's Hospital | medical | China | 188 | 825 | 2404 | 12.8 | 1.27 |
| **280** | Thai Red Cross Society | other | Thailand | 188 | 211.1 | 2355 | 12.5 | 1.03 |
| **281** | CAS - Institute of Psychology | academic | China | 187 | 122.2 | 3362 | 18 | 1.39 |
| **282** | Guizhou Medical University | academic | China | 184 | 287.5 | 2473 | 13.4 | 0.95 |
| **283** | Bombay Hospital and Medical Research Centre | medical | India | 184 | 68.8 | 1384 | 7.5 | 0.6 |
| **284** | Swinburne University of Technology | academic | Australia | 182 | 142.9 | 4592 | 25.2 | 1.52 |
| **285** | General Hospital | medical | China | 179 | 153.3 | 2491 | 13.9 | 1.1 |
| **286** | Royal Perth Hospital | medical | Australia | 178 | -44.1 | 6687 | 37.6 | 1.88 |
| **287** | Zhejiang Provincial People's Hospital | medical | China | 178 | 1100 | 1945 | 10.9 | 0.99 |
| **288** | Southwest University | academic | China | 177 | 61.5 | 3968 | 22.4 | 1.31 |
| **289** | Dankook University | academic | South Korea | 177 | 58.3 | 2162 | 12.2 | 0.79 |
| **290** | University of Wollongong | academic | Australia | 175 | 27.3 | 3497 | 20 | 1.27 |
| **291** | National Center for Geriatrics and Gerontology | government | Japan | 175 | 180 | 3236 | 18.5 | 1.29 |
| **292** | National Cancer Center Korea | medical | South Korea | 175 | 105.9 | 4086 | 23.3 | 1.83 |
| **293** | ARC Centre of Excellence in Cognition and its Disorders | academic | Australia | 174 | -90.9 | 5539 | 31.8 | 1.73 |
| **294** | Wonkwang University | academic | South Korea | 171 | -31.6 | 2181 | 12.8 | 0.81 |
| **295** | Kanazawa Medical University | academic | Japan | 170 | -68.2 | 2295 | 13.5 | 0.95 |
| **296** | Khon Kaen University | academic | Thailand | 170 | -38.9 | 2262 | 13.3 | 0.68 |
| **297** | Hangzhou First People's Hospital | medical | China | 167 | 2850 | 1985 | 11.9 | 1.35 |
| **298** | Nankai University | academic | China | 165 | 1850 | 1409 | 8.5 | 0.93 |
| **299** | Dr. Ram Manohar Lohia Hospital | medical | India | 165 | 150 | 955 | 5.8 | 0.56 |
| **300** | Shanghai University | academic | China | 164 | 328.6 | 4256 | 26 | 1.9 |
| **301** | Buddhist Tzu-Chi General Hospital Taiwan | medical | Taiwan | 164 | -36.4 | 1987 | 12.1 | 0.85 |
| **302** | Hainan Medical University | academic | China | 164 | 61.5 | 823 | 5 | 0.64 |
| **303** | Queensland Institute of Medical Research | academic | Australia | 163 | 600 | 4757 | 29.2 | 2.34 |
| **304** | Oita University | academic | Japan | 162 | -10 | 2141 | 13.2 | 0.72 |
| **305** | Nanjing University of Chinese Medicine | academic | China | 161 | 600 | 2110 | 13.1 | 1.07 |
| **306** | Tohoku Medical and Pharmaceutical University | academic | Japan | 161 | - | 2680 | 16.6 | 1.82 |
| **307** | Saga University | academic | Japan | 158 | 0 | 1930 | 12.2 | 0.83 |
| **308** | University of Occupational and Environmental Health, Japan | academic | Japan | 158 | -31.6 | 2093 | 13.2 | 0.94 |
| **309** | Chung Shan Medical University | academic | Taiwan | 158 | 45.5 | 1753 | 11.1 | 0.69 |
| **310** | CSIRO | government | Australia | 157 | 40 | 5741 | 36.6 | 2.06 |
| **311** | Akita University | academic | Japan | 157 | 91.7 | 1685 | 10.7 | 0.82 |
| **312** | Universiti Kebangsaan Malaysia | academic | Malaysia | 157 | 163.6 | 7747 | 49.3 | 3.85 |
| **313** | Amrita Vishwa Vidyapeetham | academic | India | 157 | 300 | 4229 | 26.9 | 1.87 |
| **314** | Telethon Kids Institute | academic | Australia | 157 | 175 | 3903 | 24.9 | 1.66 |
| **315** | Beijing Tsinghua Chang Gung Hospital | medical | China | 157 | - | 1112 | 7.1 | 0.71 |
| **316** | Agency for Science, Technology and Research, Singapore | government | Singapore | 156 | 0 | 3457 | 22.2 | 1.68 |
| **317** | Fu Jen Catholic University | academic | Taiwan | 156 | -29.4 | 2170 | 13.9 | 0.8 |
| **318** | James Cook University Queensland | academic | Australia | 153 | 88.9 | 2339 | 15.3 | 1.13 |
| **319** | Wannan Medical College | academic | China | 153 | 540 | 2503 | 16.4 | 1.73 |
| **320** | Seirei Hamamatsu General Hospital | medical | Japan | 153 | 50 | 1087 | 7.1 | 0.57 |
| **321** | Chosun University | academic | South Korea | 151 | 5.9 | 2005 | 13.3 | 0.93 |
| **322** | Qingdao Municipal Hospital | medical | China | 151 | 211.1 | 5538 | 36.7 | 2.14 |
| **323** | National Hospital Organization Kyushu Medical Center | medical | Japan | 150 | 7.7 | 2508 | 16.7 | 1.1 |
| **324** | Zunyi Medical University | academic | China | 149 | 1233.3 | 1178 | 7.9 | 0.75 |
| **325** | Rajiv Gandhi University of Health Sciences | academic | India | 147 | 157.1 | 880 | 6 | 0.45 |
| **326** | Guangdong Provincial Hospital of Traditional Chinese Medicine | medical | China | 146 | 1366.7 | 1726 | 11.8 | 1.3 |
| **327** | Zhejiang Chinese Medical University | academic | China | 145 | - | 1198 | 8.3 | 1.12 |
| **328** | Lady Hardinge Medical College | academic | India | 144 | 42.9 | 1278 | 8.9 | 0.75 |
| **329** | Tan Tock Seng Hospital | medical | Singapore | 144 | -16.7 | 2508 | 17.4 | 1.37 |
| **330** | Children’s Health Queensland | medical | Australia | 144 | 3100 | 2745 | 19.1 | 2.91 |
| **331** | Gangneung Asan Hospital | medical | South Korea | 142 | 7.7 | 1678 | 11.8 | 0.76 |
| **332** | National Hospital Organization Sendai Medical Center | medical | Japan | 142 | 30 | 2057 | 14.5 | 1.04 |
| **333** | University of South China | academic | China | 141 | 1100 | 1946 | 13.8 | 1.26 |
| **334** | Asahikawa Medical University | academic | Japan | 141 | 8.3 | 2160 | 15.3 | 0.9 |
| **335** | Hangzhou Normal University | academic | China | 140 | 157.1 | 3125 | 22.3 | 1.5 |
| **336** | National Hospital Organization Osaka National Hospital | medical | Japan | 139 | 350 | 1635 | 11.8 | 1.5 |
| **337** | Xidian University | academic | China | 138 | 233.3 | 2600 | 18.8 | 1.35 |
| **338** | Shimane University | academic | Japan | 137 | -10 | 1671 | 12.2 | 0.97 |
| **339** | Madras Medical College | academic | India | 137 | 166.7 | 617 | 4.5 | 0.49 |
| **340** | All India Institute of Medical Sciences, Jodhpur | academic | India | 137 | 4400 | 4737 | 34.6 | 3.72 |
| **341** | Jiangsu University | academic | China | 136 | 350 | 1650 | 12.1 | 0.85 |
| **342** | China Academy of Chinese Medical Sciences | academic | China | 136 | 400 | 1768 | 13 | 1.1 |
| **343** | SMS Medical College | medical | India | 136 | 33.3 | 595 | 4.4 | 0.41 |
| **344** | National Institutes for Quantum and Radiological Science and Technology | government | Japan | 135 | 81.8 | 2539 | 18.8 | 1.64 |
| **345** | Academia Sinica Taiwan | academic | Taiwan | 135 | 157.1 | 4416 | 32.7 | 2.31 |
| **346** | Chengdu University of Traditional Chinese Medicine | academic | China | 132 | 400 | 1757 | 13.3 | 1.2 |
| **347** | Woolcock Institute of Medical Research | academic | Australia | 132 | 166.7 | 3950 | 29.9 | 1.57 |
| **348** | Catholic University of Daegu | academic | South Korea | 131 | -9.1 | 1894 | 14.5 | 0.76 |
| **349** | Chi-Mei Medical Center | medical | Taiwan | 131 | -58.8 | 1589 | 12.1 | 1.09 |
| **350** | Banaras Hindu University | academic | India | 130 | 371.4 | 6056 | 46.6 | 4.27 |
| **351** | Kochi University | academic | Japan | 130 | 144.4 | 1619 | 12.5 | 1.07 |
| **352** | I-Shou University | academic | Taiwan | 129 | 75 | 1461 | 11.3 | 1.03 |
| **353** | South Australian Health And Medical Research Institute | academic | Australia | 129 | - | 2416 | 18.7 | 1.88 |
| **354** | Ningxia Medical University | academic | China | 128 | 91.7 | 1295 | 10.1 | 0.82 |
| **355** | Xinxiang Medical College | academic | China | 127 | 320 | 1577 | 12.4 | 0.98 |
| **356** | Changhua Christian Hospital | medical | Taiwan | 127 | 53.3 | 1295 | 10.2 | 0.75 |
| **357** | CAS - Institute of Automation | academic | China | 127 | 25 | 3658 | 28.8 | 1.53 |
| **358** | Kagawa University | academic | Japan | 126 | -70.6 | 1789 | 14.2 | 0.8 |
| **359** | Taipei City Hospital | medical | Taiwan | 126 | -73.7 | 1776 | 14.1 | 0.8 |
| **360** | University of Macau | academic | Macao | 126 | 133.3 | 3196 | 25.4 | 2.04 |
| **361** | Xinjiang Medical University | academic | China | 124 | 766.7 | 896 | 7.2 | 0.84 |
| **362** | Nippon Telegraph & Telephone | corporate | Japan | 124 | 283.3 | 926 | 7.5 | 0.66 |
| **363** | Japan Society for the Promotion of Science | government | Japan | 123 | -8.3 | 1978 | 16.1 | 0.99 |
| **364** | Korea Institute of Science and Technology | government | South Korea | 123 | 300 | 2358 | 19.2 | 1.14 |
| **365** | Jining Medical College | academic | China | 123 | 275 | 1708 | 13.9 | 1.27 |
| **366** | Xi'an Honghui Hospital | medical | China | 123 | - | 1164 | 9.5 | 0.64 |
| **367** | South China Normal University | academic | China | 122 | 220 | 1971 | 16.2 | 1.15 |
| **368** | North Sichuan Medical College | academic | China | 122 | 1050 | 879 | 7.2 | 0.64 |
| **369** | Ingham Institute | academic | Australia | 122 | 1500 | 1372 | 11.2 | 1.39 |
| **370** | Konyang University | academic | South Korea | 121 | 11.1 | 1801 | 14.9 | 0.93 |
| **371** | University of Miyazaki | academic | Japan | 120 | -10 | 1217 | 10.1 | 0.82 |
| **372** | Toranomon Hospital | medical | Japan | 120 | 62.5 | 1556 | 13 | 0.85 |
| **373** | Chiang Mai University | academic | Thailand | 120 | 211.1 | 1447 | 12.1 | 0.78 |
| **374** | Royal Melbourne Institute of Technology University | academic | Australia | 119 | 183.3 | 4241 | 35.6 | 2.49 |
| **375** | National Health Research Institutes Taiwan | government | Taiwan | 118 | 33.3 | 1935 | 16.4 | 1.19 |
| **376** | Tata Memorial Hospital | medical | India | 116 | 100 | 1089 | 9.4 | 1.03 |
| **377** | Korea Advanced Institute of Science and Technology | academic | South Korea | 116 | 333.3 | 2124 | 18.3 | 1.69 |
| **378** | Japan Science and Technology Agency | government | Japan | 115 | -96.3 | 3690 | 32.1 | 1.45 |
| **379** | All India Institute of Medical Sciences, Bhubaneswar | academic | India | 115 | 833.3 | 516 | 4.5 | 0.52 |
| **380** | Sir Ganga Ram Hospital | medical | India | 114 | 37.5 | 785 | 6.9 | 0.47 |
| **381** | Universiti Sains Malaysia | academic | Malaysia | 114 | 30 | 5876 | 51.5 | 4.42 |
| **382** | National Health Insurance Corporation Ilsan Hospital | medical | South Korea | 114 | 122.2 | 1118 | 9.8 | 0.92 |
| **383** | Chungbuk National University | academic | South Korea | 113 | -18.2 | 1368 | 12.1 | 0.63 |
| **384** | Dayanand Medical College & Hospital | medical | India | 112 | 150 | 1348 | 12 | 0.72 |
| **385** | Kosin University | academic | South Korea | 112 | 50 | 1738 | 15.5 | 1.04 |
| **386** | Thammasat University | academic | Thailand | 111 | 62.5 | 1015 | 9.1 | 0.59 |
| **387** | Japanese Red Cross Medical Center | medical | Japan | 111 | 72.7 | 3040 | 27.4 | 1.79 |
| **388** | East China Normal University | academic | China | 110 | 63.6 | 2231 | 20.3 | 1.3 |
| **389** | Capital & Coast District Health Board | government | New Zealand | 110 | 200 | 5451 | 49.6 | 3.42 |
| **390** | National Center for Global Health and Medicine | government | Japan | 109 | 40 | 1632 | 15 | 1.07 |
| **391** | Institute of Post Graduate Medical Education and Research Kolkatta | academic | India | 109 | 200 | 845 | 7.8 | 0.94 |
| **392** | Queen Elizabeth II Medical Centre Trust | medical | Australia | 109 | -36.4 | 2398 | 22 | 1.4 |
| **393** | National Hospital Organization Utano National Hospital | medical | Japan | 109 | -33.3 | 1423 | 13.1 | 1.51 |
| **394** | Walter and Eliza Hall Institute of Medical Research | academic | Australia | 108 | 650 | 1787 | 16.5 | 2.08 |
| **395** | Hangzhou Medical College | academic | China | 108 | 2600 | 697 | 6.5 | 0.7 |
| **396** | Hudson Institute of Medical Research | medical | Australia | 107 | -12.5 | 2985 | 27.9 | 1.35 |
| **397** | Prince of Songkla University | academic | Thailand | 107 | 70 | 810 | 7.6 | 0.67 |
| **398** | Barwon Health | medical | Australia | 107 | 166.7 | 2897 | 27.1 | 2.3 |
| **399** | Kuala Lumpur Hospital | medical | Malaysia | 106 | 83.3 | 957 | 9 | 0.74 |
| **400** | Western Health | medical | Australia | 106 | 1900 | 2457 | 23.2 | 1.88 |
| **401** | National Hospital Organization Murayama Medical Center | medical | Japan | 106 | 500 | 2160 | 20.4 | 1.18 |
| **402** | Kanagawa Children's Medical Center | medical | Japan | 105 | -36.4 | 1930 | 18.4 | 1.15 |
| **403** | KK Women's and Children's Hospital | medical | Singapore | 105 | 325 | 1032 | 9.8 | 0.85 |
| **404** | Kwandong University | academic | South Korea | 105 | -30.8 | 1402 | 13.4 | 0.84 |
| **405** | Raffles Hospital | medical | Singapore | 105 | 333.3 | 11158 | 106.3 | 7.93 |
| **406** | Akita Cerebrospinal and Cardiovascular Center | medical | Japan | 105 | -42.9 | 2528 | 24.1 | 1.59 |
| **407** | National Hospital Organization Hokkaido Medical Center | medical | Japan | 104 | 57.1 | 1232 | 11.8 | 0.8 |
| **408** | Liaocheng People's Hospital | medical | China | 103 | 220 | 777 | 7.5 | 1 |
| **409** | Guangdong Medical College | academic | China | 101 | 171.4 | 1452 | 14.4 | 1.15 |
| **410** | National Institutes of Natural Sciences - National Institute for Physiological Sciences | academic | Japan | 101 | -55 | 1953 | 19.3 | 1.06 |
| **411** | Jaslok Hospital and Research Centre | medical | India | 101 | 1400 | 698 | 6.9 | 0.77 |
| **412** | National Sun Yat-sen University | academic | Taiwan | 100 | -35.3 | 1245 | 12.4 | 0.81 |
| **413** | Tianjin First Central Hospital | medical | China | 100 | 900 | 1121 | 11.2 | 0.96 |
| **414** | St. George Hospital | medical | Australia | 99 | 22.2 | 2886 | 29.2 | 1.52 |
| **415** | Mackay Memorial Hospital Taiwan | medical | Taiwan | 99 | -33.3 | 1082 | 10.9 | 0.77 |
| **416** | National Development and Reform Commission of China | government | China | 99 | 4500 | 1169 | 11.8 | 1.48 |
| **417** | Hubei University of Medicine | academic | China | 98 | 400 | 1639 | 16.7 | 1.11 |
| **418** | Far Eastern Memorial Hospital | medical | Taiwan | 98 | 33.3 | 1308 | 13.3 | 0.72 |
| **419** | P.D. Hinduja National Hospital and Medical Research Centre | medical | India | 98 | 375 | 487 | 5 | 0.54 |
| **420** | Seirei Mikatahara General Hospital | medical | Japan | 98 | 28.6 | 1349 | 13.8 | 1.17 |
| **421** | Kameda Medical Center | medical | Japan | 96 | -33.3 | 952 | 9.9 | 0.6 |
| **422** | Aomori Prefectural Central Hospital | medical | Japan | 96 | -83.3 | 947 | 9.9 | 0.81 |
| **423** | Beihang University | academic | China | 95 | 675 | 828 | 8.7 | 1.18 |
| **424** | Universitas Airlangga | academic | Indonesia | 95 | - | 1453 | 15.3 | 2.66 |
| **425** | Linyi People's Hospital | medical | China | 95 | - | 1457 | 15.3 | 1.7 |
| **426** | All India Institute of Medical Sciences, Bhopal | academic | India | 95 | 400 | 353 | 3.7 | 0.53 |
| **427** | Vardhman Mahavir Medical College & Safdarjung Hospital | medical | India | 94 | 2500 | 494 | 5.3 | 0.78 |
| **428** | Waseda University | academic | Japan | 94 | 62.5 | 1063 | 11.3 | 0.93 |
| **429** | National Hospital Organization Nishi-Niigata Chuo National Hospital | medical | Japan | 94 | -16.7 | 1542 | 16.4 | 0.92 |
| **430** | ORYGEN Youth Health | academic | Australia | 93 | 966.7 | 2277 | 24.5 | 3.46 |
| **431** | Tazuke Kofukai, Medical Research Institute, Kitano Hospital | medical | Japan | 92 | 116.7 | 1420 | 15.4 | 1.19 |
| **432** | University of Canterbury | academic | New Zealand | 92 | -14.3 | 1795 | 19.5 | 1.3 |
| **433** | Shenzhen Institute of Advanced Technology | academic | China | 92 | 666.7 | 2201 | 23.9 | 3.36 |
| **434** | St. John's National Academy of Health Sciences | academic | India | 92 | 216.7 | 2139 | 23.3 | 1.52 |
| **435** | Yantai Yuhuangding Hospital | medical | China | 91 | 500 | 1412 | 15.5 | 1.38 |
| **436** | National Hospital Organization Yonezawa National Hospital | medical | Japan | 91 | 200 | 1899 | 20.9 | 2.07 |
| **437** | Peter Maccallum Cancer Centre | medical | Australia | 90 | 333.3 | 1296 | 14.4 | 1.31 |
| **438** | Daejin Medical Center | medical | South Korea | 90 | 333.3 | 966 | 10.7 | 0.86 |
| **439** | St Vincent's Clinic | medical | Australia | 90 | 166.7 | 5078 | 56.4 | 2.57 |
| **440** | Medical College, Thiruvananthapuram | medical | India | 90 | 63.6 | 400 | 4.4 | 0.39 |
| **441** | Kurashiki Central Hospital | medical | Japan | 89 | 450 | 737 | 8.3 | 0.62 |
| **442** | Krishna Institute of Medical Sciences | academic | India | 89 | 300 | 553 | 6.2 | 0.45 |
| **443** | Saitama Children's Medical Center | medical | Japan | 89 | 366.7 | 823 | 9.2 | 0.69 |
| **444** | Asia University Taiwan | academic | Taiwan | 88 | 1200 | 898 | 10.2 | 0.84 |
| **445** | Veterans General Hospital-Kaohsiung Taiwan | medical | Taiwan | 88 | 375 | 680 | 7.7 | 0.8 |
| **446** | Cogstate Ltd. | corporate | Australia | 88 | 37.5 | 4666 | 53 | 2.79 |
| **447** | Guangzhou First People's Hospital | medical | China | 87 | 400 | 1249 | 14.4 | 0.99 |
| **448** | Women's and Children's Hospital Adelaide | medical | Australia | 86 | 55.6 | 2022 | 23.5 | 1.63 |
| **449** | Binzhou Medical College | academic | China | 86 | 466.7 | 1806 | 21 | 2.83 |
| **450** | Canberra Hospital | medical | Australia | 85 | -9.1 | 959 | 11.3 | 0.79 |
| **451** | Japan Organization of Occupational Health and Safety | medical | Japan | 85 | 37.5 | 1090 | 12.8 | 0.98 |
| **452** | St. Luke's International Hospital | medical | Japan | 84 | 60 | 673 | 8 | 0.59 |
| **453** | Weifang Medical University | academic | China | 84 | 55.6 | 1311 | 15.6 | 0.93 |
| **454** | National Hospital Organization Osaka Toneyama Medical Center | medical | Japan | 84 | -44.4 | 944 | 11.2 | 0.77 |
| **455** | Tianjin University | academic | China | 83 | - | 610 | 7.3 | 0.77 |
| **456** | G.B. Pant Hospital India | medical | India | 83 | -90.6 | 851 | 10.3 | 0.49 |
| **457** | Tokyo Metropolitan University | academic | Japan | 83 | 80 | 975 | 11.7 | 0.92 |
| **458** | Institute for Basic Science | government | South Korea | 83 | 366.7 | 1541 | 18.6 | 1.43 |
| **459** | Sri Ramachandra Institute of Higher Education and Research | academic | India | 83 | 2200 | 332 | 4 | 0.31 |
| **460** | Southern Tohoku General Hospital | medical | Japan | 83 | 33.3 | 5265 | 63.4 | 2.99 |
| **461** | Hiroshima City Asa Citizens Hospital | medical | Japan | 82 | 650 | 716 | 8.7 | 1.01 |
| **462** | Fujian University of Traditional Chinese Medicine | academic | China | 81 | 1200 | 989 | 12.2 | 1.14 |
| **463** | Beijing Jishuitan Hospital | medical | China | 81 | 60 | 1150 | 14.2 | 1.04 |
| **464** | Chengdu Medical College | academic | China | 80 | 1000 | 627 | 7.8 | 0.74 |
| **465** | North China University of Science and Technology | academic | China | 80 | - | 931 | 11.6 | 0.84 |
| **466** | Japanese Red Cross Nagoya Daini Hospital | medical | Japan | 80 | -30 | 807 | 10.1 | 0.66 |
| **467** | Beijing Institute of Technology | academic | China | 79 | 225 | 973 | 12.3 | 1.1 |
| **468** | Universiti Putra Malaysia | academic | Malaysia | 79 | 1800 | 886 | 11.2 | 1.21 |
| **469** | University of Science and Technology UST | academic | South Korea | 79 | 500 | 1298 | 16.4 | 0.98 |
| **470** | Sumitomo Dainippon Pharma Co., Ltd. | corporate | Japan | 79 | 166.7 | 1265 | 16 | 1.03 |
| **471** | Cheng Hsin General Hospital | medical | Taiwan | 79 | 0 | 791 | 10 | 0.84 |
| **472** | Garvan Institute of Medical Research | medical | Australia | 78 | 750 | 2091 | 26.8 | 1.98 |
| **473** | Advanced Telecommunications Research Institute International | corporate | Japan | 78 | 450 | 1614 | 20.7 | 1 |
| **474** | St. Luke's Medical Center Quezon City | medical | Philippines | 78 | 900 | 784 | 10.1 | 1.29 |
| **475** | Liaquat National Hospital | medical | Pakistan | 78 | 350 | 926 | 11.9 | 1.02 |
| **476** | University of Indonesia | academic | Indonesia | 77 | 1200 | 1638 | 21.3 | 3.23 |
| **477** | Tianjin University of Traditional Chinese Medicine | academic | China | 77 | 366.7 | 853 | 11.1 | 0.97 |
| **478** | Westmead Millennium Institute for Medical Research | academic | Australia | 77 | 125 | 1814 | 23.6 | 1.6 |
| **479** | University of Santo Tomas | academic | Philippines | 77 | 150 | 1200 | 15.6 | 1.11 |
| **480** | Hebei General Hospital | medical | China | 77 | 850 | 1171 | 15.2 | 1 |
| **481** | Fukuoka Children's Hospital | medical | Japan | 77 | 50 | 1006 | 13.1 | 1.14 |
| **482** | Cathay General Hospital Taiwan | medical | Taiwan | 76 | -77.8 | 1085 | 14.3 | 0.8 |
| **483** | National Ageing Research Institute | academic | Australia | 76 | -14.3 | 4209 | 55.4 | 2.41 |
| **484** | Dr. D. Y. Patil Vidyapeeth, Pune | academic | India | 76 | 800 | 3564 | 46.9 | 3.78 |
| **485** | Sher-I-Kashmir Institute of Medical Sciences | academic | India | 76 | -50 | 425 | 5.6 | 0.69 |
| **486** | Royal Victorian Eye & Ear Hospital, Melbourne | medical | Australia | 75 | 140 | 1651 | 22 | 1.51 |
| **487** | PathWest Laboratory Medicine WA | academic | Australia | 75 | 160 | 1732 | 23.1 | 1.59 |
| **488** | Tasmanian Government | government | Australia | 73 | 400 | 878 | 12 | 1.02 |
| **489** | Fujian Provincial Hospital | medical | China | 73 | 400 | 941 | 12.9 | 0.86 |
| **490** | Royal Hobart Hospital | medical | Australia | 73 | 400 | 878 | 12 | 1.02 |
| **491** | Ministry of Health and Family Welfare | government | India | 73 | 450 | 220 | 3 | 0.4 |
| **492** | Tokyo Metropolitan Tama Medical Center | medical | Japan | 73 | 175 | 416 | 5.7 | 0.67 |
| **493** | Singapore National Eye Center | medical | Singapore | 72 | 12.5 | 1872 | 26 | 1.81 |
| **494** | Fortis Healthcare | corporate | India | 72 | 900 | 316 | 4.4 | 0.46 |
| **495** | Kokura Memorial Hospital | medical | Japan | 72 | - | 552 | 7.7 | 0.63 |
| **496** | Hunan Normal University | academic | China | 71 | 1200 | 779 | 11 | 1.02 |
| **497** | National Chung Hsing University | academic | Taiwan | 71 | 100 | 735 | 10.4 | 0.75 |
| **498** | Shin Kong Wu Ho-Su Memorial Hospital | medical | Taiwan | 71 | -33.3 | 818 | 11.5 | 0.91 |
| **499** | Jinzhou Medical University | academic | China | 71 | 57.1 | 1100 | 15.5 | 1 |
| **500** | Ningbo University | academic | China | 70 | 120 | 931 | 13.3 | 0.9 |
| **501** | Indian Council of Medical Research | government | India | 70 | 125 | 1546 | 22.1 | 3.16 |
| **502** | Mater Group | medical | Australia | 70 | 250 | 1163 | 16.6 | 1.23 |
| **503** | CAS - Shanghai Institute of Nutrition and Health | academic | China | 70 | -10 | 2174 | 31.1 | 1.58 |
| **504** | National Hospital Organization Kyoto Medical Center | medical | Japan | 70 | - | 718 | 10.3 | 0.83 |
| **505** | Chubu Rosai Hospital | medical | Japan | 70 | -57.1 | 1162 | 16.6 | 0.94 |
| **506** | China Pharmaceutical University | academic | China | 69 | 466.7 | 1115 | 16.2 | 1.07 |
| **507** | CAS - Institute of Biophysics | academic | China | 69 | 150 | 1041 | 15.1 | 1.26 |
| **508** | Central Queensland University | academic | Australia | 68 | 300 | 1511 | 22.2 | 1.45 |
| **509** | Changi General Hospital | medical | Singapore | 68 | -37.5 | 727 | 10.7 | 0.84 |
| **510** | National Central University | academic | Taiwan | 68 | -62.5 | 2200 | 32.4 | 1.38 |
| **511** | Guangxi University of Chinese Medicine | academic | China | 68 | 300 | 727 | 10.7 | 0.81 |
| **512** | Osaka Rosai Hospital | medical | Japan | 68 | 150 | 1228 | 18.1 | 1.2 |
| **513** | Baker Heart Research Institute | medical | Australia | 67 | 80 | 2486 | 37.1 | 4.21 |
| **514** | Queen Elizabeth Hospital Hong Kong | medical | Hong Kong | 67 | 50 | 1213 | 18.1 | 1.15 |
| **515** | Northern Health | medical | Australia | 67 | 75 | 1769 | 26.4 | 1.56 |
| **516** | Bengbu Medical College | academic | China | 67 | - | 658 | 9.8 | 0.89 |
| **517** | Sanno Hospital | medical | Japan | 67 | - | 904 | 13.5 | 1.05 |
| **518** | Foundation for Biomedical Research and Innovation at Kobe | other | Japan | 67 | -42.9 | 965 | 14.4 | 1.12 |
| **519** | Osaka Women's and Children's Hospital | medical | Japan | 67 | -44.4 | 1142 | 17 | 1.1 |
| **520** | Saiseikai Kumamoto Hospital | medical | Japan | 67 | 0 | 356 | 5.3 | 0.42 |
| **521** | Takeda Pharmaceutical Company Limited | corporate | Japan | 66 | 333.3 | 1004 | 15.2 | 1.77 |
| **522** | CSIR - Institute of Genomics and Integrative Biology | government | India | 66 | 650 | 809 | 12.3 | 0.68 |
| **523** | Anjo Kosei Hospital | medical | Japan | 65 | 0 | 518 | 8 | 0.62 |
| **524** | Tokyo Metropolitan Bokutoh Hospital | medical | Japan | 65 | -41.7 | 883 | 13.6 | 0.81 |
| **525** | Academy of Military Medical Science China | government | China | 64 | -33.3 | 970 | 15.2 | 0.97 |
| **526** | Astellas Pharma Inc. | corporate | Japan | 64 | -16.7 | 2336 | 36.5 | 1.87 |
| **527** | Shenzhen Children's Hospital | medical | China | 64 | 633.3 | 600 | 9.4 | 0.89 |
| **528** | University of the Sunshine Coast | academic | Australia | 63 | - | 834 | 13.2 | 2.25 |
| **529** | Tokyo Metropolitan Komagome Hospital | medical | Japan | 63 | -12.5 | 795 | 12.6 | 1.06 |
| **530** | Padjadjaran University | academic | Indonesia | 63 | 400 | 847 | 13.4 | 1.17 |
| **531** | National Hospital Organization Nagoya Medical Center | medical | Japan | 63 | 25 | 1141 | 18.1 | 1.04 |
| **532** | Yokohama Rosai Hospital | medical | Japan | 63 | 350 | 781 | 12.4 | 0.98 |
| **533** | Royal Women's Hospital | medical | Australia | 62 | 50 | 1463 | 23.6 | 1.4 |
| **534** | National Institute of Infectious Diseases | government | Japan | 62 | -42.9 | 5945 | 95.9 | 6.54 |
| **535** | Henan University of Chinese Medicine | academic | China | 62 | - | 846 | 13.6 | 1.19 |
| **536** | Xi'an Medical University | academic | China | 62 | 300 | 497 | 8 | 0.64 |
| **537** | Japan Community Healthcare Organization | medical | Japan | 62 | - | 746 | 12 | 0.95 |
| **538** | Shanghai University of Sport | academic | China | 61 | 1200 | 601 | 9.9 | 0.98 |
| **539** | All India Institute of Medical Sciences, Raipur | academic | India | 61 | 1500 | 140 | 2.3 | 0.49 |
| **540** | Fukuoka International University of Health and Welfare | academic | Japan | 61 | - | 536 | 8.8 | 0.71 |
| **541** | National Hospital Organization Nagasaki Medical Center | medical | Japan | 61 | -9.1 | 1319 | 21.6 | 0.79 |
| **542** | Dow University of Health Sciences | academic | Pakistan | 60 | 800 | 4165 | 69.4 | 7.32 |
| **543** | Grant Medical College | medical | India | 60 | 0 | 409 | 6.8 | 0.77 |
| **544** | Guru Teg Bahadur Hospital | medical | India | 60 | 166.7 | 258 | 4.3 | 0.47 |
| **545** | National Hospital Organization Nagasaki Kawatana Medical Center | medical | Japan | 60 | -70 | 790 | 13.2 | 0.96 |
| **546** | Kushiro Rosai Hospital | medical | Japan | 60 | -40 | 689 | 11.5 | 0.87 |
| **547** | Maulana Azad Medical College | academic | India | 59 | -33.3 | 396 | 6.7 | 0.45 |
| **548** | Chia Nan University of Pharmacy and Science | academic | Taiwan | 59 | -71.4 | 730 | 12.4 | 1.23 |
| **549** | Universitas Udayana | academic | Indonesia | 59 | - | 146 | 2.5 | 0.37 |
| **550** | Chia-Yi Christian Hospital | medical | Taiwan | 59 | -22.2 | 427 | 7.2 | 0.53 |
| **551** | Tokyo Metropolitan Children's Medical Center | medical | Japan | 59 | 100 | 394 | 6.7 | 0.55 |
| **552** | Medical College and Hospital Kolkata | medical | India | 58 | -28.6 | 231 | 4 | 0.31 |
| **553** | University of Colombo | academic | Sri Lanka | 58 | 25 | 1769 | 30.5 | 4.12 |
| **554** | Korea Institute of Oriental Medicine | government | South Korea | 58 | -75 | 1617 | 27.9 | 1.57 |
| **555** | Singapore Institute of Mental Health | medical | Singapore | 58 | 75 | 1229 | 21.2 | 2.8 |
| **556** | Monash University Malaysia | academic | Malaysia | 58 | - | 929 | 16 | 1.8 |
| **557** | Eastern Chiba Medical Center | medical | Japan | 58 | - | 652 | 11.2 | 0.99 |
| **558** | Osaka General Medical Center | medical | Japan | 58 | 33.3 | 568 | 9.8 | 1.01 |
| **559** | Henan University | academic | China | 57 | 500 | 715 | 12.5 | 0.89 |
| **560** | University of the Ryukyus | academic | Japan | 57 | 33.3 | 587 | 10.3 | 0.73 |
| **561** | National Taipei University of Nursing and Health Sciences | academic | Taiwan | 57 | 500 | 831 | 14.6 | 1.14 |
| **562** | KLE Academy of Higher Education and Research, Belagavi | academic | India | 57 | 100 | 329 | 5.8 | 0.63 |
| **563** | Hyogo Prefectural Kobe Children's Hospital | medical | Japan | 57 | - | 413 | 7.2 | 0.66 |
| **564** | Chinese Center for Disease Control and Prevention | government | China | 56 | 1600 | 3099 | 55.3 | 4.69 |
| **565** | Hong Kong Baptist University | academic | Hong Kong | 56 | 350 | 836 | 14.9 | 1.04 |
| **566** | Alexandra Hospital | medical | Singapore | 56 | -50 | 1622 | 29 | 1.75 |
| **567** | Spinal Injuries Center | medical | Japan | 56 | 60 | 729 | 13 | 0.88 |
| **568** | Northwestern Polytechnical University Xian | academic | China | 55 | 700 | 911 | 16.6 | 1.14 |
| **569** | Shaanxi Normal University | academic | China | 55 | 200 | 941 | 17.1 | 1.23 |
| **570** | Hitachi, Ltd. | corporate | Japan | 55 | 0 | 747 | 13.6 | 0.85 |
| **571** | Thailand Ministry of Public Health | government | Thailand | 55 | 175 | 470 | 8.5 | 0.81 |
| **572** | Xuzhou Central Hospital | medical | China | 55 | 800 | 1124 | 20.4 | 1.8 |
| **573** | Miyagi Children's Hospital | medical | Japan | 55 | 200 | 522 | 9.5 | 0.77 |
| **574** | Hong Kong University of Science and Technology | academic | Hong Kong | 54 | 500 | 4780 | 88.5 | 3.55 |
| **575** | Black Dog Institute | academic | Australia | 54 | 150 | 3037 | 56.2 | 2.56 |
| **576** | Shanghai University of Medicine and Health Sciences | academic | China | 54 | 125 | 1006 | 18.6 | 1.16 |
| **577** | Qinghai University | academic | China | 54 | - | 224 | 4.1 | 0.57 |
| **578** | Guangdong Second Provincial General Hospital | medical | China | 54 | 700 | 1112 | 20.6 | 1.71 |
| **579** | Kalawati Saran Children's Hospital | medical | India | 54 | 25 | 652 | 12.1 | 0.84 |
| **580** | Wakayama Rosai Hospital | medical | Japan | 54 | -63.6 | 769 | 14.2 | 1.22 |
| **581** | Princess Margaret Hospital for Children | medical | Australia | 53 | -75 | 1619 | 30.5 | 1.97 |
| **582** | Nanjing General Hospital | medical | China | 53 | 100 | 413 | 7.8 | 0.52 |
| **583** | Indraprastha Apollo Hospitals | medical | India | 53 | 0 | 426 | 8 | 0.74 |
| **584** | Bangabandhu Sheikh Mujib Medical University | academic | Bangladesh | 53 | 400 | 325 | 6.1 | 1.13 |
| **585** | Jamia Hamdard University | academic | India | 53 | 0 | 934 | 17.6 | 1.74 |
| **586** | Fuzhou General Hospital of Nanjing Military Command | medical | China | 53 | -75 | 1119 | 21.1 | 1.41 |
| **587** | Mitsui Memorial Hospital | medical | Japan | 53 | 100 | 588 | 11.1 | 0.84 |
| **588** | Fiona Stanley Hospital | medical | Australia | 53 | - | 748 | 14.1 | 1.53 |
| **589** | Aizawa Hospital | medical | Japan | 53 | 120 | 461 | 8.7 | 0.67 |
| **590** | Tribhuvan University | academic | Nepal | 52 | - | 3854 | 74.1 | 7.58 |
| **591** | Anhui University of Chinese Medicine | academic | China | 52 | 1500 | 486 | 9.3 | 0.92 |
| **592** | Southern University of Science and Technology | academic | China | 52 | 1500 | 1468 | 28.2 | 2.37 |
| **593** | University of Medicine and Pharmacy at Ho Chi Minh City | academic | Viet Nam | 52 | - | 369 | 7.1 | 0.9 |
| **594** | Central Coast Local Health District | government | Australia | 52 | 100 | 667 | 12.8 | 1.44 |
| **595** | Kyushu Rosai Hospital | medical | Japan | 52 | -25 | 463 | 8.9 | 0.55 |
| **596** | Government Medical College & Hospital Chandigarh | medical | India | 51 | 233.3 | 285 | 5.6 | 0.37 |
| **597** | Chengdu University | academic | China | 51 | 600 | 263 | 5.2 | 0.54 |
| **598** | Peninsula Health | medical | Australia | 51 | 500 | 872 | 17.1 | 1.74 |
| **599** | Shandong University of Traditional Chinese Medicine | academic | China | 51 | 375 | 471 | 9.2 | 0.79 |
| **600** | JSS Academy of Higher Education & Research | academic | India | 51 | 1200 | 2881 | 56.5 | 4.18 |
| **601** | Capital Institute of Pediatrics | medical | China | 51 | 400 | 327 | 6.4 | 0.53 |
| **602** | Tsukuba Medical Center Hospital | medical | Japan | 51 | 100 | 516 | 10.1 | 0.65 |
| **603** | Komaki City Hospital | medical | Japan | 51 | 125 | 730 | 14.3 | 1.15 |
| **604** | Japanese Red Cross Kyoto Daini Hospital | medical | Japan | 51 | 40 | 411 | 8.1 | 0.98 |
| **605** | Saiseikai Central Hospital | medical | Japan | 51 | -33.3 | 1305 | 25.6 | 1.76 |
| **606** | The Graduate University for Advanced Studies | academic | Japan | 50 | -66.7 | 1526 | 30.5 | 1.26 |
| **607** | Niigata University of Health and Welfare | academic | Japan | 50 | 50 | 503 | 10.1 | 0.95 |
| **608** | Cabrini Health | medical | Australia | 50 | 266.7 | 2051 | 41 | 1.97 |
| **609** | Chang Gung University of Science and Technology | academic | Taiwan | 50 | -66.7 | 805 | 16.1 | 0.87 |
| **610** | Sumandeep Vidyapeeth University | academic | India | 50 | 500 | 302 | 6 | 0.49 |
| **611** | Victoria University | academic | Australia | 49 | -20 | 1090 | 22.2 | 0.97 |
| **612** | Beijing University of Technology | academic | China | 49 | -50 | 720 | 14.7 | 0.71 |
| **613** | Guangdong Pharmaceutical University | academic | China | 49 | 233.3 | 488 | 10 | 0.83 |
| **614** | Kwong Wah Hospital | medical | Hong Kong | 49 | 700 | 519 | 10.6 | 0.75 |
| **615** | Otsuka Pharmaceutical Co Ltd. | corporate | Japan | 49 | -16.7 | 1107 | 22.6 | 1.49 |
| **616** | Waikato Hospital | medical | New Zealand | 49 | 1200 | 652 | 13.3 | 1.55 |
| **617** | Japan National Institute of Information and Communications Technology | government | Japan | 48 | 600 | 1027 | 21.4 | 0.87 |
| **618** | Tenri Hospital | medical | Japan | 48 | 166.7 | 472 | 9.8 | 0.77 |
| **619** | Medanta (The Medicity) | academic | India | 48 | - | 153 | 3.2 | 0.3 |
| **620** | General Hospital of Chinese People's Armed Police Forces | medical | China | 48 | -100 | 952 | 19.8 | 0.81 |
| **621** | R.G. Kar Medical College and Hospital | medical | India | 48 | 350 | 240 | 5 | 0.52 |
| **622** | Cangzhou Central Hospital | medical | China | 48 | 350 | 833 | 17.4 | 1.41 |
| **623** | China Three Gorges University | academic | China | 47 | 400 | 533 | 11.3 | 0.98 |
| **624** | Massey University | academic | New Zealand | 47 | -16.7 | 995 | 21.2 | 1.36 |
| **625** | Victoria University of Wellington | academic | New Zealand | 47 | 0 | 518 | 11 | 0.86 |
| **626** | Hungkuang University | academic | Taiwan | 47 | -25 | 502 | 10.7 | 0.65 |
| **627** | National Taiwan Normal University | academic | Taiwan | 47 | -16.7 | 891 | 19 | 0.99 |
| **628** | NITTE | academic | India | 47 | 75 | 863 | 18.4 | 1.42 |
| **629** | Cardinal Tien Hospital, Taiwan | medical | Taiwan | 47 | -37.5 | 1111 | 23.6 | 0.99 |
| **630** | PLA General Hospital | medical | China | 47 | 166.7 | 437 | 9.3 | 0.98 |
| **631** | Chiba Children's Hospital | medical | Japan | 47 | 100 | 688 | 14.6 | 1.06 |
| **632** | Zhejiang Hospital | medical | China | 47 | - | 251 | 5.3 | 0.83 |
| **633** | Peerless Hospital | medical | India | 47 | -70 | 151 | 3.2 | 0.41 |
| **634** | Indira Gandhi Medical College | medical | India | 47 | -66.7 | 222 | 4.7 | 0.41 |
| **635** | Osaka International Cancer Institute | medical | Japan | 47 | -33.3 | 848 | 18 | 1.46 |
| **636** | Hiroshima City Hiroshima Citizens Hospital | medical | Japan | 47 | 350 | 374 | 8 | 0.61 |
| **637** | University of Canberra | academic | Australia | 46 | - | 589 | 12.8 | 1.18 |
| **638** | Shenzhen People's Hospital | medical | China | 46 | 900 | 378 | 8.2 | 0.88 |
| **639** | Aichi Gakuin University | academic | Japan | 46 | 0 | 915 | 19.9 | 0.83 |
| **640** | National Institute of Advanced Industrial Science and Technology | government | Japan | 46 | - | 450 | 9.8 | 0.75 |
| **641** | Rajasthan University of Health Sciences | academic | India | 46 | - | 3597 | 78.2 | 5.74 |
| **642** | National Hospital Organization Sagamihara National Hospital | medical | Japan | 46 | 250 | 1340 | 29.1 | 1.82 |
| **643** | Saiseikai Shiga Hospital | medical | Japan | 46 | - | 91 | 2 | 0.36 |
| **644** | Hebei University | academic | China | 45 | 450 | 564 | 12.5 | 1.24 |
| **645** | National Institute of Pharmaceutical Education and Research India | government | India | 45 | 266.7 | 782 | 17.4 | 1.21 |
| **646** | Pandit Bhagwat Dayal Sharma Postgraduate Institute of Medical Sciences | academic | India | 45 | 25 | 319 | 7.1 | 0.53 |
| **647** | National Cancer Centre | medical | Singapore | 45 | 175 | 826 | 18.4 | 2.5 |
| **648** | Hanoi Medical University | academic | Viet Nam | 45 | 700 | 8756 | 194.6 | 15.17 |
| **649** | Homi Bhabha National Institute | academic | India | 45 | - | 234 | 5.2 | 0.54 |
| **650** | Guilin Medical College | academic | China | 45 | 266.7 | 689 | 15.3 | 1.27 |
| **651** | PLA General Hospital of Jinan Military Region | medical | China | 45 | -100 | 535 | 11.9 | 0.83 |
| **652** | Dongguan People's Hospital | medical | China | 45 | 80 | 417 | 9.3 | 0.8 |
| **653** | Japanese Red Cross Shizuoka Hospital | medical | Japan | 45 | 125 | 497 | 11 | 1.15 |
| **654** | Henan University of Science and Technology | academic | China | 44 | 800 | 500 | 11.4 | 1.36 |
| **655** | Wuhan General Hospital | medical | China | 44 | -100 | 608 | 13.8 | 0.82 |
| **656** | Bharati Vidyapeeth University | academic | India | 44 | 500 | 190 | 4.3 | 0.57 |
| **657** | University of Waikato | academic | New Zealand | 44 | -100 | 1252 | 28.5 | 1.16 |
| **658** | Daegu Gyeongbuk Institute of Science and Technology | academic | South Korea | 44 | 250 | 721 | 16.4 | 0.91 |
| **659** | Kementerian Kesihatan Malaysia | government | Malaysia | 44 | 1100 | 269 | 6.1 | 0.8 |
| **660** | Hiroshima Prefectural Hospital | medical | Japan | 44 | 166.7 | 711 | 16.2 | 1.02 |
| **661** | Aso Iizuka Hospital | medical | Japan | 44 | 133.3 | 281 | 6.4 | 0.57 |
| **662** | Bond University | academic | Australia | 43 | - | 549 | 12.8 | 0.96 |
| **663** | Indian Institute of Science Bangalore | academic | India | 43 | 1100 | 646 | 15 | 1.2 |
| **664** | Armed Forces Medical College | academic | India | 43 | 366.7 | 342 | 8 | 0.6 |
| **665** | Korea Institute of Radiological and Medical Sciences | medical | South Korea | 43 | -33.3 | 988 | 23 | 1.55 |
| **666** | Navy General Hospital | medical | China | 43 | -100 | 606 | 14.1 | 0.7 |
| **667** | Nepean Hospital | medical | Australia | 43 | - | 477 | 11.1 | 0.98 |
| **668** | St. Mary's Hospital, Kurume | medical | Japan | 43 | 600 | 441 | 10.3 | 0.67 |
| **669** | National Hospital Organization Tokyo Medical Center | medical | Japan | 43 | 0 | 383 | 8.9 | 0.64 |
| **670** | National Hospital Organization Sendai Nishitaga National Hospital | medical | Japan | 43 | -80 | 768 | 17.9 | 0.96 |
| **671** | Panjab University | academic | India | 42 | 0 | 1733 | 41.3 | 4.89 |
| **672** | National Chengchi University | academic | Taiwan | 42 | 200 | 561 | 13.4 | 0.97 |
| **673** | National Tsing Hua University | academic | Taiwan | 42 | 16.7 | 539 | 12.8 | 1.01 |
| **674** | ANZAC Research Institute | academic | Australia | 42 | -20 | 1318 | 31.4 | 1.9 |
| **675** | Maharishi Markandeshwar University, Mullana | academic | India | 42 | -20 | 1111 | 26.5 | 2.68 |
| **676** | Northwest University China | academic | China | 42 | 600 | 310 | 7.4 | 0.66 |
| **677** | Guizhou Provincial People's Hospital | medical | China | 42 | 250 | 272 | 6.5 | 0.6 |
| **678** | K.S. Hegde Medical Academy | medical | India | 42 | -33.3 | 593 | 14.1 | 0.78 |
| **679** | Hyogo Prefectural Amagasaki General Medical Center | medical | Japan | 42 | - | 213 | 5.1 | 0.62 |
| **680** | Aichi Developmental Disability Center | medical | Japan | 42 | 200 | 426 | 10.1 | 0.69 |
| **681** | Japanese Red Cross Nagoya Daiichi Hospital | medical | Japan | 42 | -57.1 | 415 | 9.9 | 0.63 |
| **682** | Kansai Rosai Hospital | medical | Japan | 42 | -16.7 | 515 | 12.3 | 0.84 |
| **683** | Taishan Medical University | academic | China | 41 | -33.3 | 998 | 24.3 | 1.39 |
| **684** | Tuen Mun Hospital | medical | Hong Kong | 41 | -40 | 934 | 22.8 | 1.14 |
| **685** | Kobe Gakuin University | academic | Japan | 41 | 50 | 336 | 8.2 | 0.69 |
| **686** | Sydney Hospital and Sydney Eye Hospital | medical | Australia | 41 | - | 1431 | 34.9 | 5.65 |
| **687** | Cooperative Research Centres Australia | government | Australia | 41 | 100 | 1176 | 28.7 | 1.89 |
| **688** | Gannan Medical College | academic | China | 41 | - | 352 | 8.6 | 0.76 |
| **689** | Okazaki City Hospital | medical | Japan | 41 | 25 | 246 | 6 | 0.62 |
| **690** | Japanese Red Cross Asahikawa Hospital | medical | Japan | 41 | 0 | 346 | 8.4 | 0.73 |
| **691** | Charles Sturt University | academic | Australia | 40 | -71.4 | 718 | 18 | 1.06 |
| **692** | Jiaxing University | academic | China | 40 | 900 | 283 | 7.1 | 0.8 |
| **693** | Tokyo Dental College | academic | Japan | 40 | 175 | 374 | 9.4 | 1.02 |
| **694** | Universiti Teknologi MARA | academic | Malaysia | 40 | 400 | 339 | 8.5 | 1.63 |
| **695** | SRM University | academic | India | 40 | 700 | 177 | 4.4 | 0.55 |
| **696** | Inner Mongolia University of Science and Technology | academic | China | 40 | 350 | 845 | 21.1 | 1.61 |
| **697** | The Education University of Hong Kong | academic | Hong Kong | 40 | 16.7 | 702 | 17.5 | 1.19 |
| **698** | Academia Sinica - Institute of Biomedical Sciences | academic | Taiwan | 40 | 400 | 2524 | 63.1 | 4.62 |
| **699** | Seoul Metropolitan Government | government | South Korea | 40 | - | 792 | 19.8 | 1.42 |
| **700** | Tokyo University of Science | academic | Japan | 39 | -20 | 962 | 24.7 | 1.23 |
| **701** | Kio University | academic | Japan | 39 | - | 323 | 8.3 | 0.68 |
| **702** | Korea Brain Research Institute | government | South Korea | 39 | 900 | 804 | 20.6 | 1.42 |
| **703** | Gansu Province People's Hospital | medical | China | 39 | 550 | 493 | 12.6 | 1 |
| **704** | Ehime Prefectural Central Hospital | medical | Japan | 39 | 100 | 333 | 8.5 | 0.75 |
| **705** | Kariya Toyota General Hospital | medical | Japan | 39 | 250 | 387 | 9.9 | 0.81 |
| **706** | International Centre for Diarrhoeal Disease Research Bangladesh | medical | Bangladesh | 38 | 400 | 4404 | 115.9 | 11.02 |
| **707** | University of Calcutta | academic | India | 38 | 0 | 1282 | 33.7 | 5.52 |
| **708** | Osaka Police Hospital | medical | Japan | 38 | 250 | 207 | 5.4 | 0.57 |
| **709** | Korea Research Institute of Bioscience and Biotechnology | government | South Korea | 38 | 200 | 612 | 16.1 | 0.94 |
| **710** | Soongsil University | academic | South Korea | 38 | - | 338 | 8.9 | 1.58 |
| **711** | University of Karachi | academic | Pakistan | 38 | 0 | 905 | 23.8 | 1.01 |
| **712** | National Brain Research Centre | academic | India | 38 | 50 | 351 | 9.2 | 0.65 |
| **713** | PLA No. 306 Hospital | medical | China | 38 | 0 | 497 | 13.1 | 1.11 |
| **714** | Sichuan Cancer Hospital and Institute | medical | China | 38 | 600 | 276 | 7.3 | 0.81 |
| **715** | Kawasaki Municipal Hospital | medical | Japan | 38 | 50 | 317 | 8.3 | 0.6 |
| **716** | National Hospital Organization Mito Medical Center | medical | Japan | 38 | 25 | 529 | 13.9 | 0.88 |
| **717** | National Hospital Organization Higashi Nagoya National Hospital | medical | Japan | 38 | 500 | 533 | 14 | 1.26 |
| **718** | Japanese Red Cross Musashino Hospital | medical | Japan | 38 | -42.9 | 351 | 9.2 | 0.9 |
| **719** | Japanese Red Cross Kyoto Daiichi Hospital | medical | Japan | 38 | 900 | 190 | 5 | 0.72 |
| **720** | Saiseikai Yokohamashi Tobu Hospital | medical | Japan | 38 | 250 | 257 | 6.8 | 0.65 |
| **721** | Kunming University of Science and Technology | academic | China | 37 | - | 425 | 11.5 | 1.2 |
| **722** | Repatriation General Hospital | medical | Australia | 37 | -100 | 1321 | 35.7 | 1.68 |
| **723** | Show-Chwan Memorial Hospital Taiwan | medical | Taiwan | 37 | 0 | 336 | 9.1 | 0.95 |
| **724** | Chettinad Health City | medical | India | 37 | 100 | 111 | 3 | 0.39 |
| **725** | Command Hospital Air Force | medical | India | 37 | 25 | 101 | 2.7 | 0.28 |
| **726** | Sendai City Hospital | medical | Japan | 37 | -100 | 434 | 11.7 | 0.67 |
| **727** | Japan Community Healthcare Organization Tokyo Shinjuku Medical Center | medical | Japan | 37 | 400 | 335 | 9.1 | 0.88 |
| **728** | Showa General Hospital | medical | Japan | 37 | 100 | 223 | 6 | 0.38 |
| **729** | Zhejiang Normal University | academic | China | 36 | - | 541 | 15 | 1.31 |
| **730** | Pamela Youde Nethersole Eastern Hospital | medical | Hong Kong | 36 | 400 | 828 | 23 | 1.64 |
| **731** | Naresuan University | academic | Thailand | 36 | 100 | 753 | 20.9 | 1.39 |
| **732** | Tauranga Hospital | medical | New Zealand | 36 | 300 | 552 | 15.3 | 1.5 |
| **733** | Yan'an University | academic | China | 36 | - | 271 | 7.5 | 0.96 |
| **734** | Kinghorn Cancer Centre | academic | Australia | 36 | - | 525 | 14.6 | 2.08 |
| **735** | Konan Kosei Hospital | medical | Japan | 36 | 50 | 427 | 11.9 | 0.73 |
| **736** | Kawasaki University of Medical Welfare | academic | Japan | 36 | -33.3 | 194 | 5.4 | 0.39 |
| **737** | National Hospital Organization Okayama Medical Center | medical | Japan | 36 | 100 | 232 | 6.4 | 0.51 |
| **738** | Japanese Red Cross Ashikaga Hospital | medical | Japan | 36 | -33.3 | 329 | 9.1 | 0.56 |
| **739** | Kanto Rosai Hospital | medical | Japan | 36 | 16.7 | 565 | 15.7 | 0.99 |
| **740** | Chiba Rosai Hospital | medical | Japan | 36 | 800 | 470 | 13.1 | 1.08 |
| **741** | Shin-Yurigaoka General Hospital | medical | Japan | 36 | 500 | 261 | 7.3 | 0.9 |
| **742** | Charles Darwin University | academic | Australia | 35 | 100 | 385 | 11 | 0.99 |
| **743** | Harbin Institute of Technology | academic | China | 35 | - | 456 | 13 | 1.92 |
| **744** | Doshisha University | academic | Japan | 35 | -20 | 468 | 13.4 | 0.86 |
| **745** | Gwangju Institute of Science and Technology | academic | South Korea | 35 | 25 | 507 | 14.5 | 0.88 |
| **746** | Himalayan Institute Hospital Trust | medical | India | 35 | -50 | 362 | 10.3 | 0.5 |
| **747** | Shaanxi University of Chinese Medicine | academic | China | 35 | - | 314 | 9 | 1.07 |
| **748** | En Chu Kong Hospital | medical | Taiwan | 35 | 0 | 588 | 16.8 | 1.2 |
| **749** | Ningbo First Hospital | medical | China | 35 | 1000 | 247 | 7.1 | 0.64 |
| **750** | Shizuoka Cancer Center | medical | Japan | 35 | 0 | 534 | 15.3 | 0.99 |
| **751** | Khoo Teck Puat Hospital | medical | Singapore | 35 | 200 | 592 | 16.9 | 1.13 |
| **752** | Tosei General Hospital | medical | Japan | 35 | 600 | 129 | 3.7 | 0.5 |
| **753** | Northeastern University China | academic | China | 34 | - | 1151 | 33.9 | 1.73 |
| **754** | Wuhan University of Science and Technology | academic | China | 34 | 600 | 1181 | 34.7 | 5.99 |
| **755** | Kongju National University | academic | South Korea | 34 | -50 | 464 | 13.6 | 1.09 |
| **756** | Amity University, Noida | academic | India | 34 | 300 | 707 | 20.8 | 1.44 |
| **757** | National Hospital of Sri Lanka | academic | Sri Lanka | 34 | -20 | 139 | 4.1 | 0.68 |
| **758** | Macau University of Science and Technology | academic | Macao | 34 | 100 | 633 | 18.6 | 1.02 |
| **759** | Charutar Arogya Mandal | medical | India | 34 | - | 122 | 3.6 | 0.49 |
| **760** | People's Hospital of Guangxi Zhuang Autonomous Region | medical | China | 34 | 800 | 213 | 6.3 | 0.74 |
| **761** | Asahi General Hospital | medical | Japan | 34 | 33.3 | 358 | 10.5 | 0.7 |
| **762** | Shonan Kamakura General Hospital | medical | Japan | 34 | -80 | 224 | 6.6 | 0.65 |
| **763** | First Teaching Hospital of Tianjin University of Traditional Chinese Medicine | medical | China | 34 | 350 | 374 | 11 | 1.1 |
| **764** | Indira Gandhi Institute of Child Health | medical | India | 34 | 75 | 111 | 3.3 | 0.41 |
| **765** | Toyohashi Municipal Hospital | medical | Japan | 34 | -66.7 | 690 | 20.3 | 0.91 |
| **766** | Weifang People's Hospital | medical | China | 34 | - | 248 | 7.3 | 0.59 |
| **767** | Tsuchiura Kyodo General Hospital | medical | Japan | 34 | 200 | 143 | 4.2 | 0.38 |
| **768** | Steel Memorial Yawata Hospital | medical | Japan | 34 | 200 | 346 | 10.2 | 0.83 |
| **769** | Japanese Red Cross Kumamoto Hospital | medical | Japan | 34 | 200 | 331 | 9.7 | 1.63 |
| **770** | Sri Venkateswara Institute of Medical Sciences | medical | India | 33 | 0 | 202 | 6.1 | 0.4 |
| **771** | Kuang Tien General Hospital | medical | Taiwan | 33 | 200 | 343 | 10.4 | 0.81 |
| **772** | Huilongguan Hospital | medical | China | 33 | - | 503 | 15.2 | 1.66 |
| **773** | National Rehabilitation Center for Persons with Disabilities | medical | Japan | 33 | -100 | 720 | 21.8 | 1.06 |
| **774** | Uonuma Kikan Hospital | medical | Japan | 33 | - | 238 | 7.2 | 0.87 |
| **775** | Neuropsychiatric Research Institute, Tokyo | other | Japan | 33 | -88.9 | 752 | 22.8 | 0.99 |
| **776** | National Hospital Organization Suzuka National Hospital | medical | Japan | 33 | 250 | 299 | 9.1 | 1.07 |
| **777** | Japanese Red Cross Osaka Hospital | medical | Japan | 33 | 75 | 449 | 13.6 | 1.49 |
| **778** | CAS - Kunming Institute of Zoology | academic | China | 32 | 400 | 584 | 18.3 | 1.28 |
| **779** | Ivane Javakhishvili Tbilisi State University | academic | Georgia | 32 | 50 | 624 | 19.5 | 1.78 |
| **780** | Gifu Pharmaceutical University | academic | Japan | 32 | -50 | 641 | 20 | 1.31 |
| **781** | Ritsumeikan University | academic | Japan | 32 | -33.3 | 257 | 8 | 0.79 |
| **782** | Daegu University | academic | South Korea | 32 | -100 | 986 | 30.8 | 1.55 |
| **783** | Korea Basic Science Institute | government | South Korea | 32 | 200 | 417 | 13 | 0.91 |
| **784** | Southern Taiwan University of Science and Technology | academic | Taiwan | 32 | -100 | 406 | 12.7 | 0.79 |
| **785** | Beijing Institute of Pharmacology and Toxicology | academic | China | 32 | 33.3 | 449 | 14 | 0.76 |
| **786** | Lokmanya Tilak Municipal General Hospital and Medical College | medical | India | 32 | 550 | 208 | 6.5 | 1.54 |
| **787** | Army Hospital Research and Referral | medical | India | 32 | 700 | 314 | 9.8 | 1.01 |
| **788** | Nagano Children's Hospital | medical | Japan | 32 | 166.7 | 703 | 22 | 1.24 |
| **789** | Rakuwakai Otowa Hospital | medical | Japan | 32 | 33.3 | 653 | 20.4 | 1.7 |
| **790** | Jiangnan University | academic | China | 31 | 800 | 410 | 13.2 | 1.21 |
| **791** | Nanjing University of Aeronautics and Astronautics | academic | China | 31 | 600 | 873 | 28.2 | 1.89 |
| **792** | City University of Hong Kong | academic | Hong Kong | 31 | - | 421 | 13.6 | 1.39 |
| **793** | Fukuoka Dental College | academic | Japan | 31 | 100 | 378 | 12.2 | 0.85 |
| **794** | Hunan University of Chinese Medicine | academic | China | 31 | - | 526 | 17 | 1.46 |
| **795** | Hubei University of Chinese Medicine | academic | China | 31 | - | 296 | 9.5 | 1.21 |
| **796** | Heilongjiang University of Traditional Chinese Medicine | academic | China | 31 | 300 | 351 | 11.3 | 1.68 |
| **797** | Government of Western Australia, Department of Health | government | Australia | 31 | - | 982 | 31.7 | 2.23 |
| **798** | Tungs' Taichung MetroHarbor Hospital | medical | Taiwan | 31 | 0 | 322 | 10.4 | 0.7 |
| **799** | Sin-Lau Christian Hospital, Taiwan | medical | Taiwan | 31 | -40 | 261 | 8.4 | 0.95 |
| **800** | National University of Medical Sciences | academic | Pakistan | 31 | 500 | 1166 | 37.6 | 7.13 |
| **801** | Southern District Health Board | government | New Zealand | 31 | -25 | 425 | 13.7 | 1.22 |
| **802** | Harasanshin Hospital | medical | Japan | 31 | 800 | 83 | 2.7 | 0.59 |
| **803** | National Hospital Organization Mie Chuo Medical Center | medical | Japan | 31 | 200 | 356 | 11.5 | 0.74 |
| **804** | National Hospital Organization Higashisaitama National Hospital | medical | Japan | 31 | 400 | 411 | 13.3 | 0.74 |
| **805** | Queen Elizabeth Hospital Australia | medical | Australia | 30 | -80 | 705 | 23.5 | 1.26 |
| **806** | Royal Children's Hospital Brisbane | medical | Australia | 30 | -100 | 1444 | 48.1 | 1.85 |
| **807** | Dalian University of Technology | academic | China | 30 | - | 261 | 8.7 | 0.77 |
| **808** | Ocean University of China | academic | China | 30 | -100 | 2173 | 72.4 | 3.09 |
| **809** | Yanshan University | academic | China | 30 | 50 | 592 | 19.7 | 1.26 |
| **810** | Punjabi University | academic | India | 30 | 0 | 593 | 19.8 | 0.99 |
| **811** | Chiba Cancer Center | medical | Japan | 30 | - | 512 | 17.1 | 1.4 |
| **812** | Soochow University Taiwan | academic | Taiwan | 30 | 300 | 287 | 9.6 | 1 |
| **813** | B P Koirala Institute of Health Sciences | academic | Nepal | 30 | 233.3 | 173 | 5.8 | 1.2 |
| **814** | Hospital Pulau Pinang | medical | Malaysia | 30 | 200 | 524 | 17.5 | 1.47 |
| **815** | International Medical University | academic | Malaysia | 30 | - | 364 | 12.1 | 1.86 |
| **816** | BYL Nair Charitable Hospital & TN Medical College | medical | India | 30 | - | 148 | 4.9 | 0.42 |
| **817** | Kathmandu University | academic | Nepal | 30 | - | 2251 | 75 | 5.38 |
| **818** | UnitingCare Health | medical | Australia | 30 | - | 929 | 31 | 3.11 |
| **819** | Baotou Medical College | academic | China | 30 | 800 | 313 | 10.4 | 0.84 |
| **820** | Kaohsiung Armed Forces General Hospital | medical | Taiwan | 30 | -100 | 501 | 16.7 | 0.7 |
| **821** | Shenyang Medical College | academic | China | 30 | 200 | 326 | 10.9 | 1.04 |
| **822** | Shizuoka City Shizuoka Hospital | medical | Japan | 30 | 66.7 | 252 | 8.4 | 0.73 |
| **823** | Tokyo Metropolitan Matsuzawa Hospital | medical | Japan | 30 | -60 | 275 | 9.2 | 0.64 |
| **824** | National Hospital Organization Omuta National Hospital | medical | Japan | 30 | 0 | 467 | 15.6 | 1.12 |
| **825** | Saiseikai Fukuoka General Hospital | medical | Japan | 30 | 500 | 303 | 10.1 | 1.98 |
| **826** | Indian Institute of Technology Delhi | academic | India | 29 | - | 298 | 10.3 | 0.75 |
| **827** | Chubu University | academic | Japan | 29 | -40 | 396 | 13.7 | 0.85 |
| **828** | Tokyo University of Pharmacy and Life Sciences | academic | Japan | 29 | 200 | 234 | 8.1 | 0.8 |
| **829** | Hamamatsu Medical Center | medical | Japan | 29 | 100 | 621 | 21.4 | 1.17 |
| **830** | Bharath Institute of Higher Education and Research | academic | India | 29 | - | 57 | 2 | 0.23 |
| **831** | Suzuka University of Medical Science | academic | Japan | 29 | -25 | 567 | 19.6 | 1.31 |
| **832** | Department of Health Manila | government | Philippines | 29 | 66.7 | 1204 | 41.5 | 6.7 |
| **833** | Niigata City General Hospital | medical | Japan | 29 | 0 | 406 | 14 | 1.54 |
| **834** | Wuxi People's Hospital | medical | China | 29 | 200 | 445 | 15.3 | 1.57 |
| **835** | Tokoha University | academic | Japan | 29 | 200 | 300 | 10.3 | 0.7 |
| **836** | Yodogawa Christian Hospital | medical | Japan | 29 | 0 | 402 | 13.9 | 0.93 |
| **837** | National Hospital Organization Iou National Hospital | medical | Japan | 29 | 0 | 171 | 5.9 | 0.66 |
| **838** | Kobe City Nishi-Kobe Medical Center | medical | Japan | 29 | 25 | 367 | 12.7 | 0.82 |
| **839** | Guangzhou General Hospital | medical | China | 28 | -100 | 461 | 16.5 | 0.93 |
| **840** | Jiamusi University | academic | China | 28 | - | 243 | 8.7 | 0.97 |
| **841** | The University of Electro-Communications | academic | Japan | 28 | -33.3 | 999 | 35.7 | 1.11 |
| **842** | Mitsubishi Chemical Holdings Corporation | corporate | Japan | 28 | 66.7 | 758 | 27.1 | 1.26 |
| **843** | Sapporo City General Hospital | medical | Japan | 28 | -50 | 369 | 13.2 | 0.76 |
| **844** | University of Kelaniya | academic | Sri Lanka | 28 | -71.4 | 771 | 27.5 | 1.57 |
| **845** | Burnet Institute | academic | Australia | 28 | -50 | 1119 | 40 | 2.95 |
| **846** | Jiujiang University | academic | China | 28 | 200 | 173 | 6.2 | 0.67 |
| **847** | Pelita Harapan University | academic | Indonesia | 28 | 166.7 | 522 | 18.6 | 2.38 |
| **848** | Hebei United University | academic | China | 28 | -100 | 673 | 24 | 1.25 |
| **849** | Chengde Medical University | academic | China | 28 | 300 | 307 | 11 | 0.84 |
| **850** | Teine Keijinkai Hospital | medical | Japan | 28 | 0 | 243 | 8.7 | 0.54 |
| **851** | Tokyo Teishin Hospital | medical | Japan | 28 | 50 | 346 | 12.4 | 0.81 |
| **852** | Shimane Prefectural Central Hospital | medical | Japan | 28 | 300 | 147 | 5.3 | 0.58 |
| **853** | National Hospital Organization Shimoshizu National Hospital | medical | Japan | 28 | - | 575 | 20.5 | 1.26 |
| **854** | Southern Cross University | academic | Australia | 27 | - | 272 | 10.1 | 1.17 |
| **855** | National University of Defense Technology | academic | China | 27 | 100 | 671 | 24.9 | 1.62 |
| **856** | Shanxi University | academic | China | 27 | - | 583 | 21.6 | 1.99 |
| **857** | Princess Margaret Hospital Hong Kong | medical | Hong Kong | 27 | 50 | 269 | 10 | 0.65 |
| **858** | United Christian Hospital | medical | Hong Kong | 27 | 100 | 389 | 14.4 | 1.03 |
| **859** | Aligarh Muslim University | academic | India | 27 | -25 | 360 | 13.3 | 0.55 |
| **860** | Pohang University of Science and Technology | academic | South Korea | 27 | 600 | 219 | 8.1 | 0.65 |
| **861** | Da-Yeh University | academic | Taiwan | 27 | 300 | 238 | 8.8 | 1.35 |
| **862** | Tohoku Fukushi University | academic | Japan | 27 | 0 | 149 | 5.5 | 0.77 |
| **863** | Fifth Medical Center of the General Hospital of the Chinese People's Liberation Army | medical | China | 27 | -66.7 | 396 | 14.7 | 0.91 |
| **864** | Kyungdong University | academic | South Korea | 27 | - | 607 | 22.5 | 1.19 |
| **865** | Dalian University | academic | China | 27 | - | 385 | 14.3 | 1.67 |
| **866** | National Hospital Organization Disaster Medical Center | medical | Japan | 27 | -60 | 340 | 12.6 | 1.22 |
| **867** | Shenyang Pharmaceutical University | academic | China | 26 | - | 477 | 18.3 | 1.37 |
| **868** | Yangtze University | academic | China | 26 | - | 329 | 12.7 | 1.23 |
| **869** | Shihezi University | academic | China | 26 | - | 298 | 11.5 | 1.4 |
| **870** | Gadjah Mada University | academic | Indonesia | 26 | 300 | 158 | 6.1 | 0.58 |
| **871** | University of Shizuoka | academic | Japan | 26 | 100 | 379 | 14.6 | 0.71 |
| **872** | Meiji Pharmaceutical University | academic | Japan | 26 | -100 | 547 | 21 | 1.02 |
| **873** | Al Farabi Kazakh National University | academic | Kazakhstan | 26 | - | 993 | 38.2 | 7.08 |
| **874** | Jamia Millia Islamia | academic | India | 26 | 400 | 431 | 16.6 | 1 |
| **875** | Sarawak General Hospital | medical | Malaysia | 26 | - | 156 | 6 | 0.53 |
| **876** | North Shore Hospital | medical | New Zealand | 26 | - | 757 | 29.1 | 1.72 |
| **877** | National Heart Centre Singapore | medical | Singapore | 26 | 400 | 561 | 21.6 | 1.82 |
| **878** | Sungshin Women's University | academic | South Korea | 26 | -100 | 410 | 15.8 | 0.84 |
| **879** | Chenzhou First People's Hospital | medical | China | 26 | 166.7 | 394 | 15.2 | 1.15 |
| **880** | Shizuoka General Hospital | medical | Japan | 26 | - | 172 | 6.6 | 0.62 |
| **881** | All India Institute of Medical Sciences, Patna | academic | India | 26 | - | 97 | 3.7 | 0.4 |
| **882** | Kansai University of Health Sciences | academic | Japan | 26 | -33.3 | 150 | 5.8 | 0.37 |
| **883** | Okinaka Memorial Institute for Medical Research | medical | Japan | 26 | 0 | 611 | 23.5 | 1.25 |
| **884** | Kumamoto City Hospital | medical | Japan | 26 | 200 | 114 | 4.4 | 0.43 |
| **885** | National Hospital Organization Minami-Kyoto National Hospital | medical | Japan | 26 | -33.3 | 480 | 18.5 | 1.07 |
| **886** | Japanese Red Cross Ise Hospital | medical | Japan | 26 | - | 256 | 9.8 | 0.84 |
| **887** | Gunma Children's Medical Center | medical | Japan | 26 | -100 | 506 | 19.5 | 0.96 |
| **888** | Hangzhou Dianzi University | academic | China | 25 | - | 338 | 13.5 | 1.35 |
| **889** | Nanjing Normal University | academic | China | 25 | - | 751 | 30 | 2.07 |
| **890** | Jawaharlal Nehru University | academic | India | 25 | 200 | 513 | 20.5 | 1.12 |
| **891** | Aichi Cancer Center Hospital and Research Institute | medical | Japan | 25 | 150 | 474 | 19 | 1.21 |
| **892** | Chuo University | academic | Japan | 25 | -50 | 252 | 10.1 | 0.76 |
| **893** | Kansai Electric Power Co., Inc. | corporate | Japan | 25 | - | 105 | 4.2 | 0.66 |
| **894** | Shionogi & Co., Ltd. | corporate | Japan | 25 | 0 | 512 | 20.5 | 1.28 |
| **895** | Tokyo Metropolitan Police Hospital | medical | Japan | 25 | 0 | 197 | 7.9 | 0.5 |
| **896** | International Islamic University Malaysia | academic | Malaysia | 25 | 150 | 152 | 6.1 | 0.57 |
| **897** | National Chung Cheng University | academic | Taiwan | 25 | -100 | 585 | 23.4 | 1.38 |
| **898** | Azerbaycan Tibb Universiteti | academic | Azerbaijan | 25 | - | 285 | 11.4 | 2.27 |
| **899** | Jiangxi University of Traditional Chinese Medicine | academic | China | 25 | - | 285 | 11.4 | 1.08 |
| **900** | Ibaraki Prefectural University of Health Sciences | academic | Japan | 25 | 600 | 136 | 5.4 | 1 |
| **901** | Toyota Memorial Hospital | medical | Japan | 25 | 100 | 267 | 10.7 | 1.36 |
| **902** | Teikyo Heisei University | academic | Japan | 25 | - | 169 | 6.8 | 0.52 |
| **903** | Hollywood Private Hospital | medical | Australia | 25 | -75 | 1058 | 42.3 | 2.17 |
| **904** | Hong Kong Hospital Authority | government | Hong Kong | 25 | 300 | 404 | 16.2 | 1.29 |
| **905** | Hunan Children's Hospital | medical | China | 25 | - | 127 | 5.1 | 0.98 |
| **906** | Sahmyook University | academic | South Korea | 25 | 0 | 341 | 13.6 | 0.69 |
| **907** | PLA General Hospital of Chengdu Military Region | medical | China | 25 | -100 | 473 | 18.9 | 0.78 |
| **908** | PLA No. 309 Hospital | medical | China | 25 | -100 | 220 | 8.8 | 0.62 |
| **909** | Royal Darwin Hospital | medical | Australia | 25 | - | 208 | 8.3 | 1.46 |
| **910** | CSIR - Centre for Cellular Molecular Biology | government | India | 25 | - | 287 | 11.5 | 0.77 |
| **911** | Kochi Health Sciences Center | medical | Japan | 25 | - | 185 | 7.4 | 0.83 |
| **912** | National Hospital Organization Kumamoto Saishunso National Hospital | medical | Japan | 25 | 200 | 271 | 10.8 | 0.88 |
| **913** | Japanese Red Cross Fukui Hospital | medical | Japan | 25 | 100 | 101 | 4 | 0.41 |
| **914** | Japan Community Healthcare Organization Osaka Hospital | medical | Japan | 25 | -50 | 363 | 14.5 | 0.94 |
| **915** | China Agricultural University | academic | China | 24 | 100 | 481 | 20 | 1.38 |
| **916** | Birla Institute of Technology and Science Pilani | academic | India | 24 | - | 409 | 17 | 1.39 |
| **917** | Vellore Institute of Technology | academic | India | 24 | - | 340 | 14.2 | 1.05 |
| **918** | Nagoya Institute of Technology | academic | Japan | 24 | 500 | 811 | 33.8 | 2.32 |
| **919** | Tokushima Bunri University | academic | Japan | 24 | -80 | 264 | 11 | 0.62 |
| **920** | Chukyo University | academic | Japan | 24 | 0 | 359 | 15 | 1.13 |
| **921** | Japanese Foundation for Cancer Research | medical | Japan | 24 | 50 | 364 | 15.2 | 1.23 |
| **922** | Seonam University | academic | South Korea | 24 | - | 228 | 9.5 | 0.52 |
| **923** | Central Taiwan University of Science and Technology | academic | Taiwan | 24 | 0 | 264 | 11 | 0.65 |
| **924** | Kirin Holdings Co., Ltd. | corporate | Japan | 24 | 50 | 767 | 32 | 1.86 |
| **925** | Ishikawa Central Prefectural Hospital | medical | Japan | 24 | 500 | 145 | 6 | 1.32 |
| **926** | Hebei North University | academic | China | 24 | 400 | 207 | 8.6 | 0.78 |
| **927** | Punjab Technical University | academic | India | 24 | - | 560 | 23.3 | 1.62 |
| **928** | Saveetha Institute of Medical and Technical Sciences (Deemed to be University) | academic | India | 24 | 600 | 330 | 13.8 | 1.56 |
| **929** | Medical College of Chinese People's Armed Police Forces | academic | China | 24 | 300 | 251 | 10.5 | 0.66 |
| **930** | Yantai Yantaishan Hospital | medical | China | 24 | 100 | 241 | 10 | 0.64 |
| **931** | Shandong Cancer Hospital | medical | China | 24 | 66.7 | 294 | 12.3 | 0.78 |
| **932** | Japanese Red Cross Maebashi Hospital | medical | Japan | 24 | 50 | 282 | 11.8 | 0.95 |
| **933** | Japanese Red Cross Nagaoka Hospital | medical | Japan | 24 | 300 | 99 | 4.1 | 0.28 |
| **934** | Iwate Prefectural Central Hospital | medical | Japan | 24 | - | 257 | 10.7 | 1.76 |
| **935** | Shinko Hospital | medical | Japan | 24 | 0 | 349 | 14.5 | 0.84 |
| **936** | Guizhou University | academic | China | 23 | - | 237 | 10.3 | 1.5 |
| **937** | Sichuan Normal University | academic | China | 23 | - | 194 | 8.4 | 1.57 |
| **938** | Osmania University | academic | India | 23 | -75 | 653 | 28.4 | 1.13 |
| **939** | Chang Jung Christian University | academic | Taiwan | 23 | 300 | 167 | 7.3 | 1.8 |
| **940** | Universitas Diponegoro | academic | Indonesia | 23 | - | 83 | 3.6 | 0.52 |
| **941** | Daiichi Sankyo Company, Limited | corporate | Japan | 23 | 200 | 440 | 19.1 | 1.46 |
| **942** | Ranchi University | academic | India | 23 | -33.3 | 177 | 7.7 | 0.74 |
| **943** | ShanghaiTech University | academic | China | 23 | - | 181 | 7.9 | 1.73 |
| **944** | Datta Meghe Institute of Medical Sciences | academic | India | 23 | 33.3 | 1006 | 43.7 | 8.19 |
| **945** | Guangzhou Red Cross Hospital | medical | China | 23 | - | 123 | 5.3 | 0.55 |
| **946** | Mudanjiang Medical University | academic | China | 23 | -100 | 291 | 12.7 | 0.77 |
| **947** | First People's Hospital of Foshan | medical | China | 23 | 600 | 173 | 7.5 | 0.66 |
| **948** | Changzhi Medical College | academic | China | 23 | 0 | 291 | 12.7 | 1.01 |
| **949** | Toyama Prefectural Central Hospital | medical | Japan | 23 | - | 136 | 5.9 | 0.98 |
| **950** | Sumitomo Hospital | medical | Japan | 23 | 300 | 471 | 20.5 | 1.59 |
| **951** | Gandhi Medical College | medical | India | 23 | 0 | 2765 | 120.2 | 8.4 |
| **952** | Beijing Shunyi District Hospital | medical | China | 23 | 100 | 116 | 5 | 0.47 |
| **953** | Indian Institute of Technology Banaras Hindu University | academic | India | 23 | - | 293 | 12.7 | 1.13 |
| **954** | National Hospital Organization Kanazawa Medical Center | medical | Japan | 23 | - | 41 | 1.8 | 0.19 |
| **955** | Nagano Red Cross Hospital | medical | Japan | 23 | - | 253 | 11 | 0.63 |
| **956** | Kagawa Rosai Hospital | medical | Japan | 23 | - | 107 | 4.7 | 0.35 |
| **957** | Chugoku Rosai Hospital | medical | Japan | 23 | 200 | 106 | 4.6 | 0.47 |
| **958** | The Peter Doherty Institute for Infection and Immunity | academic | Australia | 23 | - | 321 | 14 | 1.57 |
| **959** | University of New England | academic | Australia | 22 | 100 | 237 | 10.8 | 0.76 |
| **960** | Liaoning Normal University | academic | China | 22 | -25 | 344 | 15.6 | 1.28 |
| **961** | Health Sciences University of Hokkaido | academic | Japan | 22 | -33.3 | 165 | 7.5 | 0.47 |
| **962** | Daejeon University | academic | South Korea | 22 | 0 | 276 | 12.5 | 0.82 |
| **963** | Piramal Enterprises Ltd | corporate | India | 22 | - | 1120 | 50.9 | 2.79 |
| **964** | Sri Balaji Vidyapeeth University | academic | India | 22 | 100 | 139 | 6.3 | 0.59 |
| **965** | Max Healthcare | medical | India | 22 | - | 106 | 4.8 | 0.77 |
| **966** | Academy of Scientific and Innovative Research | academic | India | 22 | - | 453 | 20.6 | 1.04 |
| **967** | Shizuoka Children's Hospital | medical | Japan | 22 | - | 143 | 6.5 | 0.49 |
| **968** | Toyonaka Municipal Hospital | medical | Japan | 22 | 100 | 507 | 23 | 2.48 |
| **969** | Takatsuki General Hospital | medical | Japan | 22 | 50 | 296 | 13.5 | 0.78 |
| **970** | Kitakyushu Municipal Medical Center | medical | Japan | 22 | 0 | 234 | 10.6 | 0.95 |
| **971** | Saiseikai Utsunomiya Hospital | medical | Japan | 22 | 33.3 | 128 | 5.8 | 0.32 |
| **972** | Nara City Hospital | medical | Japan | 22 | - | 169 | 7.7 | 1.14 |
| **973** | National Hospital Organization Osaka Minami Medical Center | medical | Japan | 22 | -100 | 315 | 14.3 | 0.92 |
| **974** | National Hospital Organization Yokohama Medical Center | medical | Japan | 22 | - | 320 | 14.5 | 1.51 |
| **975** | Japanese Red Cross Akita Hospital | medical | Japan | 22 | 100 | 278 | 12.6 | 1 |
| **976** | Bharathiar University | academic | India | 21 | - | 188 | 9 | 1.03 |
| **977** | Government Medical College Srinagar | academic | India | 21 | - | 102 | 4.9 | 0.31 |
| **978** | Kyoto Institute of Technology | academic | Japan | 21 | - | 264 | 12.6 | 0.84 |
| **979** | Sogang University | academic | South Korea | 21 | -50 | 449 | 21.4 | 1.22 |
| **980** | Heart Research Institute | academic | Australia | 21 | -100 | 406 | 19.3 | 1.22 |
| **981** | Universiti Tunku Abdul Rahman (UTAR) | academic | Malaysia | 21 | - | 301 | 14.3 | 1.07 |
| **982** | CAS - Beijing Institute of Genomics | academic | China | 21 | -100 | 487 | 23.2 | 1.17 |
| **983** | West Bengal University of Health Sciences | academic | India | 21 | -87.5 | 121 | 5.8 | 0.39 |
| **984** | Yerevan State Medical University | academic | Armenia | 21 | - | 111 | 5.3 | 0.5 |
| **985** | PLA Lanzhou General Hospital | medical | China | 21 | - | 467 | 22.2 | 1.23 |
| **986** | Qiqihar Medical University | academic | China | 21 | - | 285 | 13.6 | 1.27 |
| **987** | Zhejiang Cancer Hospital | medical | China | 21 | 300 | 293 | 14 | 0.81 |
| **988** | Mejiro University | academic | Japan | 21 | - | 16 | 0.8 | 0.16 |
| **989** | Shatin Hospital | medical | Hong Kong | 21 | 66.7 | 388 | 18.5 | 0.98 |
| **990** | Aichi Children's Health and Medical Center | medical | Japan | 21 | - | 178 | 8.5 | 0.59 |
| **991** | Gauhati Medical College and Hospital | medical | India | 21 | -80 | 161 | 7.7 | 0.42 |
| **992** | Kanto Central Hospital | medical | Japan | 21 | -50 | 2371 | 112.9 | 3.25 |
| **993** | National Hospital Organization Kumamoto Medical Center | medical | Japan | 21 | - | 89 | 4.2 | 0.28 |
| **994** | Tokyo Metropolitan Ebara Hospital | medical | Japan | 21 | -75 | 380 | 18.1 | 0.9 |
| **995** | Japanese Red Cross Suwa Hospital | medical | Japan | 21 | - | 150 | 7.1 | 0.78 |
| **996** | Osaka Saiseikai Nakatsu Hospital | medical | Japan | 21 | 100 | 378 | 18 | 0.92 |
| **997** | University of Shanghai for Science and Technology | academic | China | 20 | 500 | 161 | 8.1 | 0.81 |
| **998** | Yanbian University | academic | China | 20 | - | 167 | 8.4 | 0.54 |
| **999** | Cheongju University | academic | South Korea | 20 | 0 | 156 | 7.8 | 0.66 |
| **1000** | Universiti Malaysia Sarawak | academic | Malaysia | 20 | 100 | 125 | 6.3 | 0.6 |
| **1001** | National University of Sciences and Technology Pakistan | academic | Pakistan | 20 | - | 351 | 17.5 | 1.23 |
| **1002** | CAS - Institute of Genetics and Developmental Biology | academic | China | 20 | 200 | 1820 | 91 | 4.3 |
| **1003** | Nazarbayev University | academic | Kazakhstan | 20 | - | 199 | 9.9 | 1.1 |
| **1004** | King Edward Medical University Lahore | academic | Pakistan | 20 | 600 | 319 | 15.9 | 1.04 |
| **1005** | Abay Kazakh National Pedagogical University | academic | Kazakhstan | 20 | - | 39 | 2 | 0.19 |
| **1006** | Konan Women's University | academic | Japan | 20 | - | 203 | 10.1 | 1.22 |
| **1007** | Tbilisi State Medical University | academic | Georgia | 20 | 200 | 172 | 8.6 | 0.89 |
| **1008** | Shree Guru Gobind Singh Tricentenary University | academic | India | 20 | - | 41 | 2 | 0.37 |
| **1009** | Duke Kunshan University | academic | China | 20 | - | 557 | 27.9 | 3.31 |
| **1010** | Tokushima Prefectural Central Hospital | medical | Japan | 20 | -75 | 158 | 7.9 | 0.44 |
| **1011** | Jinnah Post Graduate Medical Center | medical | Pakistan | 20 | 300 | 149 | 7.4 | 0.71 |
| **1012** | Public Health Foundation of India | other | India | 20 | - | 7727 | 386.4 | 29.8 |
| **1013** | Beijing Electric Power Hospital | medical | China | 20 | 500 | 79 | 4 | 0.36 |
| **1014** | Hamamatsu Photonics K.K. | corporate | Japan | 20 | 300 | 398 | 19.9 | 1.98 |
| **1015** | Kagawa Prefectural Central Hospital | medical | Japan | 20 | - | 174 | 8.7 | 0.67 |
| **1016** | Yamagata Prefectural University of Health Sciences | academic | Japan | 20 | - | 149 | 7.4 | 0.58 |
| **1017** | National Hospital Organization Higashihiroshima Medical Center | medical | Japan | 20 | -66.7 | 132 | 6.6 | 0.48 |
| **1018** | National Hospital Organization Matsue Medical Center | medical | Japan | 20 | - | 230 | 11.5 | 0.92 |
| **1019** | Japanese Red Cross Saitama Hospital | medical | Japan | 20 | 600 | 131 | 6.6 | 0.84 |
| **1020** | Beijing Jiaotong University | academic | China | 19 | 50 | 199 | 10.5 | 0.87 |
| **1021** | Capital Normal University | academic | China | 19 | 0 | 211 | 11.1 | 0.95 |
| **1022** | Shandong Normal University | academic | China | 19 | - | 397 | 20.9 | 1.91 |
| **1023** | Shanghai Normal University | academic | China | 19 | 400 | 385 | 20.3 | 1.59 |
| **1024** | CAS - Institute of High Energy Physics | academic | China | 19 | 300 | 260 | 13.7 | 1.38 |
| **1025** | Indian Institute of Technology Madras | academic | India | 19 | - | 107 | 5.6 | 1.38 |
| **1026** | Indian Statistical Institute | academic | India | 19 | 0 | 203 | 10.7 | 0.63 |
| **1027** | Jeonju University | academic | South Korea | 19 | -100 | 169 | 8.9 | 0.59 |
| **1028** | Middlemore Hospital | medical | New Zealand | 19 | - | 128 | 6.7 | 0.61 |
| **1029** | Tunghai University | academic | Taiwan | 19 | 150 | 110 | 5.8 | 0.67 |
| **1030** | University of Sri Jayewardenepura | academic | Sri Lanka | 19 | -66.7 | 1720 | 90.5 | 2.7 |
| **1031** | Ministry of Water Resources, P.R. China | government | China | 19 | - | 111 | 5.8 | 0.94 |
| **1032** | Hue University | academic | Viet Nam | 19 | 400 | 152 | 8 | 0.71 |
| **1033** | Kalinga Institute of Industrial Technology | academic | India | 19 | 400 | 983 | 51.7 | 9.81 |
| **1034** | Saitama Prefectural University | academic | Japan | 19 | 0 | 119 | 6.3 | 0.42 |
| **1035** | Duy Tan University | academic | Viet Nam | 19 | - | 8589 | 452.1 | 34.77 |
| **1036** | Dali University | academic | China | 19 | - | 173 | 9.1 | 1.43 |
| **1037** | Shifa International Hospital | medical | Pakistan | 19 | -100 | 107 | 5.6 | 0.65 |
| **1038** | Alice Ho Miu Ling Nethersole Hospital | medical | Hong Kong | 19 | 100 | 146 | 7.7 | 0.67 |
| **1039** | Yidu Central Hospital of Weifang | medical | China | 19 | - | 81 | 4.3 | 0.61 |
| **1040** | Hyogo Cancer Center | medical | Japan | 19 | - | 210 | 11.1 | 0.97 |
| **1041** | Sakai City Medical Center | medical | Japan | 19 | - | 280 | 14.7 | 1.38 |
| **1042** | Tachikawa Hospital | medical | Japan | 19 | -71.4 | 1100 | 57.9 | 2.04 |
| **1043** | D Y Patil Group | academic | India | 19 | 300 | 216 | 11.4 | 2.24 |
| **1044** | Lahore General Hospital | medical | Pakistan | 19 | - | 156 | 8.2 | 1.73 |
| **1045** | National Hospital Organization Shikoku Medical Center for Children and Adults | medical | Japan | 19 | - | 142 | 7.5 | 0.73 |
| **1046** | Bach Mai Hospital | medical | Viet Nam | 19 | 600 | 127 | 6.7 | 1.4 |
| **1047** | Taiyuan University of Technology | academic | China | 18 | - | 290 | 16.1 | 1.03 |
| **1048** | Huzhou University | academic | China | 18 | - | 171 | 9.5 | 0.56 |
| **1049** | Jianghan University | academic | China | 18 | 500 | 203 | 11.3 | 1.01 |
| **1050** | University of Hyderabad | academic | India | 18 | -100 | 541 | 30.1 | 1.37 |
| **1051** | The University of Lahore | academic | Pakistan | 18 | - | 1794 | 99.7 | 13.29 |
| **1052** | National Taipei University of Technology | academic | Taiwan | 18 | 0 | 180 | 10 | 0.67 |
| **1053** | Mackay Medicine, Nursing and Management College Taiwan | academic | Taiwan | 18 | 0 | 208 | 11.6 | 0.64 |
| **1054** | Ulsan National Institute of Science and Technology | academic | South Korea | 18 | 50 | 219 | 12.2 | 0.79 |
| **1055** | Okinawa Institute of Science and Technology Graduate University | academic | Japan | 18 | 0 | 616 | 34.2 | 2.59 |
| **1056** | Goa University | academic | India | 18 | -40 | 188 | 10.4 | 0.8 |
| **1057** | Shaoxing University | academic | China | 18 | - | 137 | 7.6 | 1.33 |
| **1058** | Olivia Newton-John Cancer Research Institute | medical | Australia | 18 | - | 379 | 21.1 | 1.4 |
| **1059** | Jen-Teh Junior College of Medicine, Nursing and Management | academic | Taiwan | 18 | - | 219 | 12.2 | 0.87 |
| **1060** | Pandit Bhagwat Dayal Sharma University of Health Sciences | academic | India | 18 | - | 77 | 4.3 | 0.22 |
| **1061** | Wenzhou People's Hospital | medical | China | 18 | - | 304 | 16.9 | 1.04 |
| **1062** | Aino University | academic | Japan | 18 | -80 | 140 | 7.8 | 0.46 |
| **1063** | Gifu Prefectural General Medical Center | medical | Japan | 18 | 0 | 173 | 9.6 | 1.1 |
| **1064** | National Hospital Organization Minami-Okayama Medical Center | medical | Japan | 18 | 100 | 148 | 8.2 | 0.87 |
| **1065** | Japanese Red Cross Otsu Hospital | medical | Japan | 18 | -50 | 100 | 5.6 | 0.46 |
| **1066** | Japanese Red Cross Himeji Hospital | medical | Japan | 18 | 0 | 109 | 6.1 | 0.5 |
| **1067** | Okayama Rosai Hospital | medical | Japan | 18 | - | 58 | 3.2 | 0.7 |
| **1068** | Baba Farid University of Health Sciences | academic | India | 18 | 66.7 | 78 | 4.3 | 0.29 |
| **1069** | Federation University Australia | academic | Australia | 17 | 200 | 1252 | 73.6 | 12.49 |
| **1070** | University of Dhaka | academic | Bangladesh | 17 | - | 3563 | 209.6 | 15.25 |
| **1071** | Anhui University of Science and Technology | academic | China | 17 | - | 85 | 5 | 1.16 |
| **1072** | Southwest Jiaotong University | academic | China | 17 | - | 180 | 10.6 | 0.76 |
| **1073** | Hainan University | academic | China | 17 | - | 96 | 5.6 | 1.97 |
| **1074** | Tokyo Institute of Technology | academic | Japan | 17 | - | 172 | 10.1 | 0.78 |
| **1075** | Tokyo University of Technology | academic | Japan | 17 | 500 | 188 | 11.1 | 1.2 |
| **1076** | Asahi University | academic | Japan | 17 | - | 80 | 4.7 | 0.45 |
| **1077** | Showa Pharmaceutical University | academic | Japan | 17 | -100 | 215 | 12.6 | 0.59 |
| **1078** | University of Peradeniya | academic | Sri Lanka | 17 | - | 6910 | 406.5 | 32.57 |
| **1079** | Yuan Ze University | academic | Taiwan | 17 | - | 261 | 15.4 | 1.2 |
| **1080** | CAS - Wuhan Institute of Physics and Mathematics | academic | China | 17 | -66.7 | 329 | 19.4 | 1.23 |
| **1081** | Asahi Kasei Corporation | corporate | Japan | 17 | - | 187 | 11 | 0.79 |
| **1082** | Taichung Armed Forces General Hospital | medical | Taiwan | 17 | 50 | 88 | 5.2 | 0.51 |
| **1083** | Institute of Nuclear Medicine and Allied Sciences India | government | India | 17 | 0 | 323 | 19 | 1.04 |
| **1084** | Hubei University of Arts and Science | academic | China | 17 | - | 119 | 7 | 1.05 |
| **1085** | University of the Punjab | academic | Pakistan | 17 | 200 | 156 | 9.2 | 0.76 |
| **1086** | Beijing Air Force General Hospital | medical | China | 17 | - | 170 | 10 | 0.54 |
| **1087** | Meiho University | academic | Taiwan | 17 | - | 291 | 17.1 | 0.87 |
| **1088** | Calvary Mater Newcastle | medical | Australia | 17 | 100 | 404 | 23.8 | 1.76 |
| **1089** | Nil Ratan Sircar Medical College and Hospital | medical | India | 17 | -75 | 175 | 10.3 | 0.53 |
| **1090** | Kyoto City Hospital | medical | Japan | 17 | - | 99 | 5.8 | 0.53 |
| **1091** | Gansu University of Chinese Medicine | academic | China | 17 | - | 228 | 13.4 | 1.31 |
| **1092** | Japanese Red Cross Society | medical | Japan | 17 | 100 | 78 | 4.6 | 0.51 |
| **1093** | Chiba Prefectural University of Health Sciences | academic | Japan | 17 | - | 32 | 1.9 | 0.5 |
| **1094** | Nara Prefecture General Medical Center | medical | Japan | 17 | - | 127 | 7.5 | 0.49 |
| **1095** | Mito Kyodo General Hospital | medical | Japan | 17 | - | 68 | 4 | 0.38 |
| **1096** | National Hospital Organization Matsumoto Medical Center | medical | Japan | 17 | - | 174 | 10.2 | 1.11 |
| **1097** | National Hospital Organization Asahikawa Medical Center | medical | Japan | 17 | 300 | 261 | 15.4 | 1.06 |
| **1098** | Japanese Red Cross Fukuoka Hospital | medical | Japan | 17 | -50 | 287 | 16.9 | 0.91 |
| **1099** | Tokyo Rosai Hospital | medical | Japan | 17 | -50 | 234 | 13.8 | 0.51 |
| **1100** | Yamaguchi Rosai Hospital | medical | Japan | 17 | - | 56 | 3.3 | 0.99 |
| **1101** | Rajarata University of Sri Lanka | academic | Sri Lanka | 17 | - | 1100 | 64.7 | 12.18 |
| **1102** | East China University of Science and Technology | academic | China | 16 | -66.7 | 313 | 19.6 | 1.4 |
| **1103** | Guangzhou University | academic | China | 16 | 0 | 160 | 10 | 0.89 |
| **1104** | South-Central University for Nationalities | academic | China | 16 | 0 | 325 | 20.3 | 1.11 |
| **1105** | Yunnan University | academic | China | 16 | - | 201 | 12.6 | 0.98 |
| **1106** | Anna University | academic | India | 16 | - | 122 | 7.6 | 1.19 |
| **1107** | Ochanomizu University | academic | Japan | 16 | 0 | 143 | 8.9 | 0.65 |
| **1108** | Sophia University | academic | Japan | 16 | - | 152 | 9.5 | 0.7 |
| **1109** | Fukui Prefectural University | academic | Japan | 16 | -50 | 253 | 15.8 | 0.85 |
| **1110** | National Agriculture and Food Research Organization | government | Japan | 16 | - | 294 | 18.4 | 0.85 |
| **1111** | Kyungnam University | academic | South Korea | 16 | 0 | 234 | 14.6 | 1.07 |
| **1112** | Institute of Technical Education and Research of Siksha O Anusandhan University | academic | India | 16 | - | 59 | 3.7 | 0.65 |
| **1113** | Regional Cancer Centre India | medical | India | 16 | - | 34 | 2.1 | 0.51 |
| **1114** | St. Luke's International University | academic | Japan | 16 | 100 | 93 | 5.8 | 0.54 |
| **1115** | Tsukuba University of Technology | academic | Japan | 16 | - | 100 | 6.3 | 0.44 |
| **1116** | Cochlear | corporate | Australia | 16 | -100 | 389 | 24.3 | 1.82 |
| **1117** | Wockhardt Limited | corporate | India | 16 | - | 48 | 3 | 0.48 |
| **1118** | Chitkara University | academic | India | 16 | - | 291 | 18.2 | 1.91 |
| **1119** | Yokosuka Kyosai Hospital | medical | Japan | 16 | -50 | 631 | 39.4 | 2.03 |
| **1120** | Kyoto Katsura Hospital | medical | Japan | 16 | - | 259 | 16.2 | 1.06 |
| **1121** | Mahatma Gandhi Institute of Medical Sciences | medical | India | 16 | -100 | 131 | 8.2 | 0.43 |
| **1122** | Baroda Medical College | medical | India | 16 | -75 | 133 | 8.3 | 0.52 |
| **1123** | Meiji University of Integrative Medicine | academic | Japan | 16 | -100 | 144 | 9 | 0.65 |
| **1124** | Fukui Prefectural Hospital | medical | Japan | 16 | - | 67 | 4.2 | 0.57 |
| **1125** | National Hospital Organization Kinki-Chuo Chest Medical Center | medical | Japan | 16 | - | 150 | 9.4 | 2.18 |
| **1126** | National Hospital Organization Beppu Medical Center | medical | Japan | 16 | - | 74 | 4.6 | 0.39 |
| **1127** | National Hospital Organization Niigata National Hospital | medical | Japan | 16 | - | 137 | 8.6 | 0.83 |
| **1128** | Japanese Red Cross Matsue Hospital | medical | Japan | 16 | - | 205 | 12.8 | 2.7 |
| **1129** | Japan Community Healthcare Organization Kyushu Hospital | medical | Japan | 16 | -100 | 144 | 9 | 0.73 |
| **1130** | Edogawa Hospital | medical | Japan | 16 | -100 | 93 | 5.8 | 0.4 |
| **1131** | Kyrgyz State Medical Academy | academic | Kyrgyzstan | 16 | 33.3 | 5207 | 325.4 | 32.61 |
| **1132** | Zhejiang University of Technology | academic | China | 15 | 300 | 118 | 7.9 | 0.95 |
| **1133** | Iwate University | academic | Japan | 15 | 100 | 159 | 10.6 | 0.84 |
| **1134** | Kochi University of Technology | academic | Japan | 15 | - | 155 | 10.3 | 0.68 |
| **1135** | Meijo University | academic | Japan | 15 | 0 | 79 | 5.3 | 0.35 |
| **1136** | Nara Women's University | academic | Japan | 15 | -100 | 409 | 27.3 | 0.78 |
| **1137** | Tokyo Gakugei University | academic | Japan | 15 | -75 | 78 | 5.2 | 0.28 |
| **1138** | Tamagawa University | academic | Japan | 15 | -100 | 198 | 13.2 | 0.77 |
| **1139** | University of Peshawar | academic | Pakistan | 15 | 400 | 149 | 9.9 | 0.65 |
| **1140** | Beijing Sport University | academic | China | 15 | - | 182 | 12.1 | 1.06 |
| **1141** | L.N. Gumilyov Eurasian National University | academic | Kazakhstan | 15 | - | 30 | 2 | 0.22 |
| **1142** | CSL Limited | corporate | Australia | 15 | -33.3 | 377 | 25.1 | 1.52 |
| **1143** | Defence Research and Development Organisation India | government | India | 15 | - | 281 | 18.7 | 0.93 |
| **1144** | National Natural Science Foundation of China | academic | China | 15 | -100 | 80 | 5.3 | 0.4 |
| **1145** | Aichi Shukutoku University | academic | Japan | 15 | -50 | 98 | 6.5 | 0.38 |
| **1146** | Changchun University of Chinese Medicine | academic | China | 15 | 100 | 170 | 11.3 | 0.68 |
| **1147** | Waikato District Health Board | government | New Zealand | 15 | - | 137 | 9.1 | 1.7 |
| **1148** | Allama Iqbal Medical College | academic | Pakistan | 15 | - | 142 | 9.5 | 0.84 |
| **1149** | Japan Agency for Medical Research and Development | government | Japan | 15 | - | 306 | 20.4 | 1.67 |
| **1150** | Shiga General Hospital | medical | Japan | 15 | - | 33 | 2.2 | 0.4 |
| **1151** | Kimitsu Chuo Hospital | medical | Japan | 15 | - | 269 | 17.9 | 1.44 |
| **1152** | National Hospital Organization Kurihama Medical and Addiction Center | medical | Japan | 15 | - | 439 | 29.3 | 1.13 |
| **1153** | National Hospital Organization Chiba-East National Hospital | medical | Japan | 15 | -100 | 728 | 48.5 | 2.41 |
| **1154** | National Hospital Organization Kobe Medical Center | medical | Japan | 15 | -100 | 354 | 23.6 | 1.11 |
| **1155** | Japan Community Healthcare Organization Chukyo Hospital | medical | Japan | 15 | 0 | 106 | 7.1 | 0.62 |
| **1156** | Japan Community Healthcare Organization Sendai Hospital | medical | Japan | 15 | - | 167 | 11.1 | 1.15 |
| **1157** | University of Southern Queensland | academic | Australia | 14 | - | 155 | 11.1 | 1.42 |
| **1158** | Anhui University | academic | China | 14 | - | 291 | 20.8 | 1.19 |
| **1159** | Fujian Normal University | academic | China | 14 | 0 | 305 | 21.8 | 1.28 |
| **1160** | University of Lucknow | academic | India | 14 | -100 | 183 | 13.1 | 0.67 |
| **1161** | Kanazawa Institute of Technology | academic | Japan | 14 | - | 111 | 7.9 | 1.09 |
| **1162** | National Institutes of Natural Sciences | academic | Japan | 14 | - | 115 | 8.2 | 0.69 |
| **1163** | COMSATS University Islamabad | academic | Pakistan | 14 | - | 231 | 16.5 | 0.81 |
| **1164** | Chinese Culture University | academic | Taiwan | 14 | -100 | 117 | 8.4 | 0.69 |
| **1165** | Tamkang University | academic | Taiwan | 14 | -100 | 448 | 32 | 1.79 |
| **1166** | University of Nottingham Malaysia Campus | academic | Malaysia | 14 | 200 | 162 | 11.6 | 0.62 |
| **1167** | Palmerston North Hospital | medical | New Zealand | 14 | - | 150 | 10.7 | 1.54 |
| **1168** | Vision Research Foundation India | government | India | 14 | - | 57 | 4.1 | 0.36 |
| **1169** | Nagasaki Institute of Applied Science | academic | Japan | 14 | - | 47 | 3.4 | 0.31 |
| **1170** | Kumamoto Health Science University | academic | Japan | 14 | - | 104 | 7.4 | 0.7 |
| **1171** | Shanghai International Studies University | academic | China | 14 | - | 125 | 8.9 | 1.42 |
| **1172** | Kazakh National Medical University | academic | Kazakhstan | 14 | 500 | 1662 | 118.7 | 10.08 |
| **1173** | Xiangnan University | academic | China | 14 | - | 124 | 8.9 | 1.53 |
| **1174** | Taiwanese Injury Prevention and Safety Promotion Association | other | Taiwan | 14 | - | 379 | 27.1 | 1.68 |
| **1175** | Lovely Professional University | academic | India | 14 | - | 274 | 19.6 | 1.49 |
| **1176** | Riphah International University | academic | Pakistan | 14 | - | 103 | 7.4 | 1.36 |
| **1177** | Shanxi Datong University | academic | China | 14 | - | 220 | 15.7 | 1.15 |
| **1178** | Cheng Ching Hospital | medical | Taiwan | 14 | 0 | 150 | 10.7 | 0.74 |
| **1179** | Min-Sheng General Hospital | medical | Taiwan | 14 | -100 | 170 | 12.1 | 0.7 |
| **1180** | Ziauddin University | academic | Pakistan | 14 | 100 | 233 | 16.6 | 1.35 |
| **1181** | Government Medical College Nagpur | medical | India | 14 | - | 25 | 1.8 | 0.28 |
| **1182** | Ministry of Health, Government of Singapore | government | Singapore | 14 | 100 | 97 | 6.9 | 1.13 |
| **1183** | Fujirebio Inc. | corporate | Japan | 14 | 100 | 722 | 51.6 | 3.25 |
| **1184** | CSIR - Central Institute of Medicinal Aromatic Plants | government | India | 14 | - | 111 | 7.9 | 0.51 |
| **1185** | Kibi International University | academic | Japan | 14 | - | 131 | 9.4 | 1.19 |
| **1186** | Navamindradhiraj University | academic | Thailand | 14 | - | 33 | 2.4 | 0.42 |
| **1187** | Yamagata Prefectural Central Hospital | medical | Japan | 14 | - | 108 | 7.7 | 0.56 |
| **1188** | Kagoshima City Hospital | medical | Japan | 14 | 300 | 162 | 11.6 | 2.03 |
| **1189** | National Hospital Organization Fukuoka-Higashi Medical Center | medical | Japan | 14 | - | 171 | 12.2 | 0.63 |
| **1190** | National Hospital Organization Hizen Psychiatric Center | medical | Japan | 14 | -100 | 221 | 15.8 | 0.99 |
| **1191** | JA Hiroshima General Hospital | medical | Japan | 14 | 500 | 98 | 7 | 0.47 |
| **1192** | Central China Normal University | academic | China | 13 | - | 224 | 17.2 | 1.48 |
| **1193** | Tianjin Normal University | academic | China | 13 | - | 135 | 10.4 | 0.92 |
| **1194** | CAS - Center for Excellence in Molecular Cell Science | academic | China | 13 | - | 180 | 13.8 | 1.06 |
| **1195** | Kansai University | academic | Japan | 13 | -60 | 242 | 18.6 | 0.93 |
| **1196** | Tokyo University of Agriculture and Technology | academic | Japan | 13 | - | 120 | 9.2 | 0.56 |
| **1197** | Kobe Pharmaceutical University | academic | Japan | 13 | - | 223 | 17.2 | 1.02 |
| **1198** | Gangneung-Wonju National University | academic | South Korea | 13 | - | 202 | 15.5 | 1.09 |
| **1199** | Incheon National University | academic | South Korea | 13 | - | 295 | 22.7 | 1.45 |
| **1200** | Dongseo University | academic | South Korea | 13 | - | 152 | 11.7 | 0.85 |
| **1201** | Hankuk University of Foreign Studies | academic | South Korea | 13 | - | 140 | 10.8 | 1.11 |
| **1202** | Korea National Institute of Health | government | South Korea | 13 | -50 | 303 | 23.3 | 1.38 |
| **1203** | Quaid-I-Azam University | academic | Pakistan | 13 | - | 51 | 3.9 | 0.23 |
| **1204** | Bahauddin Zakariya University | academic | Pakistan | 13 | - | 217 | 16.7 | 1.1 |
| **1205** | National Science and Technology Development Agency Thailand | government | Thailand | 13 | - | 176 | 13.5 | 0.73 |
| **1206** | Universitas Hasanuddin | academic | Indonesia | 13 | - | 86 | 6.6 | 0.77 |
| **1207** | The Wesley Research Institute | academic | Australia | 13 | - | 320 | 24.6 | 1.78 |
| **1208** | NED University of Engineering and Technology | academic | Pakistan | 13 | - | 292 | 22.5 | 0.95 |
| **1209** | West Japan Railway Company | corporate | Japan | 13 | - | 147 | 11.3 | 1.04 |
| **1210** | University of Taipei | academic | Taiwan | 13 | 100 | 218 | 16.8 | 1.13 |
| **1211** | Symbiosis International University | academic | India | 13 | - | 199 | 15.3 | 1.32 |
| **1212** | Deccan College of Medical Sciences | academic | India | 13 | - | 39 | 3 | 0.79 |
| **1213** | Korea Institute of Toxicology | government | South Korea | 13 | - | 395 | 30.4 | 1.56 |
| **1214** | Center for Medical Genetics and Primary Health Care | medical | Armenia | 13 | 200 | 55 | 4.2 | 1.75 |
| **1215** | King Edward Memorial Hospital for Women | medical | Australia | 13 | 500 | 463 | 35.6 | 1.45 |
| **1216** | Central University of Punjab, Bathinda | academic | India | 13 | - | 183 | 14.1 | 0.95 |
| **1217** | Institute of Liver and Biliary Sciences | medical | India | 13 | - | 104 | 8 | 0.96 |
| **1218** | Zhejiang University City College | academic | China | 13 | 600 | 154 | 11.8 | 1.6 |
| **1219** | Jingzhou Central Hospital | medical | China | 13 | - | 831 | 63.9 | 12.08 |
| **1220** | Shenzhen Center for Disease Control and Prevention | government | China | 13 | - | 186 | 14.3 | 1.19 |
| **1221** | Lotung Poh-Ai Hospital | medical | Taiwan | 13 | -50 | 123 | 9.5 | 0.87 |
| **1222** | Castle Peak Hospital | medical | Hong Kong | 13 | - | 269 | 20.7 | 1.41 |
| **1223** | Shifa Tameer-e-Millat University | academic | Pakistan | 13 | -100 | 50 | 3.8 | 0.39 |
| **1224** | Hiroshima Cosmopolitan University | academic | Japan | 13 | - | 51 | 3.9 | 0.94 |
| **1225** | Osaki Citizen Hospital | medical | Japan | 13 | - | 49 | 3.8 | 0.28 |
| **1226** | Japanese Red Cross Okayama Hospital | medical | Japan | 13 | 100 | 112 | 8.6 | 0.65 |
| **1227** | Japan Community Healthcare Organization Hoshigaoka Medical Center | medical | Japan | 13 | - | 176 | 13.5 | 0.98 |
| **1228** | Saiseikai Matsusaka General Hospital | medical | Japan | 13 | 100 | 133 | 10.2 | 0.59 |
| **1229** | Yokkaichi Municipal Hospital | medical | Japan | 13 | 100 | 44 | 3.4 | 0.41 |
| **1230** | Renmin University of China | academic | China | 12 | - | 108 | 9 | 1.03 |
| **1231** | University of Jinan | academic | China | 12 | -100 | 96 | 8 | 0.54 |
| **1232** | CAS - Shanghai Institute of Materia Medica | academic | China | 12 | - | 637 | 53.1 | 2.59 |
| **1233** | Guru Jambeshwar University of Science and Technology | academic | India | 12 | - | 13 | 1.1 | 0.08 |
| **1234** | L.V. Prasad Eye Institute India | medical | India | 12 | 0 | 39 | 3.3 | 0.28 |
| **1235** | Hiroshima International University | academic | Japan | 12 | 0 | 178 | 14.8 | 1.03 |
| **1236** | National Institutes of Biomedical Innovation, Health and Nutrition | government | Japan | 12 | 0 | 140 | 11.7 | 1.11 |
| **1237** | Sangmyung University | academic | South Korea | 12 | - | 172 | 14.3 | 1.31 |
| **1238** | Srinakharinwirot University | academic | Thailand | 12 | 50 | 30 | 2.5 | 0.41 |
| **1239** | Chung Yuan Christian University | academic | Taiwan | 12 | -50 | 146 | 12.2 | 0.69 |
| **1240** | National Taiwan University of Science and Technology | academic | Taiwan | 12 | -100 | 149 | 12.4 | 0.61 |
| **1241** | Hebei University of Engineering | academic | China | 12 | - | 124 | 10.3 | 1.17 |
| **1242** | Hutt Hospital | medical | New Zealand | 12 | 0 | 285 | 23.8 | 1.87 |
| **1243** | Aravind Eye Hospital | medical | India | 12 | 0 | 53 | 4.4 | 0.9 |
| **1244** | Pakistan Institute of Medical Sciences | government | Pakistan | 12 | 100 | 122 | 10.2 | 1.66 |
| **1245** | Daegu Haany University | academic | South Korea | 12 | -100 | 270 | 22.5 | 0.89 |
| **1246** | Musashino University | academic | Japan | 12 | - | 1005 | 83.8 | 11.7 |
| **1247** | Nagasaki International University | academic | Japan | 12 | - | 50 | 4.2 | 0.57 |
| **1248** | Ricoh | corporate | Japan | 12 | - | 74 | 6.2 | 0.71 |
| **1249** | National Pingtung University | academic | Taiwan | 12 | -50 | 136 | 11.3 | 0.57 |
| **1250** | Yenepoya University | academic | India | 12 | - | 54 | 4.5 | 0.18 |
| **1251** | Singapore Institute of Technology | medical | Singapore | 12 | - | 173 | 14.4 | 0.86 |
| **1252** | Kyoto Tachibana University | academic | Japan | 12 | 0 | 109 | 9.1 | 0.5 |
| **1253** | Ilia State University | academic | Georgia | 12 | - | 142 | 11.8 | 0.71 |
| **1254** | Sharda University | academic | India | 12 | - | 84 | 7 | 0.67 |
| **1255** | Yiwu Central Hospital | medical | China | 12 | - | 142 | 11.8 | 0.77 |
| **1256** | Chongqing Three Gorges University | academic | China | 12 | - | 457 | 38.1 | 3.42 |
| **1257** | Beijing Institute of Ophthalmology | medical | China | 12 | - | 3720 | 310 | 30.9 |
| **1258** | Lin Shin Hospital | medical | Taiwan | 12 | 0 | 141 | 11.8 | 0.67 |
| **1259** | Kunming General Hospital of Chengdu Military Command | medical | China | 12 | -100 | 251 | 20.9 | 0.9 |
| **1260** | Yokohama Municipal Citizen's Hospital | medical | Japan | 12 | - | 63 | 5.3 | 0.41 |
| **1261** | Hong Kong Sanatorium & Hospital | medical | Hong Kong | 12 | - | 309 | 25.8 | 1.26 |
| **1262** | Ahmedabad Civil Hospital | medical | India | 12 | - | 110 | 9.2 | 4.49 |
| **1263** | Father Muller Medical College Hospital | medical | India | 12 | 150 | 962 | 80.2 | 14.77 |
| **1264** | National Healthcare Group, Singapore | corporate | Singapore | 12 | 0 | 446 | 37.2 | 1.9 |
| **1265** | Chengdu Women's & Children's Central Hospital | medical | China | 12 | - | 24 | 2 | 0.22 |
| **1266** | Jingjiang People's Hospital | medical | China | 12 | - | 69 | 5.8 | 2.97 |
| **1267** | Ogaki Municipal Hospital | medical | Japan | 12 | -100 | 64 | 5.3 | 0.39 |
| **1268** | National Centre for Infectious Diseases | academic | Singapore | 12 | - | 193 | 16.1 | 2.86 |
| **1269** | Kagawa Prefectural University of Health Sciences | academic | Japan | 12 | - | 63 | 5.3 | 0.32 |
| **1270** | Ichinomiya Municipal Hospital | medical | Japan | 12 | -50 | 185 | 15.4 | 0.59 |
| **1271** | Jigme Dorji Wangchuck National Referral Hospital | medical | Bhutan | 12 | - | 102 | 8.5 | 0.76 |
| **1272** | National Hospital Organization Himeji Medical Center | medical | Japan | 12 | - | 122 | 10.2 | 0.53 |
| **1273** | National Hospital Organization Kure Medical Center and Chugoku Cancer Center | medical | Japan | 12 | 200 | 45 | 3.8 | 0.59 |
| **1274** | National Hospital Organization Shinshu Ueda Medical Center | medical | Japan | 12 | - | 272 | 22.7 | 1.81 |
| **1275** | Yokohama City Minato Red Cross Hospital | medical | Japan | 12 | - | 176 | 14.7 | 0.87 |
| **1276** | Narita Red Cross Hospital | medical | Japan | 12 | - | 119 | 9.9 | 1.31 |
| **1277** | Kobe Rosai Hospital | medical | Japan | 12 | -100 | 443 | 36.9 | 1.61 |
| **1278** | Ijinkai Takeda General Hospital | medical | Japan | 12 | 200 | 149 | 12.4 | 0.8 |
| **1279** | Suita Municipal Hospital | medical | Japan | 12 | 100 | 22 | 1.8 | 0.15 |
| **1280** | Iwata City Hospital | medical | Japan | 12 | 0 | 56 | 4.7 | 0.71 |
| **1281** | Omi Medical Center | medical | Japan | 12 | - | 84 | 7 | 1.05 |
| **1282** | Guangxi Normal University | academic | China | 11 | - | 166 | 15.1 | 0.97 |
| **1283** | Huazhong Agricultural University | academic | China | 11 | - | 164 | 14.9 | 1.59 |
| **1284** | CAS - Institute of Zoology | academic | China | 11 | - | 184 | 16.7 | 0.77 |
| **1285** | Changchun University | academic | China | 11 | 200 | 54 | 4.9 | 0.69 |
| **1286** | Annamalai University | academic | India | 11 | - | 103 | 9.4 | 0.35 |
| **1287** | University of Madras | academic | India | 11 | -100 | 196 | 17.8 | 0.76 |
| **1288** | University of Pune | academic | India | 11 | - | 188 | 17.1 | 2.45 |
| **1289** | Mahatma Gandhi University, Kottayam | academic | India | 11 | - | 982 | 89.3 | 16.79 |
| **1290** | Rashtrasant Tukadoji Maharaj Nagpur University | academic | India | 11 | 0 | 128 | 11.6 | 0.88 |
| **1291** | Meiji University | academic | Japan | 11 | - | 75 | 6.8 | 0.71 |
| **1292** | Saitama University | academic | Japan | 11 | 400 | 128 | 11.6 | 0.96 |
| **1293** | Azabu University | academic | Japan | 11 | -100 | 181 | 16.5 | 0.92 |
| **1294** | Canon Inc. | corporate | Japan | 11 | - | 294 | 26.7 | 2.68 |
| **1295** | Toshiba Corporation | corporate | Japan | 11 | - | 109 | 9.9 | 1.01 |
| **1296** | King Mongkut's Institute of Technology Ladkrabang | academic | Thailand | 11 | 400 | 125 | 11.4 | 1.67 |
| **1297** | Mongolian National University of Medical Sciences | academic | Mongolia | 11 | - | 362 | 32.9 | 4.97 |
| **1298** | CAS - Suzhou Institute of Biomedical Engineering and Technology | academic | China | 11 | 0 | 99 | 9 | 0.81 |
| **1299** | Institute for Medical Research Kuala Lumpur | academic | Malaysia | 11 | - | 76 | 6.9 | 0.57 |
| **1300** | University of Kashmir | academic | India | 11 | - | 191 | 17.4 | 1.12 |
| **1301** | North South University | academic | Bangladesh | 11 | - | 196 | 17.8 | 1.43 |
| **1302** | New York University Shanghai | academic | China | 11 | - | 93 | 8.5 | 1.07 |
| **1303** | New South Wales Ministry of Health | government | Australia | 11 | - | 311 | 28.3 | 2.2 |
| **1304** | Bankstown-Lidcombe Hospital | medical | Australia | 11 | -50 | 134 | 12.2 | 0.45 |
| **1305** | Semey Medical University | medical | Kazakhstan | 11 | - | 45 | 4.1 | 0.26 |
| **1306** | Mercy Hospital for Women | medical | Australia | 11 | 0 | 315 | 28.6 | 1.57 |
| **1307** | Abdul Wali Khan University Mardan | academic | Pakistan | 11 | - | 131 | 11.9 | 1.21 |
| **1308** | B.J. Medical College | academic | India | 11 | - | 70 | 6.4 | 4.49 |
| **1309** | Hebei University of Chinese Medicine | academic | China | 11 | - | 176 | 16 | 2.16 |
| **1310** | Astana Medical University | academic | Kazakhstan | 11 | - | 48 | 4.4 | 0.7 |
| **1311** | Chulabhorn Royal Academy | government | Thailand | 11 | - | 256 | 23.3 | 1.33 |
| **1312** | Shanghai Cancer Institute | medical | China | 11 | 200 | 254 | 23.1 | 1.81 |
| **1313** | Shanghai Punan Hospital | medical | China | 11 | - | 105 | 9.5 | 0.67 |
| **1314** | Sydney Cancer Centre | medical | Australia | 11 | -50 | 272 | 24.7 | 1.93 |
| **1315** | Kyoto Koka Women's University | academic | Japan | 11 | -100 | 179 | 16.3 | 0.62 |
| **1316** | Navitas | corporate | Australia | 11 | - | 432 | 39.3 | 1.93 |
| **1317** | Tokyo Ariake University of Medical and Health Sciences | academic | Japan | 11 | 200 | 87 | 7.9 | 0.82 |
| **1318** | Kansai University of Welfare Sciences | academic | Japan | 11 | 0 | 110 | 10 | 0.67 |
| **1319** | ARC Centre of Excellence for Electromaterials Science | academic | Australia | 11 | -100 | 110 | 10 | 0.7 |
| **1320** | Tokyo Metropolitan Toshima Hospital | medical | Japan | 11 | - | 24 | 2.2 | 0.19 |
| **1321** | National Hospital Organization Okinawa National Hospital | medical | Japan | 11 | - | 59 | 5.4 | 0.44 |
| **1322** | National Hospital Organization Higashi-Ohmi General Medical Center | medical | Japan | 11 | 0 | 47 | 4.3 | 0.35 |
| **1323** | Japanese Red Cross Karatsu Hospital | medical | Japan | 11 | -100 | 76 | 6.9 | 0.39 |
| **1324** | Nagasaki Rosai Hospital | medical | Japan | 11 | -50 | 34 | 3.1 | 0.5 |
| **1325** | Yokohama Minami Kyosai Hospital | medical | Japan | 11 | - | 202 | 18.4 | 1.39 |
| **1326** | Eiju General Hospital | medical | Japan | 11 | -100 | 107 | 9.7 | 0.68 |
| **1327** | Toyota Kosei Hospital | medical | Japan | 11 | - | 64 | 5.8 | 1.77 |
| **1328** | Jiangxi Normal University | academic | China | 10 | - | 47 | 4.7 | 0.43 |
| **1329** | Northwest Agriculture and Forestry University | academic | China | 10 | - | 166 | 16.6 | 1.03 |
| **1330** | Beihua University | academic | China | 10 | 100 | 113 | 11.3 | 0.81 |
| **1331** | North China University of Technology | academic | China | 10 | - | 115 | 11.5 | 0.93 |
| **1332** | Indian Institute of Technology Kanpur | academic | India | 10 | -100 | 186 | 18.6 | 1.1 |
| **1333** | Kyushu Institute of Technology | academic | Japan | 10 | 100 | 39 | 3.9 | 0.4 |
| **1334** | National Institute for Environmental Studies of Japan | government | Japan | 10 | - | 137 | 13.7 | 0.9 |
| **1335** | National Institute of Public Health of Japan | government | Japan | 10 | 100 | 113 | 11.3 | 1.43 |
| **1336** | Sunchon National University | academic | South Korea | 10 | - | 198 | 19.8 | 1.13 |
| **1337** | Hanseo University | academic | South Korea | 10 | -100 | 171 | 17.1 | 0.82 |
| **1338** | National Ilan University Taiwan | academic | Taiwan | 10 | - | 78 | 7.8 | 0.72 |
| **1339** | National Yunlin University of Science and Technology | academic | Taiwan | 10 | - | 931 | 93.1 | 17.74 |
| **1340** | Yuanpei University of Medical Technology | academic | Taiwan | 10 | - | 172 | 17.2 | 0.65 |
| **1341** | Universiti Sultan Zainal Abidin | academic | Malaysia | 10 | - | 1044 | 104.4 | 18.8 |
| **1342** | Centenary Institute | academic | Australia | 10 | - | 306 | 30.6 | 2.6 |
| **1343** | West Kazakhstan Marat Ospanov State Medical University | academic | Kazakhstan | 10 | - | 9 | 0.9 | 0.31 |
| **1344** | CAS - Guangzhou Institute of Biomedicine and Health | academic | China | 10 | - | 69 | 6.9 | 0.77 |
| **1345** | BRAC University | academic | Bangladesh | 10 | - | 5636 | 563.6 | 37.68 |
| **1346** | Jaypee University of Information Technology | academic | India | 10 | -100 | 345 | 34.5 | 1.34 |
| **1347** | Shiga University | academic | Japan | 10 | - | 123 | 12.3 | 1.23 |
| **1348** | Institute of Genetics and Hospital for Genetic Diseases, Hyderabad | academic | India | 10 | -100 | 137 | 13.7 | 0.59 |
| **1349** | Academia Sinica - Institute of Statistical Science | academic | Taiwan | 10 | 100 | 71 | 7.1 | 0.44 |
| **1350** | Nihon Fukushi University | academic | Japan | 10 | - | 69 | 6.9 | 0.46 |
| **1351** | Osaka Dental University | academic | Japan | 10 | - | 77 | 7.7 | 0.5 |
| **1352** | Tokyo International University | academic | Japan | 10 | - | 58 | 5.8 | 0.75 |
| **1353** | Xi'an Jiaotong-Liverpool University | academic | China | 10 | - | 150 | 15 | 1.47 |
| **1354** | Graylands Hospital | medical | Australia | 10 | - | 163 | 16.3 | 1.14 |
| **1355** | University of Health Sciences Lahore | academic | Pakistan | 10 | - | 29 | 2.9 | 0.28 |
| **1356** | University of North Sumatra | academic | Indonesia | 10 | - | 48 | 4.8 | 0.61 |
| **1357** | National Taiwan Sport University | academic | Taiwan | 10 | 100 | 132 | 13.2 | 0.76 |
| **1358** | Jinnah Sindh Medical University | academic | Pakistan | 10 | - | 145 | 14.5 | 0.61 |
| **1359** | Ramaiah Institute of Technology | academic | India | 10 | - | 76 | 7.6 | 0.65 |
| **1360** | National Research Institute for Family Planning, Beijing | academic | China | 10 | - | 86 | 8.6 | 0.5 |
| **1361** | St. Paul's Hospital Taoyuan | medical | Taiwan | 10 | -100 | 164 | 16.4 | 1.2 |
| **1362** | Ohta Nishinouchi Hospital | medical | Japan | 10 | -50 | 205 | 20.5 | 1.19 |
| **1363** | North District Hospital Hong Kong | medical | Hong Kong | 10 | - | 109 | 10.9 | 0.77 |
| **1364** | Shanghai Eighth People's Hospital | medical | China | 10 | - | 89 | 8.9 | 0.93 |
| **1365** | Southeast University, Dhaka | academic | Bangladesh | 10 | - | 146 | 14.6 | 1.33 |
| **1366** | Rajiv Gandhi Government General Hospital | medical | India | 10 | - | 71 | 7.1 | 0.75 |
| **1367** | Shaheed Zulfiqar Ali Bhutto Medical University | academic | Pakistan | 10 | - | 25 | 2.5 | 0.86 |
| **1368** | Hamanomachi Hospital | medical | Japan | 10 | 200 | 38 | 3.8 | 0.26 |
| **1369** | Tokyo Kasei University | academic | Japan | 10 | - | 133 | 13.3 | 1.14 |
| **1370** | Okayama City Hospital | medical | Japan | 10 | 100 | 74 | 7.4 | 0.69 |
| **1371** | Sardar Patel Medical College | medical | India | 10 | - | 18 | 1.8 | 0.24 |
| **1372** | National Hospital Organization Shizuoka Medical Center | medical | Japan | 10 | 100 | 172 | 17.2 | 1.83 |
| **1373** | National Hospital Organization Kagoshima Medical Center | medical | Japan | 10 | - | 117 | 11.7 | 0.71 |
| **1374** | National Hospital Organization Takasaki General Medical Center | medical | Japan | 10 | 0 | 84 | 8.4 | 0.57 |
| **1375** | Japanese Red Cross Nagahama Hospital | medical | Japan | 10 | 300 | 101 | 10.1 | 0.74 |
| **1376** | Sanin Rosai Hospital | medical | Japan | 10 | - | 48 | 4.8 | 0.31 |
| **1377** | Okayama Saiseikai General Hospital | medical | Japan | 10 | - | 118 | 11.8 | 0.86 |
| **1378** | Tokyo Kyosai Hospital | medical | Japan | 10 | 100 | 380 | 38 | 4.09 |
| **1379** | Peng Cheng Laboratory | government | China | 10 | - | 122 | 12.2 | 1.6 |
| **1380** | Cancer Council Victoria | academic | Australia | 9 | - | 99 | 11 | 0.84 |
| **1381** | Hunter Health | medical | Australia | 9 | - | 123 | 13.7 | 1.05 |
| **1382** | Hebei Normal University | academic | China | 9 | - | 117 | 13 | 1.41 |
| **1383** | Hubei University | academic | China | 9 | - | 71 | 7.9 | 0.88 |
| **1384** | Hunan University | academic | China | 9 | - | 147 | 16.3 | 1.32 |
| **1385** | Northeast Normal University | academic | China | 9 | - | 161 | 17.9 | 1.18 |
| **1386** | Wenzhou University | academic | China | 9 | - | 161 | 17.9 | 1.34 |
| **1387** | Yantai University | academic | China | 9 | - | 70 | 7.8 | 0.73 |
| **1388** | China Academy of Traditional Chinese Medicine | academic | China | 9 | - | 113 | 12.6 | 0.7 |
| **1389** | Chinese National Human Genome Center | government | China | 9 | -100 | 233 | 25.9 | 1.95 |
| **1390** | Hubei University for Nationalities | academic | China | 9 | - | 14 | 1.6 | 0.25 |
| **1391** | Bhabha Atomic Research Centre | government | India | 9 | -100 | 52 | 5.8 | 0.48 |
| **1392** | Tata Institute of Fundamental Research | academic | India | 9 | - | 154 | 17.1 | 0.73 |
| **1393** | University of Mumbai | academic | India | 9 | - | 52 | 5.8 | 0.47 |
| **1394** | Acharya Nagarjuna University | academic | India | 9 | - | 201 | 22.3 | 1.01 |
| **1395** | University of Hyogo | academic | Japan | 9 | - | 45 | 5 | 0.72 |
| **1396** | Hoshi University | academic | Japan | 9 | -100 | 319 | 35.4 | 1.49 |
| **1397** | Kwansei Gakuin University | academic | Japan | 9 | - | 33 | 3.7 | 0.28 |
| **1398** | Shizuoka University | academic | Japan | 9 | -100 | 84 | 9.3 | 0.42 |
| **1399** | Toyo University | academic | Japan | 9 | 33.3 | 306 | 34 | 1.28 |
| **1400** | Matsumoto Dental University | academic | Japan | 9 | -100 | 215 | 23.9 | 1.23 |
| **1401** | Korea Research Institute of Standards and Science | government | South Korea | 9 | -100 | 106 | 11.8 | 0.73 |
| **1402** | Kwangwoon University | academic | South Korea | 9 | - | 220 | 24.4 | 1.34 |
| **1403** | Sejong University | academic | South Korea | 9 | - | 96 | 10.7 | 1.19 |
| **1404** | National Changhua University of Education | academic | Taiwan | 9 | 50 | 194 | 21.6 | 0.72 |
| **1405** | Victor Chang Cardiac Research Institute | government | Australia | 9 | - | 257 | 28.6 | 3.07 |
| **1406** | Universitas Sebelas Maret | academic | Indonesia | 9 | - | 28 | 3.1 | 0.32 |
| **1407** | Fukushima University | academic | Japan | 9 | -100 | 220 | 24.4 | 1.28 |
| **1408** | Ton Duc Thang University | academic | Viet Nam | 9 | - | 125 | 13.9 | 1.18 |
| **1409** | Taylor's University Malaysia | academic | Malaysia | 9 | - | 59 | 6.6 | 0.78 |
| **1410** | Mysore Medical College | academic | India | 9 | - | 979 | 108.8 | 19.76 |
| **1411** | Tata Group India | corporate | India | 9 | -100 | 1027 | 114.1 | 3.45 |
| **1412** | Shenzhen Polytechnic | academic | China | 9 | - | 9 | 1 | 0.25 |
| **1413** | TaiZhou University | academic | China | 9 | - | 77 | 8.6 | 0.82 |
| **1414** | Centre for DNA Fingerprinting and Diagnostics | academic | India | 9 | - | 33 | 3.7 | 0.48 |
| **1415** | Ngee Ann Polytechnic | academic | Singapore | 9 | - | 1055 | 117.2 | 2.67 |
| **1416** | Rajamangala University of Technology Isan | academic | Thailand | 9 | - | 57 | 6.3 | 0.93 |
| **1417** | Rajiv Gandhi Centre for Biotechnology | academic | India | 9 | - | 116 | 12.9 | 1.19 |
| **1418** | Liaoning University of Traditional Chinese Medicine | academic | China | 9 | - | 130 | 14.4 | 1.18 |
| **1419** | Fo Guang University | academic | Taiwan | 9 | - | 60 | 6.7 | 0.46 |
| **1420** | Shanxi University of Traditional Chinese Medicine | academic | China | 9 | - | 211 | 23.4 | 1.2 |
| **1421** | China National Nuclear Corporation | corporate | China | 9 | - | 70 | 7.8 | 0.76 |
| **1422** | Baekseok University | academic | South Korea | 9 | - | 139 | 15.4 | 0.87 |
| **1423** | Sikkim Manipal University | academic | India | 9 | -33.3 | 66 | 7.3 | 0.4 |
| **1424** | Bangur Institute of Neurosciences | academic | India | 9 | - | 101 | 11.2 | 0.71 |
| **1425** | Bahria University | academic | Pakistan | 9 | - | 376 | 41.8 | 2.62 |
| **1426** | Jiangsu Provincial Center for Disease Control and Prevention | government | China | 9 | - | 101 | 11.2 | 1.01 |
| **1427** | Atma Jaya Catholic University of Indonesia | academic | Indonesia | 9 | - | 97 | 10.8 | 1.14 |
| **1428** | Bethune International Peace Hospital | medical | China | 9 | - | 123 | 13.7 | 0.7 |
| **1429** | PLA No. 401 Hospital | medical | China | 9 | -100 | 123 | 13.7 | 0.61 |
| **1430** | Gifu University of Medical Science | academic | Japan | 9 | -100 | 64 | 7.1 | 1.53 |
| **1431** | Nihon Kohden Corporation | corporate | Japan | 9 | -100 | 168 | 18.7 | 0.8 |
| **1432** | Japan Institute of Sports Sciences | government | Japan | 9 | 100 | 31 | 3.4 | 0.39 |
| **1433** | Seirei Christopher University | academic | Japan | 9 | - | 59 | 6.6 | 0.96 |
| **1434** | Tsuruga Nursing University | academic | Japan | 9 | - | 71 | 7.9 | 0.94 |
| **1435** | Oita Prefectural Hospital | medical | Japan | 9 | - | 24 | 2.7 | 0.18 |
| **1436** | CAS - Tianjin Institute of Industrial Biotechnology | academic | China | 9 | -100 | 223 | 24.8 | 3.22 |
| **1437** | National Hospital Organization Chiba Medical Center | medical | Japan | 9 | - | 49 | 5.4 | 0.54 |
| **1438** | National Hospital Organization Shimofusa Psychiatric Medical Center | medical | Japan | 9 | 200 | 62 | 6.9 | 0.67 |
| **1439** | National Hospital Organization Maizuru Medical Center | medical | Japan | 9 | 200 | 52 | 5.8 | 0.73 |
| **1440** | Tokyo Metropolitan Tama-Hokubu Medical Center | medical | Japan | 9 | -100 | 43 | 4.8 | 0.23 |
| **1441** | Japanese Red Cross Wakayama Medical Center | medical | Japan | 9 | 100 | 41 | 4.6 | 0.6 |
| **1442** | Hiroshima Red Cross Hospital & Atomic-bomb Survivors Hospital | medical | Japan | 9 | 0 | 54 | 6 | 0.59 |
| **1443** | Japanese Red Cross Tokushima Hospital | medical | Japan | 9 | 0 | 59 | 6.6 | 0.35 |
| **1444** | Moji Medical Center | medical | Japan | 9 | - | 101 | 11.2 | 0.6 |
| **1445** | Shizuoka Saiseikai General Hospital | medical | Japan | 9 | - | 138 | 15.3 | 0.68 |
| **1446** | Koseikai Takeda Hospital | medical | Japan | 9 | - | 235 | 26.1 | 1.51 |
| **1447** | Anhui Normal University | academic | China | 8 | - | 229 | 28.6 | 1.78 |
| **1448** | Beijing University of Chemical Technology | academic | China | 8 | 0 | 122 | 15.3 | 0.66 |
| **1449** | Beijing University of Posts and Telecommunications | academic | China | 8 | - | 34 | 4.3 | 0.87 |
| **1450** | Guangdong University of Technology | academic | China | 8 | - | 112 | 14 | 1.17 |
| **1451** | Hebei University of Technology | academic | China | 8 | 400 | 82 | 10.3 | 1.24 |
| **1452** | Liaocheng University | academic | China | 8 | - | 67 | 8.4 | 0.86 |
| **1453** | Nanjing University of Science and Technology | academic | China | 8 | - | 151 | 18.9 | 1.38 |
| **1454** | Jiangsu Normal University | academic | China | 8 | - | 107 | 13.4 | 2.01 |
| **1455** | Qilu University of Technology | academic | China | 8 | - | 186 | 23.3 | 1.39 |
| **1456** | Ministry of Agriculture of the People's Republic of China | government | China | 8 | 0 | 153 | 19.1 | 2.15 |
| **1457** | Bharathidasan University | academic | India | 8 | - | 85 | 10.6 | 0.74 |
| **1458** | University of Mysore | academic | India | 8 | -100 | 55 | 6.9 | 0.42 |
| **1459** | Utkal University | academic | India | 8 | -100 | 6064 | 758 | 44.52 |
| **1460** | Vidyasagar University | academic | India | 8 | -100 | 39 | 4.9 | 0.49 |
| **1461** | Nara Institute of Science and Technology | academic | Japan | 8 | - | 52 | 6.5 | 0.53 |
| **1462** | Prefectural University of Hiroshima | academic | Japan | 8 | - | 35 | 4.4 | 0.52 |
| **1463** | Mukogawa Women's University | academic | Japan | 8 | - | 45 | 5.6 | 0.9 |
| **1464** | Kookmin University | academic | South Korea | 8 | - | 78 | 9.8 | 0.48 |
| **1465** | Universiti Teknologi Malaysia | academic | Malaysia | 8 | 0 | 187 | 23.4 | 2.03 |
| **1466** | National Chiayi University | academic | Taiwan | 8 | 0 | 338 | 42.3 | 1.75 |
| **1467** | National Academy of Sciences of the Republic of Armenia | academic | Armenia | 8 | 0 | 40 | 5 | 0.35 |
| **1468** | Burapha University | academic | Thailand | 8 | 400 | 27 | 3.4 | 0.41 |
| **1469** | Brawijaya University | academic | Indonesia | 8 | - | 25 | 3.1 | 0.64 |
| **1470** | Indian Institute of Technology Gandhinagar | academic | India | 8 | 0 | 116 | 14.5 | 0.87 |
| **1471** | Universiti Sains Islam Malaysia | academic | Malaysia | 8 | 0 | 44 | 5.5 | 0.58 |
| **1472** | Vietnam National University Ho Chi Minh City | academic | Viet Nam | 8 | - | 85 | 10.6 | 1.04 |
| **1473** | Assam University | academic | India | 8 | 0 | 225 | 28.1 | 1.45 |
| **1474** | Academia Sinica - Institute of Molecular Biology | academic | Taiwan | 8 | - | 414 | 51.8 | 2.63 |
| **1475** | Southwest University for Nationalities | academic | China | 8 | - | 238 | 29.8 | 2.97 |
| **1476** | UCSI University | academic | Malaysia | 8 | - | 51 | 6.4 | 1.26 |
| **1477** | Sunway University | academic | Malaysia | 8 | 0 | 69 | 8.6 | 0.65 |
| **1478** | Yasuda Women's University | academic | Japan | 8 | - | 133 | 16.6 | 1.11 |
| **1479** | Greenslopes Private Hospital | medical | Australia | 8 | - | 69 | 8.6 | 0.65 |
| **1480** | Ballarat Health Services | medical | Australia | 8 | - | 92 | 11.5 | 0.74 |
| **1481** | Wanganui District Health Board | government | New Zealand | 8 | - | 129 | 16.1 | 0.55 |
| **1482** | Counties Manukau District Health Board | government | New Zealand | 8 | - | 146 | 18.3 | 1.34 |
| **1483** | Tashkent Medical Academy | academic | Uzbekistan | 8 | 0 | 61 | 7.6 | 0.59 |
| **1484** | MagQu Co., Ltd. | corporate | Taiwan | 8 | - | 359 | 44.9 | 2.76 |
| **1485** | SVKM's NMIMS | academic | India | 8 | - | 305 | 38.1 | 1.92 |
| **1486** | Advanced Centre for Treatment Research and Education in Cancer | academic | India | 8 | 100 | 63 | 7.9 | 0.57 |
| **1487** | Guangdong Center for Disease Control and Prevention | government | China | 8 | - | 128 | 16 | 0.95 |
| **1488** | Yunnan University of Traditional Chinese Medicine | academic | China | 8 | - | 60 | 7.5 | 0.96 |
| **1489** | Liaoning Tumor Hospital & Institute | medical | China | 8 | - | 62 | 7.8 | 1.16 |
| **1490** | Medical Research Foundation, Chennai | medical | India | 8 | 100 | 18 | 2.3 | 0.15 |
| **1491** | Nagoya Women's University | academic | Japan | 8 | 100 | 30 | 3.8 | 0.15 |
| **1492** | Tai Po Hospital | medical | Hong Kong | 8 | -100 | 116 | 14.5 | 0.51 |
| **1493** | Chengdu Sport University | academic | China | 8 | - | 38 | 4.8 | 0.66 |
| **1494** | Cancer Institute NSW | medical | Australia | 8 | - | 256 | 32 | 2.93 |
| **1495** | Meiji Holdings Co., Ltd. | corporate | Japan | 8 | - | 98 | 12.3 | 1.05 |
| **1496** | Kanagawa University of Human Services | academic | Japan | 8 | 200 | 50 | 6.3 | 1.03 |
| **1497** | Zhejiang Provincial Center for Disease Control and Prevention | other | China | 8 | - | 36 | 4.5 | 0.4 |
| **1498** | Tokyo Bay Urayasu Ichikawa Medical Center | medical | Japan | 8 | - | 87 | 10.9 | 0.87 |
| **1499** | Tzu Chi University of Science and Technology | academic | Taiwan | 8 | -50 | 80 | 10 | 0.61 |
| **1500** | Khyber Medical University | academic | Pakistan | 8 | 100 | 106 | 13.3 | 0.84 |
| **1501** | Osaka Yukioka College of Health Science | academic | Japan | 8 | - | 100 | 12.5 | 0.74 |
| **1502** | Kansai Electric Power Medical Research Institute | other | Japan | 8 | - | 75 | 9.4 | 2.91 |
| **1503** | Choithram Hospital and Research center | medical | India | 8 | - | 39 | 4.9 | 0.53 |
| **1504** | Saku Central Hospital | medical | Japan | 8 | - | 133 | 16.6 | 0.78 |
| **1505** | Sam Ratulangi University | academic | Indonesia | 8 | - | 21 | 2.6 | 0.17 |
| **1506** | Tokyo Metropolitan Ohtsuka Hospital | medical | Japan | 8 | - | 62 | 7.8 | 0.76 |
| **1507** | National Hospital Organization Kyushu Cancer Center | medical | Japan | 8 | - | 137 | 17.1 | 1.47 |
| **1508** | National Hospital Organization Mie National Hospital | medical | Japan | 8 | - | 75 | 9.4 | 1.02 |
| **1509** | National Hospital Organization Ureshino Medical Center | medical | Japan | 8 | 100 | 37 | 4.6 | 0.32 |
| **1510** | National Hospital Organization Fukuoka National Hospital | medical | Japan | 8 | -100 | 183 | 22.9 | 0.84 |
| **1511** | Japanese Red Cross Matsuyama Hospital | medical | Japan | 8 | -100 | 196 | 24.5 | 0.86 |
| **1512** | Cho Ray Hospital | medical | Viet Nam | 8 | 200 | 36 | 4.5 | 0.61 |
| **1513** | America Nepal Medical Foundation | other | Nepal | 8 | 400 | 65 | 8.1 | 1.52 |
| **1514** | Kathmandu Medical College Teaching Hospital | academic | Nepal | 8 | 100 | 70 | 8.8 | 5.92 |
| **1515** | Australian Nuclear Science and Technology Organisation | government | Australia | 7 | 0 | 413 | 59 | 3.48 |
| **1516** | Jahangirnagar University | academic | Bangladesh | 7 | 100 | 144 | 20.6 | 5.3 |
| **1517** | Northeast Agricultural University | academic | China | 7 | - | 59 | 8.4 | 0.8 |
| **1518** | Beijing Union University | academic | China | 7 | - | 189 | 27 | 0.88 |
| **1519** | CAS - Dalian Institute of Chemical Physics | academic | China | 7 | - | 138 | 19.7 | 2.5 |
| **1520** | CAS - Institute of Process Engineering | academic | China | 7 | - | 119 | 17 | 0.98 |
| **1521** | Fujian Agriculture and Forestry University | academic | China | 7 | - | 68 | 9.7 | 1.34 |
| **1522** | Jishou University | academic | China | 7 | - | 57 | 8.1 | 0.64 |
| **1523** | Lishui University | academic | China | 7 | - | 38 | 5.4 | 0.33 |
| **1524** | Hamdard University | academic | India | 7 | - | 40 | 5.7 | 0.35 |
| **1525** | Madurai Kamaraj University | academic | India | 7 | - | 218 | 31.1 | 3.2 |
| **1526** | Tamil Nadu Agricultural University | academic | India | 7 | 100 | 100 | 14.3 | 2.17 |
| **1527** | Ibaraki University | academic | Japan | 7 | - | 60 | 8.6 | 1.17 |
| **1528** | Kanagawa Dental University | academic | Japan | 7 | -50 | 46 | 6.6 | 0.23 |
| **1529** | Kyushu Dental College | academic | Japan | 7 | - | 84 | 12 | 0.62 |
| **1530** | Nippon Veterinary and Life Science University | academic | Japan | 7 | - | 68 | 9.7 | 0.49 |
| **1531** | Sony Group Corporation | corporate | Japan | 7 | - | 40 | 5.7 | 0.35 |
| **1532** | Kyonggi University | academic | South Korea | 7 | - | 57 | 8.1 | 0.47 |
| **1533** | Sang Ji University | academic | South Korea | 7 | - | 64 | 9.1 | 0.61 |
| **1534** | Woosuk University | academic | South Korea | 7 | 300 | 32 | 4.6 | 1.03 |
| **1535** | Universiti Malaysia Sabah | academic | Malaysia | 7 | - | 17 | 2.4 | 0.71 |
| **1536** | Universiti Teknologi Petronas | academic | Malaysia | 7 | 0 | 103 | 14.7 | 1.29 |
| **1537** | Kasetsart University | academic | Thailand | 7 | 0 | 86 | 12.3 | 1.03 |
| **1538** | Providence University Taiwan | academic | Taiwan | 7 | - | 56 | 8 | 0.56 |
| **1539** | Fooyin University Taiwan | academic | Taiwan | 7 | - | 81 | 11.6 | 1.3 |
| **1540** | Yerevan State University | academic | Armenia | 7 | - | 85 | 12.1 | 1.17 |
| **1541** | Ateneo de Manila University | academic | Philippines | 7 | - | 96 | 13.7 | 0.85 |
| **1542** | Children's Medical Research Institute | academic | Australia | 7 | - | 60 | 8.6 | 2.7 |
| **1543** | Walailak University | academic | Thailand | 7 | 100 | 1003 | 143.3 | 25.65 |
| **1544** | Gold Coast University Hospital | medical | Australia | 7 | -100 | 456 | 65.1 | 1.97 |
| **1545** | Beijing Information Science & Technology University | academic | China | 7 | - | 26 | 3.7 | 0.65 |
| **1546** | National Institute of Technology Calicut | academic | India | 7 | - | 319 | 45.6 | 1.96 |
| **1547** | Ono Pharmaceutical Co., Ltd. | corporate | Japan | 7 | - | 194 | 27.7 | 3.14 |
| **1548** | Rangsit University | academic | Thailand | 7 | - | 16 | 2.3 | 0.16 |
| **1549** | Shujitsu University | academic | Japan | 7 | -100 | 84 | 12 | 1.05 |
| **1550** | Himeji Dokkyo University | academic | Japan | 7 | -100 | 128 | 18.3 | 0.89 |
| **1551** | Hanson Institute | academic | Australia | 7 | -100 | 86 | 12.3 | 1.84 |
| **1552** | Institute of Chemical Technology | academic | India | 7 | - | 156 | 22.3 | 1.43 |
| **1553** | Karaganda State Medical Academy | academic | Kazakhstan | 7 | - | 19 | 2.7 | 0.17 |
| **1554** | Khoja Akhmet Yassawi International Kazakh-Turkish University | academic | Kazakhstan | 7 | -100 | 9 | 1.3 | 0.1 |
| **1555** | Inner Mongolia University for Nationalities | academic | China | 7 | - | 54 | 7.7 | 0.72 |
| **1556** | Semyung University | academic | South Korea | 7 | -100 | 288 | 41.1 | 1.54 |
| **1557** | Yichun University | academic | China | 7 | - | 35 | 5 | 0.51 |
| **1558** | Sonic Healthcare | medical | Australia | 7 | -100 | 36 | 5.1 | 0.24 |
| **1559** | University of Phayao | academic | Thailand | 7 | 100 | 118 | 16.9 | 0.92 |
| **1560** | People's University | academic | India | 7 | 100 | 4 | 0.6 | 0.26 |
| **1561** | National Kaohsiung University of Science and Technology | academic | Taiwan | 7 | -100 | 40 | 5.7 | 0.3 |
| **1562** | Zhetysu State University named after I. Zhansugurov | academic | Kazakhstan | 7 | - | 11 | 1.6 | 0.17 |
| **1563** | Putian University | academic | China | 7 | - | 17 | 2.4 | 0.45 |
| **1564** | Ministry of Justice, China | government | China | 7 | 0 | 75 | 10.7 | 0.58 |
| **1565** | Xiamen Chang Gung Hospital | medical | China | 7 | - | 41 | 5.9 | 0.51 |
| **1566** | PLA No. 101 Hospital | medical | China | 7 | - | 117 | 16.7 | 0.91 |
| **1567** | Northwest Minzu University | academic | China | 7 | - | 53 | 7.6 | 0.76 |
| **1568** | Tainan Municipal Hospital | medical | Taiwan | 7 | 0 | 38 | 5.4 | 1.08 |
| **1569** | Yuan's General Hospital | medical | Taiwan | 7 | -66.7 | 110 | 15.7 | 0.48 |
| **1570** | Taiwan Adventist Hospital | medical | Taiwan | 7 | -100 | 110 | 15.7 | 0.94 |
| **1571** | Keelung Hospital | medical | Taiwan | 7 | 500 | 70 | 10 | 1.66 |
| **1572** | Melbourne Sexual Health Centre | medical | Australia | 7 | - | 179 | 25.6 | 2.68 |
| **1573** | Changsha Medical University | academic | China | 7 | - | 80 | 11.4 | 1.77 |
| **1574** | Kidwai Memorial Institute of Oncology | medical | India | 7 | - | 35 | 5 | 0.52 |
| **1575** | Rajiv Gandhi Cancer Institute and Research Centre | medical | India | 7 | - | 2 | 0.3 | 0.05 |
| **1576** | Children's Hospital Lahore | medical | Pakistan | 7 | - | 74 | 10.6 | 1.1 |
| **1577** | Asian Institute of Gastroenterology India | academic | India | 7 | -100 | 167 | 23.9 | 1.7 |
| **1578** | West Visayas State University | academic | Philippines | 7 | -100 | 163 | 23.3 | 1.24 |
| **1579** | Caritas Medical Centre Hong Kong | medical | Hong Kong | 7 | - | 60 | 8.6 | 0.7 |
| **1580** | Tseung Kwan O Hospital | medical | Hong Kong | 7 | - | 88 | 12.6 | 0.76 |
| **1581** | Heilongjiang Provincial Hospital | medical | China | 7 | - | 104 | 14.9 | 1.31 |
| **1582** | Vietnam Military Medical University | academic | Viet Nam | 7 | - | 4201 | 600.1 | 43.45 |
| **1583** | CSIR - Central Drug Research Institute | government | India | 7 | 0 | 235 | 33.6 | 1.9 |
| **1584** | Military Hospital Secunderabad | medical | India | 7 | - | 93 | 13.3 | 0.67 |
| **1585** | Melanoma Institute Australia | other | Australia | 7 | - | 260 | 37.1 | 2.05 |
| **1586** | South Kazakhstan Medical Academy | academic | Kazakhstan | 7 | - | 26 | 3.7 | 0.36 |
| **1587** | National Skin Centre, Singapore | medical | Singapore | 7 | - | 21 | 3 | 2.03 |
| **1588** | Tashkent Pediatric Medical Institute | academic | Uzbekistan | 7 | - | 17 | 2.4 | 0.22 |
| **1589** | Showa Women's University | academic | Japan | 7 | - | 53 | 7.6 | 0.58 |
| **1590** | Seijoh University | academic | Japan | 7 | 0 | 124 | 17.7 | 1.02 |
| **1591** | Aomori University of Health and Welfare | academic | Japan | 7 | - | 34 | 4.9 | 0.39 |
| **1592** | Gifu Municipal Hospital | medical | Japan | 7 | - | 4 | 0.6 | 0.14 |
| **1593** | Ageo Central General Hospital | medical | Japan | 7 | - | 66 | 9.4 | 0.88 |
| **1594** | Sasebo City General Hospital | medical | Japan | 7 | 0 | 151 | 21.6 | 1.05 |
| **1595** | Kanagawa Cancer Center Research Institute | medical | Japan | 7 | - | 93 | 13.3 | 0.86 |
| **1596** | National Hospital Organization Hiroshima-Nishi Medical Center | medical | Japan | 7 | - | 32 | 4.6 | 0.5 |
| **1597** | Japanese Red Cross Takayama Hospital | medical | Japan | 7 | - | 60 | 8.6 | 0.38 |
| **1598** | Tohoku Rosai Hospital | medical | Japan | 7 | - | 32 | 4.6 | 0.54 |
| **1599** | Fukui-ken Saiseikai Hospital | medical | Japan | 7 | - | 40 | 5.7 | 0.35 |
| **1600** | Kasugai Municipal Hospital | medical | Japan | 7 | - | 100 | 14.3 | 1.06 |
| **1601** | Chung Jen College of Nursing Health Science and Management | academic | Taiwan | 7 | - | 19 | 2.7 | 0.13 |
| **1602** | China University of Mining and Technology | academic | China | 6 | 0 | 53 | 8.8 | 0.42 |
| **1603** | Chinese Academy of Agricultural Sciences | government | China | 6 | - | 126 | 21 | 0.93 |
| **1604** | Chongqing University of Posts and Telecommunications | academic | China | 6 | - | 27 | 4.5 | 0.41 |
| **1605** | Fuzhou University | academic | China | 6 | -100 | 55 | 9.2 | 0.61 |
| **1606** | Huaqiao University | academic | China | 6 | - | 80 | 13.3 | 1.48 |
| **1607** | Sichuan Agricultural University | academic | China | 6 | - | 51 | 8.5 | 1.12 |
| **1608** | Wuhan University of Technology | academic | China | 6 | - | 41 | 6.8 | 0.7 |
| **1609** | Jiangsu Institute of Cancer Institute & Hospital | medical | China | 6 | - | 73 | 12.2 | 1.08 |
| **1610** | Jiangxi University of Finance and Economics | academic | China | 6 | - | 134 | 22.3 | 0.91 |
| **1611** | Jinggangshan University | academic | China | 6 | - | 22 | 3.7 | 1.14 |
| **1612** | Ningxia University | academic | China | 6 | - | 71 | 11.8 | 0.86 |
| **1613** | Guru Nanak Dev University | academic | India | 6 | - | 64 | 10.7 | 0.62 |
| **1614** | Indian Institute of Technology Bombay | academic | India | 6 | - | 61 | 10.2 | 0.57 |
| **1615** | University of Allahabad | academic | India | 6 | -100 | 90 | 15 | 0.63 |
| **1616** | National Institute of Technology Rourkela | academic | India | 6 | - | 41 | 6.8 | 0.66 |
| **1617** | Aoyama Gakuin University | academic | Japan | 6 | - | 319 | 53.2 | 1.6 |
| **1618** | Tokyo City University | academic | Japan | 6 | - | 39 | 6.5 | 0.89 |
| **1619** | Rikkyo University | academic | Japan | 6 | -100 | 26 | 4.3 | 0.41 |
| **1620** | Shibaura Institute of Technology | academic | Japan | 6 | - | 37 | 6.2 | 0.72 |
| **1621** | Chugai Pharmaceutical Co. Ltd. | corporate | Japan | 6 | - | 124 | 20.7 | 2.1 |
| **1622** | Research Organization of Information and Systems, The Institute of Statistical Mathematics | academic | Japan | 6 | -100 | 90 | 15 | 0.8 |
| **1623** | Setsunan University | academic | Japan | 6 | 100 | 14 | 2.3 | 0.32 |
| **1624** | Soka University | academic | Japan | 6 | -100 | 47 | 7.8 | 0.24 |
| **1625** | Dongeui University | academic | South Korea | 6 | - | 96 | 16 | 0.94 |
| **1626** | Hannam University | academic | South Korea | 6 | 0 | 116 | 19.3 | 1.51 |
| **1627** | Sookmyung Women's University | academic | South Korea | 6 | 0 | 69 | 11.5 | 0.83 |
| **1628** | Korea National University of Transportation | academic | South Korea | 6 | - | 72 | 12 | 0.94 |
| **1629** | De La Salle University-Manila | academic | Philippines | 6 | - | 77 | 12.8 | 0.64 |
| **1630** | National Taipei University | academic | Taiwan | 6 | 0 | 54 | 9 | 0.95 |
| **1631** | National Taichung University of Science and Technology | academic | Taiwan | 6 | 0 | 39 | 6.5 | 0.47 |
| **1632** | National Institute of Biological Sciences, Beijing | academic | China | 6 | -100 | 70 | 11.7 | 0.78 |
| **1633** | University of Veterinary and Animal Sciences, Lahore, Pakistan | academic | Pakistan | 6 | - | 1019 | 169.8 | 30.53 |
| **1634** | Buketov Karaganda State University | academic | Kazakhstan | 6 | - | 3 | 0.5 | 0.05 |
| **1635** | Guangdong University of Foreign Studies | academic | China | 6 | - | 157 | 26.2 | 1.89 |
| **1636** | Fujifilm Corporation | corporate | Japan | 6 | - | 71 | 11.8 | 1.3 |
| **1637** | Ibaraki Prefectural Central Hospital | medical | Japan | 6 | - | 79 | 13.2 | 0.71 |
| **1638** | Foundation University Islamabad | academic | Pakistan | 6 | - | 5103 | 850.5 | 79.57 |
| **1639** | Tajen University | academic | Taiwan | 6 | - | 71 | 11.8 | 1.85 |
| **1640** | National Chin-Yi University of Technology Taiwan | academic | Taiwan | 6 | - | 3 | 0.5 | 0.32 |
| **1641** | Islamia University | academic | Pakistan | 6 | - | 53 | 8.8 | 0.28 |
| **1642** | University of Arid Agriculture Rawalpindi | academic | Pakistan | 6 | - | 64 | 10.7 | 0.98 |
| **1643** | International Islamic University Islamabad | academic | Pakistan | 6 | - | 13 | 2.2 | 0.36 |
| **1644** | Hokkaido University of Science | academic | Japan | 6 | - | 55 | 9.2 | 0.71 |
| **1645** | National Institute of Nutrition | academic | India | 6 | - | 74 | 12.3 | 1.3 |
| **1646** | Christ University, Bangalore | academic | India | 6 | - | 33 | 5.5 | 0.56 |
| **1647** | National Institute of Fitness and Sports in KANOYA | academic | Japan | 6 | -100 | 229 | 38.2 | 1.34 |
| **1648** | National Institutes of Natural Sciences - National Institute for Basic Biology | academic | Japan | 6 | - | 225 | 37.5 | 1.42 |
| **1649** | Academia Sinica - Genomics Research Center | academic | Taiwan | 6 | - | 183 | 30.5 | 2.36 |
| **1650** | Asian Institute of Medicine, Science & Technology | academic | Malaysia | 6 | 0 | 17 | 2.8 | 0.3 |
| **1651** | Kyoto Women's University | academic | Japan | 6 | -100 | 86 | 14.3 | 0.8 |
| **1652** | Hong Kong Metropolitan University | academic | Hong Kong | 6 | - | 66 | 11 | 1.35 |
| **1653** | Nirma University | academic | India | 6 | - | 306 | 51 | 2.8 |
| **1654** | Maebashi Institute of Technology | academic | Japan | 6 | - | 121 | 20.2 | 0.99 |
| **1655** | Ohu University | academic | Japan | 6 | -100 | 176 | 29.3 | 1.09 |
| **1656** | Takasaki University of Health and Welfare | academic | Japan | 6 | - | 35 | 5.8 | 0.37 |
| **1657** | Queensland Eye Institute | medical | Australia | 6 | - | 161 | 26.8 | 1.46 |
| **1658** | The Chinese University of Hong Kong, Shenzhen | academic | China | 6 | - | 44 | 7.3 | 1.15 |
| **1659** | Kainan University | academic | Taiwan | 6 | - | 28 | 4.7 | 0.27 |
| **1660** | Fragile X Alliance | corporate | Australia | 6 | - | 118 | 19.7 | 1.62 |
| **1661** | Government College University Faisalabad | academic | Pakistan | 6 | - | 87 | 14.5 | 1.72 |
| **1662** | Government of Western Australia | government | Australia | 6 | - | 94 | 15.7 | 2.02 |
| **1663** | Australian Government Department of Human Services | government | Australia | 6 | -100 | 86 | 14.3 | 0.98 |
| **1664** | Hanoi National University of Education | academic | Viet Nam | 6 | - | 1627 | 271.2 | 21.18 |
| **1665** | Bhandari Hospital and Research Centre | medical | India | 6 | - | 23 | 3.8 | 0.68 |
| **1666** | Bukkyo University | academic | Japan | 6 | -100 | 23 | 3.8 | 0.19 |
| **1667** | Teerthanker Mahaveer University | academic | India | 6 | - | 20 | 3.3 | 0.21 |
| **1668** | Federal Urdu University of Arts, Science and Technology | academic | Pakistan | 6 | -100 | 95 | 15.8 | 0.78 |
| **1669** | Yulin Normal University | academic | China | 6 | - | 64 | 10.7 | 0.6 |
| **1670** | Gomal University | academic | Pakistan | 6 | - | 4202 | 700.3 | 50.8 |
| **1671** | Asia Eastern University of Science and Technology | academic | Taiwan | 6 | - | 37 | 6.2 | 0.56 |
| **1672** | National Taiwan University of Sport | academic | Taiwan | 6 | 0 | 24 | 4 | 0.62 |
| **1673** | Tao-Yuan General Hospital | medical | Taiwan | 6 | -100 | 53 | 8.8 | 0.62 |
| **1674** | Xizang Minzu University | academic | China | 6 | - | 59 | 9.8 | 0.92 |
| **1675** | Central Institute for Experimental Animals | other | Japan | 6 | 0 | 85 | 14.2 | 1.02 |
| **1676** | Mayo Hospital Lahore | medical | Pakistan | 6 | - | 26 | 4.3 | 0.42 |
| **1677** | Hindu Rao Hospital | medical | India | 6 | - | 74 | 12.3 | 0.78 |
| **1678** | Honam University | academic | South Korea | 6 | - | 84 | 14 | 1.1 |
| **1679** | Tokyo University of Information Sciences | academic | Japan | 6 | - | 26 | 4.3 | 0.21 |
| **1680** | Strand Life Sciences Pvt. Ltd. | corporate | India | 6 | - | 46 | 7.7 | 0.73 |
| **1681** | Guangzhou Sport University | academic | China | 6 | - | 89 | 14.8 | 2.86 |
| **1682** | Ganesh Shankar Vidyarthi Memorial Medical College | academic | India | 6 | 0 | 25 | 4.2 | 0.21 |
| **1683** | CSIR - Institute of Himalayan Bioresource Technology | government | India | 6 | - | 77 | 12.8 | 1.04 |
| **1684** | CSIR - Indian Institute of Chemical Biology | government | India | 6 | -100 | 120 | 20 | 1.31 |
| **1685** | Catholic University of Pusan | academic | South Korea | 6 | - | 23 | 3.8 | 0.41 |
| **1686** | Victorian Infectious Diseases Reference Laboratory | medical | Australia | 6 | - | 71 | 11.8 | 1.54 |
| **1687** | Hanmi Pharmaceutical Co., Ltd. | corporate | South Korea | 6 | - | 14 | 2.3 | 0.37 |
| **1688** | Nippon Sport Science University | academic | Japan | 6 | - | 21 | 3.5 | 0.2 |
| **1689** | Tsukuba International University | academic | Japan | 6 | - | 30 | 5 | 0.56 |
| **1690** | Hokkaido Bunkyo University | academic | Japan | 6 | 0 | 24 | 4 | 0.36 |
| **1691** | Shijonawate Gakuen University | academic | Japan | 6 | 300 | 30 | 5 | 0.45 |
| **1692** | Takarazuka University of Medical and Health Care | academic | Japan | 6 | - | 30 | 5 | 0.49 |
| **1693** | Kinjo University | academic | Japan | 6 | - | 22 | 3.7 | 0.22 |
| **1694** | Gifu Prefectural Tajimi Hospital | medical | Japan | 6 | - | 13 | 2.2 | 0.29 |
| **1695** | Kishiwada Tokushukai Hospital | medical | Japan | 6 | 0 | 61 | 10.2 | 0.85 |
| **1696** | Kidney Health Australia | other | Australia | 6 | 0 | 105 | 17.5 | 2.02 |
| **1697** | Sapporo Kosei General Hospital | medical | Japan | 6 | - | 279 | 46.5 | 2.56 |
| **1698** | Yao Municipal Hospital | medical | Japan | 6 | - | 22 | 3.7 | 0.36 |
| **1699** | Matsushita Memorial Hospital | medical | Japan | 6 | -100 | 13 | 2.2 | 0.28 |
| **1700** | Tokyo Metropolitan Hiroo Hospital | medical | Japan | 6 | - | 30 | 5 | 0.19 |
| **1701** | Tokyo Medical Examiner's Office | government | Japan | 6 | -100 | 35 | 5.8 | 0.35 |
| **1702** | National Hospital Organization Nagara Medical Center | medical | Japan | 6 | 0 | 81 | 13.5 | 1.22 |
| **1703** | National Hospital Organization Shibukawa Medical Center | medical | Japan | 6 | - | 29 | 4.8 | 0.81 |
| **1704** | Osaka Psychiatric Medical Center | medical | Japan | 6 | - | 201 | 33.5 | 1.45 |
| **1705** | Japanese Red Cross Nagasaki Genbaku Hospital | medical | Japan | 6 | -100 | 22 | 3.7 | 0.18 |
| **1706** | Kumamoto Rosai Hospital | medical | Japan | 6 | - | 82 | 13.7 | 0.77 |
| **1707** | Takeda General Hospital | medical | Japan | 6 | - | 60 | 10 | 0.61 |
| **1708** | Miyagi Cancer Center | medical | Japan | 6 | 0 | 137 | 22.8 | 1.03 |
| **1709** | Sapporo Higashi Tokushukai Hospital | medical | Japan | 6 | -100 | 34 | 5.7 | 0.32 |
| **1710** | The Organization for Promoting Neurodevelopmental Disorder Research | other | Japan | 6 | - | 58 | 9.7 | 0.69 |
| **1711** | Fujisawa City Hospital | medical | Japan | 6 | 0 | 85 | 14.2 | 0.52 |
| **1712** | Fremantle Hospital | medical | Australia | 5 | -100 | 133 | 26.6 | 1.09 |
| **1713** | Hunan University of Science and Technology | academic | China | 5 | - | 35 | 7 | 0.87 |
| **1714** | Information Engineering University | academic | China | 5 | -100 | 43 | 8.6 | 0.3 |
| **1715** | Nanjing Agricultural University | academic | China | 5 | - | 59 | 11.8 | 1.58 |
| **1716** | Shanghai Institute of Technology | academic | China | 5 | - | 17 | 3.4 | 0.74 |
| **1717** | Tiangong University | academic | China | 5 | - | 431 | 86.2 | 4.8 |
| **1718** | CAS - Institute of Semiconductors | academic | China | 5 | - | 16 | 3.2 | 0.64 |
| **1719** | Jiangxi Science and Technology Normal University | academic | China | 5 | - | 49 | 9.8 | 0.56 |
| **1720** | China National Petroleum Corporation | corporate | China | 5 | 0 | 111 | 22.2 | 1.43 |
| **1721** | Andhra University | academic | India | 5 | -100 | 118 | 23.6 | 0.99 |
| **1722** | CSIR - Central Food Technological Research Institute India | government | India | 5 | - | 35 | 7 | 0.48 |
| **1723** | CSIR - Indian Institute of Chemical Technology | government | India | 5 | - | 152 | 30.4 | 6.42 |
| **1724** | Indian Institute of Technology Roorkee | academic | India | 5 | - | 27 | 5.4 | 0.78 |
| **1725** | Indian Institute of Technology Guwahati | academic | India | 5 | - | 11 | 2.2 | 0.32 |
| **1726** | Maharshi Dayanand University | academic | India | 5 | - | 54 | 10.8 | 0.77 |
| **1727** | North-Eastern Hill University India | academic | India | 5 | - | 50 | 10 | 0.4 |
| **1728** | Japan Advanced Institute of Science and Technology | academic | Japan | 5 | - | 97 | 19.4 | 1.2 |
| **1729** | National Institute of Health Sciences Tokyo | government | Japan | 5 | - | 50 | 10 | 1.04 |
| **1730** | Tokyo Denki University | academic | Japan | 5 | - | 46 | 9.2 | 0.56 |
| **1731** | Tokyo University of Agriculture | academic | Japan | 5 | - | 139 | 27.8 | 3.03 |
| **1732** | Yokohama National University | academic | Japan | 5 | - | 22 | 4.4 | 0.32 |
| **1733** | Hiroshima City University | academic | Japan | 5 | - | 43 | 8.6 | 0.5 |
| **1734** | Meikai University | academic | Japan | 5 | - | 8 | 1.6 | 0.55 |
| **1735** | Shimadzu Corporation | corporate | Japan | 5 | - | 84 | 16.8 | 1.37 |
| **1736** | Hoseo University | academic | South Korea | 5 | - | 93 | 18.6 | 0.97 |
| **1737** | Korea Institute of Industrial Technology | government | South Korea | 5 | - | 26 | 5.2 | 1.27 |
| **1738** | Pukyong National University | academic | South Korea | 5 | - | 75 | 15 | 0.83 |
| **1739** | Korea Food Research Institute | government | South Korea | 5 | - | 79 | 15.8 | 0.82 |
| **1740** | Kyungsung University | academic | South Korea | 5 | - | 116 | 23.2 | 1.46 |
| **1741** | Suwon University | academic | South Korea | 5 | - | 53 | 10.6 | 0.63 |
| **1742** | Armed Forces Research Institute of Medical Sciences, Thailand | government | Thailand | 5 | - | 31 | 6.2 | 0.9 |
| **1743** | Ming Chuan University | academic | Taiwan | 5 | -100 | 73 | 14.6 | 1.15 |
| **1744** | Universiti Brunei Darussalam | academic | Brunei Darussalam | 5 | - | 66 | 13.2 | 0.86 |
| **1745** | CAS - Wuhan Institute of Virology | academic | China | 5 | - | 117 | 23.4 | 1.71 |
| **1746** | Children's Cancer Institute Australia | academic | Australia | 5 | - | 50 | 10 | 1.75 |
| **1747** | Minzu University of China | academic | China | 5 | -100 | 420 | 84 | 2.47 |
| **1748** | Indian Institute of Science Education and Research Pune | academic | India | 5 | -100 | 55 | 11 | 0.8 |
| **1749** | Sathyabama University | academic | India | 5 | - | 26 | 5.2 | 2.23 |
| **1750** | Indian Institute of Technology Jodhpur | government | India | 5 | 100 | 64 | 12.8 | 0.92 |
| **1751** | National Defence University Malaysia | academic | Malaysia | 5 | - | 11 | 2.2 | 0.34 |
| **1752** | Zhengzhoug University of Light Industry | academic | China | 5 | - | 31 | 6.2 | 0.59 |
| **1753** | Shandong Technology and Business University | academic | China | 5 | - | 78 | 15.6 | 0.72 |
| **1754** | Ministry of Health, Indonesia | government | Indonesia | 5 | - | 3107 | 621.4 | 52.56 |
| **1755** | Liaquat University of Medical and Health Sciences | academic | Pakistan | 5 | - | 7 | 1.4 | 0.42 |
| **1756** | Karunya University | academic | India | 5 | - | 81 | 16.2 | 1.94 |
| **1757** | China University of Petroleum (East China) | academic | China | 5 | - | 15 | 3 | 1.14 |
| **1758** | Shoolini University of Biotechnology and Management Sciences | academic | India | 5 | - | 67 | 13.4 | 1.47 |
| **1759** | Perdana University | academic | Malaysia | 5 | - | 93 | 18.6 | 1 |
| **1760** | Naruto University of Education | academic | Japan | 5 | - | 44 | 8.8 | 0.63 |
| **1761** | Babasaheb Bhimrao Ambedkar University | academic | India | 5 | - | 26 | 5.2 | 0.98 |
| **1762** | Toyota Central R&D Labs., Inc. | corporate | Japan | 5 | -50 | 45 | 9 | 0.95 |
| **1763** | Medical Research Institute of New Zealand | academic | New Zealand | 5 | - | 71 | 14.2 | 2.96 |
| **1764** | Sugiyama Jogakuen University | academic | Japan | 5 | -100 | 102 | 20.4 | 0.65 |
| **1765** | Nagahama Institute of Bio-Science and Technology | academic | Japan | 5 | - | 92 | 18.4 | 0.78 |
| **1766** | Kyushu University of Health and Welfare | academic | Japan | 5 | - | 14 | 2.8 | 0.69 |
| **1767** | Wuhan Sports University | academic | China | 5 | - | 69 | 13.8 | 0.76 |
| **1768** | Sullivan Nicolaides Pathology | medical | Australia | 5 | - | 148 | 29.6 | 1.55 |
| **1769** | Independent University, Bangladesh | academic | Bangladesh | 5 | - | 1 | 0.2 | 0.34 |
| **1770** | St Vincent's Institute of Medical Research | academic | Australia | 5 | - | 130 | 26 | 1.66 |
| **1771** | Academia Sinica - Institute of Linguistics | academic | Taiwan | 5 | -100 | 101 | 20.2 | 1.18 |
| **1772** | Huaiyin Normal University | academic | China | 5 | - | 57 | 11.4 | 0.72 |
| **1773** | Fiji National University | academic | Fiji | 5 | - | 44 | 8.8 | 0.51 |
| **1774** | Charotar University of Science and Technology | academic | India | 5 | - | 3448 | 689.6 | 50.16 |
| **1775** | MEDIPOST Co., Ltd | corporate | South Korea | 5 | -100 | 304 | 60.8 | 2.55 |
| **1776** | Kissei Pharmaceutical Co., Ltd. | corporate | Japan | 5 | - | 57 | 11.4 | 0.97 |
| **1777** | Tsumura and Co. | corporate | Japan | 5 | - | 69 | 13.8 | 1.3 |
| **1778** | Dr. B.R. Ambedkar National Institute of Technology | academic | India | 5 | - | 64 | 12.8 | 1.03 |
| **1779** | Grantham Hospital Hong Kong | medical | Hong Kong | 5 | - | 29 | 5.8 | 2.58 |
| **1780** | Aditya Birla Group | corporate | India | 5 | - | 22 | 4.4 | 0.59 |
| **1781** | Osaka Shoin Women's University | academic | Japan | 5 | - | 161 | 32.2 | 3.11 |
| **1782** | Maharaja Ranjit Singh Punjab Technical University | academic | India | 5 | - | 79 | 15.8 | 1.58 |
| **1783** | Singapore University of Social Sciences | academic | Singapore | 5 | - | 515 | 103 | 2.86 |
| **1784** | Seinan Gakuin University | academic | Japan | 5 | - | 133 | 26.6 | 2.29 |
| **1785** | Chandigarh University | academic | India | 5 | - | 80 | 16 | 2.02 |
| **1786** | Health Services Academy | medical | Pakistan | 5 | - | 6446 | 1289.2 | 108.36 |
| **1787** | Pravara Institute of Medical Sciences | academic | India | 5 | - | 20 | 4 | 0.48 |
| **1788** | Jenderal Soedirman University | academic | Indonesia | 5 | - | 911 | 182.2 | 35.03 |
| **1789** | Malaghan Institute of Medical Research | academic | New Zealand | 5 | - | 81 | 16.2 | 0.9 |
| **1790** | Hubei University of Science and Technology | academic | China | 5 | - | 81 | 16.2 | 0.9 |
| **1791** | Shaoyang University | academic | China | 5 | - | 192 | 38.4 | 2.04 |
| **1792** | National Institute for Biotechnology and Genetic Engineering | academic | Pakistan | 5 | - | 84 | 16.8 | 0.94 |
| **1793** | Tzu Hui Institute of Technology | academic | Taiwan | 5 | - | 66 | 13.2 | 0.84 |
| **1794** | China Aerospace Science and Industry Corporation | government | China | 5 | -100 | 49 | 9.8 | 0.67 |
| **1795** | Woosong University | academic | South Korea | 5 | -100 | 102 | 20.4 | 0.69 |
| **1796** | Shenyang Sport University | academic | China | 5 | - | 12 | 2.4 | 0.96 |
| **1797** | The HIV Netherlands Australia Thailand Research Collaboration | academic | Bangladesh | 5 | -100 | 41 | 8.2 | 0.57 |
| **1798** | Hong Kong Eye Hospital | medical | China | 5 | - | 22 | 4.4 | 0.94 |
| **1799** | Tung Wah Eastern Hospital | medical | Hong Kong | 5 | - | 12 | 2.4 | 1.46 |
| **1800** | Araya Inc. | corporate | Japan | 5 | - | 114 | 22.8 | 11.08 |
| **1801** | Capital University of Physical Education and Sports | academic | China | 5 | - | 45 | 9 | 0.32 |
| **1802** | Morinomiya University of Medical Sciences | academic | Japan | 5 | 0 | 77 | 15.4 | 0.49 |
| **1803** | Advanced Institute of Industrial Technology | academic | Japan | 5 | - | 22 | 4.4 | 0.64 |
| **1804** | CSIR - Indian Institute of Toxicology Research | government | India | 5 | 0 | 85 | 17 | 0.81 |
| **1805** | Motilal Nehru Medical College, Allahabad | medical | India | 5 | - | 15 | 3 | 0.04 |
| **1806** | Gian Sagar Medical College and Hospital | medical | India | 5 | - | 114 | 22.8 | 0.39 |
| **1807** | SNP Genetics, Inc. | corporate | South Korea | 5 | -100 | 73 | 14.6 | 0.57 |
| **1808** | Teikyo University of Science | academic | Japan | 5 | - | 81 | 16.2 | 1.81 |
| **1809** | Nagoya University of Arts and Sciences | academic | Japan | 5 | - | 30 | 6 | 0.97 |
| **1810** | University of Human Arts and Sciences | academic | Japan | 5 | -100 | 42 | 8.4 | 0.34 |
| **1811** | Aichi University of Technology | academic | Japan | 5 | 0 | 26 | 5.2 | 0.51 |
| **1812** | Naragakuen University | academic | Japan | 5 | - | 15 | 3 | 0.6 |
| **1813** | San Beda University | academic | Philippines | 5 | - | 16 | 3.2 | 1.48 |
| **1814** | Shaheed Mohtarma Benazir Bhutto Medical University | academic | Pakistan | 5 | - | 7 | 1.4 | 0.18 |
| **1815** | Rinku General Medical Center | medical | Japan | 5 | - | 33 | 6.6 | 1.47 |
| **1816** | Institute for Health Economics and Policy | other | Japan | 5 | - | 98 | 19.6 | 1.1 |
| **1817** | ARC Centre of Excellence for Nanoscale BioPhotonics | academic | Australia | 5 | - | 112 | 22.4 | 1.53 |
| **1818** | National Hospital Organization Iwakuni Clinical Center | medical | Japan | 5 | - | 9 | 1.8 | 0.11 |
| **1819** | National Hospital Organization Kokura Medical Center | medical | Japan | 5 | -100 | 65 | 13 | 0.87 |
| **1820** | National Hospital Organization Kanmon Medical Center | medical | Japan | 5 | 0 | 28 | 5.6 | 0.49 |
| **1821** | National Hospital Organization Toyohashi Medical Center | medical | Japan | 5 | - | 42 | 8.4 | 0.46 |
| **1822** | Sasebo Chuo Hospital | medical | Japan | 5 | - | 4 | 0.8 | 0.04 |
| **1823** | Suzuka General Hospital | medical | Japan | 5 | - | 37 | 7.4 | 0.42 |
| **1824** | Chulabhorn Graduate Institute | academic | Thailand | 5 | - | 166 | 33.2 | 1.55 |
| **1825** | Miyazaki Medical Association Hospital | medical | Japan | 5 | - | 12 | 2.4 | 0.83 |
| **1826** | KKR Sapporo Medical Center | medical | Japan | 5 | - | 59 | 11.8 | 0.86 |
| **1827** | Kathmandu Medical College | academic | Nepal | 5 | - | 20 | 4 | 0.49 |
| **1828** | Yong In University | academic | South Korea | 5 | - | 58 | 11.6 | 0.72 |
| **1829** | Rajshahi University | academic | Bangladesh | 4 | - | 934 | 233.5 | 44.38 |
| **1830** | China Jiliang University | academic | China | 4 | - | 53 | 13.3 | 0.63 |
| **1831** | China University of Geosciences, Wuhan | academic | China | 4 | - | 29 | 7.3 | 0.65 |
| **1832** | Chongqing Institute of Technology | academic | China | 4 | - | 35 | 8.8 | 0.7 |
| **1833** | Harbin Engineering University | academic | China | 4 | - | 55 | 13.8 | 0.98 |
| **1834** | Heilongjiang University | academic | China | 4 | - | 4 | 1 | 0.2 |
| **1835** | Changzhou University | academic | China | 4 | - | 64 | 16 | 0.94 |
| **1836** | Jimei University | academic | China | 4 | - | 22 | 5.5 | 0.32 |
| **1837** | Lanzhou University of Technology | academic | China | 4 | - | 11 | 2.8 | 0.83 |
| **1838** | Nanjing University of Posts and Telecommunications | academic | China | 4 | - | 78 | 19.5 | 0.98 |
| **1839** | Northwest Normal University | academic | China | 4 | -100 | 40 | 10 | 0.52 |
| **1840** | Shaanxi University of Science and Technology | academic | China | 4 | - | 7 | 1.8 | 0.37 |
| **1841** | Shenyang Aerospace University | academic | China | 4 | - | 3 | 0.8 | 0.03 |
| **1842** | Southwest University of Science and Technology | academic | China | 4 | - | 59 | 14.8 | 1.69 |
| **1843** | Tianjin University of Technology | academic | China | 4 | - | 93 | 23.3 | 1.2 |
| **1844** | University of Science and Technology Beijing | academic | China | 4 | - | 11 | 2.8 | 0.28 |
| **1845** | Zhejiang Sci-Tech University | academic | China | 4 | -100 | 26 | 6.5 | 0.51 |
| **1846** | Anhui Agricultural University | academic | China | 4 | - | 24 | 6 | 1.07 |
| **1847** | CAS - Shanghai Institute of Organic Chemistry | academic | China | 4 | - | 52 | 13 | 0.78 |
| **1848** | Chengdu University of Information Technology | academic | China | 4 | - | 19 | 4.8 | 1 |
| **1849** | Bangalore University | academic | India | 4 | - | 16 | 4 | 0.3 |
| **1850** | Indian Institute of Technology Kharagpur | academic | India | 4 | 100 | 21 | 5.3 | 0.7 |
| **1851** | Jadavpur University | academic | India | 4 | - | 58 | 14.5 | 0.78 |
| **1852** | Jawaharlal Nehru Centre for Advanced Scientific Research | academic | India | 4 | -100 | 106 | 26.5 | 0.83 |
| **1853** | Pondicherry University | academic | India | 4 | - | 19 | 4.8 | 0.41 |
| **1854** | University of Kerala | academic | India | 4 | - | 98 | 24.5 | 1.46 |
| **1855** | Alagappa University | academic | India | 4 | - | 57 | 14.3 | 0.45 |
| **1856** | Deen Dayal Upadhyay Gorakhpur University India | academic | India | 4 | - | 21 | 5.3 | 0.26 |
| **1857** | Dr. Reddy's Laboratories Ltd. | corporate | India | 4 | - | 57 | 14.3 | 0.83 |
| **1858** | Regional Institute of Medical Science India | academic | India | 4 | - | 27 | 6.8 | 0.6 |
| **1859** | Chiba Institute of Technology | academic | Japan | 4 | - | 62 | 15.5 | 1.03 |
| **1860** | Kyoto Pharmaceutical University | academic | Japan | 4 | - | 71 | 17.8 | 1.17 |
| **1861** | Obihiro University of Agriculture and Veterinary Medicine | academic | Japan | 4 | - | 21 | 5.3 | 0.21 |
| **1862** | Gakushuin University | academic | Japan | 4 | - | 91 | 22.8 | 1.84 |
| **1863** | Hokuriku University | academic | Japan | 4 | -100 | 74 | 18.5 | 0.76 |
| **1864** | Josai University | academic | Japan | 4 | - | 23 | 5.8 | 0.49 |
| **1865** | Kyoto Prefectural University | academic | Japan | 4 | - | 21 | 5.3 | 0.17 |
| **1866** | Electronics and Telecommunications Research Institute | government | South Korea | 4 | - | 83 | 20.8 | 1.38 |
| **1867** | Hanbat National University | academic | South Korea | 4 | -100 | 337 | 84.3 | 2.02 |
| **1868** | Korea Institute of Machinery and Materials | government | South Korea | 4 | - | 14 | 3.5 | 0.36 |
| **1869** | Mokpo National University | academic | South Korea | 4 | - | 16 | 4 | 0.39 |
| **1870** | Dongshin University | academic | South Korea | 4 | - | 22 | 5.5 | 0.3 |
| **1871** | Korea Electric Power | corporate | South Korea | 4 | - | 49 | 12.3 | 0.68 |
| **1872** | Silla University | academic | South Korea | 4 | - | 75 | 18.8 | 0.58 |
| **1873** | University of Engineering and Technology Lahore | academic | Pakistan | 4 | - | 177 | 44.3 | 2.58 |
| **1874** | Suranaree University of Technology | academic | Thailand | 4 | - | 36 | 9 | 0.56 |
| **1875** | Mahasarakham University | academic | Thailand | 4 | - | 33 | 8.3 | 0.41 |
| **1876** | Industrial Technology Research Institute Hsinchu | government | Taiwan | 4 | - | 49 | 12.3 | 1.19 |
| **1877** | National Dong Hwa University | academic | Taiwan | 4 | - | 61 | 15.3 | 0.65 |
| **1878** | National Pingtung University of Science and Technology | academic | Taiwan | 4 | 100 | 23 | 5.8 | 0.37 |
| **1879** | Chung Hwa University of Medical Technology | academic | Taiwan | 4 | - | 29 | 7.3 | 1.51 |
| **1880** | National Research Institute of Chinese Medicine Taiwan | academic | Taiwan | 4 | - | 64 | 16 | 1.33 |
| **1881** | Vietnam National University, Hanoi | academic | Viet Nam | 4 | - | 907 | 226.8 | 43.77 |
| **1882** | CAS - Hefei Institutes of Physical Sciences | academic | China | 4 | - | 22 | 5.5 | 0.53 |
| **1883** | University of Ruhuna | academic | Sri Lanka | 4 | -100 | 26 | 6.5 | 0.39 |
| **1884** | Universitas Syiah Kuala | academic | Indonesia | 4 | - | 39 | 9.8 | 0.71 |
| **1885** | Institut Teknologi Sepuluh Nopember | academic | Indonesia | 4 | 100 | 59 | 14.8 | 0.33 |
| **1886** | South Kazakhstan State University (SKSU) | academic | Kazakhstan | 4 | - | 8 | 2 | 0.22 |
| **1887** | Kim Il Sung University | academic | North Korea | 4 | - | 14 | 3.5 | 0.19 |
| **1888** | Indian Institute of Technology Hyderabad | government | India | 4 | - | 22 | 5.5 | 0.6 |
| **1889** | Malaysian Agricultural Research and Development Institute | academic | Malaysia | 4 | - | 14 | 3.5 | 1.35 |
| **1890** | Azerbaijan State Advanced Training Institute for Doctors | academic | Azerbaijan | 4 | - | 50 | 12.5 | 1.04 |
| **1891** | Heilongjiang Bayi Agricultural University | academic | China | 4 | -100 | 105 | 26.3 | 1.27 |
| **1892** | Sichuan University of Science & Engineering | academic | China | 4 | - | 105 | 26.3 | 1.22 |
| **1893** | DENSO Corporation | corporate | Japan | 4 | - | 41 | 10.3 | 0.41 |
| **1894** | Radiation Effects Research Foundation Hiroshima | other | Japan | 4 | - | 27 | 6.8 | 0.66 |
| **1895** | Japan Anti-Tuberculosis Association | medical | Japan | 4 | -100 | 29 | 7.3 | 0.57 |
| **1896** | Guru Gobind Singh Indraprastha University | academic | India | 4 | 0 | 3 | 0.8 | 0.03 |
| **1897** | Chongqing University of Science and Technology | academic | China | 4 | - | 18 | 4.5 | 0.57 |
| **1898** | Thapar University | academic | India | 4 | - | 35 | 8.8 | 0.6 |
| **1899** | Mae Fah Luang University | academic | Thailand | 4 | - | 23 | 5.8 | 0.56 |
| **1900** | Birla Institute of Technology, Mesra | academic | India | 4 | - | 206 | 51.5 | 1.88 |
| **1901** | Academia Sinica - Institute of Cellular and Organismic Biology | academic | Taiwan | 4 | - | 98 | 24.5 | 1.23 |
| **1902** | Penang Medical College | academic | Malaysia | 4 | - | 54 | 13.5 | 0.82 |
| **1903** | Matsuyama University | academic | Japan | 4 | -100 | 21 | 5.3 | 0.2 |
| **1904** | Central University of Karnataka | academic | India | 4 | - | 23 | 5.8 | 0.6 |
| **1905** | Shandong University of Technology | academic | China | 4 | - | 94 | 23.5 | 1.27 |
| **1906** | Kokushikan University | academic | Japan | 4 | - | 17 | 4.3 | 0.1 |
| **1907** | United International College (UIC) | academic | China | 4 | - | 42 | 10.5 | 0.96 |
| **1908** | New Zealand Rugby | government | New Zealand | 4 | - | 16 | 4 | 0.7 |
| **1909** | Lakes District Health Board | government | New Zealand | 4 | - | 4 | 1 | 0.16 |
| **1910** | Macrogen Inc | corporate | South Korea | 4 | -100 | 127 | 31.8 | 0.84 |
| **1911** | Indian Institute of Science Education and Research, Kolkata | academic | India | 4 | - | 149 | 37.3 | 1.19 |
| **1912** | Guangdong University of Finance | academic | China | 4 | - | 6 | 1.5 | 0.43 |
| **1913** | Ube Industries Ltd | corporate | Japan | 4 | 0 | 0 | 0 | 0 |
| **1914** | Ministry of Health, New Zealand | government | New Zealand | 4 | -100 | 267 | 66.8 | 1.88 |
| **1915** | Karachi Medical and Dental College | medical | Pakistan | 4 | - | 14 | 3.5 | 0.22 |
| **1916** | Justice Health and Forensic Mental Health Network | government | Australia | 4 | - | 96 | 24 | 1.04 |
| **1917** | The Open University of Japan | academic | Japan | 4 | - | 18 | 4.5 | 0.36 |
| **1918** | CHA Medical Group | medical | South Korea | 4 | -100 | 19 | 4.8 | 0.33 |
| **1919** | Sime Darby Berhad | corporate | Malaysia | 4 | - | 31 | 7.8 | 0.58 |
| **1920** | University of Management and Technology | academic | Pakistan | 4 | - | 806 | 201.5 | 14.34 |
| **1921** | Jissen Women's University | academic | Japan | 4 | - | 4 | 1 | 0.58 |
| **1922** | Australian and New Zealand Intensive Care Society | medical | Australia | 4 | - | 68 | 17 | 1.18 |
| **1923** | Kazakh State Women's Pedagogical University | academic | Kazakhstan | 4 | - | 15 | 3.8 | 0.36 |
| **1924** | GLA University | academic | India | 4 | - | 23 | 5.8 | 0.71 |
| **1925** | Korea National Sport University | academic | South Korea | 4 | - | 134 | 33.5 | 2.66 |
| **1926** | M.S. Ramaiah University of Applied Sciences | academic | India | 4 | - | 36 | 9 | 0.26 |
| **1927** | Shanghai Institute of Planned Parenthood Research | government | China | 4 | - | 68 | 17 | 1.7 |
| **1928** | Jiangsu Institute of Nuclear Medicine | government | China | 4 | - | 59 | 14.8 | 0.97 |
| **1929** | PLA No. 117 Hospital | medical | China | 4 | - | 143 | 35.8 | 2.07 |
| **1930** | PLA No. 85 Hospital | medical | China | 4 | - | 51 | 12.8 | 0.58 |
| **1931** | National Institute of Science Education and Research | academic | India | 4 | 0 | 36 | 9 | 0.37 |
| **1932** | Raja Isteri Pengiran Anak Saleha Hospital | medical | Brunei Darussalam | 4 | - | 46 | 11.5 | 0.75 |
| **1933** | Tianjin Eye Hospital | medical | China | 4 | - | 38 | 9.5 | 0.82 |
| **1934** | Dr. A.L.M. Postgraduate Institute of Basic Medical Sciences | academic | India | 4 | - | 121 | 30.3 | 1.28 |
| **1935** | Taiho Pharmaceutical Co., Ltd. | corporate | Japan | 4 | - | 28 | 7 | 0.63 |
| **1936** | Galgotias University | academic | India | 4 | - | 107 | 26.8 | 1.63 |
| **1937** | Cenderawasih University | academic | Indonesia | 4 | - | 10 | 2.5 | 0.18 |
| **1938** | Shaukat Khanum Memorial Cancer Hospital and Research Centre | medical | Pakistan | 4 | - | 48 | 12 | 1.53 |
| **1939** | International Institute for Population Sciences | academic | India | 4 | - | 4869 | 1217.3 | 105.5 |
| **1940** | Services Institute of Medical Sciences Lahore | medical | Pakistan | 4 | 0 | 47 | 11.8 | 0.82 |
| **1941** | Otemon Gakuin University | academic | Japan | 4 | - | 19 | 4.8 | 0.69 |
| **1942** | Kangnam University | academic | South Korea | 4 | - | 27 | 6.8 | 0.97 |
| **1943** | Hokkaido Information University | academic | Japan | 4 | - | 27 | 6.8 | 0.55 |
| **1944** | Bunkyo University | academic | Japan | 4 | -100 | 59 | 14.8 | 1.03 |
| **1945** | APJ Abdul Kalam Technological University | academic | India | 4 | - | 62 | 15.5 | 1.07 |
| **1946** | M.Kh. Dulaty Taraz State University | academic | Kazakhstan | 4 | - | 0 | 0 | 0 |
| **1947** | Eastern University, Sri Lanka | academic | Sri Lanka | 4 | - | 39 | 9.8 | 0.35 |
| **1948** | Shenzhen Eye Hospital | medical | China | 4 | - | 10 | 2.5 | 0.55 |
| **1949** | Yan Chai Hospital | medical | Hong Kong | 4 | - | 153 | 38.3 | 2.24 |
| **1950** | Kwai Chung Hospital | medical | Hong Kong | 4 | - | 17 | 4.3 | 0.19 |
| **1951** | Jilin Central General Hospital | medical | China | 4 | - | 181 | 45.3 | 4.96 |
| **1952** | PLA No. 210 Hospital | medical | China | 4 | -100 | 108 | 27 | 1.47 |
| **1953** | Jaipur National University | academic | India | 4 | - | 0 | 0 | 0 |
| **1954** | Nguyen Tat Thanh University | academic | Viet Nam | 4 | - | 4370 | 1092.5 | 107.42 |
| **1955** | National Institute for Research in Tuberculosis | academic | India | 4 | - | 86 | 21.5 | 1.48 |
| **1956** | Nippon Shinyaku Co., Ltd. | corporate | Japan | 4 | -100 | 78 | 19.5 | 1.54 |
| **1957** | Santosh Deemed to be University | academic | India | 4 | - | 1 | 0.3 | 0.01 |
| **1958** | Base Hospital Delhi Cantt | medical | India | 4 | 0 | 15 | 3.8 | 0.21 |
| **1959** | Taiyuan Iron and Steel (Group) Company Ltd. | corporate | China | 4 | -100 | 26 | 6.5 | 0.2 |
| **1960** | Dosmukhamedov Atyrau University | academic | Kazakhstan | 4 | - | 12 | 3 | 0.29 |
| **1961** | Sakakibara Heart Institute | medical | Japan | 4 | - | 1 | 0.3 | 0 |
| **1962** | Daiichi University of Pharmacy | academic | Japan | 4 | - | 18 | 4.5 | 0.24 |
| **1963** | Komatsu University | academic | Japan | 4 | - | 74 | 18.5 | 2.09 |
| **1964** | Bunkyo Gakuin University | academic | Japan | 4 | - | 59 | 14.8 | 1.01 |
| **1965** | Nagoya Gakuin University | academic | Japan | 4 | - | 10 | 2.5 | 0.16 |
| **1966** | Sagami Women's University | academic | Japan | 4 | -100 | 130 | 32.5 | 1.56 |
| **1967** | Ibaraki Christian University | academic | Japan | 4 | - | 54 | 13.5 | 0.63 |
| **1968** | The Cardiovascular Institute | medical | Japan | 4 | - | 29 | 7.3 | 0.74 |
| **1969** | Osaka Center for Cancer and Cardiovascular Disease Prevention | medical | Japan | 4 | - | 33 | 8.3 | 0.64 |
| **1970** | Health Science University, Yamanashi | academic | Japan | 4 | - | 71 | 17.8 | 1.35 |
| **1971** | Akita Industrial Technology Center | government | Japan | 4 | -100 | 126 | 31.5 | 0.9 |
| **1972** | National Stroke Foundation, Australia | other | Australia | 4 | -100 | 109 | 27.3 | 1.36 |
| **1973** | TAFE Queensland | government | Australia | 4 | - | 5 | 1.3 | 0.21 |
| **1974** | Okinawa Chubu Hospital | medical | Japan | 4 | -100 | 28 | 7 | 0.29 |
| **1975** | Cancer Institute India | academic | India | 4 | - | 48 | 12 | 1.21 |
| **1976** | RPH Research Foundation | other | Australia | 4 | -100 | 708 | 177 | 5.74 |
| **1977** | Nepal Health Research Council | government | Nepal | 4 | - | 1692 | 423 | 57.87 |
| **1978** | Chemo-Sero-Therapeutic Research Institute | other | Japan | 4 | - | 62 | 15.5 | 0.64 |
| **1979** | National Hospital Organization Tochigi Medical Center | medical | Japan | 4 | - | 17 | 4.3 | 0.7 |
| **1980** | Japanese Red Cross Toyama Hospital | medical | Japan | 4 | - | 35 | 8.8 | 0.25 |
| **1981** | Japanese Red Cross Yamaguchi Hospital | medical | Japan | 4 | - | 28 | 7 | 0.39 |
| **1982** | Hamamatsu Rosai Hospital | medical | Japan | 4 | -100 | 21 | 5.3 | 0.91 |
| **1983** | Aomori Rosai Hospital | medical | Japan | 4 | -100 | 7 | 1.8 | 0.12 |
| **1984** | Japanese Red Cross Kyushu International College of Nursing | academic | Japan | 4 | - | 4 | 1 | 0.23 |
| **1985** | Vietnam National Children's Hospital | medical | Viet Nam | 4 | - | 22 | 5.5 | 0.82 |
| **1986** | Jikei University of Health Care Sciences | academic | Japan | 4 | -100 | 99 | 24.8 | 0.96 |
| **1987** | Shenzhen Bay Laboratory | government | China | 4 | - | 18 | 4.5 | 1.21 |
| **1988** | University of Okara | academic | Pakistan | 4 | - | 22 | 5.5 | 0.55 |
| **1989** | Sir Salimullah Medical College | medical | Bangladesh | 4 | - | 19 | 4.8 | 2.4 |
| **1990** | Defence Science & Technology Group | government | Australia | 3 | -100 | 49 | 16.3 | 0.9 |
| **1991** | South Australian Research and Development Institute | government | Australia | 3 | - | 32 | 10.7 | 0.43 |
| **1992** | Changsha University of Science and Technology | academic | China | 3 | - | 29 | 9.7 | 0.86 |
| **1993** | China West Normal University | academic | China | 3 | -100 | 35 | 11.7 | 0.58 |
| **1994** | Donghua University | academic | China | 3 | - | 21 | 7 | 0.68 |
| **1995** | Jiangsu Ocean University | academic | China | 3 | - | 28 | 9.3 | 1.31 |
| **1996** | Nanjing Forestry University | academic | China | 3 | - | 133 | 44.3 | 1.68 |
| **1997** | Nanjing Tech University | academic | China | 3 | - | 87 | 29 | 1.02 |
| **1998** | North China Electric Power University | academic | China | 3 | - | 18 | 6 | 1.32 |
| **1999** | North University of China | academic | China | 3 | - | 105 | 35 | 2.04 |
| **2000** | Qingdao University of Science and Technology | academic | China | 3 | - | 48 | 16 | 1.19 |
| **2001** | Shanxi Normal University | academic | China | 3 | - | 15 | 5 | 0.77 |
| **2002** | South China Agricultural University | academic | China | 3 | - | 121 | 40.3 | 2.06 |
| **2003** | PLA No. 302 Hospital | medical | China | 3 | - | 43 | 14.3 | 0.85 |
| **2004** | Beijing Technology and Business University | academic | China | 3 | - | 25 | 8.3 | 0.82 |
| **2005** | CAS - Institute of Microbiology | academic | China | 3 | - | 64 | 21.3 | 1.54 |
| **2006** | Chongqing Technology and Business University | academic | China | 3 | 0 | 91 | 30.3 | 1.67 |
| **2007** | Guangdong Ocean University | academic | China | 3 | - | 48 | 16 | 0.84 |
| **2008** | Shandong Jianzhu University | academic | China | 3 | - | 12 | 4 | 2.14 |
| **2009** | Shanghai University of Finance and Economics | academic | China | 3 | - | 30 | 10 | 0.51 |
| **2010** | Wuhan Polytechnic University | academic | China | 3 | -100 | 113 | 37.7 | 1.47 |
| **2011** | Xinyang Normal University | academic | China | 3 | - | 139 | 46.3 | 1.26 |
| **2012** | Bose Institute | academic | India | 3 | - | 50 | 16.7 | 0.51 |
| **2013** | Cochin University of Science and Technology | academic | India | 3 | -100 | 51 | 17 | 0.38 |
| **2014** | PSG College of Technology India | academic | India | 3 | - | 38 | 12.7 | 0.66 |
| **2015** | Saha Institute of Nuclear Physics | academic | India | 3 | - | 29 | 9.7 | 0.59 |
| **2016** | Devi Ahilya University of Indore | academic | India | 3 | - | 15 | 5 | 0.87 |
| **2017** | Gauhati University | academic | India | 3 | - | 3 | 1 | 0.04 |
| **2018** | International Institute of Information Technology Hyderabad | academic | India | 3 | - | 123 | 41 | 2.17 |
| **2019** | National Centre for Cell Science | government | India | 3 | 0 | 110 | 36.7 | 1.36 |
| **2020** | National Institute of Cholera and Enteric Diseases India | government | India | 3 | - | 38 | 12.7 | 0.57 |
| **2021** | National Institute of Immunology India | government | India | 3 | -100 | 24 | 8 | 0.46 |
| **2022** | National Institute of Technology Tiruchirappalli | academic | India | 3 | - | 26 | 8.7 | 0.72 |
| **2023** | Japan Aerospace Exploration Agency | government | Japan | 3 | - | 45 | 15 | 0.81 |
| **2024** | Kitami Institute of Technology | academic | Japan | 3 | -100 | 99 | 33 | 1.25 |
| **2025** | National Institute for Materials Science Tsukuba | government | Japan | 3 | - | 236 | 78.7 | 3.53 |
| **2026** | Research Organization of Information and Systems, National Institute of Genetics Mishima | academic | Japan | 3 | - | 283 | 94.3 | 6.71 |
| **2027** | Nippon Dental University | academic | Japan | 3 | - | 17 | 5.7 | 0.28 |
| **2028** | Osaka Electro-Communication University | academic | Japan | 3 | -100 | 14 | 4.7 | 0.61 |
| **2029** | Fujitsu | corporate | Japan | 3 | - | 42 | 14 | 2.77 |
| **2030** | Hitotsubashi University | academic | Japan | 3 | - | 1 | 0.3 | 0.06 |
| **2031** | Japan Women's University | academic | Japan | 3 | - | 9 | 3 | 0.48 |
| **2032** | Kanto Gakuin University | academic | Japan | 3 | - | 3 | 1 | 0.92 |
| **2033** | Kyoto Sangyo University | academic | Japan | 3 | - | 18 | 6 | 0.33 |
| **2034** | Niigata University of Pharmacy and Applied Life Sciences | academic | Japan | 3 | - | 85 | 28.3 | 1.26 |
| **2035** | Saitama Cancer Center | medical | Japan | 3 | - | 54 | 18 | 1.08 |
| **2036** | Toyota Motor | corporate | Japan | 3 | - | 23 | 7.7 | 0.46 |
| **2037** | Hongik University | academic | South Korea | 3 | - | 28 | 9.3 | 0.68 |
| **2038** | Korea Research Institute of Chemical Technology | government | South Korea | 3 | - | 98 | 32.7 | 2.02 |
| **2039** | University of Seoul | academic | South Korea | 3 | - | 57 | 19 | 0.72 |
| **2040** | Universiti Malaysia Perlis | academic | Malaysia | 3 | - | 29 | 9.7 | 0.49 |
| **2041** | Singapore Management University | academic | Singapore | 3 | - | 10 | 3.3 | 0.72 |
| **2042** | DSO National Laboratory, Singapore | government | Singapore | 3 | -100 | 63 | 21 | 0.83 |
| **2043** | Feng Chia University | academic | Taiwan | 3 | - | 27 | 9 | 1.07 |
| **2044** | National Defense University Taiwan | academic | Taiwan | 3 | -100 | 9 | 3 | 0.17 |
| **2045** | National University of Tainan Taiwan | academic | Taiwan | 3 | - | 9 | 3 | 0.2 |
| **2046** | Shu-Te University | academic | Taiwan | 3 | - | 18 | 6 | 0.31 |
| **2047** | Te Whare Wananga o Awanuiarangi | academic | New Zealand | 3 | -100 | 39 | 13 | 0.47 |
| **2048** | Universitas Andalas | academic | Indonesia | 3 | - | 17 | 5.7 | 0.45 |
| **2049** | University of Nottingham Ningbo China | academic | China | 3 | - | 37 | 12.3 | 2.03 |
| **2050** | SASTRA | academic | India | 3 | - | 100 | 33.3 | 2.04 |
| **2051** | Panasonic Holdings Corporation | corporate | Japan | 3 | -100 | 83 | 27.7 | 1.3 |
| **2052** | Fukuoka Prefectural University | academic | Japan | 3 | - | 14 | 4.7 | 0.26 |
| **2053** | Zhongnan University of Economics and Law | academic | China | 3 | - | 116 | 38.7 | 1.22 |
| **2054** | Lahore University of Management Sciences | academic | Pakistan | 3 | - | 31 | 10.3 | 0.96 |
| **2055** | Indian Institute of Technology Indore | academic | India | 3 | - | 17 | 5.7 | 0.73 |
| **2056** | Teijin Ltd. | corporate | Japan | 3 | - | 55 | 18.3 | 0.99 |
| **2057** | Indian Institute of Technology Mandi | academic | India | 3 | - | 16 | 5.3 | 0.41 |
| **2058** | Sultan Idris Education University | academic | Malaysia | 3 | - | 130 | 43.3 | 1.4 |
| **2059** | Jilin Agricultural University | academic | China | 3 | - | 36 | 12 | 0.61 |
| **2060** | Zhejiang University of Finance and Economics | academic | China | 3 | - | 17 | 5.7 | 0.09 |
| **2061** | Yancheng Institute of Technology | academic | China | 3 | - | 53 | 17.7 | 0.76 |
| **2062** | Chiba Institute of Science | academic | Japan | 3 | - | 41 | 13.7 | 0.84 |
| **2063** | Manonmaniam Sundaranar University | academic | India | 3 | 0 | 774 | 258 | 21.1 |
| **2064** | Khulna University | academic | Bangladesh | 3 | - | 24 | 8 | 0.97 |
| **2065** | Aletheia University | academic | Taiwan | 3 | 0 | 33 | 11 | 0.8 |
| **2066** | Manav Rachna International University | academic | India | 3 | - | 8 | 2.7 | 0.17 |
| **2067** | Kumaun University India | academic | India | 3 | - | 58 | 19.3 | 1.79 |
| **2068** | University of Sargodha | academic | Pakistan | 3 | - | 81 | 27 | 10.48 |
| **2069** | Senshu University | academic | Japan | 3 | - | 36 | 12 | 0.75 |
| **2070** | Indian Institute of Management Bangalore | academic | India | 3 | - | 20 | 6.7 | 0.27 |
| **2071** | Australian Institute of Sport | government | Australia | 3 | - | 35 | 11.7 | 0.58 |
| **2072** | Hyogo University of Teacher Education | academic | Japan | 3 | -100 | 51 | 17 | 0.64 |
| **2073** | Gujarat University | academic | India | 3 | - | 30 | 10 | 1.03 |
| **2074** | Indian Institute of Science Education and Research Bhopal | academic | India | 3 | - | 18 | 6 | 0.67 |
| **2075** | Universitas Muhammadiyah Malang | academic | Indonesia | 3 | - | 30 | 10 | 2.08 |
| **2076** | Academia Sinica - Institute of Information Science | academic | Taiwan | 3 | -100 | 28 | 9.3 | 0.53 |
| **2077** | Academia Sinica - Institute of Mathematics | academic | Taiwan | 3 | - | 267 | 89 | 5.29 |
| **2078** | Academia Sinica - Institute of Sociology | academic | Taiwan | 3 | - | 222 | 74 | 3.08 |
| **2079** | SEGi University | academic | Malaysia | 3 | - | 15 | 5 | 0.38 |
| **2080** | Prefectural University of Kumamoto | academic | Japan | 3 | - | 35 | 11.7 | 0.39 |
| **2081** | Kinjo Gakuin University | academic | Japan | 3 | -100 | 85 | 28.3 | 1.1 |
| **2082** | State Islamic University Of Malang | academic | Indonesia | 3 | - | 16 | 5.3 | 1.6 |
| **2083** | Fakir Mohan University | academic | India | 3 | - | 29 | 9.7 | 3.3 |
| **2084** | Beijing Language and Culture University | academic | China | 3 | - | 29 | 9.7 | 1.4 |
| **2085** | Shih Chien University | academic | Taiwan | 3 | -100 | 99 | 33 | 1.08 |
| **2086** | Maulana Abul Kalam Azad University of Technology | academic | India | 3 | - | 0 | 0 | 0 |
| **2087** | Cumberland Hospital | medical | Australia | 3 | 0 | 86 | 28.7 | 3.45 |
| **2088** | Indira Gandhi National Open University | academic | India | 3 | - | 3 | 1 | 0.79 |
| **2089** | Ming Chi University of Technology | academic | Taiwan | 3 | - | 7 | 2.3 | 0.04 |
| **2090** | Hainan Normal University | academic | China | 3 | -100 | 24 | 8 | 0.59 |
| **2091** | Australian Genome Research Facility Ltd. | corporate | Australia | 3 | - | 65 | 21.7 | 1.94 |
| **2092** | Tencent | corporate | China | 3 | - | 45 | 15 | 2.43 |
| **2093** | Qinghai Normal University | academic | China | 3 | - | 5 | 1.7 | 0.96 |
| **2094** | University of Balochistan | academic | Pakistan | 3 | -100 | 5 | 1.7 | 0.06 |
| **2095** | Tibet University | academic | China | 3 | 100 | 50 | 16.7 | 2.02 |
| **2096** | KYORIN Pharmaceutical Co., Ltd. | corporate | Japan | 3 | - | 2 | 0.7 | 0.16 |
| **2097** | Mochida Pharmaceutical Co, Ltd. | corporate | Japan | 3 | - | 40 | 13.3 | 0.89 |
| **2098** | TauRX Therapeutics Ltd | corporate | Singapore | 3 | - | 36 | 12 | 1.31 |
| **2099** | Shiv Nadar University | academic | India | 3 | - | 106 | 35.3 | 2.36 |
| **2100** | National Institute of Malaria Research India | academic | India | 3 | - | 17 | 5.7 | 0.46 |
| **2101** | Chittagong University of Engineering and Technology | academic | Bangladesh | 3 | - | 24 | 8 | 0.83 |
| **2102** | Jilin Medical College | academic | China | 3 | -100 | 65 | 21.7 | 0.86 |
| **2103** | Dr. A.P.J. Abdul Kalam Technical University | academic | India | 3 | -50 | 46 | 15.3 | 0.63 |
| **2104** | Duksung Women's University | academic | South Korea | 3 | - | 54 | 18 | 1.13 |
| **2105** | Chhattisgarh Swami Vivekanand Technical University | academic | India | 3 | -100 | 43 | 14.3 | 0.78 |
| **2106** | Maharashtra University of Health Sciences | academic | India | 3 | -100 | 87 | 29 | 1.06 |
| **2107** | Parul University | academic | India | 3 | - | 16 | 5.3 | 0.62 |
| **2108** | Faisalabad Medical University | medical | Pakistan | 3 | - | 9 | 3 | 0.25 |
| **2109** | University of Malakand | academic | Pakistan | 3 | - | 63 | 21 | 2.79 |
| **2110** | Daito Bunka University | academic | Japan | 3 | -100 | 39 | 13 | 1.58 |
| **2111** | Meenakshi Academy of Higher Education and Research | academic | India | 3 | - | 16 | 5.3 | 0.7 |
| **2112** | Westlake University | academic | China | 3 | - | 16 | 5.3 | 1.52 |
| **2113** | Phenikaa University | academic | Viet Nam | 3 | - | 26 | 8.7 | 2.15 |
| **2114** | Hanshin University | academic | South Korea | 3 | - | 17 | 5.7 | 0.49 |
| **2115** | Panzhihua University | academic | China | 3 | - | 8 | 2.7 | 0.71 |
| **2116** | Hong Kong Shue Yan University | academic | China | 3 | - | 53 | 17.7 | 1.94 |
| **2117** | Hunan First Normal University | academic | China | 3 | - | 57 | 19 | 1.22 |
| **2118** | Guizhou University of Traditional Chinese Medicine | academic | China | 3 | - | 53 | 17.7 | 0.91 |
| **2119** | Guangxi Center for Disease Prevention and Control | government | China | 3 | - | 25 | 8.3 | 0.65 |
| **2120** | PLA No. 303 Hospital | medical | China | 3 | -100 | 56 | 18.7 | 0.95 |
| **2121** | Nanjing Xiaozhuang College | academic | China | 3 | - | 13 | 4.3 | 0.32 |
| **2122** | Hazara University | academic | Pakistan | 3 | - | 13 | 4.3 | 0.44 |
| **2123** | University of Gujrat | academic | Pakistan | 3 | - | 18 | 6 | 0.45 |
| **2124** | Shahjalal University of Science and Technology | academic | Bangladesh | 3 | - | 24 | 8 | 0.52 |
| **2125** | Sriwijaya University | academic | Indonesia | 3 | - | 3 | 1 | 0.18 |
| **2126** | National Tainan Junior College of Nursing | academic | Taiwan | 3 | - | 14 | 4.7 | 0.29 |
| **2127** | Cardinal Tien Junior College of Healthcare and Management | academic | Taiwan | 3 | - | 33 | 11 | 0.76 |
| **2128** | Ping Tung Christian Hospital, Taiwan | medical | Taiwan | 3 | - | 9 | 3 | 0.17 |
| **2129** | Feng-Yuan Hospital | medical | Taiwan | 3 | - | 19 | 6.3 | 0.59 |
| **2130** | Dhaka Medical College and Hospital | medical | Bangladesh | 3 | - | 13 | 4.3 | 0.39 |
| **2131** | University of Asia Pacific | academic | Bangladesh | 3 | - | 62 | 20.7 | 0.61 |
| **2132** | Kiang Wu Nursing College of Macau | academic | Macao | 3 | - | 42 | 14 | 1.07 |
| **2133** | Niigata Cancer Center Hospital | medical | Japan | 3 | - | 27 | 9 | 0.41 |
| **2134** | Rajavithi Hospital | medical | Thailand | 3 | - | 62 | 20.7 | 0.7 |
| **2135** | Namseoul University | academic | South Korea | 3 | - | 19 | 6.3 | 0.31 |
| **2136** | Calcutta School of Tropical Medicine | academic | India | 3 | -100 | 39 | 13 | 0.54 |
| **2137** | Joongbu University | academic | South Korea | 3 | - | 70 | 23.3 | 1.11 |
| **2138** | Islamia College Peshawar | academic | Pakistan | 3 | - | 11 | 3.7 | 0.56 |
| **2139** | Haiphong University | academic | Viet Nam | 3 | - | 15 | 5 | 0.78 |
| **2140** | Baqai Medical University | medical | Pakistan | 3 | - | 154 | 51.3 | 2.44 |
| **2141** | Okinawa International University | academic | Japan | 3 | - | 34 | 11.3 | 0.72 |
| **2142** | Sax Institute | other | Australia | 3 | -100 | 127 | 42.3 | 2.01 |
| **2143** | YARSI University | academic | Indonesia | 3 | - | 17 | 5.7 | 1.12 |
| **2144** | Shenzhen Technology University | academic | China | 3 | - | 15 | 5 | 0.63 |
| **2145** | Jilin Provincial Cancer Hospital | medical | China | 3 | - | 32 | 10.7 | 0.42 |
| **2146** | Ruttonjee Hospital | medical | Hong Kong | 3 | - | 26 | 8.7 | 0.77 |
| **2147** | PLA No. 211 Hospital | medical | China | 3 | - | 111 | 37 | 1.62 |
| **2148** | Al-Shifa Trust Eye Hospital | medical | Pakistan | 3 | - | 4338 | 1446 | 141.29 |
| **2149** | Rawalpindi Medical College | academic | Pakistan | 3 | - | 8 | 2.7 | 0.34 |
| **2150** | National Center for AIDS/STD Control and Prevention | government | China | 3 | - | 15 | 5 | 0.4 |
| **2151** | KLS Gogte Institute of Technology | academic | India | 3 | - | 26 | 8.7 | 0.84 |
| **2152** | CSIR - National Chemical Laboratory | government | India | 3 | - | 26 | 8.7 | 1.05 |
| **2153** | Bharat Electronics | corporate | India | 3 | - | 4 | 1.3 | 0.06 |
| **2154** | Islamic University, Kushtia | academic | Bangladesh | 3 | - | 24 | 8 | 0.93 |
| **2155** | NIMS University | academic | India | 3 | - | 4 | 1.3 | 0.27 |
| **2156** | Ulsan College | academic | South Korea | 3 | - | 65 | 21.7 | 1.12 |
| **2157** | Gunma Paz University | academic | Japan | 3 | - | 109 | 36.3 | 1.79 |
| **2158** | Jashore University of Science and Technology | academic | Bangladesh | 3 | - | 22 | 7.3 | 0.72 |
| **2159** | Bangladesh University of Health Sciences | academic | Bangladesh | 3 | - | 21 | 7 | 0.48 |
| **2160** | National Cancer Institute Sri Lanka | government | Sri Lanka | 3 | - | 11 | 3.7 | 0.57 |
| **2161** | Academy of Public Administration under the President of the Republic of Kazakhstan | academic | Kazakhstan | 3 | - | 2 | 0.7 | 0.07 |
| **2162** | Korkyt Ata Kyzylorda State University | academic | Kazakhstan | 3 | - | 1 | 0.3 | 0.03 |
| **2163** | Mitsubishi Kyoto Hospital | medical | Japan | 3 | - | 18 | 6 | 0.87 |
| **2164** | Sendai Kousei Hospital | medical | Japan | 3 | - | 30 | 10 | 0.68 |
| **2165** | Mount Fuji Research Institute | government | Japan | 3 | - | 12 | 4 | 0.56 |
| **2166** | Teijin Pharma Limited | corporate | Japan | 3 | - | 59 | 19.7 | 1.23 |
| **2167** | National Institute for Minamata Disease | government | Japan | 3 | - | 52 | 17.3 | 0.84 |
| **2168** | Aomori University | academic | Japan | 3 | - | 2 | 0.7 | 0.04 |
| **2169** | Osaka Kawasaki Rehabilitation University | academic | Japan | 3 | - | 12 | 4 | 0.71 |
| **2170** | Seitoku University | academic | Japan | 3 | - | 11 | 3.7 | 0.17 |
| **2171** | Japan Women's College of Physical Education | academic | Japan | 3 | 100 | 71 | 23.7 | 0.67 |
| **2172** | Hokusho University | academic | Japan | 3 | - | 21 | 7 | 0.51 |
| **2173** | Kyoto Industrial Health Association | other | Japan | 3 | - | 26 | 8.7 | 0.33 |
| **2174** | Nishikyushu University | academic | Japan | 3 | - | 18 | 6 | 0.2 |
| **2175** | Matsumoto University | academic | Japan | 3 | - | 14 | 4.7 | 0.19 |
| **2176** | Osaka University of Health and Sport Sciences | academic | Japan | 3 | - | 17 | 5.7 | 0.32 |
| **2177** | Kyushu University of Nursing and Social Welfare | academic | Japan | 3 | - | 15 | 5 | 0.14 |
| **2178** | Ryotokuji University | academic | Japan | 3 | - | 9 | 3 | 1.53 |
| **2179** | Gunma Prefectural Cancer Center | medical | Japan | 3 | - | 14 | 4.7 | 0.69 |
| **2180** | Shaheed Mohtarma Benazir Bhutto Medical College | academic | Pakistan | 3 | - | 2 | 0.7 | 0.12 |
| **2181** | Japan Ministry of Land, Infrastructure and Transport | government | Japan | 3 | - | 2 | 0.7 | 0.07 |
| **2182** | Japan Ministry of Health, Labour and Welfare | government | Japan | 3 | -100 | 66 | 22 | 1.48 |
| **2183** | Public Health Concern Trust, Nepal | other | Nepal | 3 | - | 44 | 14.7 | 0.64 |
| **2184** | CRECON Medical Assessment Inc. | corporate | Japan | 3 | - | 8 | 2.7 | 0.38 |
| **2185** | Institute for Health Outcomes and Process Evaluation Research (iHope International) | other | Japan | 3 | - | 28 | 9.3 | 1.16 |
| **2186** | Rajamangala University of Technology Suvarnabhumi | academic | Thailand | 3 | - | 36 | 12 | 1.49 |
| **2187** | ARC Centre of Excellence for the Dynamics of Language | academic | Australia | 3 | - | 39 | 13 | 0.48 |
| **2188** | TechnoSuruga Laboratory Co., Ltd. | corporate | Japan | 3 | - | 11 | 3.7 | 0.93 |
| **2189** | Fukuyama City Hospital | medical | Japan | 3 | - | 0 | 0 | 0 |
| **2190** | Miyazaki Prefectural Miyazaki Hospital | medical | Japan | 3 | - | 43 | 14.3 | 0.98 |
| **2191** | Iwate Tohoku Medical Megabank Organization | medical | Japan | 3 | - | 77 | 25.7 | 2.17 |
| **2192** | Adamas University | academic | India | 3 | - | 29 | 9.7 | 3.3 |
| **2193** | Tokyo Metropolitan Ohkubo Hospital | medical | Japan | 3 | -100 | 32 | 10.7 | 0.5 |
| **2194** | Yamanashi Prefectural Central Hospital | medical | Japan | 3 | -100 | 91 | 30.3 | 1.55 |
| **2195** | Hi-Tech Medical College and Hospital | medical | India | 3 | - | 28 | 9.3 | 2.9 |
| **2196** | National Hospital Organization Tokyo National Hospital | medical | Japan | 3 | -100 | 97 | 32.3 | 1.7 |
| **2197** | National Hospital Organization Nanao National Hospital | medical | Japan | 3 | - | 9 | 3 | 0.19 |
| **2198** | Japanese Red Cross Takamatsu Hospital | medical | Japan | 3 | - | 3 | 1 | 0.08 |
| **2199** | Nasu Red Cross Hospital | medical | Japan | 3 | 0 | 54 | 18 | 0.72 |
| **2200** | Japanese Red Cross College of Nursing | academic | Japan | 3 | - | 5 | 1.7 | 1.16 |
| **2201** | Japanese Red Cross Hiroshima College of Nursing | academic | Japan | 3 | - | 22 | 7.3 | 0.59 |
| **2202** | Saiseikai Suita Hospital | medical | Japan | 3 | - | 20 | 6.7 | 1.11 |
| **2203** | Saiseikai Yokohamashi Nanbu Hospital | medical | Japan | 3 | - | 17 | 5.7 | 0.69 |
| **2204** | Can Tho University of Medicine and Pharmacy | academic | Viet Nam | 3 | - | 46 | 15.3 | 0.91 |
| **2205** | Tokai Central Hospital | medical | Japan | 3 | - | 40 | 13.3 | 1.09 |
| **2206** | StaGen Co., Ltd. | corporate | Japan | 3 | -100 | 38 | 12.7 | 0.94 |
| **2207** | Mitoyo General Hospital | medical | Japan | 3 | - | 5 | 1.7 | 0 |
| **2208** | Fatima Jinnah Medical University | academic | Pakistan | 3 | - | 16 | 5.3 | 1.5 |
| **2209** | Japanese Organisation for Research and Treatment of Cancer | other | Japan | 3 | - | 27 | 9 | 1.47 |
| **2210** | Hamamatsu Pharma Research, Inc. | corporate | Japan | 3 | - | 43 | 14.3 | 1.44 |
| **2211** | Kyrgyz-Russian Slavic University | academic | Kyrgyzstan | 3 | - | 66 | 22 | 10.27 |
| **2212** | Sylhet MAG Osmani Medical College | academic | Bangladesh | 3 | - | 12 | 4 | 1.3 |
| **2213** | Institute of Medical and Veterinary Science Australia | government | Australia | 2 | 0 | 21 | 10.5 | 0.38 |
| **2214** | NSW Department of Primary Industries | government | Australia | 2 | - | 16 | 8 | 2.15 |
| **2215** | Beijing Forestry University | academic | China | 2 | - | 12 | 6 | 0.47 |
| **2216** | Chang'an University | academic | China | 2 | - | 8 | 4 | 0.37 |
| **2217** | China Academy of Engineering Physics | government | China | 2 | - | 31 | 15.5 | 1.14 |
| **2218** | Guangxi University | academic | China | 2 | - | 11 | 5.5 | 0.77 |
| **2219** | Harbin University of Science and Technology | academic | China | 2 | - | 22 | 11 | 0.54 |
| **2220** | Hebei Agricultural University | academic | China | 2 | 0 | 37 | 18.5 | 0.54 |
| **2221** | Qufu Normal University | academic | China | 2 | - | 11 | 5.5 | 0.63 |
| **2222** | Shandong University of Science and Technology | academic | China | 2 | - | 7 | 3.5 | 0.54 |
| **2223** | Shanghai Maritime University | academic | China | 2 | - | 22 | 11 | 0.91 |
| **2224** | Wuhan Institute of Technology | academic | China | 2 | - | 3 | 1.5 | 0.27 |
| **2225** | Xi'an University of Architecture and Technology | academic | China | 2 | - | 9 | 4.5 | 0.59 |
| **2226** | Xihua University | academic | China | 2 | - | 14 | 7 | 2.8 |
| **2227** | Xinjiang University | academic | China | 2 | - | 101 | 50.5 | 3.34 |
| **2228** | Yunnan Normal University | academic | China | 2 | - | 23 | 11.5 | 0.78 |
| **2229** | Zhejiang Agriculture and Forestry University | academic | China | 2 | - | 6 | 3 | 0.86 |
| **2230** | Bohai University | academic | China | 2 | - | 261 | 130.5 | 1.7 |
| **2231** | CAS - Institute of Chemistry | academic | China | 2 | - | 50 | 25 | 3.07 |
| **2232** | CAS - Institute of Software | academic | China | 2 | - | 55 | 27.5 | 1.88 |
| **2233** | CAS - Shanghai Institute of Ceramics | academic | China | 2 | - | 12 | 6 | 0.56 |
| **2234** | Changshu Institute of Technology | academic | China | 2 | - | 29 | 14.5 | 0.66 |
| **2235** | East China Jiaotong University | academic | China | 2 | - | 30 | 15 | 1.55 |
| **2236** | Foshan University | academic | China | 2 | -100 | 28 | 14 | 0.59 |
| **2237** | IAPCM | academic | China | 2 | - | 29 | 14.5 | 1.48 |
| **2238** | Ludong University | academic | China | 2 | - | 54 | 27 | 0.91 |
| **2239** | Luoyang Normal University | academic | China | 2 | 0 | 21 | 10.5 | 0.25 |
| **2240** | National Institute of Metrology China | government | China | 2 | - | 0 | 0 | 0 |
| **2241** | Northeast Dianli University | academic | China | 2 | - | 35 | 17.5 | 0.99 |
| **2242** | Southwest Petroleum University China | academic | China | 2 | - | 14 | 7 | 0.61 |
| **2243** | State Oceanic Administration China | government | China | 2 | - | 8 | 4 | 0.8 |
| **2244** | Lingnan University | academic | Hong Kong | 2 | - | 12 | 6 | 0.82 |
| **2245** | Indian Space Research Organization | government | India | 2 | - | 2 | 1 | 0.32 |
| **2246** | Jawaharlal Nehru Technological University | academic | India | 2 | - | 24 | 12 | 1.03 |
| **2247** | Karnatak University | academic | India | 2 | - | 8 | 4 | 0.4 |
| **2248** | Mangalore University | academic | India | 2 | - | 14 | 7 | 0.39 |
| **2249** | Institute of Physics Bhubaneswar | government | India | 2 | -100 | 64 | 32 | 1.78 |
| **2250** | University of Calicut | academic | India | 2 | - | 2 | 1 | 0.18 |
| **2251** | Visva-Bharati University | academic | India | 2 | - | 3 | 1.5 | 0.2 |
| **2252** | Hosei University | academic | Japan | 2 | - | 56 | 28 | 1.15 |
| **2253** | Nagaoka University of Technology | academic | Japan | 2 | - | 3 | 1.5 | 0.04 |
| **2254** | National Defense Academy of Japan | academic | Japan | 2 | - | 226 | 113 | 5.46 |
| **2255** | Okayama University of Science | academic | Japan | 2 | -100 | 25 | 12.5 | 0.59 |
| **2256** | Osaka Institute of Technology | academic | Japan | 2 | -100 | 4 | 2 | 0.11 |
| **2257** | Research Organization of Information and Systems, National Institute of Informatics | academic | Japan | 2 | - | 31 | 15.5 | 1.85 |
| **2258** | Toyohashi University of Technology | academic | Japan | 2 | - | 192 | 96 | 2.27 |
| **2259** | Utsunomiya University | academic | Japan | 2 | - | 15 | 7.5 | 0.74 |
| **2260** | Ajinomoto Co Inc | corporate | Japan | 2 | - | 18 | 9 | 1.54 |
| **2261** | Fukuoka Institute of Technology | academic | Japan | 2 | - | 52 | 26 | 1.95 |
| **2262** | Kao Corporation | corporate | Japan | 2 | - | 18 | 9 | 0.79 |
| **2263** | Kazusa DNA Research Institute | other | Japan | 2 | - | 15 | 7.5 | 0.52 |
| **2264** | Meisei University | academic | Japan | 2 | - | 6 | 3 | 0.26 |
| **2265** | NEC Corporation | corporate | Japan | 2 | - | 225 | 112.5 | 4.62 |
| **2266** | Okayama Prefectural University | academic | Japan | 2 | - | 1 | 0.5 | 0.09 |
| **2267** | Rakuno Gakuen University | academic | Japan | 2 | - | 3 | 1.5 | 0.05 |
| **2268** | Saitama Institute of Technology | academic | Japan | 2 | - | 7 | 3.5 | 0.59 |
| **2269** | Seikei University | academic | Japan | 2 | - | 2 | 1 | 0.72 |
| **2270** | Suntory Holdings Limited | corporate | Japan | 2 | - | 26 | 13 | 0.74 |
| **2271** | Tsurumi University | academic | Japan | 2 | - | 30 | 15 | 2.46 |
| **2272** | Korea Institute of Ocean Science & Technology | academic | South Korea | 2 | - | 34 | 17 | 0.6 |
| **2273** | Kumoh National Institute of Technology | academic | South Korea | 2 | - | 18 | 9 | 0.88 |
| **2274** | Kunsan National University | academic | South Korea | 2 | - | 10 | 5 | 0.97 |
| **2275** | Food and Drug Administration of Korea | government | South Korea | 2 | - | 24 | 12 | 0.71 |
| **2276** | Korea Electrotechnology Research Institute | government | South Korea | 2 | - | 15 | 7.5 | 0.48 |
| **2277** | Rural Development Administration | government | South Korea | 2 | - | 42 | 21 | 1.1 |
| **2278** | AgResearch | government | New Zealand | 2 | - | 26 | 13 | 0.85 |
| **2279** | ESR | government | New Zealand | 2 | - | 143 | 71.5 | 11.54 |
| **2280** | University of Agriculture Faisalabad | academic | Pakistan | 2 | - | 5 | 2.5 | 0.96 |
| **2281** | Pakistan Institute of Engineering and Applied Sciences | academic | Pakistan | 2 | - | 58 | 29 | 1.53 |
| **2282** | King Mongkut's University of Technology Thonburi | academic | Thailand | 2 | 0 | 51 | 25.5 | 1.54 |
| **2283** | Chaoyang University of Technology | academic | Taiwan | 2 | -100 | 10 | 5 | 0.29 |
| **2284** | Cheng Shiu University Taiwan | academic | Taiwan | 2 | - | 63 | 31.5 | 1.19 |
| **2285** | National Kaohsiung Normal University | academic | Taiwan | 2 | - | 37 | 18.5 | 1 |
| **2286** | Shanghai Ocean University | academic | China | 2 | - | 4 | 2 | 0.37 |
| **2287** | TianJin University of Technology and Education | academic | China | 2 | - | 17 | 8.5 | 1.05 |
| **2288** | MingDao University | academic | Taiwan | 2 | - | 2 | 1 | 0.16 |
| **2289** | CAS - Shenyang Institute of Automation | academic | China | 2 | - | 30 | 15 | 0.47 |
| **2290** | CAS - Xi'an Institute of Optics and Precision Mechanics | academic | China | 2 | - | 32 | 16 | 1.11 |
| **2291** | CAS - Institute of Intelligent Machines | academic | China | 2 | - | 102 | 51 | 3.15 |
| **2292** | CAS - Northwest Institute of Plateau Biology | academic | China | 2 | - | 52 | 26 | 1.68 |
| **2293** | Singapore University of Technology and Design | academic | Singapore | 2 | - | 19 | 9.5 | 0.96 |
| **2294** | National Fusion Research Institute | government | South Korea | 2 | - | 4 | 2 | 0.17 |
| **2295** | China University of Political Science and Law | academic | China | 2 | - | 7 | 3.5 | 0.58 |
| **2296** | Lions Eye Institute | academic | Australia | 2 | - | 1864 | 932 | 67.83 |
| **2297** | Institut Pasteur du Cambodge | academic | Cambodia | 2 | - | 8 | 4 | 0.25 |
| **2298** | Institut Pasteur de Nouvelle-Caledonie Noumea | academic | New Caledonia | 2 | - | 49 | 24.5 | 2.53 |
| **2299** | Mongolian Academy of Sciences | academic | Mongolia | 2 | - | 14 | 7 | 0.62 |
| **2300** | Universiti Teknikal Malaysia Melaka | academic | Malaysia | 2 | - | 27 | 13.5 | 0.81 |
| **2301** | Korea Institute of Science and Technology Information | government | South Korea | 2 | - | 34 | 17 | 0.93 |
| **2302** | Southwestern University of Finance and Economics | academic | China | 2 | - | 57 | 28.5 | 1.18 |
| **2303** | Balochistan University of Information Technology, Engineering and Management Sciences | academic | Pakistan | 2 | - | 120 | 60 | 2.25 |
| **2304** | Rajiv Gandhi Technical University | academic | India | 2 | - | 59 | 29.5 | 1.17 |
| **2305** | Indian Institute of Technology Ropar | academic | India | 2 | -100 | 134 | 67 | 3.37 |
| **2306** | Saken Seifullin Kazakh Agrotechnical University | academic | Kazakhstan | 2 | - | 5 | 2.5 | 0.26 |
| **2307** | Muhammadiyah University of Surakarta | academic | Indonesia | 2 | - | 44 | 22 | 3.79 |
| **2308** | Guizhou Normal University | academic | China | 2 | - | 62 | 31 | 1.22 |
| **2309** | Guangxi University of Technology | academic | China | 2 | - | 2 | 1 | 0.29 |
| **2310** | Shandong University of Finance and Economics | academic | China | 2 | - | 17 | 8.5 | 0.76 |
| **2311** | Sri Sivasubramaniya Nadar College of Engineering, Chennai | academic | India | 2 | - | 9 | 4.5 | 0.28 |
| **2312** | Peoples Education Society, Bangalore | academic | India | 2 | - | 68 | 34 | 4.26 |
| **2313** | Santen Pharmaceutical Co., Ltd. | corporate | Japan | 2 | - | 6 | 3 | 0.85 |
| **2314** | Sumitomo Heavy Industries, Ltd. | corporate | Japan | 2 | -100 | 25 | 12.5 | 0.64 |
| **2315** | Toray Industries, Inc. | corporate | Japan | 2 | - | 57 | 28.5 | 2.19 |
| **2316** | Taisho Pharmaceutical Co., Ltd. | corporate | Japan | 2 | - | 42 | 21 | 1.69 |
| **2317** | FAW Group Corporation | corporate | China | 2 | - | 11 | 5.5 | 0.67 |
| **2318** | State Government of Victoria | government | Australia | 2 | -100 | 7 | 3.5 | 0 |
| **2319** | Chinese Academy of Social Sciences | government | China | 2 | - | 4 | 2 | 0.75 |
| **2320** | Karnataka Veterinary, Animal & Fisheries Sciences University | academic | India | 2 | - | 14 | 7 | 0.95 |
| **2321** | Nepal Medical College | academic | Nepal | 2 | - | 5 | 2.5 | 0.35 |
| **2322** | Ayub Medical College | academic | Pakistan | 2 | - | 11 | 5.5 | 0.5 |
| **2323** | Bangladesh Institute of Research and Rehabilitation in Diabetes, Endocrine and Metabolic Disorders | government | Bangladesh | 2 | - | 9 | 4.5 | 0.11 |
| **2324** | Kalasalingam University | academic | India | 2 | - | 0 | 0 | 0 |
| **2325** | National Institute of Technology Karnataka | academic | India | 2 | - | 14 | 7 | 1.2 |
| **2326** | Gandhi Institute of Technology and Management | academic | India | 2 | 0 | 4 | 2 | 0.38 |
| **2327** | University of Chittagong | academic | Bangladesh | 2 | - | 1 | 0.5 | 0.14 |
| **2328** | Ningbo University of Technology | academic | China | 2 | - | 23 | 11.5 | 0.56 |
| **2329** | Huaiyin Institute of Technology | academic | China | 2 | - | 28 | 14 | 1.22 |
| **2330** | Xuchang University | academic | China | 2 | - | 9 | 4.5 | 0.55 |
| **2331** | Taishan University | academic | China | 2 | - | 148 | 74 | 3.82 |
| **2332** | Dezhou University | academic | China | 2 | - | 11 | 5.5 | 0.52 |
| **2333** | Kokugakuin University | academic | Japan | 2 | - | 0 | 0 | 0 |
| **2334** | Komazawa University | academic | Japan | 2 | -100 | 9 | 4.5 | 0.24 |
| **2335** | National Graduate Institute for Policy Studies | academic | Japan | 2 | - | 11 | 5.5 | 1.01 |
| **2336** | Guangdong Institute of Microbiology | academic | China | 2 | - | 68 | 34 | 2.92 |
| **2337** | PEC University of Technology | academic | India | 2 | - | 4 | 2 | 0.17 |
| **2338** | Hokkaido University of Education | academic | Japan | 2 | -100 | 15 | 7.5 | 1.31 |
| **2339** | Joetsu University of Education | academic | Japan | 2 | - | 0 | 0 | 0 |
| **2340** | Nara University of Education | academic | Japan | 2 | -100 | 12 | 6 | 0.19 |
| **2341** | Manipur University | academic | India | 2 | - | 19 | 9.5 | 0.86 |
| **2342** | Jiwaji University | academic | India | 2 | - | 91 | 45.5 | 1.46 |
| **2343** | Heze University | academic | China | 2 | - | 133 | 66.5 | 2.78 |
| **2344** | Dongduk Women's University | academic | South Korea | 2 | - | 28 | 14 | 1.28 |
| **2345** | Academia Sinica - Institute of Chemistry | academic | Taiwan | 2 | - | 59 | 29.5 | 1.98 |
| **2346** | Academia Sinica - Institute of Physics | academic | Taiwan | 2 | - | 19 | 9.5 | 0.26 |
| **2347** | Academia Sinica - Institute of European and American Studies | academic | Taiwan | 2 | - | 28 | 14 | 0.74 |
| **2348** | Academia Sinica - Research Center for Humanities and Social Science | academic | Taiwan | 2 | - | 195 | 97.5 | 3.67 |
| **2349** | University of Cyberjaya | academic | Malaysia | 2 | - | 6 | 3 | 0.88 |
| **2350** | Quest International University Perak (QIUP) | academic | Malaysia | 2 | - | 27 | 13.5 | 0.5 |
| **2351** | Animal and Plant Quarantine Agency | government | South Korea | 2 | - | 13 | 6.5 | 0.79 |
| **2352** | University of Jember | academic | Indonesia | 2 | - | 3 | 1.5 | 0.09 |
| **2353** | Central University of Tamil Nadu | academic | India | 2 | - | 5 | 2.5 | 0.24 |
| **2354** | Yokohama University of Pharmacy | academic | Japan | 2 | - | 75 | 37.5 | 1.25 |
| **2355** | Kyoto University of Advanced Science | academic | Japan | 2 | - | 8 | 4 | 1.26 |
| **2356** | Doshisha Women's College of Liberal Arts | academic | Japan | 2 | -100 | 37 | 18.5 | 0.71 |
| **2357** | Changchun Normal University | academic | China | 2 | - | 7 | 3.5 | 0.3 |
| **2358** | Beijing Foreign Studies University | academic | China | 2 | - | 2 | 1 | 0.18 |
| **2359** | Tianjin University of Sport | academic | China | 2 | - | 11 | 5.5 | 0.14 |
| **2360** | Ubon Rachathani University | academic | Thailand | 2 | - | 140 | 70 | 3.09 |
| **2361** | Suan Sunandha Rajabhat University | academic | Thailand | 2 | - | 25 | 12.5 | 0.79 |
| **2362** | Boe Technology Group | corporate | China | 2 | - | 2 | 1 | 0.32 |
| **2363** | Hubei University of Technology | academic | China | 2 | - | 9 | 4.5 | 1.46 |
| **2364** | Mesoblast | corporate | Australia | 2 | - | 37 | 18.5 | 1.38 |
| **2365** | Patna University | academic | India | 2 | - | 94 | 47 | 6.33 |
| **2366** | Latrobe Regional Hospital | medical | Australia | 2 | - | 3 | 1.5 | 0.19 |
| **2367** | Department of Health and Human Services Tasmania | government | Australia | 2 | - | 8 | 4 | 0.44 |
| **2368** | Nelson Marlborough District Health Board | government | New Zealand | 2 | - | 19 | 9.5 | 2.24 |
| **2369** | South Canterbury District Health Board | government | New Zealand | 2 | - | 38 | 19 | 1.57 |
| **2370** | Tamil University | academic | India | 2 | - | 31 | 15.5 | 0.32 |
| **2371** | Koneru Lakshmaiah Education Foundation | academic | India | 2 | - | 0 | 0 | 0 |
| **2372** | Torrens University Australia | academic | Australia | 2 | - | 2 | 1 | 0.65 |
| **2373** | S. Toraighyrov Pavlodar State University | academic | Kazakhstan | 2 | - | 5 | 2.5 | 0.21 |
| **2374** | Rajamangala University of Technology Thanyaburi | academic | Thailand | 2 | - | 8 | 4 | 0.14 |
| **2375** | O.P. Jindal Global University | academic | India | 2 | - | 5 | 2.5 | 0.26 |
| **2376** | Tainan University of Technology | academic | Taiwan | 2 | - | 0 | 0 | 0 |
| **2377** | Sher-e-Kashmir University of Agricultural Sciences and Technology of Jammu | academic | India | 2 | - | 113 | 56.5 | 2.15 |
| **2378** | TOPCON Corporation | corporate | Japan | 2 | - | 116 | 58 | 4.47 |
| **2379** | Guangxi Teachers Education University | academic | China | 2 | - | 220 | 110 | 2.66 |
| **2380** | Jiangsu University of Technology | academic | China | 2 | - | 15 | 7.5 | 0.21 |
| **2381** | Xi'an International Studies University | academic | China | 2 | - | 16 | 8 | 1.19 |
| **2382** | J.F. Oberlin University | academic | Japan | 2 | - | 49 | 24.5 | 1.72 |
| **2383** | Riddet Institute | academic | New Zealand | 2 | 0 | 33 | 16.5 | 0.43 |
| **2384** | Acer Inc. | corporate | Taiwan | 2 | - | 19 | 9.5 | 0.6 |
| **2385** | WuXi AppTec | corporate | China | 2 | - | 33 | 16.5 | 1.81 |
| **2386** | Takara Bio Inc. | corporate | Japan | 2 | 0 | 12 | 6 | 1.73 |
| **2387** | Morinaga Milk Industry Co., Ltd. | corporate | Japan | 2 | - | 78 | 39 | 3.12 |
| **2388** | Seikagaku Corporation | corporate | Japan | 2 | - | 67 | 33.5 | 1.75 |
| **2389** | Shino-Test Corporation | corporate | Japan | 2 | - | 18 | 9 | 0.32 |
| **2390** | ANBAS Corporation | corporate | Japan | 2 | - | 45 | 22.5 | 0.39 |
| **2391** | Central Japan Railway Company | corporate | Japan | 2 | -100 | 3 | 1.5 | 0.03 |
| **2392** | NTN Corporation | corporate | Japan | 2 | -100 | 10 | 5 | 0.12 |
| **2393** | Delhi Technological University | academic | India | 2 | - | 11 | 5.5 | 0.59 |
| **2394** | St John of God Health Care | medical | Australia | 2 | - | 8 | 4 | 0.31 |
| **2395** | Korea Occupational Safety and Health Agency | government | South Korea | 2 | - | 60 | 30 | 2.13 |
| **2396** | Ministry of Health, Nutrition and Indigenous Medicine | government | Sri Lanka | 2 | - | 170 | 85 | 4.09 |
| **2397** | Waikato Institute of Technology | academic | New Zealand | 2 | - | 6 | 3 | 0.72 |
| **2398** | Handong Global University | academic | South Korea | 2 | -100 | 14 | 7 | 0.52 |
| **2399** | Kazakh National Agrarian University | academic | Kazakhstan | 2 | - | 10 | 5 | 0.22 |
| **2400** | Vels Institute of Science, Technology & Advanced Studies | academic | India | 2 | - | 9 | 4.5 | 1.7 |
| **2401** | Karpagam Academy of Higher Education | academic | India | 2 | - | 24 | 12 | 0.38 |
| **2402** | Dr. M.G.R Educational and Research Institute | academic | India | 2 | - | 14 | 7 | 0.83 |
| **2403** | Swami Vivekanand Subharti University | academic | India | 2 | - | 2 | 1 | 0.1 |
| **2404** | Rayalaseema University | academic | India | 2 | - | 33 | 16.5 | 0.6 |
| **2405** | Babu Banarasi Das Group of Educational Institutions | academic | India | 2 | - | 23 | 11.5 | 2.01 |
| **2406** | Indian Institute of Food Processing Technology | academic | India | 2 | - | 31 | 15.5 | 0.32 |
| **2407** | Family Planning NSW | other | Australia | 2 | - | 29 | 14.5 | 1.05 |
| **2408** | Hiroshima Shudo University | academic | Japan | 2 | - | 25 | 12.5 | 0.6 |
| **2409** | Nakamura Gakuen University | academic | Japan | 2 | - | 24 | 12 | 0.76 |
| **2410** | DIT University | academic | India | 2 | - | 6 | 3 | 0.31 |
| **2411** | Ministry of Science and Technology, India | government | India | 2 | - | 229 | 114.5 | 5.16 |
| **2412** | China Disabled Persons' Federation | other | China | 2 | - | 10 | 5 | 3.04 |
| **2413** | Regional Centre for Biotechnology | academic | India | 2 | - | 15 | 7.5 | 0.41 |
| **2414** | IFLYTEK Co., Ltd. | corporate | China | 2 | - | 4 | 2 | 0.41 |
| **2415** | Hisamitsu Pharmaceutical Co., Inc. | corporate | Japan | 2 | - | 11 | 5.5 | 0.7 |
| **2416** | IIHMR University, Jaipur | academic | India | 2 | - | 1861 | 930.5 | 65.24 |
| **2417** | Suresh Gyan Vihar University | academic | India | 2 | - | 25 | 12.5 | 6.85 |
| **2418** | Zhejiang Shuren University | academic | China | 2 | - | 1 | 0.5 | 0.35 |
| **2419** | Zhaoqing University | academic | China | 2 | - | 3 | 1.5 | 0.23 |
| **2420** | China National GeneBank | government | China | 2 | - | 24 | 12 | 0.95 |
| **2421** | Hengyang Normal University | academic | China | 2 | - | 22 | 11 | 0.5 |
| **2422** | Guiyang University | academic | China | 2 | - | 11 | 5.5 | 0.32 |
| **2423** | Sanming University | academic | China | 2 | - | 10 | 5 | 0.48 |
| **2424** | Ningde Normal University | academic | China | 2 | - | 0 | 0 | 0 |
| **2425** | Shanghai University of Political Science and Law | academic | China | 2 | - | 18 | 9 | 0.69 |
| **2426** | Sichuan International Studies University | academic | China | 2 | - | 9 | 4.5 | 0.24 |
| **2427** | Beijing University of Agriculture | academic | China | 2 | - | 28 | 14 | 1.11 |
| **2428** | Noorul Islam University | academic | India | 2 | -100 | 67 | 33.5 | 1.44 |
| **2429** | Integral University | academic | India | 2 | - | 11 | 5.5 | 0.5 |
| **2430** | Rajshahi University of Engineering and Technology | academic | Bangladesh | 2 | - | 1 | 0.5 | 0.26 |
| **2431** | Sichuan Academy of Agricultural Sciences | academic | China | 2 | - | 8 | 4 | 0.2 |
| **2432** | Indian Institute of Information Technology, Allahabad | academic | India | 2 | - | 57 | 28.5 | 1.49 |
| **2433** | Yessenov University | academic | Kazakhstan | 2 | - | 8 | 4 | 0.4 |
| **2434** | Kuo General Hospital | medical | Taiwan | 2 | - | 15 | 7.5 | 0.43 |
| **2435** | Tian-Sheng Memorial Hospital | medical | Taiwan | 2 | -100 | 9 | 4.5 | 0.81 |
| **2436** | Krida Wacana Christian University | academic | Indonesia | 2 | - | 3 | 1.5 | 0.34 |
| **2437** | Christian University of Indonesia | academic | Indonesia | 2 | - | 0 | 0 | 0 |
| **2438** | Maranatha Christian Unviersity | academic | Indonesia | 2 | - | 6 | 3 | 0.55 |
| **2439** | Heilongjiang Institute of Technology | academic | China | 2 | - | 52 | 26 | 0.52 |
| **2440** | Handan College | academic | China | 2 | - | 6 | 3 | 1.9 |
| **2441** | Government Medical College, Amritsar | government | India | 2 | - | 1 | 0.5 | 0.27 |
| **2442** | JSS Science and Technology University | academic | India | 2 | - | 15 | 7.5 | 0.65 |
| **2443** | Shandong Eye Institute and Hospital | medical | China | 2 | - | 17 | 8.5 | 0.31 |
| **2444** | Mulawarman University | academic | Indonesia | 2 | - | 18 | 9 | 0.72 |
| **2445** | South Asian University | academic | India | 2 | - | 30 | 15 | 0.75 |
| **2446** | Fuzzy Logic Systems Institute | other | Japan | 2 | - | 28 | 14 | 0.46 |
| **2447** | Siam University | academic | Thailand | 2 | - | 3 | 1.5 | 0.27 |
| **2448** | Vaxine Pty Ltd. | corporate | Australia | 2 | - | 65 | 32.5 | 2.38 |
| **2449** | Kyungil University | academic | South Korea | 2 | - | 25 | 12.5 | 0.72 |
| **2450** | Institute of Nano Science and Technology | academic | India | 2 | - | 147 | 73.5 | 3.96 |
| **2451** | Gujarat Cancer and Research Institute | medical | India | 2 | 0 | 4 | 2 | 0.05 |
| **2452** | Tanjungpura University | academic | Indonesia | 2 | - | 0 | 0 | 0 |
| **2453** | World University of Bangladesh | academic | Bangladesh | 2 | - | 25 | 12.5 | 1 |
| **2454** | Tohoku Bunka Gakuen University | academic | Japan | 2 | -100 | 22 | 11 | 0.35 |
| **2455** | Vongchavalitkul University | academic | Thailand | 2 | - | 41 | 20.5 | 0.64 |
| **2456** | Uiduk University | academic | South Korea | 2 | - | 8 | 4 | 0.43 |
| **2457** | Nambu University | academic | South Korea | 2 | - | 28 | 14 | 0.63 |
| **2458** | Mawlana Bhashani Science and Technology University | academic | Bangladesh | 2 | - | 31 | 15.5 | 1.88 |
| **2459** | Tianjin People's Hospital | medical | China | 2 | - | 0 | 0 | 0 |
| **2460** | Pok Oi Hospital | medical | Hong Kong | 2 | - | 2 | 1 | 0.25 |
| **2461** | PLA No. 254 Hospital | medical | China | 2 | -100 | 121 | 60.5 | 1.51 |
| **2462** | PLA No. 252 Hospital | medical | China | 2 | -100 | 4 | 2 | 0.2 |
| **2463** | PLA No. 305 Hospital | medical | China | 2 | - | 4 | 2 | 0.06 |
| **2464** | Southern University Bangladesh | academic | Bangladesh | 2 | - | 638 | 319 | 6.94 |
| **2465** | Sapthagiri Institute of Medical Science and Research Center | academic | India | 2 | - | 18 | 9 | 0.35 |
| **2466** | University of Swabi | academic | Pakistan | 2 | - | 46 | 23 | 0.95 |
| **2467** | Yamaguchi Prefectural University | academic | Japan | 2 | -100 | 8 | 4 | 1.09 |
| **2468** | Patanjali Research Foundation | academic | India | 2 | - | 20 | 10 | 0.56 |
| **2469** | Henan University of Animal Husbandry and Economy | academic | China | 2 | - | 5 | 2.5 | 0.34 |
| **2470** | National University of Computer and Emerging Sciences, Lahore | academic | Pakistan | 2 | - | 2 | 1 | 0.06 |
| **2471** | Sir Syed University of Engineering and Technology | academic | Pakistan | 2 | - | 33 | 16.5 | 0.68 |
| **2472** | CSIR - National Environmental Engineering Research Institute | government | India | 2 | -100 | 31 | 15.5 | 0.85 |
| **2473** | CSIR - Central Scientific Instruments Organisation | government | India | 2 | - | 189 | 94.5 | 2.14 |
| **2474** | South Western Sydney Area Health Service | medical | Australia | 2 | - | 46 | 23 | 0.85 |
| **2475** | National Institutes for Food and Drug Control | academic | China | 2 | - | 21 | 10.5 | 1.11 |
| **2476** | Zhengzhou Railway Vocational and Technical College | academic | China | 2 | - | 1 | 0.5 | 0.25 |
| **2477** | Noakhali Science and Technology University | academic | Bangladesh | 2 | - | 18 | 9 | 0.54 |
| **2478** | Hang Tuah University | academic | Indonesia | 2 | - | 3 | 1.5 | 0.14 |
| **2479** | Forensicare | medical | Australia | 2 | - | 18 | 9 | 0.61 |
| **2480** | Rehman Medical Institute | corporate | Pakistan | 2 | - | 3 | 1.5 | 0.74 |
| **2481** | Suven Life Sciences Ltd. | corporate | India | 2 | - | 5 | 2.5 | 1.29 |
| **2482** | Shanghai ChemPartner | corporate | China | 2 | - | 6 | 3 | 1.12 |
| **2483** | Pavlodar State Pedagogical University | academic | Kazakhstan | 2 | - | 3 | 1.5 | 0.15 |
| **2484** | Silkway International University | academic | Kazakhstan | 2 | - | 0 | 0 | 0 |
| **2485** | Ualikhanov Kokshetau State University | academic | Kazakhstan | 2 | - | 0 | 0 | 0 |
| **2486** | Osaka Institute of Public Health | government | Japan | 2 | - | 3 | 1.5 | 0.12 |
| **2487** | Minamisoma Municipal General Hospital | medical | Japan | 2 | - | 21 | 10.5 | 0.54 |
| **2488** | New Tokyo Hospital | medical | Japan | 2 | - | 40 | 20 | 1.35 |
| **2489** | Meiji Yasuda Life Foundation of Health and Welfare | other | Japan | 2 | - | 20 | 10 | 1.06 |
| **2490** | Andijan State Medical Institute | academic | Uzbekistan | 2 | - | 0 | 0 | 0 |
| **2491** | Eternal University | academic | India | 2 | - | 4 | 2 | 0.43 |
| **2492** | Gunma Prefectural College of Health Sciences | academic | Japan | 2 | - | 1 | 0.5 | 0.13 |
| **2493** | University of Kochi | academic | Japan | 2 | - | 36 | 18 | 1.4 |
| **2494** | Chiba Prefectural Institute of Public Health | government | Japan | 2 | 0 | 40 | 20 | 0.57 |
| **2495** | Sapporo City University | academic | Japan | 2 | -100 | 22 | 11 | 0.56 |
| **2496** | Shonan University of Medical Sciences | academic | Japan | 2 | - | 4 | 2 | 0.18 |
| **2497** | Shigakkan University | academic | Japan | 2 | - | 7 | 3.5 | 0.22 |
| **2498** | Ehime Prefectural University of Health Sciences | academic | Japan | 2 | - | 6 | 3 | 0.36 |
| **2499** | Taisho University | academic | Japan | 2 | - | 0 | 0 | 0 |
| **2500** | Toyohashi Sozo University | academic | Japan | 2 | - | 25 | 12.5 | 0.66 |
| **2501** | Wayo Women's University | academic | Japan | 2 | - | 12 | 6 | 3.24 |
| **2502** | Morioka University | academic | Japan | 2 | - | 6 | 3 | 0.4 |
| **2503** | Ishikawa Prefectural Nursing University | academic | Japan | 2 | - | 3 | 1.5 | 0.64 |
| **2504** | Shukutoku University | academic | Japan | 2 | - | 4 | 2 | 0.24 |
| **2505** | Baika Women's University | academic | Japan | 2 | 0 | 9 | 4.5 | 0.25 |
| **2506** | Gunma Institute of Public Health and Environmental Sciences | government | Japan | 2 | - | 13 | 6.5 | 0.66 |
| **2507** | Uekusa Gakuen University | academic | Japan | 2 | - | 15 | 7.5 | 0.45 |
| **2508** | Tokyo University and Graduate School of Social Welfare | academic | Japan | 2 | - | 26 | 13 | 0.78 |
| **2509** | Japan Ministry of Education, Culture, Sports, Science and Technology | government | Japan | 2 | - | 105 | 52.5 | 2.39 |
| **2510** | College of Healthcare Management | academic | Japan | 2 | - | 9 | 4.5 | 0.46 |
| **2511** | SBI Pharmaceuticals Co., Ltd. | corporate | Japan | 2 | - | 6 | 3 | 0.28 |
| **2512** | Tokiwakai Group | medical | Japan | 2 | - | 41 | 20.5 | 1.58 |
| **2513** | Human Metabolome Technologies, Inc. | corporate | Japan | 2 | - | 41 | 20.5 | 1.23 |
| **2514** | ARC Centre of Excellence in Advanced Molecular Imaging | academic | Australia | 2 | - | 200 | 100 | 3.22 |
| **2515** | Japan Ministry of Finance | government | Japan | 2 | - | 8 | 4 | 0.27 |
| **2516** | Nipro Corporation | corporate | Japan | 2 | -100 | 13 | 6.5 | 0.56 |
| **2517** | Bureau of Social Welfare and Public Health | government | Japan | 2 | - | 12 | 6 | 0.46 |
| **2518** | Immuno-Biological Laboratories Co., Ltd. | corporate | Japan | 2 | -100 | 64 | 32 | 1.04 |
| **2519** | Maxell, Ltd. | corporate | Japan | 2 | - | 1 | 0.5 | 0.29 |
| **2520** | Mizuno Corporation | corporate | Japan | 2 | - | 4 | 2 | 0 |
| **2521** | Daqing Oilfield Company Ltd. | corporate | China | 2 | -100 | 26 | 13 | 0.59 |
| **2522** | National Hospital Organization Saitama National Hospital | medical | Japan | 2 | - | 14 | 7 | 0.57 |
| **2523** | National Hospital Organization Kochi National Hospital | medical | Japan | 2 | - | 4 | 2 | 1.08 |
| **2524** | National Hospital Organization Yonago Medical Center | medical | Japan | 2 | - | 27 | 13.5 | 1.3 |
| **2525** | National Hospital Organization Hirosaki National Hospital | medical | Japan | 2 | - | 2 | 1 | 1.17 |
| **2526** | National Hospital Organization Ehime Medical Center | medical | Japan | 2 | - | 7 | 3.5 | 0.37 |
| **2527** | National Hospital Organization Saga National Hospital | medical | Japan | 2 | - | 14 | 7 | 0.57 |
| **2528** | Osaka Habikino Medical Center | medical | Japan | 2 | 0 | 34 | 17 | 5.87 |
| **2529** | Kobe City Medical Center West Hospital | medical | Japan | 2 | -100 | 28 | 14 | 0.8 |
| **2530** | Japanese Red Cross Kochi Hospital | medical | Japan | 2 | - | 3 | 1.5 | 0.21 |
| **2531** | Japanese Red Cross Oita Hospital | medical | Japan | 2 | - | 7 | 3.5 | 0.03 |
| **2532** | Cricket Australia | other | Australia | 2 | - | 13 | 6.5 | 0.43 |
| **2533** | Kumamoto Chuo Hospital | medical | Japan | 2 | - | 5 | 2.5 | 0.52 |
| **2534** | TXP Medical Co. Ltd. | corporate | Japan | 2 | - | 2 | 1 | 0.25 |
| **2535** | Chulabhorn Research Institute | academic | Thailand | 2 | - | 127 | 63.5 | 2.14 |
| **2536** | Japanese Drug Organization of Appropriate Use and Research | other | Japan | 2 | - | 17 | 8.5 | 0.67 |
| **2537** | GN Corporation Co. Ltd. | corporate | Japan | 2 | - | 6 | 3 | 1.55 |
| **2538** | Tokyo Metropolitan Government | government | Japan | 2 | -100 | 33 | 16.5 | 0.9 |
| **2539** | Kishiwada City Hospital | medical | Japan | 2 | - | 38 | 19 | 0.7 |
| **2540** | New Vision University | academic | Georgia | 2 | - | 3 | 1.5 | 0.95 |
| **2541** | JCR Pharmaceuticals Co., Ltd. | corporate | Japan | 2 | - | 11 | 5.5 | 1.03 |
| **2542** | Korea Nazarene University | academic | South Korea | 2 | - | 6 | 3 | 0.12 |
| **2543** | State University of Bangladesh | academic | Bangladesh | 2 | - | 3 | 1.5 | 0.16 |
| **2544** | Osh State University | academic | Kyrgyzstan | 2 | - | 5 | 2.5 | 0.14 |
| **2545** | Sher-e-Bangla Medical College | academic | Bangladesh | 2 | - | 12 | 6 | 0.28 |
| **2546** | Australian Antarctic Division | government | Australia | 1 | - | 8 | 8 | 0.59 |
| **2547** | Bangladesh University of Engineering and Technology | academic | Bangladesh | 1 | - | 10 | 10 | 1.16 |
| **2548** | Bangladesh Agricultural University | academic | Bangladesh | 1 | - | 43 | 43 | 2.34 |
| **2549** | Chengdu University of Technology | academic | China | 1 | - | 19 | 19 | 2.35 |
| **2550** | Dalian Maritime University | academic | China | 1 | - | 1 | 1 | 0 |
| **2551** | Guilin University of Electronic Technology | academic | China | 1 | - | 24 | 24 | 1.09 |
| **2552** | Hebei University of Science and Technology | academic | China | 1 | - | 33 | 33 | 1.04 |
| **2553** | Hefei University of Technology | academic | China | 1 | - | 9 | 9 | 0.83 |
| **2554** | Henan Agricultural University | academic | China | 1 | - | 7 | 7 | 0.26 |
| **2555** | Henan University of Technology | academic | China | 1 | - | 3 | 3 | 0.51 |
| **2556** | Hohai University | academic | China | 1 | - | 9 | 9 | 1.14 |
| **2557** | Liaoning University | academic | China | 1 | - | 5 | 5 | 0.92 |
| **2558** | Nanjing University of Information Science & Technology | academic | China | 1 | - | 22 | 22 | 2.54 |
| **2559** | Naval University of Engineering Wuhan | academic | China | 1 | - | 4 | 4 | 0.45 |
| **2560** | Shandong Agricultural University | academic | China | 1 | - | 18 | 18 | 0.89 |
| **2561** | Xi'an University of Technology | academic | China | 1 | - | 1 | 1 | 0.18 |
| **2562** | Academy of Armored Force Engineering China | academic | China | 1 | - | 0 | 0 | 0 |
| **2563** | Beijing Institute of Petrochemical Technology | academic | China | 1 | - | 3 | 3 | 0.55 |
| **2564** | CAS - Academy of Mathematics and System Sciences | academic | China | 1 | - | 14 | 14 | 1.49 |
| **2565** | CAS - Guangzhou Institute of Geochemistry | academic | China | 1 | - | 47 | 47 | 2.08 |
| **2566** | CAS - Institute of Geology and Geophysics | academic | China | 1 | - | 1 | 1 | 0.6 |
| **2567** | CAS - Institute of Metal Research | academic | China | 1 | - | 8 | 8 | 0.5 |
| **2568** | CAS - Institute of Physics | academic | China | 1 | - | 2 | 2 | 1.45 |
| **2569** | CAS - Research Center for Eco-Environmental Sciences | academic | China | 1 | - | 5 | 5 | 0.29 |
| **2570** | Central South University of Forestry & Technology | academic | China | 1 | - | 2 | 2 | 1.16 |
| **2571** | Chongqing Normal University | academic | China | 1 | - | 1 | 1 | 0.52 |
| **2572** | Harbin University of Commerce | academic | China | 1 | - | 26 | 26 | 1.54 |
| **2573** | Henan Institute of Science and Technology | academic | China | 1 | -100 | 38 | 38 | 0.67 |
| **2574** | Hunan Agricultural University | academic | China | 1 | - | 18 | 18 | 0.61 |
| **2575** | Jilin Normal University | academic | China | 1 | - | 4 | 4 | 0.09 |
| **2576** | Logistical Engineering University China | academic | China | 1 | - | 10 | 10 | 1.39 |
| **2577** | National Center for Nanoscience and Technology | academic | China | 1 | - | 3 | 3 | 0.51 |
| **2578** | National Meteorological Center | government | China | 1 | - | 10 | 10 | 5.16 |
| **2579** | North China University of Water Resources and Electric Power | academic | China | 1 | - | 6 | 6 | 0.59 |
| **2580** | Qiqihar University | academic | China | 1 | - | 3 | 3 | 0.65 |
| **2581** | Shanghai Academy of Agricultural Sciences | government | China | 1 | - | 3 | 3 | 0.39 |
| **2582** | Shenyang University | academic | China | 1 | - | 5 | 5 | 0.46 |
| **2583** | Suzhou University of Science and Technology | academic | China | 1 | - | 16 | 16 | 0.68 |
| **2584** | Wuyi University | academic | China | 1 | - | 23 | 23 | 1.72 |
| **2585** | Changchun University of Science and Technology | academic | China | 1 | - | 2 | 2 | 1.06 |
| **2586** | Bandung Institute of Technology | academic | Indonesia | 1 | - | 9 | 9 | 1.25 |
| **2587** | Dr. Harisingh Gour University, Sagar | academic | India | 1 | - | 13 | 13 | 0.64 |
| **2588** | M.S. University of Baroda | academic | India | 1 | - | 13 | 13 | 0.71 |
| **2589** | University of Rajasthan | academic | India | 1 | - | 0 | 0 | 0 |
| **2590** | Assam Agricultural University India | academic | India | 1 | - | 0 | 0 | 0 |
| **2591** | Defence Research and Development Establishment | government | India | 1 | -100 | 18 | 18 | 0.92 |
| **2592** | Gulbarga University | academic | India | 1 | -100 | 11 | 11 | 0.34 |
| **2593** | Institute for Plasma Research | government | India | 1 | - | 4 | 4 | 0.39 |
| **2594** | Inter University Accelerator Centre India | government | India | 1 | - | 0 | 0 | 0 |
| **2595** | International Centre for Genetic Engineering and Biotechnology India | government | India | 1 | - | 22 | 22 | 0.95 |
| **2596** | Mohan Lal Sukhadia University | academic | India | 1 | - | 48 | 48 | 1.13 |
| **2597** | Saurashtra University | academic | India | 1 | - | 1 | 1 | 0.43 |
| **2598** | Sri Krishnadevaraya University India | academic | India | 1 | - | 0 | 0 | 0 |
| **2599** | University of Agricultural Sciences, Bangalore | academic | India | 1 | - | 12 | 12 | 0.71 |
| **2600** | University of Jammu | academic | India | 1 | - | 5 | 5 | 2.24 |
| **2601** | Aichi Institute of Technology | academic | Japan | 1 | - | 3 | 3 | 1.66 |
| **2602** | Akita Prefectural University | academic | Japan | 1 | - | 3 | 3 | 0.39 |
| **2603** | Central Research Institute of Electric Power Industry | other | Japan | 1 | - | 1 | 1 | 0.09 |
| **2604** | Japan Fisheries Research and Education Agency | government | Japan | 1 | - | 8 | 8 | 4.32 |
| **2605** | Japan Agency for Marine-Earth Science and Technology | government | Japan | 1 | - | 0 | 0 | 0 |
| **2606** | Kanagawa University | academic | Japan | 1 | - | 2 | 2 | 0.21 |
| **2607** | Wakayama University | academic | Japan | 1 | - | 20 | 20 | 0.89 |
| **2608** | Fukui University of Technology | academic | Japan | 1 | - | 22 | 22 | 0.95 |
| **2609** | Fukuyama University | academic | Japan | 1 | - | 1 | 1 | 0.37 |
| **2610** | Iwate Prefectural University | academic | Japan | 1 | - | 11 | 11 | 0.73 |
| **2611** | Nissan Motor Co., Ltd. | corporate | Japan | 1 | - | 22 | 22 | 0.61 |
| **2612** | Osaka Kyoiku University | academic | Japan | 1 | - | 1 | 1 | 0.12 |
| **2613** | Tokyo Polytechnic University | academic | Japan | 1 | - | 0 | 0 | 0 |
| **2614** | The University of Shiga Prefecture | academic | Japan | 1 | - | 167 | 167 | 5.67 |
| **2615** | Andong National University | academic | South Korea | 1 | - | 14 | 14 | 1.1 |
| **2616** | Korea Atomic Energy Research Institute | government | South Korea | 1 | - | 24 | 24 | 2.07 |
| **2617** | Myongji University | academic | South Korea | 1 | - | 8 | 8 | 1.31 |
| **2618** | Sun Moon University | academic | South Korea | 1 | - | 16 | 16 | 2.39 |
| **2619** | Hankyong National University | academic | South Korea | 1 | -100 | 5 | 5 | 0.26 |
| **2620** | Korean Agency for Defense Development | government | South Korea | 1 | - | 2 | 2 | 0.08 |
| **2621** | Paichai University | academic | South Korea | 1 | - | 1 | 1 | 0.5 |
| **2622** | Seoul Women's University | academic | South Korea | 1 | - | 7 | 7 | 0.27 |
| **2623** | SK Corporation | corporate | South Korea | 1 | - | 0 | 0 | 0 |
| **2624** | Multimedia University | academic | Malaysia | 1 | - | 8 | 8 | 0.42 |
| **2625** | Lincoln University | academic | New Zealand | 1 | - | 31 | 31 | 1.55 |
| **2626** | Chienkuo Technology University Taiwan | academic | Taiwan | 1 | - | 5 | 5 | 0.31 |
| **2627** | National Chi Nan University | academic | Taiwan | 1 | - | 3 | 3 | 0.1 |
| **2628** | National Formosa University | academic | Taiwan | 1 | - | 2 | 2 | 1.03 |
| **2629** | National Taiwan Ocean University | academic | Taiwan | 1 | - | 4 | 4 | 2.89 |
| **2630** | National United University Taiwan | academic | Taiwan | 1 | - | 0 | 0 | 0 |
| **2631** | Tatung University | academic | Taiwan | 1 | - | 0 | 0 | 0 |
| **2632** | Huafan University | academic | Taiwan | 1 | - | 3 | 3 | 0.24 |
| **2633** | Kao Yuan University | academic | Taiwan | 1 | - | 0 | 0 | 0 |
| **2634** | National Synchrotron Radiation Research Center Taiwan | government | Taiwan | 1 | - | 22 | 22 | 1.36 |
| **2635** | Baku State University | academic | Azerbaijan | 1 | - | 16 | 16 | 1.91 |
| **2636** | University of the South Pacific | academic | Fiji | 1 | - | 21 | 21 | 1.91 |
| **2637** | National University of Uzbekistan named after Mirzo Ulugbek | academic | Uzbekistan | 1 | -100 | 2 | 2 | 1.06 |
| **2638** | Academy of Sciences of the Republic of Uzbekistan | academic | Uzbekistan | 1 | - | 4 | 4 | 0.72 |
| **2639** | Hanoi University of Science and Technology | academic | Viet Nam | 1 | - | 20 | 20 | 1.56 |
| **2640** | Ministry of Education and Science of the Republic of Kazakhstan | government | Kazakhstan | 1 | - | 0 | 0 | 0 |
| **2641** | Shanghai University of Electric Power | academic | China | 1 | - | 0 | 0 | 0 |
| **2642** | Tianjin University of Science & Technology | academic | China | 1 | - | 59 | 59 | 2.61 |
| **2643** | Tianjin University of Commerce | academic | China | 1 | - | 0 | 0 | 0 |
| **2644** | Mandalay University | academic | Myanmar | 1 | - | 4 | 4 | 0.28 |
| **2645** | CAS - Fujian Institute of Research on the Structure of Matter | academic | China | 1 | - | 23 | 23 | 1.96 |
| **2646** | CAS - Institute of Hydrobiology | academic | China | 1 | - | 8 | 8 | 0.59 |
| **2647** | CAS - Institute of Electronics | academic | China | 1 | - | 1 | 1 | 0.58 |
| **2648** | CAS - Chengdu Institute of Biology | academic | China | 1 | - | 15 | 15 | 0.48 |
| **2649** | CAS - Chengdu Institute of Organic Chemistry | academic | China | 1 | - | 1 | 1 | 0.52 |
| **2650** | CAS - Institute of Computing Technology | academic | China | 1 | - | 11 | 11 | 1.08 |
| **2651** | Otago Polytechnic | academic | New Zealand | 1 | - | 30 | 30 | 1.27 |
| **2652** | Medical Research Institute Colombo | academic | Sri Lanka | 1 | - | 3 | 3 | 0.58 |
| **2653** | Sabaragamuwa University of Sri Lanka | academic | Sri Lanka | 1 | - | 1 | 1 | 0.13 |
| **2654** | Sri Lanka Institute of Information Technology | academic | Sri Lanka | 1 | - | 4 | 4 | 2.19 |
| **2655** | Institute of Fundamental Studies Kandy | academic | Sri Lanka | 1 | - | 11 | 11 | 0.57 |
| **2656** | University of Moratuwa | academic | Sri Lanka | 1 | - | 1 | 1 | 0.53 |
| **2657** | Institut Pertanian Bogor | academic | Indonesia | 1 | - | 4 | 4 | 0.41 |
| **2658** | Beijing International Studies University | academic | China | 1 | - | 26 | 26 | 0.53 |
| **2659** | Central University of Finance and Economics | academic | China | 1 | - | 9 | 9 | 1.73 |
| **2660** | Kanagawa Institute of Technology | academic | Japan | 1 | - | 1 | 1 | 0.47 |
| **2661** | Institut Pasteur Korea | other | South Korea | 1 | - | 4 | 4 | 1.45 |
| **2662** | International University of Japan | academic | Japan | 1 | - | 15 | 15 | 1.07 |
| **2663** | Xi'an Polytechnic University | academic | China | 1 | - | 7 | 7 | 4.19 |
| **2664** | Satbayev University | academic | Kazakhstan | 1 | - | 0 | 0 | 0 |
| **2665** | King Mongkut's University of Technology North Bangkok | academic | Thailand | 1 | - | 0 | 0 | 0 |
| **2666** | University of Guam | academic | Guam | 1 | - | 1 | 1 | 0.17 |
| **2667** | University of Papua New Guinea | academic | Papua New Guinea | 1 | - | 18 | 18 | 0.85 |
| **2668** | New Energy and Industrial Technology Development Organization | government | Japan | 1 | -100 | 12 | 12 | 0.64 |
| **2669** | National Research Institute of Brewing | government | Japan | 1 | - | 2 | 2 | 0.19 |
| **2670** | Kazakh Ablai Khan University of International Relations and World Languages | academic | Kazakhstan | 1 | - | 3 | 3 | 0.21 |
| **2671** | Lincoln University College | academic | Malaysia | 1 | - | 0 | 0 | 0 |
| **2672** | CAS - Institut Pasteur of Shanghai | academic | China | 1 | - | 28 | 28 | 1.8 |
| **2673** | CAS - Ningbo Institute of Material Technology and Engineering | academic | China | 1 | - | 2 | 2 | 0.37 |
| **2674** | CAS - Suzhou Institute of Nano-Tech and Nano-Bionics | academic | China | 1 | - | 2 | 2 | 1.03 |
| **2675** | Bina Nusantara University | academic | Indonesia | 1 | - | 1 | 1 | 0.26 |
| **2676** | Universiti Kuala Lumpur | academic | Malaysia | 1 | - | 1 | 1 | 0.56 |
| **2677** | Malaysian Palm Oil Board | government | Malaysia | 1 | - | 17 | 17 | 0.98 |
| **2678** | Inner Mongolia Agricultural University | academic | China | 1 | - | 18 | 18 | 8.68 |
| **2679** | Inner Mongolia University of Technology | academic | China | 1 | - | 5 | 5 | 2.49 |
| **2680** | Minnan Normal University | academic | China | 1 | - | 1 | 1 | 0.18 |
| **2681** | Nanchang Hangkong University | academic | China | 1 | - | 3 | 3 | 0.32 |
| **2682** | Shanxi Agricultural University | academic | China | 1 | - | 27 | 27 | 1.81 |
| **2683** | Dongguan University of Technology | academic | China | 1 | - | 2 | 2 | 1.45 |
| **2684** | Dalian Ocean University | academic | China | 1 | - | 8 | 8 | 0.99 |
| **2685** | Fujian University of Technology | academic | China | 1 | -100 | 18 | 18 | 0.8 |
| **2686** | Quanzhou Normal University | academic | China | 1 | - | 35 | 35 | 1.5 |
| **2687** | Nanyang Normal University | academic | China | 1 | - | 3 | 3 | 1.5 |
| **2688** | Jining University | academic | China | 1 | - | 7 | 7 | 0.65 |
| **2689** | Binzhou University | academic | China | 1 | - | 1 | 1 | 0.17 |
| **2690** | Yancheng Teachers University | academic | China | 1 | - | 0 | 0 | 0 |
| **2691** | Bijapur Lingayat District Educational Association | academic | India | 1 | - | 905 | 905 | 174.03 |
| **2692** | Huawei Technologies Co., Ltd. | corporate | China | 1 | - | 0 | 0 | 0 |
| **2693** | MediaTek | corporate | Taiwan | 1 | - | 22 | 22 | 3.58 |
| **2694** | Bharat Heavy Electricals Ltd | corporate | India | 1 | - | 5 | 5 | 0.37 |
| **2695** | Mazda Motor | corporate | Japan | 1 | - | 15 | 15 | 2.12 |
| **2696** | ULVAC, Inc. | corporate | Japan | 1 | - | 2 | 2 | 0.09 |
| **2697** | IHI Corporation | corporate | Japan | 1 | - | 0 | 0 | 0 |
| **2698** | ENEOS Holdings, Inc. | corporate | Japan | 1 | - | 22 | 22 | 1.13 |
| **2699** | Honda Motor Co., Ltd. | corporate | Japan | 1 | - | 24 | 24 | 2.12 |
| **2700** | China Telecommunications | corporate | China | 1 | - | 53 | 53 | 2.21 |
| **2701** | China National Offshore Oil Corp | corporate | China | 1 | - | 21 | 21 | 1.51 |
| **2702** | Government of Kerala | government | India | 1 | - | 2 | 2 | 0.43 |
| **2703** | Japan Ministry of Agriculture, Forestry and Fisheries | government | Japan | 1 | - | 2 | 2 | 0.19 |
| **2704** | Sir Hurkisondas Nurrotumdas Hospital & Research Centre | medical | India | 1 | - | 7 | 7 | 0.31 |
| **2705** | Tropical Botanic Garden and Research Institute India | government | India | 1 | - | 28 | 28 | 1.22 |
| **2706** | Vietnamese Academy of Science and Technology | academic | Viet Nam | 1 | - | 0 | 0 | 0 |
| **2707** | National Applied Research Laboratories Taiwan | government | Taiwan | 1 | - | 8 | 8 | 0.57 |
| **2708** | Pakistan Institute of Development Economics | government | Pakistan | 1 | - | 1 | 1 | 0.21 |
| **2709** | Microbial Chemistry Research Foundation | other | Japan | 1 | - | 5 | 5 | 0.36 |
| **2710** | Bannari Amman Institute of Technology | academic | India | 1 | - | 1 | 1 | 0.29 |
| **2711** | Periyar University | academic | India | 1 | - | 22 | 22 | 1.18 |
| **2712** | Tamil Nadu Veterinary and Animal Sciences University | academic | India | 1 | - | 0 | 0 | 0 |
| **2713** | National Institute of Technology Warangal | academic | India | 1 | - | 84 | 84 | 1.68 |
| **2714** | Sardar Vallabhbhai National Institute of Technology Surat | academic | India | 1 | -100 | 4 | 4 | 0.35 |
| **2715** | Pt. Ravishankar Shukla University | academic | India | 1 | - | 0 | 0 | 0 |
| **2716** | Far East University | academic | Taiwan | 1 | - | 17 | 17 | 0.54 |
| **2717** | Southwest Forestry University | academic | China | 1 | - | 24 | 24 | 1.95 |
| **2718** | Isra University | academic | Pakistan | 1 | - | 10 | 10 | 0.95 |
| **2719** | Banasthali University | academic | India | 1 | - | 0 | 0 | 0 |
| **2720** | Netaji Subhas University of Technology | academic | India | 1 | - | 22 | 22 | 0.88 |
| **2721** | Ch. Charan Singh University | academic | India | 1 | - | 1 | 1 | 0.51 |
| **2722** | The University of Agriculture, Peshawar | academic | Pakistan | 1 | - | 1 | 1 | 0.52 |
| **2723** | Sher-e-Kashmir University of Agricultural Sciences and Technology of Kashmir | academic | India | 1 | - | 99 | 99 | 3.88 |
| **2724** | Weifang University | academic | China | 1 | - | 24 | 24 | 0.83 |
| **2725** | Communication University of China | academic | China | 1 | - | 2 | 2 | 0.96 |
| **2726** | Amorepacific Corporation | corporate | South Korea | 1 | - | 3 | 3 | 0.27 |
| **2727** | Indian Institute of Management Indore | academic | India | 1 | - | 11 | 11 | 0.2 |
| **2728** | Australian Research Council | government | Australia | 1 | - | 3 | 3 | 0 |
| **2729** | National Research Institute of Police Science | government | Japan | 1 | - | 36 | 36 | 1.48 |
| **2730** | Universitas Islam Indonesia | academic | Indonesia | 1 | - | 3 | 3 | 0.54 |
| **2731** | Kobe City University of Foreign Studies | academic | Japan | 1 | - | 21 | 21 | 0.76 |
| **2732** | Otaru University of Commerce | academic | Japan | 1 | - | 0 | 0 | 0 |
| **2733** | Rissho University | academic | Japan | 1 | - | 0 | 0 | 0 |
| **2734** | International Institute of Information Technology Bangalore | academic | India | 1 | - | 53 | 53 | 2.37 |
| **2735** | National Law School of India University | academic | India | 1 | - | 0 | 0 | 0 |
| **2736** | National Institute of Fashion Technology India | academic | India | 1 | - | 3 | 3 | 0.31 |
| **2737** | Shiseido Company, Limited | corporate | Japan | 1 | - | 24 | 24 | 1.19 |
| **2738** | Guangdong Institute of Eco-Environment and Soil Science | academic | China | 1 | - | 0 | 0 | 0 |
| **2739** | Institute of Life Sciences | academic | India | 1 | -100 | 31 | 31 | 0.96 |
| **2740** | Papua New Guinea Institute of Medical Research | academic | Papua New Guinea | 1 | - | 10 | 10 | 0.65 |
| **2741** | University of Teacher Education Fukuoka | academic | Japan | 1 | -100 | 0 | 0 | 0 |
| **2742** | Sri Sathya Sai University | academic | India | 1 | - | 3 | 3 | 0.55 |
| **2743** | Central University of Kerala | academic | India | 1 | - | 4 | 4 | 0.31 |
| **2744** | Central University of Jammu | academic | India | 1 | - | 4 | 4 | 0.73 |
| **2745** | Indian Institute of Science Education and Research Thiruvananthapuram | academic | India | 1 | - | 0 | 0 | 0 |
| **2746** | University of Central Punjab | academic | Pakistan | 1 | - | 32 | 32 | 5.79 |
| **2747** | Swami Ramanand Teerth Marathwada University | academic | India | 1 | - | 2 | 2 | 0.06 |
| **2748** | National Taipei University of Education | academic | Taiwan | 1 | - | 11 | 11 | 2.44 |
| **2749** | Eijkman Institute for Molecular Biology | academic | Indonesia | 1 | - | 12 | 12 | 0.74 |
| **2750** | Daikin Industries Ltd. | corporate | Japan | 1 | - | 3 | 3 | 0.29 |
| **2751** | Central University of Rajasthan | academic | India | 1 | - | 4 | 4 | 0.84 |
| **2752** | Zaozhuang University | academic | China | 1 | - | 26 | 26 | 1.46 |
| **2753** | National Institutes of Natural Sciences - Okazaki Institute for Integrative Bioscience | academic | Japan | 1 | -100 | 37 | 37 | 1.04 |
| **2754** | JAXA Institute of Space and Astronautical Science | government | Japan | 1 | - | 5 | 5 | 0.27 |
| **2755** | Institute of Advanced Study in Science and Technology India | academic | India | 1 | - | 0 | 0 | 0 |
| **2756** | Academia Sinica - Institute of Atomic and Molecular Sciences | academic | Taiwan | 1 | - | 6 | 6 | 0.39 |
| **2757** | Academia Sinica - Biodiversity Research Center | academic | Taiwan | 1 | - | 0 | 0 | 0 |
| **2758** | Academia Sinica - Agricultural Biotechnology Research Center | academic | Taiwan | 1 | - | 1 | 1 | 0.39 |
| **2759** | Kagawa Nutrition University | academic | Japan | 1 | -100 | 13 | 13 | 0.4 |
| **2760** | University of San Carlos - Philippines | academic | Philippines | 1 | - | 6 | 6 | 0.65 |
| **2761** | State Intellectual Property Office of the Peoples Republic of China | government | China | 1 | - | 1 | 1 | 0 |
| **2762** | Pharmaceuticals and Medical Devices Agency | government | Japan | 1 | - | 7 | 7 | 0.24 |
| **2763** | Pandit Dwarka Prasad Mishra Indian Institute of Information Technology, Design and Manufacturing Jabalpur | academic | India | 1 | - | 47 | 47 | 5.04 |
| **2764** | Indian Institute of Science Education and Research Mohali | academic | India | 1 | - | 0 | 0 | 0 |
| **2765** | Indian Institute of Management Rohtak | academic | India | 1 | - | 0 | 0 | 0 |
| **2766** | Central University of South Bihar | academic | India | 1 | - | 47 | 47 | 1.46 |
| **2767** | Central University of Haryana | academic | India | 1 | - | 24 | 24 | 0.82 |
| **2768** | Future University Hakodate | academic | Japan | 1 | -100 | 96 | 96 | 1.05 |
| **2769** | Fukuoka Women's University | academic | Japan | 1 | -100 | 12 | 12 | 0.41 |
| **2770** | Iryo Sosei University | academic | Japan | 1 | - | 1 | 1 | 0.74 |
| **2771** | Meiji Gakuin University | academic | Japan | 1 | - | 11 | 11 | 0.64 |
| **2772** | Niigata Institute of Technology | academic | Japan | 1 | -100 | 20 | 20 | 0.58 |
| **2773** | Shizuoka Institute of Science and Technology | academic | Japan | 1 | - | 16 | 16 | 1.41 |
| **2774** | Vel Tech University | academic | India | 1 | - | 1 | 1 | 0.58 |
| **2775** | Mehran University of Engineering & Technology | academic | Pakistan | 1 | - | 0 | 0 | 0 |
| **2776** | State University of Malang | academic | Indonesia | 1 | - | 1 | 1 | 0.56 |
| **2777** | Universitas Islam Malang | academic | Indonesia | 1 | - | 9 | 9 | 0.65 |
| **2778** | American University of Armenia | academic | Armenia | 1 | - | 5 | 5 | 0.96 |
| **2779** | Hokkaido Research Organization | government | Japan | 1 | - | 10 | 10 | 0.21 |
| **2780** | University Visvesvaraya College of Engineering | academic | India | 1 | - | 4 | 4 | 0.39 |
| **2781** | Mirpur University of Science and Technology | academic | Pakistan | 1 | - | 2 | 2 | 0.36 |
| **2782** | Chizhou University | academic | China | 1 | - | 3 | 3 | 1.5 |
| **2783** | Southwest University of Political Science and Law | academic | China | 1 | - | 8 | 8 | 0.26 |
| **2784** | TDK Corporation | corporate | Japan | 1 | - | 29 | 29 | 2.35 |
| **2785** | Yamaha Corporation | corporate | Japan | 1 | - | 1 | 1 | 0 |
| **2786** | National Institute of Development Administration | academic | Thailand | 1 | - | 1 | 1 | 0.52 |
| **2787** | Olympus Corporation | corporate | Japan | 1 | - | 15 | 15 | 1.53 |
| **2788** | Terumo Corporation | corporate | Japan | 1 | - | 0 | 0 | 0 |
| **2789** | Dai Nippon Printing Co., Ltd. | corporate | Japan | 1 | - | 13 | 13 | 2.56 |
| **2790** | Xuzhou Institute of Technology | academic | China | 1 | - | 3 | 3 | 1.55 |
| **2791** | Jiangsu University of Science and Technology | academic | China | 1 | - | 39 | 39 | 1.57 |
| **2792** | Berhampur University India | academic | India | 1 | - | 6 | 6 | 0.12 |
| **2793** | Barkatullah University | academic | India | 1 | - | 1 | 1 | 0.29 |
| **2794** | Ravenshaw University | academic | India | 1 | - | 2 | 2 | 0.13 |
| **2795** | Indraprastha Institute of Information Technology Delhi | academic | India | 1 | - | 0 | 0 | 0 |
| **2796** | Tripura University | academic | India | 1 | - | 61 | 61 | 1.6 |
| **2797** | Shri Mata Vaishno Devi University | academic | India | 1 | - | 2 | 2 | 0.47 |
| **2798** | Gurukula Kangri Vishwavidyalaya | academic | India | 1 | - | 9 | 9 | 1.65 |
| **2799** | Chhatrapati Shahu Ji Maharaj Kanpur University | academic | India | 1 | - | 1 | 1 | 0.19 |
| **2800** | Cancer Council NSW | academic | Australia | 1 | - | 1 | 1 | 0.47 |
| **2801** | Taranaki District Health Board | government | New Zealand | 1 | - | 18 | 18 | 1.51 |
| **2802** | The LifeFlight Foundation | other | Australia | 1 | - | 3 | 3 | 0.55 |
| **2803** | Chaudhary Devi Lal University, Sirsa | academic | India | 1 | - | 1 | 1 | 0.11 |
| **2804** | Dharmsinh Desai University | academic | India | 1 | - | 3 | 3 | 0.42 |
| **2805** | Hindustan Institute of Technology and Science | academic | India | 1 | - | 4 | 4 | 0.35 |
| **2806** | University of International Business | academic | Kazakhstan | 1 | - | 0 | 0 | 0 |
| **2807** | Ministry of Health of the Republic of Kazakhstan | government | Kazakhstan | 1 | - | 2 | 2 | 0.11 |
| **2808** | Takming University of Science and Technology | academic | Taiwan | 1 | - | 11 | 11 | 0.71 |
| **2809** | Bionics Institute | academic | Australia | 1 | -100 | 10 | 10 | 0.66 |
| **2810** | China University of Mining & Technology, Beijing | academic | China | 1 | - | 7 | 7 | 0.47 |
| **2811** | Taiyo Kagaku Co. Ltd. | corporate | Japan | 1 | - | 13 | 13 | 0.46 |
| **2812** | MLR Institute of Technology | academic | India | 1 | - | 1 | 1 | 0.35 |
| **2813** | Sai Life Sciences | corporate | India | 1 | - | 13 | 13 | 0.46 |
| **2814** | Avondale College | academic | Australia | 1 | -100 | 30 | 30 | 1.22 |
| **2815** | ICAR - National Institute of Veterinary Epidemiology and Disease Informatics, Hebbal | government | India | 1 | - | 4 | 4 | 0.39 |
| **2816** | Guru Angad Dev Veterinary and Animal Sciences University | academic | India | 1 | - | 12 | 12 | 0.87 |
| **2817** | Kerala University of Fisheries & Ocean Studies | academic | India | 1 | - | 7 | 7 | 0.33 |
| **2818** | AMET University | academic | India | 1 | - | 2 | 2 | 0.22 |
| **2819** | Academia Sinica - Research Center for Environmental Changes | academic | Taiwan | 1 | - | 27 | 27 | 2.19 |
| **2820** | Academia Sinica - Research Center for Information Technology Innovation | academic | Taiwan | 1 | - | 3 | 3 | 0.57 |
| **2821** | Shanxi University of Finance and Economics | academic | China | 1 | - | 1 | 1 | 0.56 |
| **2822** | Siam Cement Group | corporate | Thailand | 1 | - | 4 | 4 | 0.17 |
| **2823** | University of Petroleum and Energy Studies | academic | India | 1 | - | 11 | 11 | 1.18 |
| **2824** | Anhui Polytechnic University | academic | China | 1 | - | 19 | 19 | 0.25 |
| **2825** | Fuyang Normal University | academic | China | 1 | - | 14 | 14 | 0.54 |
| **2826** | Shaoguan University | academic | China | 1 | - | 8 | 8 | 5.7 |
| **2827** | Australian Wine Research Institute | academic | Australia | 1 | - | 74 | 74 | 2.22 |
| **2828** | Tissupath Pty Ltd. | corporate | Australia | 1 | - | 8 | 8 | 0.76 |
| **2829** | University Of Palangka Raya | academic | Indonesia | 1 | - | 0 | 0 | 0 |
| **2830** | Queensland Medical Laboratories Pathology | corporate | Australia | 1 | - | 12 | 12 | 1.09 |
| **2831** | Toppan Printing Co., Ltd. | corporate | Japan | 1 | - | 6 | 6 | 0.75 |
| **2832** | VCS Foundation | corporate | Australia | 1 | - | 0 | 0 | 0 |
| **2833** | Contech International Health Consultants | corporate | Pakistan | 1 | - | 1344 | 1344 | 64.51 |
| **2834** | Bened Biomedical Co., Ltd. | corporate | Taiwan | 1 | - | 68 | 68 | 4.23 |
| **2835** | Grape King Bio | corporate | Taiwan | 1 | - | 54 | 54 | 1.48 |
| **2836** | Abion Inc. | corporate | South Korea | 1 | - | 13 | 13 | 0.92 |
| **2837** | DNA Link | corporate | South Korea | 1 | - | 23 | 23 | 1.64 |
| **2838** | Gencurix, Inc. | corporate | South Korea | 1 | - | 7 | 7 | 0.54 |
| **2839** | Yokogawa Electric Corporation | corporate | Japan | 1 | -100 | 35 | 35 | 1.28 |
| **2840** | Zenyaku Kogyo Co., Ltd. | corporate | Japan | 1 | - | 18 | 18 | 0.91 |
| **2841** | JVCKenwood Corporation | corporate | Japan | 1 | - | 5 | 5 | 0.92 |
| **2842** | Kaken Pharmaceutical Co., Ltd. | corporate | Japan | 1 | - | 5 | 5 | 0.91 |
| **2843** | NIDEK Co., Ltd. | corporate | Japan | 1 | - | 12 | 12 | 0.96 |
| **2844** | Hyundai Heavy Industries Co., Ltd | corporate | South Korea | 1 | - | 10 | 10 | 2.29 |
| **2845** | Yuhan Corporation | corporate | South Korea | 1 | - | 2 | 2 | 1.22 |
| **2846** | Glory Ltd. | corporate | Japan | 1 | - | 1 | 1 | 0.53 |
| **2847** | HOYA Corporation | corporate | Japan | 1 | - | 4 | 4 | 0.74 |
| **2848** | New Zealand Defence Force | government | New Zealand | 1 | -100 | 109 | 109 | 3.15 |
| **2849** | Yakult Honsha Co., Ltd. | corporate | Japan | 1 | - | 39 | 39 | 3.85 |
| **2850** | Australian Red Cross Blood Service | other | Australia | 1 | - | 32 | 32 | 1.08 |
| **2851** | Ministry of Environment, Forest and Climate Change | government | India | 1 | - | 86 | 86 | 1.11 |
| **2852** | Cancer Council Queensland | academic | Australia | 1 | - | 0 | 0 | 0 |
| **2853** | Yuxi Normal University | academic | China | 1 | - | 0 | 0 | 0 |
| **2854** | Vinayaka Mission's Research Foundation | academic | India | 1 | - | 4 | 4 | 0.12 |
| **2855** | Hong Kong Government | government | Hong Kong | 1 | - | 57 | 57 | 2.58 |
| **2856** | Mymensingh Medical College Hospital | medical | Bangladesh | 1 | - | 0 | 0 | 0 |
| **2857** | National Institute of Technical Teachers' Training and Research, Chandigarh | academic | India | 1 | - | 6 | 6 | 0.26 |
| **2858** | Shin Nippon Biomedical Laboratories, Ltd. | corporate | Japan | 1 | - | 131 | 131 | 5.53 |
| **2859** | Semarang State University | academic | Indonesia | 1 | - | 1856 | 1856 | 130.04 |
| **2860** | National Heart Foundation of Australia | other | Australia | 1 | - | 7 | 7 | 0.18 |
| **2861** | Larsen and Toubro Limited | corporate | India | 1 | - | 0 | 0 | 0 |
| **2862** | Government College of Engineering and Ceramic Technology | academic | India | 1 | - | 2 | 2 | 0.67 |
| **2863** | Ramakrishna Mission Vivekananda University | academic | India | 1 | - | 0 | 0 | 0 |
| **2864** | Northern Territory Government of Australia | government | Australia | 1 | - | 14 | 14 | 0.78 |
| **2865** | Australian Radiation Protection and Nuclear Safety Agency | government | Australia | 1 | - | 26 | 26 | 2.01 |
| **2866** | National Heart Institute | academic | Malaysia | 1 | -100 | 256 | 256 | 9.93 |
| **2867** | Tamil Nadu Dr. J. Jayalalithaa Fisheries University | academic | India | 1 | - | 0 | 0 | 0 |
| **2868** | Nelson Marlborough Institute of Technology | other | New Zealand | 1 | - | 6 | 6 | 1.53 |
| **2869** | The Energy and Resources Institute India | academic | India | 1 | -100 | 5 | 5 | 0 |
| **2870** | State Tobacco Monopoly Administration, China Tobacco Corporation | government | China | 1 | - | 8 | 8 | 0.64 |
| **2871** | B.S. Abdur Rahman Crescent Institute of Science and Technology | academic | India | 1 | - | 3 | 3 | 0.28 |
| **2872** | Avinashilingam Institute for Home Science and Higher Education for Women | academic | India | 1 | - | 27 | 27 | 1.53 |
| **2873** | Uttarakhand Technical University | academic | India | 1 | - | 6 | 6 | 0.62 |
| **2874** | M.V. Hospital for Diabetes | medical | India | 1 | - | 12 | 12 | 3.8 |
| **2875** | Ministry of National Health Services, Regulations and Coordination | government | Pakistan | 1 | - | 32 | 32 | 1.85 |
| **2876** | Jaya Educational Trust | academic | India | 1 | - | 17 | 17 | 0.91 |
| **2877** | Rajiv Gandhi University of Knowledge and Technology | academic | India | 1 | - | 5 | 5 | 0.54 |
| **2878** | St. Aloysius College | academic | India | 1 | - | 3 | 3 | 0.19 |
| **2879** | Bayside Health | medical | Australia | 1 | - | 184 | 184 | 7.01 |
| **2880** | Institute of Space Technology | academic | Pakistan | 1 | - | 6 | 6 | 0.32 |
| **2881** | Daffodil International University | academic | Bangladesh | 1 | - | 0 | 0 | 0 |
| **2882** | Shikoku University | academic | Japan | 1 | -100 | 161 | 161 | 2.65 |
| **2883** | Mongolian University of Life Sciences | academic | Mongolia | 1 | - | 7 | 7 | 0.6 |
| **2884** | Alibaba Group Holding Ltd. | corporate | China | 1 | - | 0 | 0 | 0 |
| **2885** | Baba Ghulam Shah Badshah University | academic | India | 1 | - | 16 | 16 | 1.34 |
| **2886** | Ashoka University | academic | India | 1 | - | 5 | 5 | 0.9 |
| **2887** | Sarsen Amanzholov East Kazakhstan State University | academic | Kazakhstan | 1 | - | 0 | 0 | 0 |
| **2888** | RK University | academic | India | 1 | - | 2 | 2 | 0.17 |
| **2889** | Indira Gandhi National Tribal University | academic | India | 1 | - | 3 | 3 | 0.15 |
| **2890** | St. Peter's Institute of Pharmaceutical Sciences | academic | India | 1 | -100 | 8 | 8 | 0.21 |
| **2891** | Osaka Ohtani University | academic | Japan | 1 | - | 30 | 30 | 2.22 |
| **2892** | The Australian Psychological Society Ltd. | other | Australia | 1 | - | 7 | 7 | 0.59 |
| **2893** | K.R. Mangalam University | academic | India | 1 | - | 37 | 37 | 3.31 |
| **2894** | Ministry of Ayurveda, Yoga and Naturopathy, Unani, Siddha, Sowa Rigpa and Homoeopathy | government | India | 1 | - | 39 | 39 | 1.54 |
| **2895** | K. Zhubanov Aktobe Regional State University | academic | Kazakhstan | 1 | - | 3 | 3 | 0.32 |
| **2896** | BPS Govt. Medical College for Women | academic | India | 1 | - | 2 | 2 | 1.07 |
| **2897** | Shaheed Zulfikar Ali Bhutto Institute of Science and Technology, Karachi | academic | Pakistan | 1 | - | 2 | 2 | 0.1 |
| **2898** | Delhi Institute of Pharmaceutical Sciences and Research | academic | India | 1 | - | 0 | 0 | 0 |
| **2899** | Mokwon University | academic | South Korea | 1 | - | 23 | 23 | 1.06 |
| **2900** | Discovery Technology | corporate | Australia | 1 | - | 0 | 0 | 0 |
| **2901** | B.M.S. College of Engineering | academic | India | 1 | - | 23 | 23 | 1.59 |
| **2902** | Nanyang Institute of Technology | academic | China | 1 | - | 0 | 0 | 0 |
| **2903** | Guangdong Polytechnic Normal University | academic | China | 1 | - | 8 | 8 | 0.45 |
| **2904** | National University of Computer and Emerging Sciences, Islamabad | academic | Pakistan | 1 | - | 0 | 0 | 0 |
| **2905** | Riau University | academic | Indonesia | 1 | - | 3 | 3 | 0.42 |
| **2906** | Haluoleo University | academic | Indonesia | 1 | - | 24 | 24 | 1.24 |
| **2907** | Jiangsu Institute of Parasitic Diseases | government | China | 1 | - | 6 | 6 | 0.48 |
| **2908** | Shanghai Center for Bioinformation Technology | government | China | 1 | - | 7 | 7 | 0.69 |
| **2909** | Wenzhou Polytechnic | academic | China | 1 | - | 39 | 39 | 2.63 |
| **2910** | PLA No. 105 Hospital | medical | China | 1 | -100 | 7 | 7 | 0.39 |
| **2911** | SanJiang University | academic | China | 1 | - | 26 | 26 | 0.81 |
| **2912** | PLA No. 180 Hospital | medical | China | 1 | - | 7 | 7 | 0.57 |
| **2913** | Chongqing University of Arts and Science | academic | China | 1 | - | 15 | 15 | 0.21 |
| **2914** | Xinjiang Agriculture University | academic | China | 1 | - | 9 | 9 | 0.49 |
| **2915** | Zhoukou Normal University | academic | China | 1 | - | 36 | 36 | 0.7 |
| **2916** | Kohat University of Science and Technology | academic | Pakistan | 1 | - | 3 | 3 | 1.3 |
| **2917** | Henan Academy of Agricultural Sciences | academic | China | 1 | - | 11 | 11 | 0.63 |
| **2918** | Asia Pacific Center for Theoretical Physics | government | South Korea | 1 | - | 38 | 38 | 2.03 |
| **2919** | National Quemoy University | academic | Taiwan | 1 | - | 4 | 4 | 0 |
| **2920** | Hwa Hsia University of Technology | academic | Taiwan | 1 | - | 193 | 193 | 2.67 |
| **2921** | Central Police University, Taiwan | academic | Taiwan | 1 | - | 3 | 3 | 0.09 |
| **2922** | Toko University | academic | Taiwan | 1 | - | 11 | 11 | 0.13 |
| **2923** | Hsin Sheng College of Medical Care and Management | academic | Taiwan | 1 | - | 4 | 4 | 0 |
| **2924** | Ministry of Labor, Taiwan | government | Taiwan | 1 | - | 5 | 5 | 0.48 |
| **2925** | University of Surabaya | academic | Indonesia | 1 | - | 1 | 1 | 0 |
| **2926** | Tarumanagara University | academic | Indonesia | 1 | - | 9 | 9 | 0.61 |
| **2927** | Gunadarma University | academic | Indonesia | 1 | - | 0 | 0 | 0 |
| **2928** | Sanata Dharma University | academic | Indonesia | 1 | - | 7 | 7 | 0.29 |
| **2929** | Eastern Liaoning University | academic | China | 1 | - | 3 | 3 | 0.51 |
| **2930** | East West University | academic | Bangladesh | 1 | - | 22 | 22 | 1.69 |
| **2931** | Jiaying University | academic | China | 1 | - | 2 | 2 | 0.58 |
| **2932** | Tongmyong University | academic | South Korea | 1 | - | 2 | 2 | 0.43 |
| **2933** | Mianyang Normal University | academic | China | 1 | - | 14 | 14 | 0.49 |
| **2934** | Hexi University | academic | China | 1 | - | 2 | 2 | 1.45 |
| **2935** | Huizhou University | academic | China | 1 | - | 1 | 1 | 0.09 |
| **2936** | Hanshan Normal University | academic | China | 1 | - | 33 | 33 | 1.66 |
| **2937** | Beijing Electronic Science and Technology Institute | academic | China | 1 | - | 1 | 1 | 0.17 |
| **2938** | Oil and Natural Gas Corporation Ltd. | corporate | India | 1 | - | 5 | 5 | 0.57 |
| **2939** | Tadulako University | academic | Indonesia | 1 | - | 1 | 1 | 0.11 |
| **2940** | Rajendra Memorial Research Institute of Medical Sciences | medical | India | 1 | - | 2 | 2 | 0.33 |
| **2941** | Japan Automobile Research Institute | other | Japan | 1 | - | 19 | 19 | 1.93 |
| **2942** | Madhav Institute of Technology and Science | academic | India | 1 | - | 151 | 151 | 4.77 |
| **2943** | Chengdu Normal University | academic | China | 1 | - | 6 | 6 | 1.04 |
| **2944** | Neusoft Corporation | corporate | China | 1 | - | 0 | 0 | 0 |
| **2945** | International Islamic University Chittagong | academic | Bangladesh | 1 | - | 17 | 17 | 1.62 |
| **2946** | Aichi University | academic | Japan | 1 | - | 73 | 73 | 4.8 |
| **2947** | Satya Wacana Christian University | academic | Indonesia | 1 | - | 24 | 24 | 1.24 |
| **2948** | Syngene International Ltd. | corporate | India | 1 | - | 56 | 56 | 4.4 |
| **2949** | Xi'an Peihua University | academic | China | 1 | - | 54 | 54 | 3.52 |
| **2950** | Chinese People's Police University | academic | China | 1 | - | 5 | 5 | 1.24 |
| **2951** | Sindh Institute of Urology and Transplantation | medical | Pakistan | 1 | - | 33 | 33 | 13.54 |
| **2952** | Cadila Healthcare Ltd. | corporate | India | 1 | - | 1 | 1 | 0.11 |
| **2953** | Pabna University of Science and Technology | academic | Bangladesh | 1 | - | 905 | 905 | 174.03 |
| **2954** | Adikavi Nannaya University | academic | India | 1 | -100 | 6 | 6 | 0.19 |
| **2955** | Pokhara University | academic | Nepal | 1 | - | 8 | 8 | 0.7 |
| **2956** | Baddi University of Emerging Sciences and Technologies | academic | India | 1 | - | 52 | 52 | 4.17 |
| **2957** | National University of Modern Languages | academic | Pakistan | 1 | - | 7 | 7 | 1.5 |
| **2958** | Tokyo Keizai University | academic | Japan | 1 | - | 13 | 13 | 2.53 |
| **2959** | Ryutsu Keizai University | academic | Japan | 1 | - | 1 | 1 | 0.26 |
| **2960** | Nara University | academic | Japan | 1 | - | 0 | 0 | 0 |
| **2961** | Otsuma Women's University | academic | Japan | 1 | - | 17 | 17 | 0.56 |
| **2962** | University of Mataram | academic | Indonesia | 1 | - | 0 | 0 | 0 |
| **2963** | Tsan Yuk Hospital | medical | Hong Kong | 1 | - | 17 | 17 | 1.34 |
| **2964** | Wuhan Asia Heart Hospital | medical | China | 1 | - | 4 | 4 | 0.2 |
| **2965** | Taixing People's Hospital | medical | China | 1 | - | 11 | 11 | 0.35 |
| **2966** | PLA No. 202 Hospital | medical | China | 1 | - | 10 | 10 | 0.7 |
| **2967** | PLA No. 411 Hospital | medical | China | 1 | -100 | 54 | 54 | 1.74 |
| **2968** | PLA No. 463 Hospital | medical | China | 1 | -100 | 36 | 36 | 1.09 |
| **2969** | University of Human Environments | academic | Japan | 1 | - | 1 | 1 | 0.6 |
| **2970** | Bushfire and Natural Hazards CRC | other | Australia | 1 | - | 38 | 38 | 0.91 |
| **2971** | Vuno, Inc. | corporate | South Korea | 1 | - | 21 | 21 | 2.14 |
| **2972** | St John Ambulance Western Australia | other | Australia | 1 | - | 20 | 20 | 1.23 |
| **2973** | Silverline Research | corporate | Australia | 1 | - | 19 | 19 | 1.17 |
| **2974** | Yamanashi Gakuin University | academic | Japan | 1 | - | 1 | 1 | 0.61 |
| **2975** | The Women University Multan | academic | Pakistan | 1 | - | 7 | 7 | 1.5 |
| **2976** | Tama University | academic | Japan | 1 | - | 17 | 17 | 0.38 |
| **2977** | University of Jambi | academic | India | 1 | - | 0 | 0 | 0 |
| **2978** | Caritas Institute of Higher Education | academic | Hong Kong | 1 | - | 6 | 6 | 1.05 |
| **2979** | Sandor Life Sciences Pvt. Ltd. | corporate | India | 1 | - | 13 | 13 | 0.71 |
| **2980** | Shangrao Normal University | academic | China | 1 | - | 8 | 8 | 0.64 |
| **2981** | Lambung Mangkurat University | academic | Indonesia | 1 | - | 0 | 0 | 0 |
| **2982** | Lingnan Normal University | academic | China | 1 | - | 51 | 51 | 1.29 |
| **2983** | National Forensic Service | government | South Korea | 1 | -100 | 4 | 4 | 0.11 |
| **2984** | Gyeongju University | academic | South Korea | 1 | -100 | 6 | 6 | 0.46 |
| **2985** | Rajagiri School of Engineering & Technology | academic | India | 1 | - | 26 | 26 | 1.89 |
| **2986** | Shaheed Zulfikar Ali Bhutto Institute of Science and Technology, Islamabad | academic | Pakistan | 1 | - | 39 | 39 | 1.57 |
| **2987** | Institute of Engineering and Technology, Lucknow | academic | India | 1 | - | 0 | 0 | 0 |
| **2988** | Sri Padmavati Women's University | academic | India | 1 | - | 48 | 48 | 2.45 |
| **2989** | PA Engineering College Mangalore | academic | India | 1 | - | 13 | 13 | 0.71 |
| **2990** | Sir M. Visvesvaraya Institute of Technology | academic | India | 1 | -100 | 8 | 8 | 0.26 |
| **2991** | Yung Ta Institute of Technology and Commerce | academic | Taiwan | 1 | -100 | 2 | 2 | 0 |
| **2992** | CSIR - Institute of Microbial Technology | government | India | 1 | - | 17 | 17 | 0.79 |
| **2993** | CSIR - National Botanical Research Institute | government | India | 1 | - | 2 | 2 | 0.92 |
| **2994** | CSIR - Indian Institute of Integrative Medicine | government | India | 1 | - | 23 | 23 | 0.64 |
| **2995** | CSIR - North-East Institute of Science and Technology | government | India | 1 | - | 8 | 8 | 0.86 |
| **2996** | CSIR - National Institute of Science, Technology and Development Studies | government | India | 1 | - | 0 | 0 | 0 |
| **2997** | Kaifeng University | academic | China | 1 | - | 8 | 8 | 0.43 |
| **2998** | International University of Business, Agriculture and Technology | academic | Bangladesh | 1 | - | 3 | 3 | 1.5 |
| **2999** | Commission on Science and Technology for Sustainable Development in the South | academic | Pakistan | 1 | - | 5 | 5 | 2.9 |
| **3000** | Fuji Xerox Co., Ltd. | corporate | Japan | 1 | - | 1 | 1 | 0.18 |

**Table 3:**	The list of 3000 universities with sector, scholarly output, growth rate (%), total citations, citations per publications and field-weighted citation impact.

| **S#** | **Name** | **Scholarly Output** | **Most recent publication** | **Citations** | **Citations per Publication** | **Field-Weighted Citation Impact** | **h-index** |
| --- | --- | --- | --- | --- | --- | --- | --- |
| **1** | Masters, Colin L. | 155 | 2022 | 9421 | 60.8 | 3.48 | 135 |
| **2** | Hodges, John R. | 139 | 2020 | 5775 | 41.5 | 2.28 | 133 |
| **3** | Berkovic, Samuel F. | 159 | 2022 | 8684 | 54.6 | 2.62 | 117 |
| **4** | Feigin, Valery L. | 110 | 2022 | 15920 | 144.7 | 9.44 | 115 |
| **5** | Sachdev, Perminder S. | 114 | 2022 | 9670 | 84.8 | 4.92 | 115 |
| **6** | Donnan, Geoffrey Alan | 129 | 2022 | 6564 | 50.9 | 3.35 | 115 |
| **7** | Scheffer, Ingrid E. | 225 | 2022 | 18328 | 81.5 | 4.4 | 112 |
| **8** | Halliday, Glenda Margaret | 191 | 2022 | 14222 | 74.5 | 3.3 | 107 |
| **9** | Davis, Stephen M. | 146 | 2022 | 6883 | 47.1 | 3.26 | 103 |
| **10** | Anderson, Craig S. | 146 | 2022 | 5338 | 36.6 | 1.9 | 94 |
| **11** | Rowe, Christopher C. | 107 | 2022 | 15271 | 142.7 | 5.92 | 90 |
| **12** | Sobue, Gen | 165 | 2021 | 4165 | 25.2 | 1.39 | 90 |
| **13** | Villemagne, Victor L. | 106 | 2022 | 7281 | 68.7 | 4.04 | 88 |
| **14** | Gong, Qiyong | 154 | 2022 | 3234 | 21 | 1.44 | 86 |
| **15** | Hattori, Nobutaka | 320 | 2022 | 4868 | 15.2 | 1.14 | 85 |
| **16** | Kiernan, Matthew C. | 292 | 2022 | 8846 | 30.3 | 2.09 | 83 |
| **17** | Tuji, Shoji | 101 | 2022 | 2036 | 20.2 | 1.49 | 83 |
| **18** | Venketasubramanian, Narayanaswamy | 117 | 2022 | 11836 | 101.2 | 7.61 | 78 |
| **19** | Parsons, Mark W. | 151 | 2022 | 3921 | 26 | 1.96 | 77 |
| **20** | O'Brien, Terence J. | 259 | 2022 | 5849 | 22.6 | 1.69 | 76 |
| **21** | Kuwabara, S. | 177 | 2022 | 4158 | 23.5 | 1.4 | 74 |
| **22** | Nishino, Ichizo | 203 | 2022 | 2817 | 13.9 | 1.3 | 74 |
| **23** | Wong, Kasing Lawrence | 156 | 2022 | 4930 | 31.6 | 1.46 | 73 |
| **24** | Levi, Christopher Royce | 137 | 2022 | 4468 | 32.6 | 1.88 | 73 |
| **25** | Dale, Russell C. | 133 | 2022 | 8213 | 61.8 | 3.56 | 71 |
| **26** | Wang, Shuu Jiun | 169 | 2022 | 6723 | 39.8 | 1.97 | 70 |
| **27** | Tan, Eng King | 147 | 2022 | 2942 | 20 | 1.8 | 70 |
| **28** | Ponsford, Jennie L. | 113 | 2022 | 3376 | 29.9 | 1.33 | 69 |
| **29** | Takahashi, Ryousuke | 148 | 2022 | 2100 | 14.2 | 0.86 | 68 |
| **30** | Chen, Christopher P. | 132 | 2022 | 7493 | 56.8 | 2.79 | 67 |
| **31** | Wang, Yongjun | 387 | 2022 | 9046 | 23.4 | 1.78 | 65 |
| **32** | Mok, Chung Tong Vincent | 168 | 2022 | 7690 | 45.8 | 2.23 | 65 |
| **33** | Nakamura, Masaya | 111 | 2022 | 1320 | 11.9 | 0.89 | 65 |
| **34** | Lewis, Simon John Geoffrey | 138 | 2022 | 6766 | 49 | 2.37 | 64 |
| **35** | Abe, Kohji | 160 | 2021 | 2001 | 12.5 | 0.76 | 64 |
| **36** | Kwan, Patrick K.L. | 155 | 2022 | 4028 | 26 | 1.9 | 63 |
| **37** | Houkin, Kiyohiro | 134 | 2022 | 2461 | 18.4 | 1.04 | 63 |
| **38** | Campbell, Bruce C.V. | 184 | 2022 | 6004 | 32.6 | 3.52 | 62 |
| **39** | Kim, Jong-sung | 120 | 2022 | 2654 | 22.1 | 1.06 | 62 |
| **40** | Ugawa, Yoshikazu | 107 | 2022 | 4829 | 45.1 | 1.89 | 61 |
| **41** | Chu, Kon | 103 | 2022 | 2533 | 24.6 | 1.35 | 61 |
| **42** | Piguet, Olivier | 108 | 2022 | 3766 | 34.9 | 1.99 | 60 |
| **43** | Kira, Junichi | 195 | 2022 | 2531 | 13 | 0.88 | 60 |
| **44** | Vucic, Steve | 182 | 2022 | 6741 | 37 | 2.34 | 59 |
| **45** | Kakita, Akiyoshi | 142 | 2022 | 2386 | 16.8 | 1.16 | 59 |
| **46** | Ji, Xunming | 264 | 2022 | 5604 | 21.2 | 1.46 | 58 |
| **47** | Katsuno, Masahisa | 158 | 2022 | 2742 | 17.4 | 1.42 | 58 |
| **48** | Butzkueven, Helmut | 144 | 2022 | 4533 | 31.5 | 2.14 | 57 |
| **49** | Taylor, Bruce V.M. | 141 | 2022 | 3526 | 25 | 1.4 | 57 |
| **50** | Bang, Oh-young | 124 | 2022 | 3410 | 27.5 | 1.71 | 57 |
| **51** | Lee, Sang kun | 124 | 2022 | 2688 | 21.7 | 1.3 | 57 |
| **52** | Mochizuki, Hideki | 137 | 2022 | 2028 | 14.8 | 0.97 | 57 |
| **53** | Tominaga, Teiji | 401 | 2022 | 5205 | 13 | 0.86 | 56 |
| **54** | Aoki, Masashi | 135 | 2022 | 3755 | 27.8 | 2.15 | 56 |
| **55** | Chen, Shengdi | 157 | 2022 | 2986 | 19 | 1.21 | 56 |
| **56** | Na, Duk-lyul | 117 | 2022 | 2744 | 23.5 | 1.52 | 56 |
| **57** | Lee, Soon-tae ‑t | 118 | 2022 | 2554 | 21.6 | 1.42 | 56 |
| **58** | Matsumoto, Morio | 142 | 2022 | 1970 | 13.9 | 0.98 | 56 |
| **59** | Ihara, Masafumi | 107 | 2022 | 1746 | 16.3 | 1.31 | 56 |
| **60** | Ando, Yukio | 115 | 2021 | 1242 | 10.8 | 0.73 | 56 |
| **61** | Cook, Mark James | 111 | 2022 | 3642 | 32.8 | 1.95 | 54 |
| **62** | Miyamoto, Susumu | 192 | 2022 | 3261 | 17 | 1.18 | 54 |
| **63** | Kaji, Ryuji | 136 | 2020 | 2360 | 17.4 | 0.92 | 54 |
| **64** | Kuroda, Satoshi | 150 | 2022 | 2255 | 15 | 1.29 | 54 |
| **65** | Yoshida, Mari | 129 | 2022 | 2100 | 16.3 | 1.47 | 54 |
| **66** | Nakashima, Ichiro | 142 | 2022 | 4530 | 31.9 | 2.72 | 53 |
| **67** | Saitsu, Hirotomo | 105 | 2022 | 2457 | 23.4 | 1.57 | 53 |
| **68** | Jung, Keun-hwa ‑h | 107 | 2022 | 2224 | 20.8 | 1.09 | 53 |
| **69** | Sharma, Vijay Kumar | 127 | 2022 | 2564 | 20.2 | 1.72 | 52 |
| **70** | Fujimura, Miki | 147 | 2022 | 2130 | 14.5 | 1.22 | 52 |
| **71** | Ikeda, Akio | 117 | 2022 | 1127 | 9.6 | 0.64 | 52 |
| **72** | Toyoda, Kazunori | 223 | 2022 | 4279 | 19.2 | 1.55 | 51 |
| **73** | Heo, Ji-hoe | 118 | 2022 | 2635 | 22.3 | 1.7 | 51 |
| **74** | Hirata, Koichi | 100 | 2022 | 951 | 9.5 | 0.88 | 51 |
| **75** | Wang, Yilong | 254 | 2022 | 6499 | 25.6 | 1.68 | 50 |
| **76** | Barnett, Michael H. | 137 | 2022 | 4924 | 35.9 | 2.22 | 50 |
| **77** | Misra, Ushakant | 125 | 2022 | 1911 | 15.3 | 0.78 | 50 |
| **78** | Kim, Seung-ki | 134 | 2022 | 1856 | 13.9 | 0.88 | 50 |
| **79** | Nakada, Mitsutoshi | 131 | 2022 | 1524 | 11.6 | 0.76 | 50 |
| **80** | Zhou, Dong | 234 | 2022 | 3905 | 16.7 | 1.19 | 49 |
| **81** | Kim, Ji-soon | 240 | 2022 | 3561 | 14.8 | 1.1 | 49 |
| **82** | Chung, Chun Kee | 158 | 2022 | 2591 | 16.4 | 0.93 | 49 |
| **83** | Tang, Beisha | 190 | 2022 | 2418 | 12.7 | 1.09 | 49 |
| **84** | Kalita, Jayantee | 127 | 2022 | 1937 | 15.3 | 0.77 | 49 |
| **85** | Kwon, Sun-uck | 109 | 2022 | 1811 | 16.6 | 1.07 | 49 |
| **86** | Date, Isao | 125 | 2022 | 1458 | 11.7 | 0.98 | 49 |
| **87** | Wang, Kyuchang | 118 | 2022 | 1367 | 11.6 | 0.68 | 49 |
| **88** | Liu, Liping | 217 | 2022 | 4635 | 21.4 | 1.44 | 48 |
| **89** | Phan, Kevin | 144 | 2022 | 4070 | 28.3 | 1.52 | 48 |
| **90** | Mitchell, Peter J. | 120 | 2022 | 3470 | 28.9 | 2.95 | 48 |
| **91** | Bae, Hee-joon | 159 | 2022 | 3362 | 21.1 | 1.5 | 48 |
| **92** | Seo, Sang-won | 108 | 2022 | 2937 | 27.2 | 1.86 | 48 |
| **93** | Lee, Phil-hyu | 133 | 2022 | 2047 | 15.4 | 1.07 | 48 |
| **94** | Kimura, Kazumi | 162 | 2022 | 1966 | 12.1 | 0.72 | 48 |
| **95** | Yoshimura, Shinichi | 140 | 2022 | 1454 | 10.4 | 0.91 | 48 |
| **96** | Churilov, Leonid | 141 | 2022 | 3169 | 22.5 | 1.9 | 47 |
| **97** | Kusunoki, Susumu | 125 | 2022 | 2646 | 21.2 | 1.43 | 47 |
| **98** | Saito, Nobuhito | 206 | 2022 | 2624 | 12.7 | 0.86 | 47 |
| **99** | Goel, Atul H. | 262 | 2022 | 2354 | 9 | 1.34 | 47 |
| **100** | Park, Sung-hye Hye | 109 | 2022 | 1834 | 16.8 | 1.3 | 47 |
| **101** | Zhao, Xingquan Qquan | 259 | 2022 | 4747 | 18.3 | 1.38 | 46 |
| **102** | Fung, Victor S.C. | 111 | 2022 | 4610 | 41.5 | 2.45 | 46 |
| **103** | Liu, M. | 115 | 2022 | 3433 | 29.9 | 1.5 | 46 |
| **104** | Jeon, Beomseok | 218 | 2022 | 3094 | 14.2 | 1.08 | 46 |
| **105** | Matsuyama, Yukihiro | 116 | 2022 | 1647 | 14.2 | 1.1 | 46 |
| **106** | Lechner-Scott, Jeannette S. | 120 | 2022 | 3731 | 31.1 | 2.53 | 45 |
| **107** | Takahashi, Toshiyuki | 169 | 2022 | 3088 | 18.3 | 1.43 | 45 |
| **108** | McCombe, Pamela A. | 100 | 2022 | 2430 | 24.3 | 1.66 | 45 |
| **109** | Kurisu, Kaoru | 158 | 2022 | 1518 | 9.6 | 0.74 | 45 |
| **110** | Ogasawara, Kuniaki | 136 | 2022 | 1453 | 10.7 | 1.22 | 45 |
| **111** | Morita, Akio | 171 | 2022 | 2666 | 15.6 | 0.9 | 44 |
| **112** | Imagama, Shiro | 172 | 2022 | 2001 | 11.6 | 0.99 | 44 |
| **113** | Watanabe, Kota | 124 | 2022 | 1653 | 13.3 | 0.98 | 43 |
| **114** | Tsuboi, Yoshio | 108 | 2022 | 1112 | 10.3 | 0.98 | 43 |
| **115** | Hong, Keun-sik | 104 | 2022 | 2317 | 22.3 | 1.37 | 42 |
| **116** | Kumabe, Toshihiro | 105 | 2022 | 1366 | 13 | 0.86 | 42 |
| **117** | Yan, Bernard | 125 | 2022 | 2677 | 21.4 | 1.94 | 41 |
| **118** | Qiu, Yong | 176 | 2022 | 1984 | 11.3 | 0.82 | 41 |
| **119** | Matsumura, Akira | 105 | 2020 | 1319 | 12.6 | 0.59 | 41 |
| **120** | Cui, Liying | 311 | 2022 | 3321 | 10.7 | 0.82 | 40 |
| **121** | Pal, Pramod Kumar | 221 | 2022 | 2562 | 11.6 | 0.89 | 40 |
| **122** | Nam, Hyo-suk | 100 | 2022 | 2169 | 21.7 | 1.54 | 40 |
| **123** | Kim, Jeong Eun | 125 | 2022 | 2077 | 16.6 | 0.92 | 40 |
| **124** | Sohn, Youngho | 126 | 2022 | 1885 | 15 | 1.04 | 40 |
| **125** | Garg, Ravindra Kumar | 122 | 2022 | 1604 | 13.1 | 0.97 | 40 |
| **126** | Suzuki, Hidenori | 105 | 2022 | 1360 | 13 | 1.4 | 40 |
| **127** | Shang, Huifang | 221 | 2022 | 3184 | 14.4 | 0.9 | 39 |
| **128** | Han, Moon Ku | 117 | 2022 | 2098 | 17.9 | 1 | 39 |
| **129** | Mikuni, Nobuhiro | 157 | 2022 | 1456 | 9.3 | 0.67 | 39 |
| **130** | Sharma, Mehar Chand | 116 | 2022 | 1010 | 8.7 | 0.57 | 39 |
| **131** | Tripathi, Manjari | 203 | 2022 | 2988 | 14.7 | 1.68 | 38 |
| **132** | Lee, Byung-chul | 105 | 2022 | 2209 | 21 | 1.4 | 38 |
| **133** | Takahashi, Yukitoshi | 100 | 2022 | 1200 | 12 | 0.61 | 38 |
| **134** | Li, Hao | 125 | 2022 | 3918 | 31.3 | 2.11 | 37 |
| **135** | Zhao, Jizong | 178 | 2022 | 2630 | 14.8 | 1.25 | 37 |
| **136** | Kim, Beom Joon | 112 | 2022 | 2098 | 18.7 | 1.42 | 37 |
| **137** | Chandra, Ronil V. | 104 | 2022 | 1539 | 14.8 | 1.25 | 37 |
| **138** | Mahadevan, Anita | 138 | 2022 | 1097 | 7.9 | 0.61 | 37 |
| **139** | Pan, Yuesong | 163 | 2022 | 3095 | 19 | 1.47 | 36 |
| **140** | Li, Youxiang | 127 | 2022 | 1568 | 12.3 | 0.68 | 36 |
| **141** | Izumi, Yuishin | 123 | 2022 | 1556 | 12.7 | 0.88 | 36 |
| **142** | Sharma, Bhawani Shankar | 129 | 2022 | 1496 | 11.6 | 0.67 | 36 |
| **143** | Matsumoto, Riki | 100 | 2022 | 932 | 9.3 | 0.68 | 36 |
| **144** | Kishima, Haruhiko | 104 | 2022 | 799 | 7.7 | 0.78 | 36 |
| **145** | Hongo, Kazuhiro | 122 | 2022 | 782 | 6.4 | 0.5 | 36 |
| **146** | Iihara, Koji | 166 | 2022 | 2025 | 12.2 | 0.91 | 35 |
| **147** | Yamagami, Hiroshi | 118 | 2022 | 1896 | 16.1 | 1.81 | 35 |
| **148** | Yang, Xinjian | 122 | 2022 | 1692 | 13.9 | 1.11 | 35 |
| **149** | Kang, Hyun-seung | 122 | 2022 | 1624 | 13.3 | 0.8 | 35 |
| **150** | Kohmura, Eiji | 101 | 2022 | 902 | 8.9 | 0.66 | 35 |
| **151** | Kalincik, Tomas | 119 | 2022 | 3238 | 27.2 | 2.07 | 34 |
| **152** | Fan, Dongsheng | 112 | 2022 | 1812 | 16.2 | 1.24 | 34 |
| **153** | Xu, Yuming | 108 | 2022 | 1531 | 14.2 | 1.31 | 34 |
| **154** | Yoshida, Kazunari | 149 | 2022 | 1453 | 9.8 | 0.68 | 34 |
| **155** | Taly, Arun B. | 109 | 2022 | 986 | 9 | 0.53 | 34 |
| **156** | Nakase, Hiroyuki | 137 | 2022 | 846 | 6.2 | 0.63 | 34 |
| **157** | Kim, Joon-tae | 121 | 2022 | 2439 | 20.2 | 1.53 | 33 |
| **158** | Zhang, Jianguo | 126 | 2022 | 1948 | 15.5 | 1.21 | 33 |
| **159** | Liu, Xinfeng | 108 | 2022 | 1883 | 17.4 | 1.58 | 33 |
| **160** | Zhu, Zezhang | 143 | 2022 | 1570 | 11 | 0.75 | 33 |
| **161** | Sinha, Sanjib | 139 | 2022 | 1453 | 10.5 | 0.56 | 33 |
| **162** | Kawamata, Takakazu | 200 | 2022 | 1418 | 7.1 | 0.68 | 33 |
| **163** | Suri, Ashish K. | 103 | 2022 | 1165 | 11.3 | 0.74 | 33 |
| **164** | Yamashita, Tohru | 110 | 2022 | 1060 | 9.6 | 0.72 | 33 |
| **165** | Behari, Sanjay S. | 130 | 2022 | 798 | 6.1 | 0.52 | 33 |
| **166** | Sakai, Nobuyuki | 181 | 2022 | 2033 | 11.2 | 1.26 | 32 |
| **167** | Lee, Chengchia | 132 | 2022 | 1988 | 15.1 | 1.41 | 32 |
| **168** | Kim, Chi-hwa | 104 | 2022 | 1779 | 17.1 | 1.06 | 32 |
| **169** | Koga, Masatoshi | 106 | 2022 | 1775 | 16.7 | 1.65 | 32 |
| **170** | Yamasaki, Ryo | 110 | 2022 | 1541 | 14 | 0.78 | 32 |
| **171** | Phi, Ji-hoon | 107 | 2022 | 1273 | 11.9 | 0.86 | 32 |
| **172** | Bhatia, Rohit S. | 103 | 2022 | 1214 | 11.8 | 0.8 | 32 |
| **173** | Huang, Qianghai | 136 | 2022 | 1470 | 10.8 | 0.82 | 31 |
| **174** | Garg, Ajay | 153 | 2022 | 1238 | 8.1 | 0.62 | 30 |
| **175** | Gulati, Sheffali K. | 110 | 2022 | 927 | 8.4 | 0.66 | 30 |
| **176** | Wiwanitit, V. A. | 111 | 2022 | 109 | 1 | 0.46 | 30 |
| **177** | Yu, Shengyuan | 143 | 2022 | 1885 | 13.2 | 0.84 | 29 |
| **178** | Yoshimoto, Koji | 100 | 2022 | 1498 | 15 | 0.83 | 29 |
| **179** | Zhang, Liwei | 117 | 2022 | 1339 | 11.4 | 0.75 | 29 |
| **180** | Ishikawa, Eiichi | 113 | 2022 | 837 | 7.4 | 0.5 | 29 |
| **181** | Wang, Shuo | 180 | 2022 | 1895 | 10.5 | 0.88 | 28 |
| **182** | Hong, Bo | 103 | 2022 | 1069 | 10.4 | 0.8 | 28 |
| **183** | Hirose, Yuichi | 101 | 2022 | 933 | 9.2 | 0.69 | 28 |
| **184** | Zhang, Junting | 140 | 2022 | 1820 | 13 | 0.78 | 27 |
| **185** | Kim, Han-joon | 148 | 2022 | 1691 | 11.4 | 0.93 | 27 |
| **186** | Kim, Hyo-jung | 116 | 2022 | 1383 | 11.9 | 0.98 | 27 |
| **187** | Miao, Zhongrong | 162 | 2022 | 1829 | 11.3 | 1.44 | 26 |
| **188** | Kale, Shashank Sharad | 155 | 2022 | 1459 | 9.4 | 0.64 | 26 |
| **189** | Saini, Jitender S. | 156 | 2022 | 1455 | 9.3 | 0.71 | 26 |
| **190** | Ando, Kei | 123 | 2022 | 1102 | 9 | 0.91 | 26 |
| **191** | Devi, Bhagavathula Indira | 136 | 2022 | 1080 | 7.9 | 0.62 | 26 |
| **192** | Wang, Zhaoxia | 122 | 2022 | 735 | 6 | 0.7 | 26 |
| **193** | Srivastava, Achal Kumar | 120 | 2022 | 718 | 6 | 0.6 | 26 |
| **194** | Cao, Bei | 120 | 2022 | 1756 | 14.6 | 0.89 | 25 |
| **195** | Wu, Zhen | 140 | 2022 | 1766 | 12.6 | 0.8 | 24 |
| **196** | Wei, Qianqian | 124 | 2022 | 1444 | 11.6 | 0.87 | 24 |
| **197** | Ma, Ning | 102 | 2022 | 1181 | 11.6 | 1.39 | 24 |
| **198** | Yang, Pengfei | 106 | 2022 | 1105 | 10.4 | 1.16 | 24 |
| **199** | Peng, Bin | 117 | 2022 | 888 | 7.6 | 0.67 | 24 |
| **200** | Goyal, Vinay | 102 | 2022 | 775 | 7.6 | 0.77 | 24 |
| **201** | Zhang, Dong | 126 | 2022 | 1608 | 12.8 | 0.99 | 23 |
| **202** | Wang, Rong | 100 | 2022 | 1524 | 15.2 | 1.11 | 23 |
| **203** | You, Chao | 192 | 2022 | 1383 | 7.2 | 0.57 | 23 |
| **204** | Ou, Ruwei | 104 | 2022 | 1201 | 11.5 | 0.89 | 23 |
| **205** | Zhang, Hongqi | 132 | 2022 | 993 | 7.5 | 0.71 | 23 |
| **206** | Shah, Abhidha Harshad | 114 | 2022 | 973 | 8.5 | 0.89 | 23 |
| **207** | Lal, Vivek | 122 | 2022 | 607 | 5 | 0.54 | 22 |
| **208** | Horiuchi, Tetsuyoshi | 111 | 2022 | 548 | 4.9 | 0.57 | 22 |
| **209** | Agrawal, Amit R. | 101 | 2022 | 1198 | 11.9 | 1.23 | 21 |
| **210** | Cao, Yong | 115 | 2022 | 1169 | 10.2 | 0.83 | 21 |
| **211** | Chandra, Poodipedi Sarat | 205 | 2022 | 1127 | 5.5 | 0.56 | 21 |
| **212** | Yadav, Ravi | 141 | 2022 | 1111 | 7.9 | 0.76 | 21 |
| **213** | Inoue, Tooru | 103 | 2022 | 946 | 9.2 | 0.76 | 21 |
| **214** | Salunke, Pravin Shashikant | 183 | 2022 | 933 | 5.1 | 0.56 | 20 |
| **215** | Singh, Manmohan | 106 | 2022 | 765 | 7.2 | 0.57 | 20 |
| **216** | Nakagawa, Ichiro | 115 | 2022 | 618 | 5.4 | 0.44 | 20 |
| **217** | Pandey, Sanjay | 124 | 2022 | 991 | 8 | 0.75 | 18 |
| **218** | Shukla, Dhaval Premchand | 158 | 2022 | 920 | 5.8 | 0.45 | 18 |
| **219** | Bhat, Dhananjaya Ishwar | 106 | 2022 | 790 | 7.5 | 0.47 | 18 |
| **220** | Jiao, Liqun | 117 | 2022 | 749 | 6.4 | 0.74 | 18 |
| **221** | Tripathi, Manjul | 150 | 2022 | 629 | 4.2 | 0.77 | 14 |
| **222** | Garg, Kanwaljeet | 175 | 2022 | 785 | 4.5 | 0.6 | 13 |
| **223** | Das, Kuntal Kanti | 113 | 2022 | 354 | 3.1 | 0.37 | 11 |
| **224** | Sharawat, Indar Kumar | 104 | 2022 | 211 | 2 | 0.46 | 11 |
| **225** | Yagi, Mitsuru | 99 | 2022 | 1422 | 14.4 | 0.99 | 29 |
| **226** | Lv, Xianli | 99 | 2022 | 1094 | 11.1 | 0.64 | 30 |
| **227** | Nagoshi, Narihito | 99 | 2022 | 1379 | 13.9 | 1.15 | 36 |
| **228** | Zhao, Yuanli | 99 | 2022 | 1349 | 13.6 | 1.01 | 27 |
| **229** | Hankey, Graeme J. | 99 | 2022 | 11236 | 113.5 | 8.75 | 139 |
| **230** | Jackson, Graeme D. | 99 | 2022 | 2603 | 26.3 | 1.87 | 68 |
| **231** | Liu, Chunfeng | 98 | 2022 | 1185 | 12.1 | 0.95 | 44 |
| **232** | Bhidayasiri, Roongroj | 98 | 2022 | 1605 | 16.4 | 1.34 | 31 |
| **233** | Kim, Young-dae | 98 | 2022 | 2189 | 22.3 | 1.66 | 40 |
| **234** | Mobbs, Ralph Jasper | 98 | 2022 | 2940 | 30 | 1.51 | 43 |
| **235** | Wakabayashi, Toshishiko | 98 | 2019 | 1230 | 12.6 | 0.65 | 46 |
| **236** | Kang, Dong-wha | 98 | 2022 | 2256 | 23 | 1.28 | 54 |
| **237** | Ahuja, Chirag Kamal | 97 | 2022 | 422 | 4.4 | 0.44 | 17 |
| **238** | Hyun, Seung-jae | 97 | 2022 | 1960 | 20.2 | 1.24 | 31 |
| **239** | Singh, Pankaj Kumar | 97 | 2022 | 559 | 5.8 | 0.49 | 20 |
| **240** | Kobayashi, Kazuyoshi | 97 | 2022 | 868 | 8.9 | 0.93 | 22 |
| **241** | Matsumaru, Yuji | 97 | 2022 | 842 | 8.7 | 1.41 | 24 |
| **242** | Malhotra, Hardeep Singh | 97 | 2022 | 865 | 8.9 | 0.52 | 24 |
| **243** | Prasad, Kameshwar Nagendra | 97 | 2022 | 1219 | 12.6 | 0.89 | 42 |
| **244** | Koda, Masao | 96 | 2022 | 650 | 6.8 | 0.53 | 39 |
| **245** | Yu, Jintai | 96 | 2022 | 4353 | 45.3 | 2.61 | 74 |
| **246** | Mikami, Takeshi | 96 | 2022 | 911 | 9.5 | 0.67 | 20 |
| **247** | Srivastava, Arun Kumar | 96 | 2022 | 484 | 5 | 0.45 | 16 |
| **248** | Guan, Hongzhi | 96 | 2022 | 783 | 8.2 | 0.59 | 23 |
| **249** | Qiu, Wei | 96 | 2022 | 1370 | 14.3 | 1.08 | 31 |
| **250** | Saito, Ryuta | 96 | 2022 | 943 | 9.8 | 0.75 | 29 |
| **251** | Minematsu, Kazuo | 96 | 2022 | 2170 | 22.6 | 1.49 | 53 |
| **252** | Wang, Yuping | 96 | 2022 | 1149 | 12 | 0.89 | 38 |
| **253** | Arai, Hajime | 95 | 2022 | 1286 | 13.5 | 1.14 | 55 |
| **254** | Zhao, Bi | 95 | 2022 | 1218 | 12.8 | 0.85 | 24 |
| **255** | Murayama, Shigeo | 95 | 2022 | 2534 | 26.7 | 2.19 | 65 |
| **256** | Lee, Ji-yeoun | 95 | 2022 | 1048 | 11 | 0.72 | 29 |
| **257** | van der Mei, Ingrid A.F. | 95 | 2022 | 2441 | 25.7 | 2.12 | 42 |
| **258** | Chung, Sun-ju | 95 | 2022 | 1400 | 14.7 | 1.09 | 33 |
| **259** | Murayama, Yuichi | 95 | 2022 | 1475 | 15.5 | 1.24 | 45 |
| **260** | Watanabe, Hirohisa | 95 | 2022 | 1717 | 18.1 | 1.04 | 45 |
| **261** | Martins, Ralph Nigel | 94 | 2022 | 5945 | 63.2 | 3.81 | 91 |
| **262** | Fujii, Yukihiko | 94 | 2022 | 1090 | 11.6 | 0.78 | 37 |
| **263** | Mao, Ying | 94 | 2022 | 2169 | 23.1 | 1.29 | 54 |
| **264** | Okada, Yoshikazu | 94 | 2022 | 1121 | 11.9 | 0.59 | 31 |
| **265** | Xu, Yi | 94 | 2022 | 1030 | 11 | 0.83 | 25 |
| **266** | Kuwabara, Satoshi | 94 | 2022 | 848 | 9 | 1.33 | 17 |
| **267** | Kitazono, Takanari | 94 | 2022 | 1534 | 16.3 | 1.14 | 53 |
| **268** | Fujihara, Kazuo | 94 | 2020 | 10664 | 113.4 | 4.44 | 65 |
| **269** | Mori, Masahiro | 94 | 2022 | 1551 | 16.5 | 1.44 | 49 |
| **270** | Park, Jong-moo | 94 | 2022 | 1809 | 19.2 | 1.21 | 33 |
| **271** | Kong, Doo-sik | 93 | 2022 | 1179 | 12.7 | 1.07 | 38 |
| **272** | Iguchi, Yasuyuki | 93 | 2022 | 591 | 6.4 | 0.66 | 34 |
| **273** | Endo, Hidenori | 93 | 2022 | 1021 | 11 | 0.92 | 27 |
| **274** | Nagappa, Madhu | 93 | 2022 | 708 | 7.6 | 0.56 | 16 |
| **275** | Okumura, Akihisa | 93 | 2022 | 884 | 9.5 | 0.68 | 38 |
| **276** | Tanaka, Fumiaki | 93 | 2022 | 1461 | 15.7 | 1.14 | 56 |
| **277** | Gao, Feng | 93 | 2022 | 1160 | 12.5 | 1.51 | 22 |
| **278** | Mo, Dapeng | 92 | 2022 | 991 | 10.8 | 1.38 | 20 |
| **279** | Suri, Vaishali S. | 92 | 2022 | 904 | 9.8 | 0.65 | 27 |
| **280** | Yeo, Leonard Leong Litt | 92 | 2022 | 1203 | 13.1 | 1.52 | 25 |
| **281** | Niizuma, Kuniyasu | 92 | 2022 | 1492 | 16.2 | 1.08 | 36 |
| **282** | Song, Wei | 92 | 2022 | 1559 | 16.9 | 0.9 | 26 |
| **283** | Cho, Won-sang | 92 | 2022 | 1608 | 17.5 | 1.02 | 22 |
| **284** | Anderson, Vicki A. | 92 | 2022 | 1914 | 20.8 | 1.07 | 78 |
| **285** | Hasegawa, Yasuhiro | 92 | 2022 | 1229 | 13.4 | 0.76 | 36 |
| **286** | Chan, Piu | 92 | 2022 | 6483 | 70.5 | 2.18 | 63 |
| **287** | Yu, Kyungho | 92 | 2022 | 1899 | 20.6 | 1.34 | 35 |
| **288** | Sylaja, Padmavathyamma Narayanapillai | 91 | 2022 | 4097 | 45 | 3.94 | 33 |
| **289** | Imamura, Hirotoshi | 91 | 2022 | 700 | 7.7 | 0.69 | 14 |
| **290** | Ohata, Kenji | 91 | 2022 | 767 | 8.4 | 0.63 | 29 |
| **291** | Kim, Joong-seok | 91 | 2022 | 756 | 8.3 | 0.86 | 29 |
| **292** | Suzuki, Keisuke | 91 | 2022 | 912 | 10 | 0.8 | 34 |
| **293** | Rajshekhar, Vedantam | 91 | 2022 | 806 | 8.9 | 0.58 | 39 |
| **294** | Dhandapani, Sivashanmugam | 91 | 2022 | 911 | 10 | 0.83 | 20 |
| **295** | Hishikawa, Nozomi | 91 | 2021 | 968 | 10.6 | 0.79 | 27 |
| **296** | Thijs, Vincent N.S. | 91 | 2022 | 1961 | 21.5 | 1.85 | 72 |
| **297** | Sasaki, Masayuki | 91 | 2022 | 931 | 10.2 | 0.71 | 40 |
| **298** | Choi, Jeong-yoon | 90 | 2022 | 985 | 10.9 | 0.93 | 20 |
| **299** | Nakashima, Hiroaki | 90 | 2022 | 1660 | 18.4 | 1.31 | 33 |
| **300** | Toda, Tatsushi | 90 | 2022 | 1125 | 12.5 | 1.16 | 59 |
| **301** | Ha, Yoon | 90 | 2022 | 1457 | 16.2 | 1.07 | 36 |
| **302** | Liu, Mingsheng | 90 | 2022 | 853 | 9.5 | 0.78 | 21 |
| **303** | Leung, Thomas Wai Hong | 89 | 2022 | 2824 | 31.7 | 2.5 | 35 |
| **304** | Kikuchi, Takayuki | 89 | 2022 | 1111 | 12.5 | 0.67 | 22 |
| **305** | Hao, Dingjun | 89 | 2022 | 916 | 10.3 | 0.63 | 35 |
| **306** | Lou, Min | 89 | 2022 | 1554 | 17.5 | 1.61 | 32 |
| **307** | Lu, Zhengqi | 89 | 2022 | 1341 | 15.1 | 0.86 | 32 |
| **308** | Kim, Byungmoon | 88 | 2022 | 1750 | 19.9 | 1.3 | 41 |
| **309** | Mahale, Rohan R. | 88 | 2022 | 299 | 3.4 | 0.46 | 10 |
| **310** | Arima, Hisatomi | 88 | 2022 | 2235 | 25.4 | 1.55 | 71 |
| **311** | Wanibuchi, Masahiko | 88 | 2022 | 707 | 8 | 0.61 | 21 |
| **312** | Mukasa, Akitake | 88 | 2022 | 1455 | 16.5 | 0.96 | 35 |
| **313** | Takahashi, Jun C. | 88 | 2022 | 1942 | 22.1 | 1.76 | 29 |
| **314** | Tan, Louis Chew Seng | 88 | 2022 | 2195 | 24.9 | 1.65 | 42 |
| **315** | Jaiswal, Awadhesh Kumar | 87 | 2022 | 450 | 5.2 | 0.43 | 17 |
| **316** | Takashima, Hiroshi | 87 | 2022 | 886 | 10.2 | 0.62 | 37 |
| **317** | Yuki, Nobuhiro | 87 | 2021 | 2478 | 28.5 | 1.15 | 71 |
| **318** | Meng, Xia | 87 | 2022 | 1437 | 16.5 | 1.32 | 26 |
| **319** | Yamamoto, Tetsuya | 87 | 2022 | 966 | 11.1 | 0.67 | 29 |
| **320** | Deora, Harsh | 87 | 2022 | 397 | 4.6 | 0.71 | 11 |
| **321** | Liu, Jianmin | 87 | 2022 | 842 | 9.7 | 0.75 | 22 |
| **322** | Fitzgerald, Paul B. | 87 | 2022 | 4922 | 56.6 | 2.58 | 82 |
| **323** | Mizuno, Toshiki | 87 | 2022 | 934 | 10.7 | 0.79 | 33 |
| **324** | Chae, Jong-hee | 86 | 2022 | 879 | 10.2 | 0.95 | 35 |
| **325** | Tanikawa, Rokuya | 86 | 2022 | 804 | 9.3 | 0.62 | 16 |
| **326** | Furuya, Takeo | 86 | 2022 | 707 | 8.2 | 0.8 | 26 |
| **327** | Lindley, Richard Iain | 86 | 2022 | 5605 | 65.2 | 2.18 | 54 |
| **328** | Bivard, Andrew | 86 | 2022 | 2020 | 23.5 | 1.75 | 32 |
| **329** | Kim, Dae-hyun | 86 | 2022 | 1534 | 17.8 | 1.2 | 25 |
| **330** | Cadilhac, Dominique A. | 86 | 2022 | 1232 | 14.3 | 1.31 | 38 |
| **331** | Hong, Zhen | 86 | 2022 | 1501 | 17.5 | 0.95 | 47 |
| **332** | Toda, Masahiro | 86 | 2022 | 799 | 9.3 | 0.75 | 31 |
| **333** | Hegde, Alangar Sathya | 86 | 2022 | 384 | 4.5 | 0.28 | 15 |
| **334** | Arivazhagan, Arimappamagan | 85 | 2022 | 600 | 7.1 | 0.5 | 24 |
| **335** | Kim, Dong-eog | 85 | 2022 | 1549 | 18.2 | 1.22 | 38 |
| **336** | Kanda, Takashi | 85 | 2022 | 1116 | 13.1 | 0.79 | 41 |
| **337** | Matsumoto, Naomichi | 85 | 2022 | 801 | 9.4 | 0.96 | 29 |
| **338** | Muragaki, Yoshihiro | 85 | 2022 | 1605 | 18.9 | 1.11 | 36 |
| **339** | Ren, Haitao | 85 | 2022 | 900 | 10.6 | 0.7 | 23 |
| **340** | Okada, Yashushi | 85 | 2022 | 2012 | 23.7 | 1.33 | 42 |
| **341** | Zhu, Yicheng | 84 | 2022 | 1244 | 14.8 | 1.08 | 33 |
| **342** | Srinivas, Dwarakanath | 84 | 2022 | 387 | 4.6 | 0.48 | 15 |
| **343** | Ishiguro, Naoki | 84 | 2021 | 1237 | 14.7 | 0.93 | 65 |
| **344** | Cho, Young-dae | 84 | 2022 | 863 | 10.3 | 0.74 | 21 |
| **345** | Ma, Lu | 84 | 2022 | 681 | 8.1 | 0.72 | 17 |
| **346** | Liu, Jianmin | 84 | 2022 | 904 | 10.8 | 1.49 | 24 |
| **347** | Song, Tae-jin | 84 | 2022 | 1555 | 18.5 | 1.47 | 29 |
| **348** | Jang, Sungho | 84 | 2020 | 1113 | 13.3 | 0.94 | 45 |
| **349** | Suzuki, Satoshi Satoshi | 84 | 2020 | 1431 | 17 | 0.91 | 43 |
| **350** | Kim, Heung-dong | 84 | 2022 | 1213 | 14.4 | 0.9 | 37 |
| **351** | Doddamani, Ramesh Sharanappa | 84 | 2022 | 262 | 3.1 | 0.53 | 8 |
| **352** | Park, Chul Kee | 84 | 2022 | 1314 | 15.6 | 1.03 | 47 |
| **353** | Li, Zixiao | 84 | 2022 | 1214 | 14.5 | 1.69 | 22 |
| **354** | Zetterberg, Henrik H. | 84 | 2022 | 823 | 9.8 | 4.19 | 136 |
| **355** | Poon, Wai sang | 84 | 2022 | 1411 | 16.8 | 0.95 | 54 |
| **356** | Inoue, Yushi | 84 | 2022 | 891 | 10.6 | 0.65 | 37 |
| **357** | Okanishi, T. | 83 | 2022 | 353 | 4.3 | 0.46 | 17 |
| **358** | Fujimoto, Ayataka | 83 | 2022 | 291 | 3.5 | 0.36 | 13 |
| **359** | Wang, Kai | 83 | 2022 | 1065 | 12.8 | 1.21 | 39 |
| **360** | Li, Hao | 83 | 2022 | 887 | 10.7 | 0.74 | 20 |
| **361** | Lee, Jung-il | 83 | 2022 | 1041 | 12.5 | 0.99 | 56 |
| **362** | Fuh, Jongling | 83 | 2022 | 1830 | 22 | 1.24 | 65 |
| **363** | Ichimura, Koichi | 83 | 2022 | 2301 | 27.7 | 1.94 | 53 |
| **364** | Nishizawa, Masatoyo | 83 | 2021 | 1724 | 20.8 | 1.25 | 61 |
| **365** | Endo, Toshiki | 83 | 2022 | 709 | 8.5 | 0.72 | 23 |
| **366** | Sun, Bomin | 83 | 2022 | 1064 | 12.8 | 1.26 | 27 |
| **367** | Komori, Takashi | 82 | 2022 | 2009 | 24.5 | 2.43 | 34 |
| **368** | Wu, Bo | 82 | 2022 | 1936 | 23.6 | 1.25 | 32 |
| **369** | Yan, Xiaoling | 82 | 2020 | 47 | 0.6 | 0.04 | 12 |
| **370** | Kim, Kijeong | 82 | 2022 | 1599 | 19.5 | 1.26 | 30 |
| **371** | Jiang, Tao | 82 | 2022 | 2257 | 27.5 | 1.3 | 67 |
| **372** | Takahashi, Hitoshi | 82 | 2019 | 2367 | 28.9 | 1.55 | 84 |
| **373** | Kim, Bum-joon | 82 | 2022 | 812 | 9.9 | 0.83 | 20 |
| **374** | Sakakibara, Ryuuji | 82 | 2022 | 1186 | 14.5 | 0.84 | 47 |
| **375** | Narita, Yoshitaka | 82 | 2022 | 2177 | 26.5 | 1.63 | 44 |
| **376** | Maruyama, Hirofumi | 82 | 2022 | 913 | 11.1 | 1.07 | 34 |
| **377** | Kitagawa, Kazuo | 82 | 2022 | 1334 | 16.3 | 1.16 | 53 |
| **378** | Uzawa, Akiyuki | 81 | 2022 | 995 | 12.3 | 0.95 | 28 |
| **379** | Seo, Woo-keun | 81 | 2022 | 1284 | 15.9 | 1.04 | 28 |
| **380** | Aoki, Junya | 81 | 2022 | 905 | 11.2 | 0.7 | 30 |
| **381** | Kang, Hoon Chul | 81 | 2022 | 1456 | 18 | 1.21 | 40 |
| **382** | Onodera, Osamu | 81 | 2022 | 1367 | 16.9 | 1.12 | 55 |
| **383** | Ohta, Yasuyuki | 81 | 2022 | 740 | 9.1 | 0.73 | 31 |
| **384** | Lee, Jun | 81 | 2022 | 1650 | 20.4 | 1.37 | 29 |
| **385** | Nam, Do-hyun | 81 | 2022 | 1407 | 17.4 | 1.12 | 65 |
| **386** | Hatano, Taku | 81 | 2022 | 1309 | 16.2 | 1.49 | 31 |
| **387** | Maehara, Taketoshi | 81 | 2022 | 904 | 11.2 | 0.96 | 27 |
| **388** | Lee, Juneyoung | 80 | 2021 | 1749 | 21.9 | 1.23 | 46 |
| **389** | Goyal, Manoj Kumar | 80 | 2022 | 443 | 5.5 | 0.37 | 15 |
| **390** | Suzuki, Michiyasu | 80 | 2022 | 930 | 11.6 | 0.7 | 17 |
| **391** | Kim, Sehoon | 80 | 2022 | 1442 | 18 | 1.61 | 45 |
| **392** | Maruff, Paul T. | 80 | 2022 | 4756 | 59.5 | 3.08 | 84 |
| **393** | Murai, Hiroyuki | 80 | 2022 | 2583 | 32.3 | 1.94 | 33 |
| **394** | Cha, Jae-kwan | 80 | 2022 | 1446 | 18.1 | 1.11 | 26 |
| **395** | Boyd, Roslyn N. | 80 | 2022 | 2205 | 27.6 | 1.59 | 62 |
| **396** | Simpson-Yap, Steve L. | 79 | 2022 | 1409 | 17.8 | 1.51 | 29 |
| **397** | Mehrotra, Anant Kumar | 79 | 2022 | 391 | 4.9 | 0.4 | 14 |
| **398** | Kim, Suhyun | 79 | 2022 | 2390 | 30.3 | 2.3 | 32 |
| **399** | Gupta, Sunil Kumar | 79 | 2022 | 636 | 8.1 | 0.76 | 31 |
| **400** | Kim, Hojin | 79 | 2022 | 2583 | 32.7 | 1.96 | 34 |
| **401** | Kleinig, Timothy J. | 79 | 2022 | 1864 | 23.6 | 3.12 | 36 |
| **402** | Taira, Takaomi | 79 | 2022 | 1077 | 13.6 | 1.35 | 24 |
| **403** | Sato, Noriko O. | 79 | 2022 | 895 | 11.3 | 0.82 | 33 |
| **404** | Bharath, Rose Dawn | 78 | 2022 | 789 | 10.1 | 0.6 | 22 |
| **405** | Aoki, Shigeki | 78 | 2022 | 1525 | 19.6 | 1.84 | 58 |
| **406** | Deng, Xiaofeng | 78 | 2022 | 961 | 12.3 | 0.8 | 17 |
| **407** | Guo, Jifeng | 78 | 2022 | 925 | 11.9 | 1.06 | 32 |
| **408** | Nakamura, Hiroaki | 78 | 2022 | 899 | 11.5 | 0.87 | 36 |
| **409** | Thrift, Amanda G. | 78 | 2022 | 7989 | 102.4 | 7.86 | 72 |
| **410** | Mahapatra, Ashok Kumar | 78 | 2021 | 860 | 11 | 0.5 | 40 |
| **411** | Hwang, Yangha | 78 | 2022 | 1639 | 21 | 1.96 | 25 |
| **412** | Yamazaki, Masashi | 78 | 2022 | 716 | 9.2 | 0.67 | 48 |
| **413** | Song, Yueming | 78 | 2022 | 709 | 9.1 | 0.67 | 27 |
| **414** | Singh, Mamta Bhushan | 78 | 2022 | 491 | 6.3 | 0.47 | 20 |
| **415** | Lee, Jong Min | 77 | 2022 | 2061 | 26.8 | 1.13 | 52 |
| **416** | Liu, Xinfeng | 77 | 2022 | 1539 | 20 | 1.3 | 43 |
| **417** | Oh, Chang-wan | 77 | 2022 | 1568 | 20.4 | 1 | 38 |
| **418** | Li, Da | 77 | 2021 | 1107 | 14.4 | 0.9 | 18 |
| **419** | Matsumoto, Naomichi | 77 | 2022 | 2267 | 29.4 | 1.88 | 66 |
| **420** | Thomas, Bejoy | 77 | 2022 | 888 | 11.5 | 0.68 | 26 |
| **421** | Hodges, Paul William | 77 | 2022 | 1717 | 22.3 | 1.73 | 97 |
| **422** | Phan, Thanh G. | 77 | 2022 | 1577 | 20.5 | 1.54 | 53 |
| **423** | Sonoda, Yukihiko | 77 | 2022 | 1056 | 13.7 | 0.83 | 29 |
| **424** | Dong, Qiang | 77 | 2022 | 2583 | 33.5 | 2.33 | 35 |
| **425** | Yokota, Takanori | 77 | 2022 | 852 | 11.1 | 1.05 | 43 |
| **426** | Mori, Kentaro | 77 | 2022 | 481 | 6.2 | 0.44 | 27 |
| **427** | Kim, Sangyun | 77 | 2022 | 1038 | 13.5 | 1.08 | 34 |
| **428** | Kashiwazaki, Daina | 77 | 2022 | 746 | 9.7 | 0.77 | 18 |
| **429** | Somanna, Sampath | 76 | 2022 | 477 | 6.3 | 0.35 | 21 |
| **430** | Yoon, Byoungwoo | 76 | 2022 | 2341 | 30.8 | 1.46 | 53 |
| **431** | Wang, Shuang | 76 | 2022 | 984 | 12.9 | 0.82 | 30 |
| **432** | Jung, Ki-young | 76 | 2022 | 1448 | 19.1 | 0.96 | 32 |
| **433** | Satyarthee, Guru Dutta | 76 | 2022 | 300 | 3.9 | 0.56 | 13 |
| **434** | Wang, Rengzhi | 76 | 2022 | 775 | 10.2 | 0.68 | 34 |
| **435** | Nakatomi, Hirofumi | 76 | 2022 | 840 | 11.1 | 0.82 | 21 |
| **436** | Motoyama, Yasushi | 76 | 2022 | 475 | 6.3 | 0.52 | 15 |
| **437** | Arita, Kazunori | 76 | 2022 | 756 | 9.9 | 0.5 | 39 |
| **438** | Iwaki, Toru | 76 | 2022 | 1332 | 17.5 | 0.95 | 51 |
| **439** | Shen, Lu | 76 | 2022 | 1489 | 19.6 | 1.7 | 35 |
| **440** | Yamada, Masahito | 76 | 2022 | 2227 | 29.3 | 1.67 | 60 |
| **441** | Chu, Min-Kyung | 75 | 2022 | 1075 | 14.3 | 1.12 | 24 |
| **442** | Hao, Shuyu | 75 | 2021 | 1007 | 13.4 | 0.72 | 23 |
| **443** | Jian, Feng Zeng | 75 | 2022 | 255 | 3.4 | 0.58 | 12 |
| **444** | Vishnu, Venugopalan Yamuna | 75 | 2022 | 177 | 2.4 | 0.39 | 9 |
| **445** | Zhang, Wei | 75 | 2022 | 494 | 6.6 | 0.64 | 23 |
| **446** | Lee, Sang Ahm | 75 | 2022 | 649 | 8.7 | 0.73 | 27 |
| **447** | Sonoo, Masahiro | 75 | 2022 | 818 | 10.9 | 0.61 | 27 |
| **448** | Shiokawa, Yoshiaki | 75 | 2022 | 1311 | 17.5 | 1.47 | 34 |
| **449** | Sarkar, Chitra | 75 | 2022 | 1631 | 21.7 | 1.64 | 44 |
| **450** | Koike, Haruki | 75 | 2022 | 1645 | 21.9 | 1.49 | 45 |
| **451** | Kim, Kijoong | 75 | 2022 | 713 | 9.5 | 0.7 | 30 |
| **452** | Chung, Joonho | 74 | 2022 | 741 | 10 | 0.86 | 19 |
| **453** | Wu, Jau-Ching | 74 | 2022 | 942 | 12.7 | 1.3 | 31 |
| **454** | Sakamoto, Yuki | 74 | 2021 | 738 | 10 | 0.57 | 21 |
| **455** | Sohn, Seung Il I. | 74 | 2022 | 1720 | 23.2 | 2.53 | 32 |
| **456** | Youn, Jinyoung | 74 | 2022 | 957 | 12.9 | 0.78 | 22 |
| **457** | Park, Kyung-min | 74 | 2022 | 637 | 8.6 | 1.13 | 15 |
| **458** | Wang, Liang | 74 | 2022 | 751 | 10.1 | 0.77 | 19 |
| **459** | Ohtori, Seiji | 74 | 2022 | 1356 | 18.3 | 1.04 | 49 |
| **460** | Lee, Mi-ji | 74 | 2022 | 1340 | 18.1 | 1.24 | 27 |
| **461** | Akiyama, Yukinori | 74 | 2022 | 659 | 8.9 | 0.68 | 19 |
| **462** | Matsuda, Hiroslri | 74 | 2022 | 1415 | 19.1 | 1.09 | 58 |
| **463** | Hou, Yanbing | 74 | 2022 | 570 | 7.7 | 0.91 | 15 |
| **464** | Lee, Hye-sun | 74 | 2022 | 1492 | 20.2 | 1.48 | 39 |
| **465** | Kim, Jin-sung Luke | 74 | 2022 | 1131 | 15.3 | 1.25 | 29 |
| **466** | Saini, Lokesh | 74 | 2022 | 184 | 2.5 | 0.78 | 9 |
| **467** | Delatycki, Martin B. | 74 | 2022 | 1626 | 22 | 1.47 | 58 |
| **468** | Halmagyi, Michael G. | 74 | 2022 | 1712 | 23.1 | 1.57 | 55 |
| **469** | Yuan, Yun | 74 | 2022 | 479 | 6.5 | 0.44 | 26 |
| **470** | Choi, Jay-chol | 74 | 2022 | 1363 | 18.4 | 1.2 | 24 |
| **471** | Mohindra, Sandeep | 74 | 2022 | 335 | 4.5 | 0.51 | 18 |
| **472** | Yoshii, Toshitaka | 73 | 2022 | 890 | 12.2 | 1.19 | 28 |
| **473** | Kanamori, Masayuki | 73 | 2022 | 1123 | 15.4 | 0.92 | 29 |
| **474** | Wang, Chunxue | 73 | 2022 | 2293 | 31.4 | 1.25 | 39 |
| **475** | Kim, Hyeun-sung | 73 | 2022 | 831 | 11.4 | 1.2 | 23 |
| **476** | Rajasekaran, S. B. | 73 | 2022 | 1632 | 22.4 | 1.51 | 40 |
| **477** | Cho, Jin-whan | 73 | 2022 | 949 | 13 | 0.71 | 24 |
| **478** | Takeuchi, Satoru | 73 | 2021 | 670 | 9.2 | 0.53 | 19 |
| **479** | Paek, Sun-ha | 73 | 2022 | 1413 | 19.4 | 0.94 | 49 |
| **480** | Lee, Jee-young | 73 | 2022 | 1203 | 16.5 | 1.18 | 31 |
| **481** | Choi, Kangho | 73 | 2022 | 935 | 12.8 | 1.12 | 23 |
| **482** | Liu, Zhen | 73 | 2022 | 941 | 12.9 | 0.84 | 28 |
| **483** | Mizusawa, Hidehiro | 73 | 2022 | 1578 | 21.6 | 1.38 | 64 |
| **484** | Morioka, Takato | 73 | 2022 | 541 | 7.4 | 0.68 | 30 |
| **485** | Yu, Xinguang | 73 | 2022 | 954 | 13.1 | 0.95 | 22 |
| **486** | Zhang, Yan | 73 | 2022 | 982 | 13.5 | 1.09 | 20 |
| **487** | Nalini, Atchayaram | 73 | 2022 | 601 | 8.2 | 0.69 | 28 |
| **488** | Misu, Tatsurou | 73 | 2022 | 3476 | 47.6 | 3.71 | 51 |
| **489** | Khandelwal, Niranjan K. | 73 | 2022 | 862 | 11.8 | 0.79 | 43 |
| **490** | Misawa, Sonoko | 73 | 2022 | 1124 | 15.4 | 1.05 | 44 |
| **491** | Mizoguchi, Masahiro | 73 | 2022 | 1351 | 18.5 | 1 | 29 |
| **492** | MacKay, Mark T. | 73 | 2022 | 7513 | 102.9 | 6.05 | 56 |
| **493** | Yan, Chuanzhu | 73 | 2022 | 452 | 6.2 | 0.71 | 23 |
| **494** | Yoshida, Kazumichi | 73 | 2022 | 783 | 10.7 | 0.74 | 22 |
| **495** | Vyas, Sameer | 72 | 2022 | 231 | 3.2 | 0.52 | 12 |
| **496** | Moseley, Lorimer Lorimer | 72 | 2022 | 3224 | 44.8 | 1.92 | 77 |
| **497** | Drummond, Katharine Jann | 72 | 2022 | 977 | 13.6 | 1.14 | 32 |
| **498** | Grunstein, Ronald R. | 72 | 2022 | 2104 | 29.2 | 1.48 | 64 |
| **499** | Morita, Tatsuya | 72 | 2022 | 1177 | 16.3 | 1.28 | 59 |
| **500** | Jiang, Chuhan | 72 | 2022 | 730 | 10.1 | 0.71 | 27 |

**Table 4:**	The list of top 100 researchers with scholarly output, most recent publications, total citations, citations per publications, field-weighted citation impact and h-index.

| **S#** | **CiteScore quartile** | **Overall** | **2013** | **2014** | **2015** | **2016** | **2017** | **2018** | **2019** | **2020** | **2021** | **2022** |
| --- | --- | --- | --- | --- | --- | --- | --- | --- | --- | --- | --- | --- |
| **1** | Q1 (top 25%) | 37407 | 3137 | 3265 | 3359 | 3259 | 3264 | 3661 | 4205 | 3996 | 4514 | 4747 |
| **2** | Q2 (top 26% - 50%) | 38752 | 2748 | 2568 | 2804 | 3456 | 3888 | 3402 | 3128 | 4163 | 5632 | 6963 |
| **3** | Q3 (top 51% - 75%) | 33218 | 2370 | 2921 | 2705 | 2616 | 3015 | 3883 | 4809 | 4295 | 3583 | 3021 |
| **4** | Q4 (top 76% - 100%) | 15364 | 1362 | 1262 | 1294 | 1338 | 1168 | 1180 | 1217 | 2016 | 2490 | 2037 |
| **5** | Total | 124741 | 9617 | 10016 | 10162 | 10669 | 11335 | 12126 | 13359 | 14470 | 16219 | 16768 |

**Table 5:**	The publications distribution in four quartile group (Q1-Q4).

| **S#** | **Scopus Source** | **Publications** | **Citations** | **Citations per Publication** | **Source-Normalized Impact per Paper (SNIP)** | **CiteScore 2022** | **SCImago Journal Rank (SJR)** |
| --- | --- | --- | --- | --- | --- | --- | --- |
| **1** | World Neurosurgery | 6439 | 48138 | 7.5 | 1.032 | 3.9 | 0.591 |
| **2** | Frontiers in Neurology | 5011 | 37694 | 7.5 | 1.151 | 4.8 | 0.978 |
| **3** | Neurology India | 3462 | 10238 | 3 | 0.764 | 1.6 | 0.448 |
| **4** | Journal of Clinical Neuroscience | 2707 | 24865 | 9.2 | 0.836 | 3.9 | 0.54 |
| **5** | Journal of Stroke and Cerebrovascular Diseases | 2636 | 26240 | 10 | 0.893 | 4.3 | 0.713 |
| **6** | Brain Research | 2092 | 37302 | 17.8 | 0.766 | 6.6 | 0.854 |
| **7** | Journal of the Neurological Sciences | 2015 | 25653 | 12.7 | 1.128 | 6.5 | 0.983 |
| **8** | Spine | 1975 | 35203 | 17.8 | 1.804 | 5.7 | 1.197 |
| **9** | Stroke | 1931 | 58918 | 30.5 | 2.734 | 12.9 | 2.746 |
| **10** | Chinese Journal of Contemporary Neurology and Neurosurgery | 1861 | 1055 | 0.6 | 0.084 | 0.3 | 0.112 |
| **11** | Neurology | 1685 | 55098 | 32.7 | 2.47 | 12.4 | 2.537 |
| **12** | Neurological Sciences | 1653 | 14742 | 8.9 | 1.075 | 5.1 | 0.765 |
| **13** | BMC Neurology | 1572 | 15202 | 9.7 | 1.092 | 4 | 0.771 |
| **14** | Annals of Indian Academy of Neurology | 1566 | 6584 | 4.2 | 0.654 | 2.3 | 0.334 |
| **15** | Clinical Neurology and Neurosurgery | 1544 | 12730 | 8.2 | 0.785 | 3.2 | 0.538 |
| **16** | Chinese Journal of Neurology | 1390 | 1032 | 0.7 | 0.214 | 0.7 | 0.137 |
| **17** | Journal of Neurosurgery | 1378 | 23644 | 17.2 | 1.896 | 8.1 | 1.138 |
| **18** | Clinical Neurology | 1337 | 2042 | 1.5 | 0.227 | 0.4 | 0.138 |
| **19** | Journal of Neurosciences in Rural Practice | 1179 | 5081 | 4.3 | 0.717 | 2.2 | 0.343 |
| **20** | Journal of Korean Neurosurgical Society | 1153 | 9285 | 8.1 | 1.099 | 3 | 0.522 |
| **21** | Brain and Development | 1131 | 10443 | 9.2 | 1.002 | 3.8 | 0.624 |
| **22** | Surgical Neurology International | 1131 | 3378 | 3 | 0.509 | 1.1 | 0.256 |
| **23** | Human Brain Mapping | 1112 | 29507 | 26.5 | 1.395 | 9.1 | 1.688 |
| **24** | Spine Journal | 1071 | 17375 | 16.2 | 2.087 | 7.5 | 1.562 |
| **25** | Acta Neurochirurgica | 1044 | 9472 | 9.1 | 1.302 | 4.2 | 0.718 |
| **26** | American Journal of Neuroradiology | 983 | 19006 | 19.3 | 1.553 | 6.7 | 1.167 |
| **27** | Epilepsy and Behavior | 977 | 10369 | 10.6 | 0.899 | 5.1 | 0.923 |
| **28** | Child's Nervous System | 941 | 5650 | 6 | 0.9 | 2.6 | 0.504 |
| **29** | Neurologia Medico-Chirurgica | 935 | 9835 | 10.5 | 1.15 | 3.5 | 0.691 |
| **30** | Clinical Neurophysiology | 923 | 19460 | 21.1 | 1.482 | 7.6 | 1.212 |
| **31** | Parkinsonism and Related Disorders | 901 | 15010 | 16.7 | 1.19 | 7.4 | 1.05 |
| **32** | Journal of Neurology | 890 | 14359 | 16.1 | 1.874 | 8.8 | 1.556 |
| **33** | Psychiatry and Clinical Neurosciences | 852 | 12842 | 15.1 | 1.754 | 9.7 | 1.389 |
| **34** | Neural Plasticity | 829 | 11404 | 13.8 | 0.836 | 5.7 | 0.766 |
| **35** | Journal of Neuro-Oncology | 825 | 14165 | 17.2 | 1.338 | 7.3 | 1.178 |
| **36** | Brain and Nerve | 817 | 1460 | 1.8 | - | - | 0.145 |
| **37** | Neurological Research | 799 | 9450 | 11.8 | 0.707 | 4 | 0.543 |
| **38** | Japanese Journal of Neurosurgery | 794 | 220 | 0.3 | 0.015 | 0 | 0.101 |
| **39** | Neurobiology of Aging | 793 | 22476 | 28.3 | 1.058 | 8.7 | 1.521 |
| **40** | Chinese Journal of Cerebrovascular Diseases | 792 | 246 | 0.3 | 0.024 | 0.3 | 0.114 |
| **41** | European Journal of Neurology | 788 | 11707 | 14.9 | 1.61 | 8.7 | 1.554 |
| **42** | Otology and Neurotology | 759 | 7957 | 10.5 | 1.482 | 3.5 | 0.887 |
| **43** | Sleep and Breathing | 744 | 7971 | 10.7 | 1.096 | 4.9 | 0.685 |
| **44** | British Journal of Neurosurgery | 712 | 3796 | 5.3 | 0.55 | 2.5 | 0.272 |
| **45** | Journal of Neurosurgery: Spine | 710 | 10503 | 14.8 | 1.597 | 5.4 | 1.192 |
| **46** | Journal of Clinical Neurology (Korea) | 699 | 6633 | 9.5 | 1.082 | 4.2 | 0.76 |
| **47** | NeuroImage: Clinical | 699 | 13769 | 19.7 | 1.324 | 8.1 | 1.395 |
| **48** | Journal of Neuroimmunology | 695 | 11319 | 16.3 | 0.804 | 6.2 | 0.919 |
| **49** | Brain Imaging and Behavior | 694 | 8621 | 12.4 | 1.07 | 6.2 | 1.002 |
| **50** | Movement Disorders | 680 | 25703 | 37.8 | 1.981 | 13.7 | 2.602 |
| **51** | Journal of Neurology, Neurosurgery and Psychiatry | 679 | 22270 | 32.8 | 2.878 | 15.9 | 3.178 |
| **52** | Metabolic Brain Disease | 679 | 9854 | 14.5 | 0.853 | 5.7 | 0.761 |
| **53** | Seizure : the journal of the British Epilepsy Association | 679 | 8579 | 12.6 | 1.126 | 5.3 | 0.868 |
| **54** | Neuroradiology | 663 | 7976 | 12 | 1.367 | 4.7 | 0.823 |
| **55** | Epilepsy Research | 662 | 9275 | 14 | 0.815 | 4.5 | 0.727 |
| **56** | Epilepsia | 649 | 28760 | 44.3 | 1.893 | 10.6 | 1.966 |
| **57** | Pain | 641 | 22829 | 35.6 | 3.151 | 12.5 | 2.445 |
| **58** | Interdisciplinary Neurosurgery: Advanced Techniques and Case Management | 641 | 659 | 1 | 0.302 | 0.7 | 0.177 |
| **59** | Neurology Asia | 620 | 898 | 1.4 | 0.112 | 0.3 | 0.14 |
| **60** | Muscle and Nerve | 611 | 7369 | 12.1 | 1.236 | 5.7 | 0.909 |
| **61** | Journal of Pain and Symptom Management | 606 | 10969 | 18.1 | 1.67 | 7.4 | 1.306 |
| **62** | Neurourology and Urodynamics | 595 | 8142 | 13.7 | 1.247 | 4.5 | 0.721 |
| **63** | Spinal Cord | 592 | 7619 | 12.9 | 1.359 | 3.9 | 0.716 |
| **64** | Sleep | 580 | 16798 | 29 | 1.591 | 8.7 | 1.58 |
| **65** | Journal of Neurogastroenterology and Motility | 578 | 7440 | 12.9 | 1.383 | 6.9 | 0.979 |
| **66** | Aging and Disease | 567 | 13781 | 24.3 | 1.494 | 13.6 | 1.682 |
| **67** | Neuropathology | 554 | 5644 | 10.2 | 0.628 | 3.5 | 0.666 |
| **68** | Neurosurgical Review | 546 | 5192 | 9.5 | 1.671 | 4.8 | 0.808 |
| **69** | Developmental Medicine and Child Neurology | 543 | 13037 | 24 | 1.998 | 8.2 | 1.146 |
| **70** | Journal of Cerebral Blood Flow and Metabolism | 543 | 17266 | 31.8 | 1.519 | 11.4 | 1.925 |
| **71** | Chinese Journal of Neuromedicine | 542 | 127 | 0.2 | 0.017 | 0.2 | 0.11 |
| **72** | Multiple Sclerosis and Related Disorders | 536 | 4635 | 8.6 | 0.997 | 5.6 | 0.92 |
| **73** | Chinese Journal of Urology | 522 | 93 | 0.2 | 0.043 | 0.1 | 0.112 |
| **74** | Acupuncture in Medicine | 519 | 4890 | 9.4 | 1.117 | 3.9 | 0.468 |
| **75** | Journal of NeuroInterventional Surgery | 519 | 6935 | 13.4 | 2.454 | 10.7 | 2.01 |
| **76** | Clinical and Experimental Neuroimmunology | 517 | 1256 | 2.4 | 0.212 | 1.1 | 0.195 |
| **77** | Brain | 507 | 26446 | 52.2 | 3.147 | 20.7 | 4.437 |
| **78** | Chinese Journal of Neurosurgery | 490 | 48 | 0.1 | 0.045 | 0.1 | 0.103 |
| **79** | Global Spine Journal | 485 | 4785 | 9.9 | 1.572 | 5 | 1.043 |
| **80** | Brain Injury | 479 | 5594 | 11.7 | 0.912 | 3.4 | 0.567 |
| **81** | Journal of Neurotrauma | 469 | 11066 | 23.6 | 1.288 | 9.4 | 1.31 |
| **82** | International Neurourology Journal | 465 | 4170 | 9 | 0.853 | 4.5 | 0.612 |
| **83** | Interventional Neuroradiology | 463 | 2814 | 6.1 | 0.931 | 2.8 | 0.478 |
| **84** | Turkish Neurosurgery | 461 | 2463 | 5.3 | 0.47 | 1.6 | 0.264 |
| **85** | Annals of Neurology | 453 | 18745 | 41.4 | 2.769 | 18.4 | 3.977 |
| **86** | Journal of Clinical Sleep Medicine | 452 | 9481 | 21 | 1.329 | 5.5 | 0.988 |
| **87** | Brain Stimulation | 447 | 12479 | 27.9 | 1.816 | 12.9 | 2.184 |
| **88** | Clinical Spine Surgery | 445 | 4364 | 9.8 | 0.94 | 2.9 | 0.682 |
| **89** | Neurosurgery | 436 | 8707 | 20 | 2.325 | 7.4 | 1.221 |
| **90** | Annals of Clinical and Translational Neurology | 427 | 6968 | 16.3 | 1.383 | 8.8 | 1.885 |
| **91** | European Neurology | 415 | 4254 | 10.3 | 0.981 | 3.8 | 0.492 |
| **92** | Equilibrium Research | 415 | 279 | 0.7 | 0.181 | 0.2 | 0.16 |
| **93** | Pediatric Neurology | 410 | 3485 | 8.5 | 1.152 | 5.2 | 0.876 |
| **94** | Pain Medicine | 410 | 5762 | 14.1 | 1.207 | 5.9 | 0.854 |
| **95** | Cerebrovascular Diseases | 399 | 5630 | 14.1 | 1.027 | 4.8 | 0.757 |
| **96** | Acta Neurologica Belgica | 398 | 1667 | 4.2 | 0.931 | 3.7 | 0.55 |
| **97** | Current Alzheimer Research | 398 | 6393 | 16.1 | 0.58 | 5.1 | 0.637 |
| **98** | JAMA Neurology | 398 | 20104 | 50.5 | 5.982 | 40.7 | 6.697 |
| **99** | Neuro-Ophthalmology Japan | 373 | 38 | 0.1 | 0.019 | 0 | 0.101 |
| **100** | Journal of Child Neurology | 367 | 3686 | 10 | 1.025 | 3.9 | 0.614 |

**Table 6:**	The list of 100 sources with most publications, total citations, citations per publication, Source-Normalized Impact per Paper (SNIP), CiteScore 2022 and SCImago Journal Rank (SJR).
